# Revised nomenclature of avian quadrate morphology and a detailed survey of clade-specific anatomical features

**DOI:** 10.64898/2026.02.07.704559

**Authors:** Pei-Chen Kuo, Roger Benson, Daniel J. Field

## Abstract

In birds, the quadrate bone serves as a hinge articulating with the lower jaw and the skull, playing an important mechanical role in the feeding apparatus. Avian cranial kinesis is dependent on the streptostylic quadrate transferring force from the adductor muscles at the back of the skull toward the beak, as part of a four-bar mechanical linkage to elevate and depress the bill. The complex morphology of the bird quadrate has led to authors adopting a range of alternative terminologies to describe the same anatomical structures and character states, impeding clarity of communication and presenting a barrier to progress in our understanding of the evolution of this important component of the avian feeding apparatus. Here, we reconcile terminological discord among previous studies on avian quadrate morphology and propose a stable nomenclature for future work. To characterise the considerable variation in quadrate form across crown bird diversity, we present an extensive anatomical atlas of the avian quadrate and summarise major patterns of quadrate morphological variation across extant avian phylogeny. In addition, we investigate macroevolutionary patterns in avian quadrate morphology, incorporating comparisons of crown birds and Late Cretaceous near-crown stem birds. We demonstrate that quadrate characters are useful for diagnosing a range of major avian subclades, and suggest that numerous distinctive features are likely to be associated with important biomechanical consequences. This investigation has implications for resolving the unsettled phylogenetic relationships of extinct bird clades such as Pelagornithidae and Gastornithiformes, as well as controversial relationships within several extant groups.

## Introduction

In birds, the quadrate bone (*os quadratum*, hereafter ‘quadrate’; Baumel and Witmer, 1993) serves an important role in cranial kinesis and the feeding apparatus as it directly articulates with the lower jaw (mandible), the skull (cranium), the quadratojugal, and the pterygoid-palatine complex (which comprises a pair of pterygoids and palatines, and a single vomer) (Figure 1 A-B; Bock, 1964; Burton, 1974; Bühler, 1981; Dawson et al., 2011; Bailleul et al., 2017). The quadrate transfers forces from the adductor muscles at the back of the skull through two set of pushrods (the pterygoid-palatine complex and the jugal bars; Figure 1B) to the mandible and rostrum as part of a four-bar mechanical linkage system which eventually bends the beak at a flexible position situated at its base and/or near its tip (Kaiser, 2010). The quadrate provides attachment sites for several key adductor muscles involved in cranial kinesis, such as *M. pseudotemporalis profundus*, *M. adductor mandibulae posterior*, and *M. protractor pars quadrati* (Bühler, 1981).

**Fig. 1.**
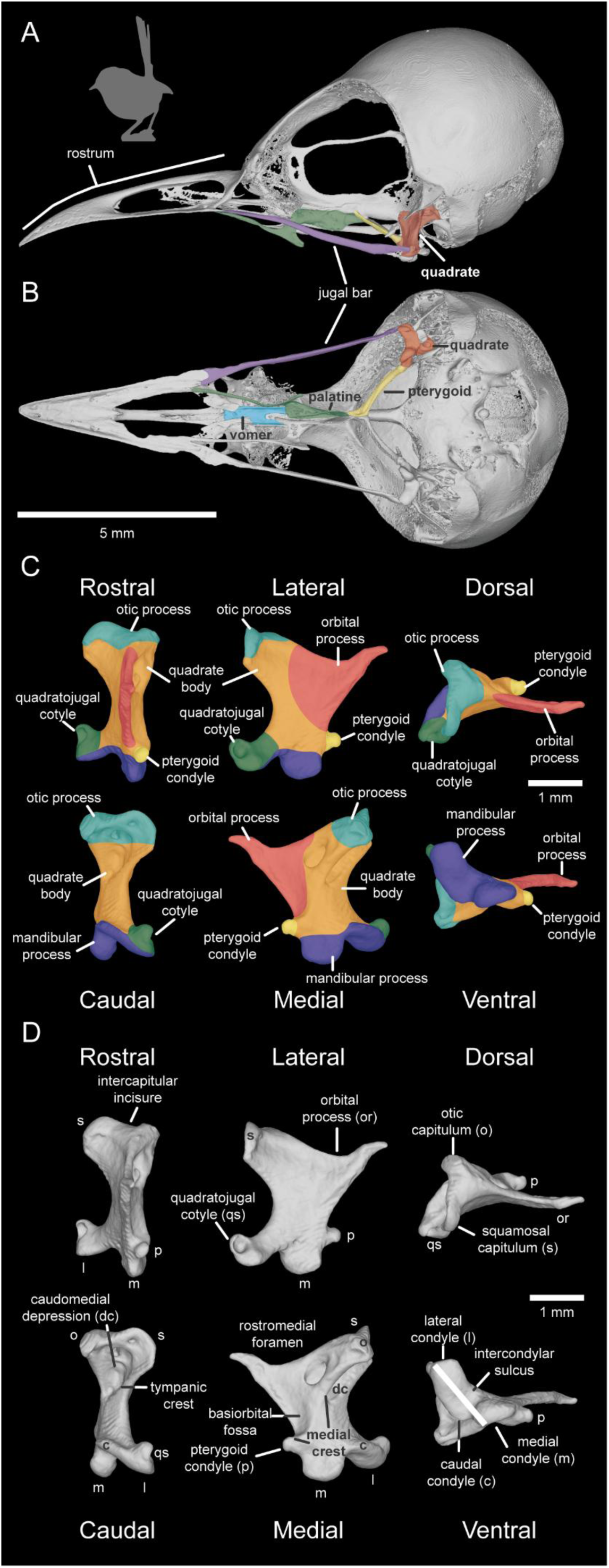
Quadrate anatomy in Neornithes. The skull of *Malurus melanocephalus* (UMMZ 224775), the species exhibiting a quadrate shape closest to the mean shape in our dataset in lateral (A) and ventral view (B), and its quadrate in rostral, lateral dorsal, caudal, medial, and ventral views (C and D). The quadrate of *Malurus melanocephalus* is highlighted in red in (A) and (B). The shape variance of the bird quadrate is separated into six different portions: otic process (*processus oticus*, coloured in light blue in C), orbital process (*processus orbitalis*, coloured in red in C), quadratojugal cotyle (*cotyla quadratojugalis*, coloured in green in C), pterygoid condyle (*condylus pterygoideus*, coloured in yellow in C), mandibular process (*processus mandibularis*, coloured in dark blue in C), and quadrate body (*corpus quadrati*, coloured in orange in C).

Studies on the morphological variation of bird quadrates extend back to the late 19th Century (Walker, 1888), and the morphological diversity of avian quadrates has at times been suggested to encode key information useful for bird classification (Lowe, 1926). Several studies have provided substantial qualitative descriptions of bird quadrate anatomy covering several major crown bird subclades (Galloanserae [Elzanowski and Stidham, 2010], Caprimulgidae [Bühler, 1970; Mayr, 2002; Zusi, 2013], Apodiformes [Mayr, 2002; Zusi, 2013], Cuculiformes [Posso and Donatelli, 2001], some clades in Ardeae [Saiff, 1978; Piro and Hospitaleche, 2019a, b; Hood et al., 2019], Accipitriformes [Saiff, 2006], some clades in Coraciimorphae [Elzanowski and Boles, 2015], or Phorusrhacidae [Degrange, 2021], and a wider diversity of crown birds [Walker, 1888; Samejima and Otsuka, 1987; Mayr, 2025]). Although Baumel and Witmer (1993) proposed systematic terminology for several aspects of avian quadrate morphology, the sheer diversity of form encompassed by avian quadrates has led to inconsistent nomenclatural approaches applied across prior studies, confounding large-scale investigations of this anatomically complex component of the feeding apparatus.

For instance, previous studies have labelled major anatomical components of the quadrate, such as the otic capitulum (*capitulum oticum*), squamosal capitulum (*capitulum squamosum*), quadratojugal cotyle (*cotyla quadratojugalis*), pterygoid condyle (*condylus pterygoideus*), mandibular process (*processus mandibularis*), medial mandibular condyle (*condylus mandibularis medialis*), lateral mandibular condyle (*condylus mandibularis lateralis*), and caudal mandibular condyle (*condylus mandibularis caudalis*) with their own idiosyncratic naming systems (Table 1), complicating subsequent studies and efforts to assess homologies and macroevolutionary changes among different avian lineages. As an example, several publications applied the term ‘lateral process’ to describe the laterally protruding quadratojugal cotyle articulating with the jugal bar (e.g., Elzanowski et al., 2001; Elzanowski and Stidham, 2010; Watanabe and Matsuoka, 2014; Mayr, 2015; Hood et al., 2019; Bell and Chiappe, 2020; Heingård et al., 2021). At the same time, these studies also used either quadratojugal cotyle (Hood et al., 2019; Heingård et al., 2021), quadratojugal socket (Elzanowski et al., 2001; Elzanowski and Stidham, 2010), or pars quadratojugalis (Elzanowski and Stidham, 2010; Watanabe and Matsuoka, 2014) to discuss the same character, multiplying the number of terminological options for a single anatomical feature. Furthermore, the nomenclatural system proposed by Baumel and Witmer (1993) lacks certain details necessary for detailed interclade comparisons of quadrate morpohology, such as pneumatic features. As such, revising and standardizing terminology for investigations of the avian quadrate is necessary for undertaking meaningful large-scale anatomical comparisons across bird diversity, and unambiguously scoring characters for phylogenetic analysis.

**Table 1.** Proposed terminology and abbreviations of the avian quadrate applied in this thesis, and alternative nomenclatures for equivalent structures from other publications. Standardized terminology and abbreviations for the avian quadrate adopted here are mainly based on Baumel and Witmer (1993) and Elzanowski and Stidham (2010). *We favour the term otic process (*processus oticus*) over quadrate head (*caput quadrati*) (Baumel and Witmer, 1993) for two reasons: first of all, ‘quadrate head’ does not precisely describe the articular connection with the braincase. Secondly, most previous studies have appled the term otic process rather than quadrate head, so otic process provides a more stable option (e.g., Norberg, 1977; Bühler et al., 1988; Elzanowski et al., 2001; Posso and Donatelli, 2001; Pascotto and Donatelli, 2003; Clarke, 2004; Saff, 2006; Ksepka et al., 2008; Mayr and Rubilar-Roger, 2010; Olson, 2011; Donatelli, 2012; Ksepka et al., 2012; Mayr and Zvonok, 2012; Zusi, 2013; Mayr, 2018; Mayr and Walsh, 2018; Hassan, 2019; Piro and Hospitaleche, 2019a-b; Mayr, 2020; Musser and Clarke, 2020; Degrange, 2021; Heingård et al., 2021; Mayr and Kitchener, 2022a-d; Mayr and Kitchener, 2023 a-b; Crane et al., 2025).

| Anatomical term applied here | Abb. | Alternative nomenclature in previous publications |
| --- | --- | --- |
| Quadrate (*Os quadratum*) | q |  |
| Quadrate body (*Corpus quadrati*) | qb | quadrate shaft, os quadratum shaft (Saiff, 1978; Saiff, 2006) |
| Caudomedial depression (*Depressio caudomedialis*) | dc | incisura intercapitularis (Piro and Hospitaleche, 2019a, b) |
| Subcapitular tubercle (*Tuberculum subcapitulare*) | t | eminentia articularis (Mayr and Rubilar-Roger, 2010), C1 crest (Donatelli, 2012), dorsal tubercle (Ksepka et al., 2008) |
| Lateral crest (*Crista lateralis*) | cl |  |
| Medial crest (*Crista medialis*) | cm | crista tympanica (Piro and Hospitaleche, 2019a, b) |
| Tympanic crest (*Crista tympanica*) | ct |  |
| Otic process* (processus oticus) | op | Caput quadrati (Baumel and Witmer, 1993), quadrate head (Walker, 1888; Saiff, 1978; Elzanowski et al., 2001; Nesbitt and Clarke, 2016), otic head (Saiff, 1988; Bell and Chiappe, 2020) |
| Squamosal capitulum (*Capitulum squamosum*) | s | squamosal head (Lowe, 1943; Saiff, 1982; Field et al., 2020); external capitulum (Walker, 1888; Saiff, 1974; Saiff, 1978; Saiff, 1988), squamosal articulation (Ono, 1989), squamosal facet (Elzanowski and Stidham, 2010; Hood et al., 2019); squamosal articular head (Saiff, 1976b); external otic process (Norberg, 1977); squamosal head (Bell and Chiappe, 2020) |
| Otic capitulum (*Capitulum oticum*) | o | internal capitulum (Walker, 1888; Saiff, 1974; Saiff, 1978; Saiff, 1988), prootic head (Field et al., 2020), medial facet (Ono, 1989), otic facet (Elzanowski and Stidham, 2010; Hood et al., 2019); prootic articular heads (Saiff, 1976); internal otic process (Norberg, 1977); otic head (Bell and Chiappe, 2020) |
| Intercapitular incisure<br>(*Incisura intercapitularis*) | ic | capitular groove (Saiff, 1976; Stiff, 1978; Saiff, 1982), intercapitular vallecule (Elzanowski and Stidham, 2010), intercapitular groove (Nesbitt et al, 2011) |
| Orbital process<br>(*Processus orbitalis*) | or | anterior process (Walker, 1888), postorbital process (Agnolin and Chafrat, 2015) |
| Orbital crest<br>(*Crista orbitalis*) | oc | orbitocotylar crest (Bertelli et al., 2006) |
| Basiorbital fossa<br>(*Fossa basiorbitalis*) | b | facies tympanica (Piro and Hospitaleche, 2019a, b); rostromedial depression (Pascotto and Donatelli, 2003) |
| Quadratojugal cotyle<br>(*Cotyla quadratojugal*) | qs | quadratojugal articulation (Clarke, 2004; Bertelli et al., 2006), quadratojugal socket (Elzanowski et al., 2001; Elzanowski and Stidham, 2010), lateral cotyle (Bühler et al., 1988), quadratojugal cup (Walker, 1888; Saiff, 1974; Saiff, 1978; Saiff, 1982), pars quadratojugal (Elzanowski and Stidham, 2010; Watanabe and Matsuoka, 2014), fovea quadratojugal (Tambussi et al., 2019); quadratojugal articulatory surface (Saiff, 1976; Saiff, 1978) |
| Submeatic process/<br>submeatic prominence<br>(*Processus submeaticus/*<br>*Prominentia submeatica*) | ps |  |
| Pterygoid condyle<br>(*Condylus pterygoideus*) | p | pterygoid articulation (Field et al., 2020; Ono, 1989; Elzanowski and Stidham, 2010; Hood et al., 2019), pterygoid facet (Elzanowski and Stidham, 2010; Elzanowski and Boles, 2015), pterygoquadrate articulation (Elzanowski and Stidham, 2010); pterygoid articulatory (Saiff, 1976) |
| Orbitopterygoid facet<br>(*Facet orbitopterygoidea*) | po | Pterygoid articulation facets of the orbital process (Elzanowski and Stidham, 2010); Pterygoid facet (Bertelli et al., 2006); pterygoid articular facet (Hood et al., 2019) |
| Mandibular process<br>(*Processus mandibularis*) | m | mandibular condyles (Saiff, 1981; Field et al., 2020; Elzanowski and Stidham, 2010), mandibular articulatory (Saiff, 1976; Saiff, 1978; Saiff, 1988; Saiff, 2006), ventral articulation (Ono, 1989), mandibular articulation (Elzanowski and Stidham, 2010; Hood et al., 2019), quadratomandibular articulation (Zusi, 2013) |
| Mandibular medial condyle<br>(*Condyles mandibularis medialis*) | mc | anterior condyle (Saiff, 1988), anterior condylar pairs (Saiff, 1978), inner condylar pairs (Saiff, 1976), inner condyle (Saiff, 1978) |
| Lateral trochlea<br>(*Trochlea lateralis*) | ltc | concave articulation facet lateral of the condylus medialis (Mayr, 2015); convex articular facet of medial condyle (Hood et al., 2019); trochlea of the medial condyle (Elzanowski and Zelenkov, 2015) |
| Mandibular lateral condyle<br>(*Condylus mandibularis lateralis*) | lc | posterior condyle (Saiff, 1981; Saiff, 1988), posterior condyle pairs (Saiff, 1976a; Saiff, 1978), outer condylar pairs (Saiff, 1976b), outer condyle (Saiff, 1978) |
| Mandibular caudal condyle<br>(*Condylus mandibularis caudalis*) | cc | posterior condyle (Saiff, 1978; Saiff, 1981; Saiff, 1988), posterior condyle pairs (Saiff, 1976a; Saiff, 1978), outer condylar pairs (Saiff, 1976b), outer condyle (Saiff, 1978), condylus rostralis (Saiff, 2006) |
| Intercondylar sulcus<br>(*Sulcus intercondylaris*) | is | trochlear groove (Saiff, 1974; Saiff, 1978); vallecule intercondylaris (Scofield et al., 2010) |
| Postsquamosal capitulum foramen<br>(*Foramen pneumaticum postcapitulum squamosum*) | fpps |  |
| Postotic capitulum foramen<br>(*Foramen pneumaticum postcapitulum oticum*) | fppo |  |
| Rostromedial foramen<br>(*Foramen pneumaticum rostromediale*) | fpr |  |
| Caudomedial foramen | fpc | foramen pneumaticum mediale (Scofield et al., |
| (*Foramen pneumaticum caudomediale*) |  | 2010; Elzanowski and Boles, 2015); caudal pneumatic foramen (Degrange et al., 2015); rostromedial pneumatic foramina (Mayr, 2015b; Hood et al., 2019) |
| Basiorbital foramen<br>(*Foramen pneumaticum basiorbitale*) | fpb | foramen pneumaticum mediale (Mayr, 2020) |
| Postcapitular foramen<br>(*Foramen pneumaticum postcapitulare*) | fppc |  |
| Dorsal foramen<br>(*Foramen pneumaticum dorsale*) | fpd |  |

Here, we investigated 250 bird quadrates across extant bird diversity as well as representatives of key extinct taxa (e.g., *Ichthyornis*, *Lithornis*, Pelagornithidae, *Palaelodus*, and Phorusrhacidae). We sample representatives of all major extant avian clades to build a comprehensive descriptive and comparative framework, in order to achieve three goals: Firstly, derived from the nomenclature systems previously proposed by Baumel and Witmer (1993), Elzanowski and Stidham (2010) and Mayr (2025), we propose a standardization of anatomical nomenclature for the avian quadrate (Table 1), clarifying the state of the literature for future anatomical descriptions. Secondly, following the classification of different quadrate shapes proposed by by Samejima and Otsuka (1987), we categorized different anatomical characters based on overall shape and the relative position of structures to facilitate comparisons among diverse avian lineages. Finally, we compared the morphological disparity of bird quadrates among crown and near-crown Mesozoic stem group birds to clarify the evolutionary history of avian quadrate morphology.

### Anatomical features of avian quadrates Otic process (*Processus oticus*)

The otic process of the bird quadrate connects with the skull via two capitula, the squamosal capitulum (*capitulum squamosum*) and otic capitulum (*capitulum oticum*) (Figure 1A, C, and D). The squamosal capitulum is positioned dorsolaterally on the otic process, while the otic capitulum is positioned dorsomedially. In most crown birds (Neornithes), the squamosal capitulum is either slightly elevated with respect to the otic capitulum or exhibits a comparable dorsal extent to the otic capitulum in rostral view (Figure 2B). The squamosal capitulum is significantly elevated with respect to the otic capitulum in only a few extant taxa, such as the skimmer *Rynchops niger*, the puffbird *Bucco capensis*, and Falconiformes (see Plate 26, 38, 40, and 41). However, the contrasting case, whereby the otic capitulum is substantially elevated with respect to the squamosal capitulum, is observed in the stem bird *Ichthyornis* and a handful of extant taxa (e.g., the pigeon *Columba livia*, the limpkin *Aramus guarauna*, the tropicbird *Phaethon lepturus*, the frigatebird *Fregata aquila*, and the pelican *Pelecanus occidentalis*; see Plate 1, 20, 22, 27, 30, and 32). In some representatives of Neoaves (e.g., the oilbird *Steatornis caripensis*, the barn owl *Tyto alba*, and some Coraciiformes), either the squamosal capitulum or the otic capitulum may be bifurcated at their articulation with the cranium, such that either the squamosal or otic capitulum are much wider lateromedially than is normally the case (Figure 2C). In some avian groups (mostly in Neognathae), a distinct cleavage or groove of variable depth, the intercapitular incisure (*incisura intercapitularis*), exists between two capitula on the otic process (Figure 2D). Based on the relative position of these two capitula and the direction of their articular surfaces, we have identified five categories of shape variation in the otic process:

**Fig. 2.**
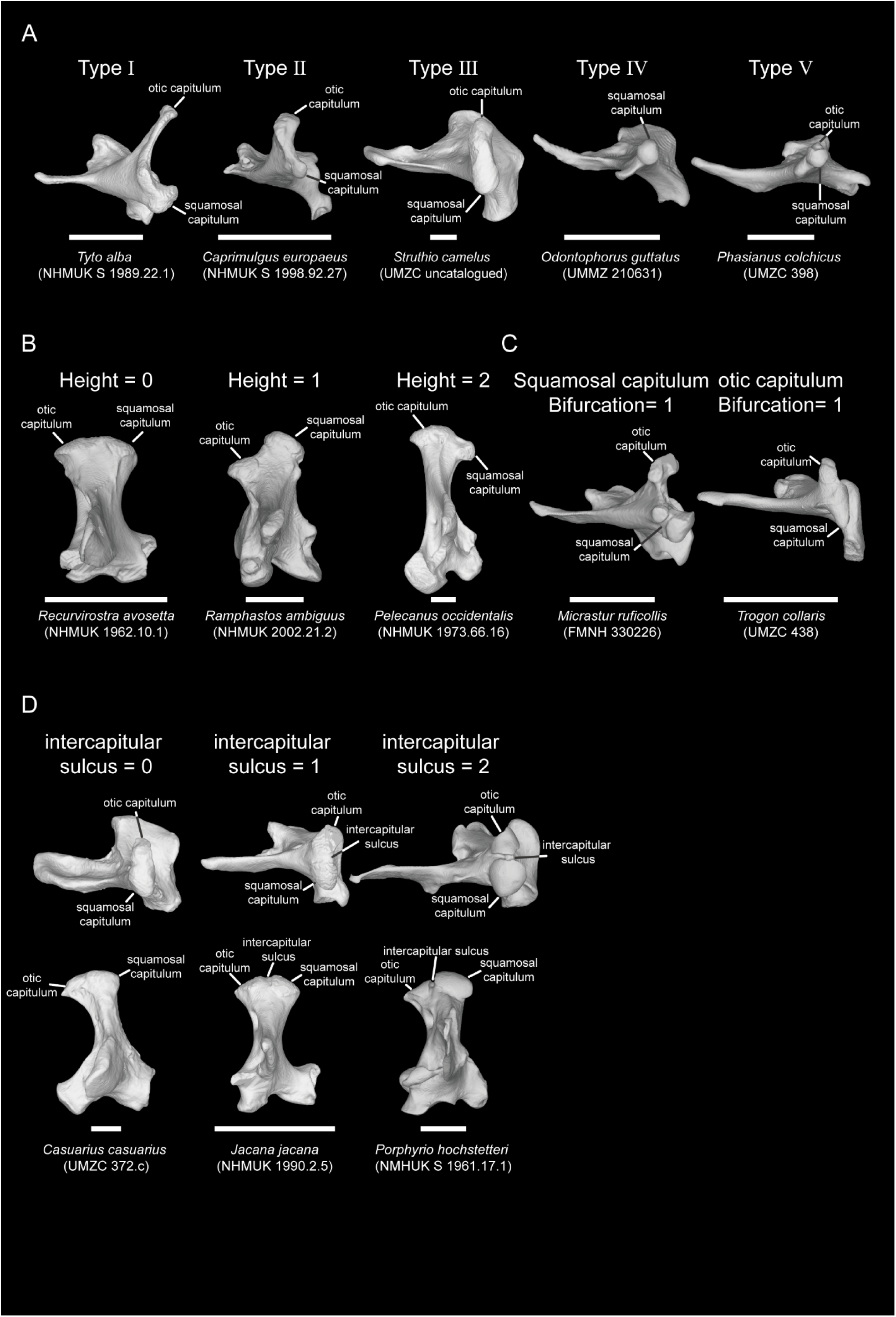
Classification for the types of the otic process (A) in dorsal view, the relative height between two capitula (B) in rostral view, the bifurcation of the articular surface on each capitulum (C) in dorsal view, and the absence/presence of the intercapitular sulcus on otic process (D) in dorsal (top) and rostral (bottom) view. Scale bar: 5 mm.

**Type** Ⅰ: The two capitula are widely separated with each other (Figure 2A). Extant taxa exhibiting this condition include the frogmouth *Podargus strigoides*, owls (Strigiformes), and two extinct representatives of ^†^Phorusrhacidae (^†^*Llallawavis scagliai* and ^†^*Psilopterus lemoinei*; Table 2 and Table S1). In owls, especially the barn owl (*Tyto alba*), the otic capitulum is greatly extended caudomedially into a structure that has been termed the internal otic process (see Plate 35; Norberg, 1977).

**Type** Ⅱ: The two capitula are situated closer to one another but are still distinctly separated (Figure 2A). This condition is seen in ^†^*Ichthyornis*, ^†^*Lithornis plebius*, kiwis (Apterygiformes), the tinamou *Nothoprocta ornata*, ^†^*Asteriornis maastrichtensis*, some Galliformes (Megapodiidae and Cracidae), ^†^*Conflicto antarcticus*, ^†^*Presbyornis*, some Anseriformes (Anhimidae, Anseranatidae, *Oxyura jamaicensis*, *Cereopsis novaehollandiae*, *Clangula hyemalis*, and *Merganetta armata*), most Strisores (except the frogmouth *Podargus strigoides* and the treeswift *Hemiprocne comata*), bustards and kin (Otidimorphae), pigeons and kin (Columbimorphae), most Gruiformes (with exceptions including the swamphen *Porphyrio hochstetteri*, the limpkin *Aramus guarauna*, and the crowned crane *Balearica pavonina*), most Mirandornithes (exceptions include the grebe *Podiceps taczanowskii*), most Charadriiformes (exceptions include the avocet *Recurvirostra avosetta*, the oystercatcher *Haematopus ostralegus*, and the woodcock *Scolopax rusticola*), Eurypygidae and Phaethontidae (collectively Phaethontiformes), loons (Gaviiformes), petrels and kin (Procellariiformes), some storks including *Ciconia ciconia*, boobies and kin (Suliformes), most Pelecaniformes (exceptions include the ibis *Eudocimus ruber*), diurnal raptors (Accipitriformes), mousebirds (Coliiformes), the courol *Leptosomus discolor*, Trogoniformes, hornbills and hoopoes (Bucerotiformes), two lineages of Coraciiformes (rollers such as *Coracias benghalensis* and motmots such as *Momotus momota*), most Piciformes (excepting wrynecks, *Jynx*), the seriema *Chunga burmeisteri* but not *Cariama cristata*, Falconiformes, New Zealand Wrens (Acanthisittidae), some, but not most representatives of Tyranni (including the asiti *Neodrepanis coruscans* and the manakin *Ceratopipra erythrocephala*), and most Passeri (except *Edolisoma tenuirostre*) (Table 2 and Table S1).

**Type** Ⅲ: The two capitula are situated close to each other, and in some groups, are almost continuous without a clear intercapitular incisure between them, rendering the boundaries of the two capitula unclear (Figure 2A). This condition characterises ^†^*Lithornis celetius*, most Palaeognathae (except kiwis Apterygiformes and the tinamou *Nothoprocta ornata*), the galliform *Ptilopachus petrosus*, the pelagornithids *Dasornis toliapica* and *Osteodontornis sp*., most waterfowl (Anatidae, except *Oxyura jamaicensis*, *Cereopsis novaehollandiae*, *Clangula hyemalis*, *Mergus merganser*, and *Merganetta armata*), *Hemiprocne comata*, select Gruiformes (e.g., *Porphyrio hochstetteri*, *Aramus guarauna*, and *Balearica pavonina*), the grebe *Podiceps taczanowskii*, some Charadriiformes (e.g., *Recurvirostra avosetta*, *Haematopus ostralegus*, and *Scolopax rusticola*), penguins (Sphenisciformes), some storks e.g., *Leptoptilos crumenifer*, the ibis *Eudocimus ruber*, the hoatzin *Opisthocomus hoazin*, most Coraciiformes (excepting rollers and motmots), wrynecks (e.g., *Jynx torquilla*), the seriema *Cariama cristata*, the parrot *Psittacus erithacus*, most Tyranni (exceptions include *Neodrepanis coruscans* and *Ceratopipra erythrocephala*) and the oscine *Edolisoma tenuirostre* (Table 2 and Table S1).

**Type** Ⅳ: the squamosal capitulum comprises the entire otic process (Figure 2A), a condition observed in a small number of galliform taxa, including *Odontophorus guttatus* and *Rollulus rouloul* (Table 2 and Table S1).

**Type** Ⅴ: the articular surface of the otic capitulum turns to face medially (Figure 2A). This condition characterises most Phasianidae (except the crested partridge *Rollulus rouloul*), some piscivorous anseriforms (e.g., *Mergus merganser*), and most parrots (Psittaciformes, except *Psittacus erithacus*; Table 2 and Table S1).

### Subcapitular tubercle (*Tuberculum subcapitulare*)

The subcapitular tubercle is a muscle attachment positioned laterally or rostrolaterally relative to the squamosal capitulum, which exhibits different shapes in different groups (Figure 8B; Elzanowski and Stidham, 2010). It has been proposed as a synapomorphy of Galloanserae (Galliformes + Anseriformes; Elzanowski and Stidham, 2010), but it is absent in all Pelagornithidae (extinct bony-tooth birds, which have been proposed to be members of Galloanserae). In most Galliformes, the subcapitular tubercle has a mound-like shape, while it forms a platform-like protrusion in most Anseriformes. Nevertheless, in some anseriform quadrates (e.g., *Anseranas semipalmata*, *Dendrocygna bicolor* and *Thalassornis leuconotus*), the subcapitular tubercle is positioned ventral to the squamosal capitulum (see Plate 11). In addition, the subcapitular tubercle is also present on the quadrates of some neoavians, such as the bustard *Ardeotis australis*, the mesite *Monias benschi*, some Gruiformes (rails such as *Lewinia striata* and *Atlantisia rogersi*, the trumpeter *Psophia crepitans*, the limpkin *Aramus guarauna*, the crowned crane *Balearica pavonina*, and the crane *Leucogeranus leucogeranus*), most Mirandornithes (except some grebes such as *Rollandia rolland* and *Podilymbus gigas*), some Charadriiformes (including the thick-knee *Burhinus senegalensis*, the plover *Charadrius vociferus*, *Recurvirostra avosetta*, *Haematopus ostralegus*, the snipe *Lymnocryptes minimus*, and the godwit *Limosa lapponica*), the loon *Gavia arctica*, Spheniscidae, some Procellariiformes (the petrels *Pelagodroma marina*, *Fulmarus glacialis*, *Puffinus puffinus*, *Pelecanoides urinatrix*, and *Macronectes giganteus*), storks (Ciconiiformes), most Suliformes (except *Fregata aquila*), four Pelecaniformes (*Eudocimus ruber*, and the herons *Tigrisoma lineatum*, *Ardea alba*, and *Ixobrychus minutus*), *Opisthocomus hoazin*, most Accipitriformes (except the osprey *Pandion haliaetus* and the secretarybird *Sagittarius serpentarius*), Coliiformes, *Leptosomus discolor*, most Coraciiformes (except the tody *Todus mexicanus*), three Piciformes (the woodpecker *Picus viridis*, the barbet *Psilopogon chrysopogon*, and the toucan *Ramphastos ambiguus*), the falcon *Micrastur ruficollis*, and the manakin *Ceratopipra erythrocephala*.

It is worth mentioning that the subcapitular tubercle is positioned far ventral to the squamosal capitulum with a prominent linear ridge in Coliiformes, *Picus viridis* and other Pici (Donatelli, 2012). This might be related to a distinctive biomechanical function, as an important adductor muscle, *M. adductor mandibulae externus caudalis lateralis*, originates on this ridge (Donatelli, 2012). In most Passeri (with uncommon exceptions including the shriketit *Falcunculus frontatus*, the chough *Pyrrhocorax pyrrhocorax*, the kokako *Callaeas cinereus*, the rockfowl *Picathartes gymnocephalus*, and the laughingthrush *Leiothrix lutea*), a distinct ridge is located ventral to the squamosal capitulum, but shows little morphological affinity with the linear shape of the subcapitular tubercle in other bird groups. Instead, this ridge appears to extend from the lateral crest (see the lateral view of Passeri in Plate 43-46). Though this character in Passeri might not be homologous with the subcapitular tubercle in other avian groups, it potentially serves a comparable role as a muscle attachment for the *M. adductor mandibulae externus caudalis lateralis*, indicating that it may serve the same function as the subcapitular tubercle does in other groups. Future work investigating soft tissue relationships may clarify the nature of this structure.

### Orbital process (Processus orbitalis)

The orbital process is a distinct projection toward the orbit originating on the quadrate body (Figure 1C-D), and serves as the origin and lever arm of some adductor muscles of the feeding apparatus, such as the *M. pseudotemporalis profundus* (Baumel and Witmer, 1993). Based on its aspect ratio, the shape of its tip and the direction it faces, we have identified six main categories of shape variation for the orbital process:

**Type**: the orbital process shows a high aspect ratio with a flat and hammer-shaped tip (Figure 3A). This condition characterises ^†^*Ichthyornis*, tinamous (Tinamiformes), the coucal *Centropus goliath*, and the extinct penguin ^†^*Madrynornis mirandus* (Table 2 and Table S1).

**Fig. 3.**
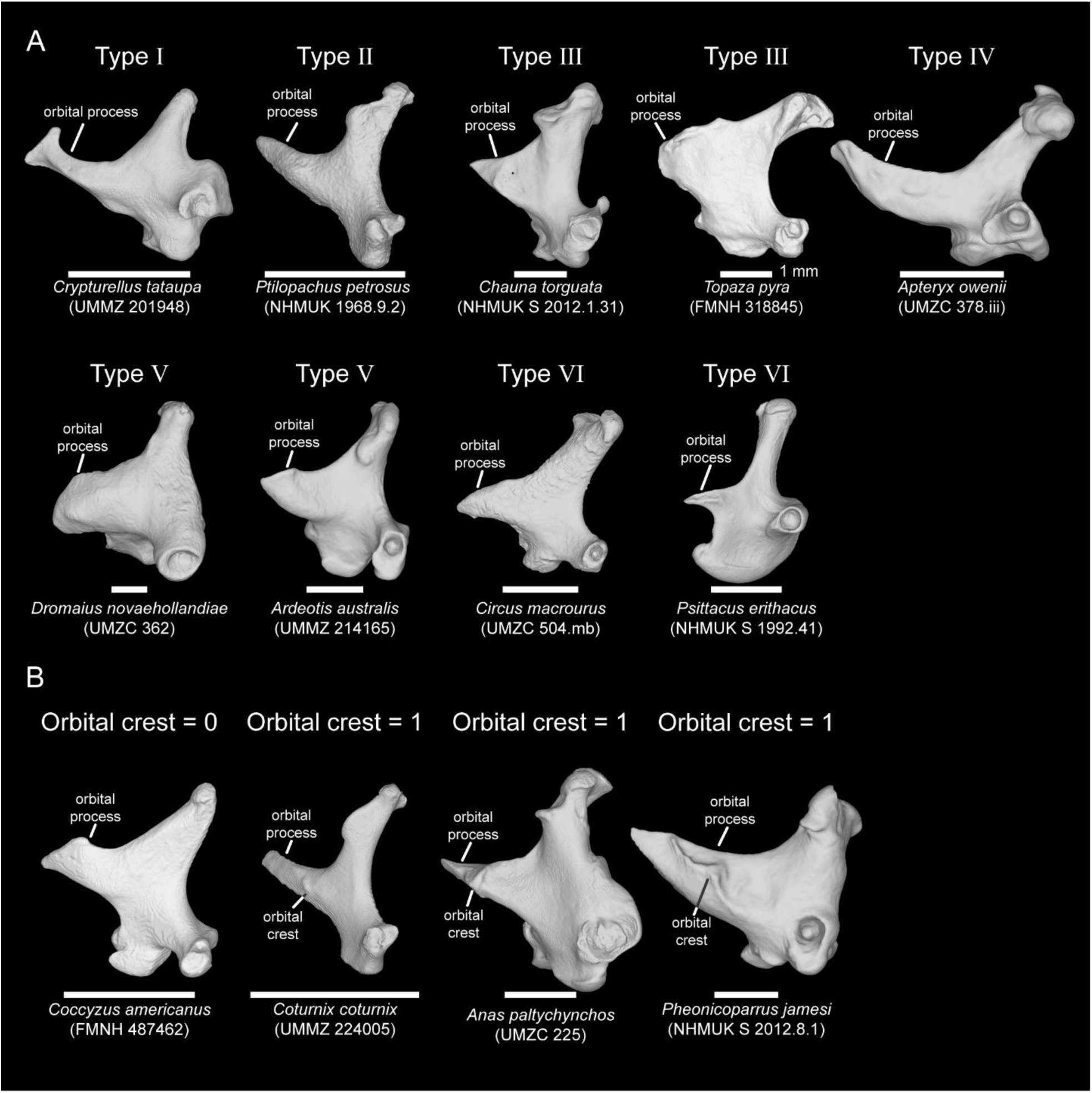
Classification for the types of the orbital process (A) and the absence/presence of the orbital crest (B) in lateral view. Scale bar: 5 mm. (Except for *Topaza pyra*.)

**Type**: the orbital process exhibits a high aspect ratio with a pointed tip (Figure 3A). This includes most Galliformes (except the curassow *Mitu mitu*, the guan *Pipile pipile*, and the pheasant *Lophophorus impejanus*), most Anatidae (exceptions include *Dendrocygna bicolor*, *Thalassornis leuconotus*, *Biziura lobata*, *Cereopsis novaehollandiae*, *Anser fabalis*, *Histrionicus histrionicus*, and *Plectropterus gambensis*), two Strisores (*Podargus strigoides*, and the owlet-nightjar *Aegotheles cristatus*), most Columbidae (except the dodo ^†^*Raphus cucullatus*), three Charadriiformes (the jacana *Jacana jacana*, *Scolopax rusticola*, and the buttonquail *Turnix varius*), Gaviiformes, extant Sphenisciformes, most Suliformes (except *Fregata aquila*), *Opisthocomus hoazin*, Strigidae, *Momotus momota*, *Jynx torquilla*, ^†^*Andalgalornis steulleti*, ^†^*Psilopterus lemoinei*, the parrot *Nestor notabilis*, some Tyranni including the crescentchest *Melanopareia torquata*, and a few representatives of Passeri (including the fairywren *Malurus melanocephalus*, *Callaeas cinereus*, *Chlorodrepanis virens*, and the bunting *Emberiza calandra*; Table 2 and Table S1).

**Type**: the orbital process displays a low aspect ratio with a pointed or a sharp tip (Figure 3A). This condition describes ^†^*Osteodontornis* sp., Anhimidae, *Anseranas semipalmata*, several Anatidae (*Dendrocygna bicolor*, *Thalassornis leuconotus*, *Biziura lobata*, *Cereopsis novaehollandiae*, *Anser fabalis*, and *Plectropterus gambensis*), most Strisores (exceptions include *Steatornis caripensis*, *Podargus strigoides*, and *Aegotheles cristatus*), *Rynchops niger*, *Pelecanoides urinatrix*, some Coraciiformes (including the bee-eater *Merops orientalis*, *Todus mexicanus*, and the kingfisher *Alcedo atthis*), *Bucco capensis*, *Micrastur ruficollis*, and Acanthisittidae (Table 2 and Table S1).

**Type**: the orbital process appears to have a high aspect ratio with a blunt and flat tip (Figure 3A). This condition characerises Apterygiformes, some Galliformes (*Mitu mitu*, *Pipile pipile*, and *Lophophorus impejanus*), the diving duck *Histrionicus histrionicus*, *Steatornis caripensis*, most Otidimorphae (except *Ardeotis australis* and *Centropus goliath*), some Columbimorphae (*Monias benschi*, the sandgrouse *Syrrhaptes paradoxus*, and ^†^*Raphus cucullatus*), most Gruiformes (except the flufftail *Sarothrura elegans*), Mirandornithes, most Charadriiformes (*Recurvirostra avosetta*, *Haematopus ostralegus*, *Lymnocryptes minimus*, *Limosa lapponica*, the whimbrel *Numenius phaeopus*, the skua *Stercorarius skua*, the pratincole *Glareola pratincola*, the auk *Uria aalge*, and the gull *Chroicocephalus novaehollandiae*), Eurypygiformes, most Procellariiformes (except *Pelecanoides urinatrix*), Ciconiiformes, *Fregata aquila*, most Pelecaniformes (except the shoebill *Balaeniceps rex* and *Pelecanus occidentalis*), some Accipitriformes (including the New World vulture *Cathartes burrovianus*, *Pandion haliaetus*, and the Old World vulture *Necrosyrtes monachus*), Tytonidae, Coliiformes, *Leptosomus discolor*, Trogoniformes, Bucerotiformes, Coraciiformes belonging to the roller and ground-roller clade, Coracii (represented by *Coracias benghalensis* and the ground-roller *Atelornis pittoides* in our dataset), most Piciformes (except the jacamar *Galbula dea*, *Bucco capensis*, and *Jynx torquilla*), ^†^*Llallawavis scagliai*, Cariamidae, Falconiformes belonging to the caracara clade Polyborinae (represented by *Caracara cheriway* and *Daptrius ater* in our dataset), two Psittaciformes (the kakapo *Strigops habroptilus* and the cockatoo *Probosciger aterrimus*), most Tyranni (except *Melanopareia torquata* in our dataset), and most Passeri (except *Malurus melanocephalus*, *Callaeas cinereus*, *Chlorodrepanis virens*, and *Emberiza calandra*) (Table 2 and Table S1).

**Type**: the orbital process has a low aspect ratio with a blunt or flat tip (Figure 3A). This type includes ^†^Lithornithidae, most extant Palaeognathae (except Apterygiformes and Tinamiformes), the moa ^†^*Megalapteryx didinus*, *Ardeotis australis*, *Sarothrura elegans*, three Charadriiformes (*Burhinus senegalensis*, *Charadrius vociferus*, and the plains-wanderer *Pedionomus torquatus*), *Phaethon lepturus*, two Pelecaniformes (*Balaeniceps rex* and *Pelecanus occidentalis*), *Galbula dea*, and the falcon *Falco sparverius* (Table 2 and Table S1).

**Type**: Unusually, the tip of the orbital process faces ventrally (Figure 3A). This condition is only observed in most Accipitriformes (except *Cathartes burrovianus*, *Pandion haliaetus* and *Necrosyrtes monachus*) and *Psittacus erithacus* (Table 2 and Table S1).

### Orbital crest (*Crista orbitalis*)

The orbital crest is a small structure on the lateral surface of the orbital process (Figure 3B) and is considered one of the synapomorphic features of galloanseran quadrates (Elzanowski and Stidham, 2010). In Galliformes, the orbital crest is a linear ridge positioned close to the ventral surface of the orbital process, while it is a tilted U- or V-shaped structure with two branches extending dorsally and ventrally on anseriform quadrates (Elzanowski and Stidham, 2010). The orbital crest is also observed in some Tinamidae (including *Nothoprocta ornata* and *Crypturellus tataupa*), *Porphyrio hochstetteri*, some Mirandornithes (the flamingos *Phoenicopterus roseus* and *Phoenicoparrus jamesi*, and the grebe *Podilymbus gigas*), *Recurvirostra avosetta*, *Haematopus ostralegus*, Gaviidae, Spheniscidae, some Procellariiformes (the shearwater *Ardenna tenuirostris*, and the petrels *Puffinus puffinus*, *Pterodroma lessonii*, and *Halobaena caerulea*), *Cariama cristata*, *Psittacus erithacus*, and two Passeri (*Callaeas cinereus* and *Chlorodrepanis virens*). It remains in question whether the orbitocotylar crest on ^†^*Potamornis skutchi* (UCMP 73103, Hesperornithes) (Elzanowski et al., 2000), a prominent ridge extending from the rim of the quadratojugal cotyle to the base of the orbital process, is homologous with the orbital crest in Neornithes.

### Basiorbital fossa (*Fossa basiorbitalis*)

The basiorbital fossa is a shallow depression positioned medially at the base of the orbital process and dorsally above the pterygoid condyle (Figure 1D; Elzanowski and Stidham, 2010). It varies in size and depth among taxa examined. For instance, it is broad and distinctively deep in most Anseriformes, Gruiformes, and Mirandornithes, while it is shallow in most major subclades of Neoaves.

### Quadratojugal cotyle (*Cotyla quadratojugalis*)

The quadratojugal cotyle is an articular joint connecting the jugal bar to the quadrate, an important connection in the avian cranial kinetic system (Figure 1). Generally, it forms a cup- or socket-like fossa, but in some Galliformes (e.g., *Gallus varius*) and Tinamiformes, it has a saddle-like articular surface without clear dorsal or ventral margins. In some Strisores (e.g., *Hemiprocne comata*) and Coraciimorphae (e.g., *Trogon melanurus*), the margin of the quadratojugal cotyle does not develop distinctly, instead forming a flat articular surface. Based on how the jugal connects with the quadrate and the shape of its articular surface, the quadratojugal cotyle can be categorised into four different types:

**Type**: the quadratojugal cotyle exhibits a shallow joint with a complete margin (Figure 4A). This condition applies to ^†^*Ichthyornis*, the quail *Odontophorus guttatus*, five examind Anseriformes (^†^*Conflicto antarcticus*, *Anseranas semipalmata*, *Dendrocygna bicolor*, *Biziura lobata*, *Tachyeres brachyptera*), four examined Strisores (the nightjars *Eurostopodus mysticalis*, *Caprimulgus europaeus*, and *Chordeiles minor*, and *Steatornis caripensis*), two observed Otidimorphae (the turaco *Corythaeola cristata* and *Ardeotis australis*), ^†^*Raphus cucullatus*, *Opisthocomus hoazin*, *Sagittarius serpentarius*, the mousebird *Colius striatus*, the hoopoe *Upupa epops*, ^†^*Andalgalornis steulleti*, Acanthisittidae, and two Passeri (the swallow *Hirundo rustica* and the weaver *Plocepasser mahali*) (Table 2 and Table S1).

**Fig. 4.**
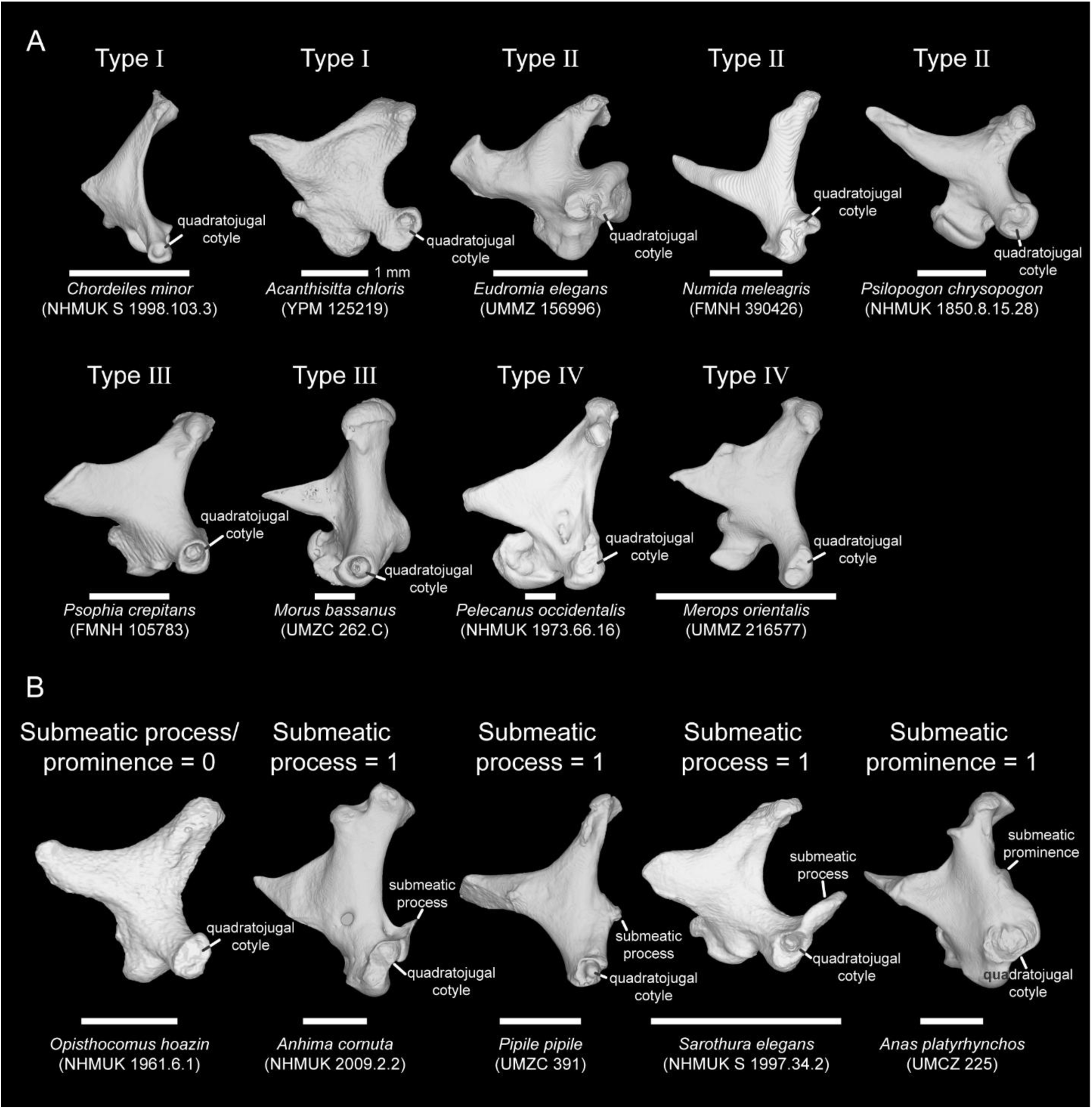
Classification for the types of the quadratojugal cotyle (A) and the absence/presence of the submeatic process/prominence (B) in lateral view. Scale bar: 5 mm.

**Type:** the dorsal or ventral margin of the quadratojugal cotyle is not well-developed (Figure 4A). This condition incorporates the kiwi *Apteryx australis*, Tinamiformes, most Galliformes (except Megapodiidae, *Mitu mitu*, and *Odontophorus guttatus*), most Anseriformes (except ^†^*Conflicto*, ^†^*Presbyornis*, *Anseranas semipalmata*, *Dendrocygna bicolor*, *Biziura lobata*, and *Tachyeres brachyptera*), most Strisores (the potoo *Nyctibius griseus*, *Aegotheles cristatus*, and hummingbirds Trochilidae), most Columbidae (except ^†^*Raphus cucullatus*), *Porphyrio hochstetteri*, *Pelagodroma marina*, the kite *Elanus caeruleus*, most Strigiformes (except the bay owl *Phodilus badius*), two Coraciiformes (*Todus mexicanus* and *Alcedo atthis*), two Piciformes (the honeyguide *Indicator exilis* and the barbet *Psilopogon chrysopogon*), and Tyranni (Table 2 and Table S1).

**Type**: the quadratojugal cotyle exhibits a relatively deep joint with a complete margin (Figure 4A). This condition is observed in most Palaeognathae (Lithornithiformes, *Struthio camelus*, *Rhea americana*, *Apteryx owenii*, Casuariiformes, and ^†^*Megalapteryx didinus*), ^†^*Asteriornis maastrichtensis*, Pelagornithidae, a few Galliformes (Megapodiidae and *Mitu mitu*), ^†^*Presbyornis*, *Podargus strigoides*, three Otidimorphae (the cuckoos *Crotophaga ani*, *Tapera naevia*, and *Centropus goliath*), two Columbimorphae (*Monias benschi* and *Syrrhaptes paradoxus*), most Gruiformes (except for *Porphyrio hochstetteri*), Mirandornithes, Charadriiformes, Eurypygiformes, *Phaethon lepturus*, Gaviiformes, Sphenisciformes, most Procellariiformes (except *Pelagodroma marina*), Ciconiiformes, Suliformes, most Pelecaniformes (except *Pelecanus occidentalis*), most Accipitriformes (except *Sagittarius serpentarius* and *Elanus caeruleus*), *Leptosomus discolor*, most Bucerotiformes (except *Upupa epops*), three Coraciiformes (*Coracias benghalensis*, *Atelornis pittoides*, and *Momotus momota*), four examined Piciformes (*Galbula dea*, *Bucco capensis*, *Picus viridis*, and *Ramphastos ambiguus*), ^†^*Llallawavis scagliai*, ^†^*Psilopterus lemoinei*, extent Cariamiformes, Falconiformes, Psittaciformes, and most observed Passeri (except *Falcunculus frontatus*, *Hirundo rustica*, and *Plocepasser mahali*) (Table 2 and Table S1).

**Type:** the quadratojugal cotyle has a flat articular surface, and may not exhibit clear margins (Figure 4A). This is observed in three Strisores (*Hemiprocne comata*, and the swifts Apodidae), two Otidimorphae (the cuckoos *Coccyzus americanus* and *Hierococcyx fugax*), *Pelecanus occidentalis*, *Phodilus badius*, the mousebird *Urocolius macrourus*, Trogoniformes, *Merops orientalis*, two Piciformes (*Jynx torquilla* and the barbet *Lybius dubius*), and *Falcunculus frontatus* (Table 2 and Table S1).

### Submeatic process/submeatic prominence (Processus submeaticus/Prominentia submeatica)

The submeatic process or submeatic prominence is a bulge-like muscle attachment located at the caudal or dorsal margin of the quadratojugal cotyle (Figure 4B). This character appears in Cracidae, most Anseriformes (except *Anseranas semipalmata* and *Biziura lobata*), *Sarothrura elegans*, *Picus viridis*, and Falconiformes, but it differs in shape and position on those quadrates in specific clades. For instance, in Cracidae the submeatic process is positioned caudodorsal of the quadratojugal cotyle, forming a discrete protrusion with a round or square shape (see the lateral view of Cracidae quadrate in Plate 6-7), while it is ridge-like in most Anseriformes (see the lateral view of anseriform quadrate in Plate 10-14).

### Pterygoid condyle (Condylus pterygoideus)

The pterygoid condyle is located medially, close to the basiorbital fossa and dorsal to the medial condyle, with an oval articular surface connecting rostrally with the pterygoid (Figure 1C-D). Based on how the pterygoid articulates with the quadrate and the shape of this articular surface, we have discretised the shape variance of the pterygoid condyle into four types.

**Type**: the pterygoid condyle is bulbous and prominently directed rostrally (Figure 5A). This condition applies to ^†^*Ichthyornis*, ^†^*Dasornis toliapica*, some Anseriformes (*Chauna*, *Oxyura jamaicensis*, *Malacorhynchus membranaceus*, *Cygnus olor*, *Branta bernicla*, *Anser fabalis*, *Melanitta nigra*, *Mergellus albellus*, and *Tachyeres brachyptera*), most Apodiformes (except *Hemiprocne comata*, *Streptoprocne zonaris*, and *Phaethornis superciliosus*), some Cuculidae (*Corythaeola cristata*, *Crotophaga ani*, *Tapera naevia*, and *Coccyzus americanus*), the rail *Tribonyx mortierii*, most Mirandornithes (except ^†^*Palaelodus ambiguus* and *Phoenicoparrus jamesi*), four examined Charadriiformes (*Burhinus senegalensis*, *Pedionomus torquatus*, *Stercorarius skua*, and *Uria*), *Eurypyga helias*, *Phaethon lepturus*, Sphenisciformes, most Procellariiformes (except *Pelecanoides urinatrix*), *Ciconia ciconia*, two Sulliformes (the cormorants *Leucocarbo atriceps* and *Phalacrocorax carbo*), *Eudocimus ruber*, *Leptosomus discolor*, Trogoniformes, most Coraciiformes (except *Merops orientalis* and *Momotus momota*), most Piciformes (except *Indicator exilis*, *Jynx torquilla*, and *Picus viridis*), ^†^*Llallawavis scagliai*, two Falconiformes (*Micrastur ruficollis* and *Falco sparverius*), Psittaciformes, and Passeriformes (Table 2 and Table S1).

**Fig. 5.**
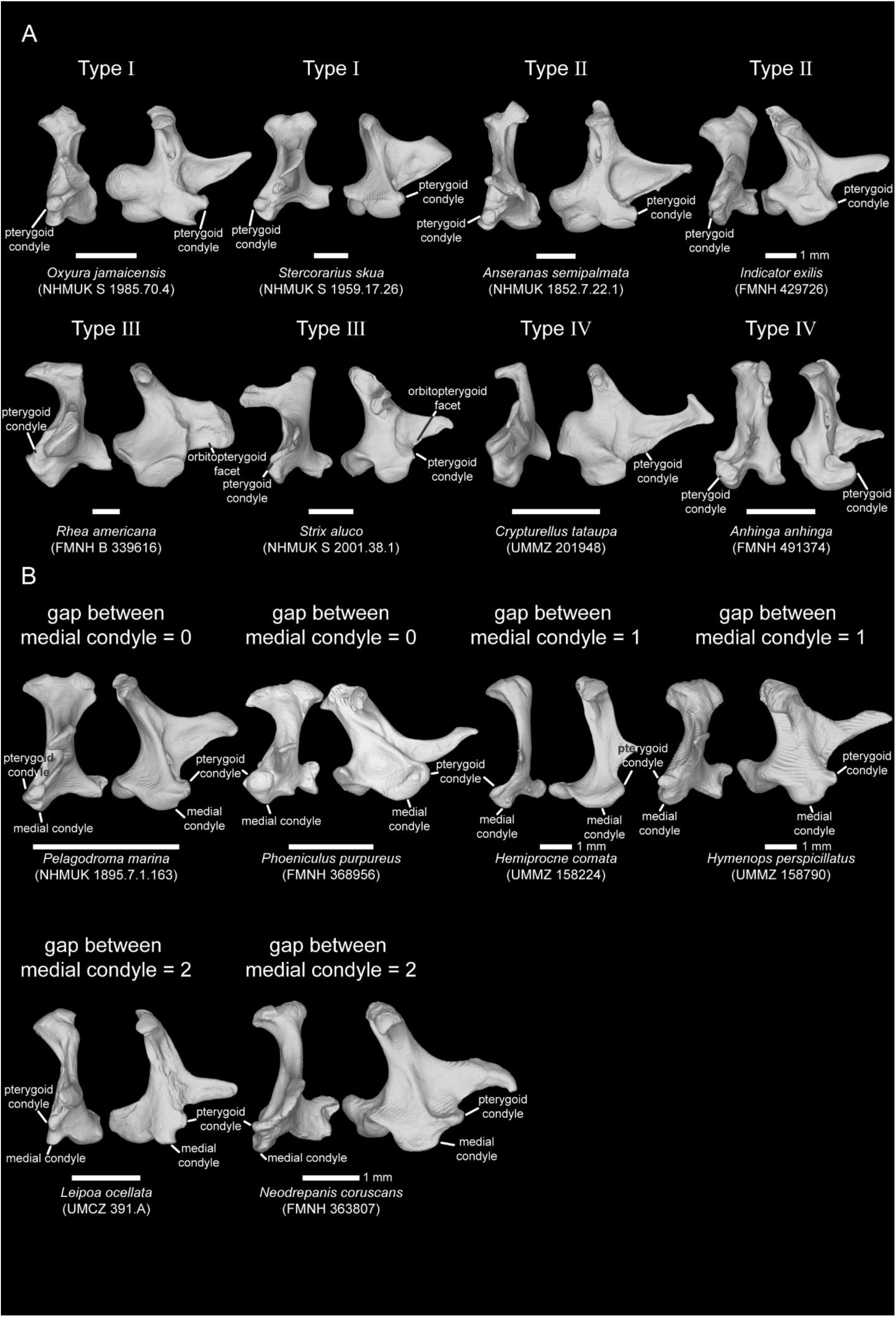
Classification for the types of the pterygoid condyle (A) and the gap between the pterygoid condyle and medial condyle (B) in rostral (right) and medial (left) view. Scale bar: 5 mm. (Except for *Indicator exilis*, *Hymenops perspicillatus*, and *Neodrepanis coruscans*.)

**Type**: the pterygoid condyle is bulbous or ovoid in shape, but only projects shallowly, therefore falling between between Type (prominently rostrally protruding) and Type (a flat articular surface; full description below) (Figure 5A). This condition applies to Apterygiformes, most Anseriformes (*Anseranas semipalmata*, *Dendrocygna bicolor*, *Thalassornis leuconotus*, *Cereopsis novaehollandiae*, *Clangula hyemalis*, *Histrionicus histrionicus*, *Mergus merganser*, *Chenonetta jubata*, *Plectropterus gambensis*, *Sarkidiornis melanotos*, *Aythya ferina*, *Anas platyrhynchos*, and *Anas aucklandica*), Caprimulgiformes, a few Apodiformes (*Hemiprocne comata*, *Streptoprocne zonaris*, and *Phaethornis superciliosus*), a few Otidimorphae (*Ardeotis australis*, *Centropus goliath*, and *Hierococcyx fugax*), some Columbimorphae (*Monias benschi*, *Syrrhaptes paradoxus*, *Didunculus strigirostris*, ^†^*Raphus cucullatus*, and *Leptotila rufaxilla*), two Gruiformes (*Porphyrio hochstetteri* and *Balearica pavonina*), *Phoenicoparrus jamesi*, *Haematopus ostralegus*, *Pelecanoides urinatrix*, two Suliformes (*Sula dactylatra* and *Morus bassanus*), three Pelecaniformes (*Tigrisoma lineatum*, *Ardea alba*, and *Scopus umbretta*), *Opisthocomus hoazin*, some Accipitrimorphae (*Elanus caeruleus*, *Buteo rufofuscus*, *Necrosyrtes monachus*, *Accipiter nisus*, and *Circus macrourus*), Coliiformes, Bucerotiformes, two Coraciiformes (*Merops orientalis* and *Momotus momota*), three Piciformes (*Indicator exilis*, *Jynx torquilla*, and *Picus viridis*), ^†^*Andalgalornis steulleti*, ^†^*Psilopterus lemoinei*, and extent Cariamiformes (Table 2 and Table S1).

**Type:** The quadrate exhibits two articular surfaces for the pterygoid: the pterygoid condyle and the orbitopterygoid facet, which is located medially at the ventral margin of the orbital process (Figure 5A). This condition applies to three members of Palaeognathae (*Struthio camelus*, *Rhea americana*, and ^†^*Megalapteryx didinus*), Galliformes, ^†^*Presbyornis*, most examined Columbidae (except ^†^*Raphus cucullatus* and *Leptotila rufaxilla*), most examined Gruiformes (except *Porphyrio hochstetteri*, *Tribonyx mortierii*, and *Balearica pavonina*), ^†^*Palaelodus ambiguus*, most examined Charadriiformes (except *Burhinus senegalensis*, *Haematopus ostralegus*, *Pedionomus torquatus*, *Stercorarius skua*, and *Uria aalge*), *Rhynochetos jubatus*, Gaviiformes, *Leptoptilos crumenifer*, *Fregata aquila*, two examined Pelecaniformes (*Ixobrychus minutus* and *Balaeniceps rex*) some Accipitrimorphae (e.g., *Cathartes burrovianus*, *Pandion haliaetus*, *Sagittarius serpentarius*, and *Parabuteo unicinctus*), Strigiformes, and the examined polyborine Falconiformes (*Caracara cheriway* and *Daptrius ater*) (Table 2 and Table S1). It is worth noting that the pterygoid condyle in the moa ^†^*Megalapteryx didinus* shares distinctive features with Casuariiformes and Tinamiformes (see the rostral and medial view of *Megalapteryx* quadrate in Plate 3): the pterygoid articular surface on the quadrate has two flat facies; one is located along the orbital process and faces medially (Tinamiformes-like feature), while the other is located above the medial condyle with a smooth surface (Casuariiformes-like feature). This observation is potentially relevant for assessing the controversial phylogenetic affinities of these palaeognath lineages (Faux and Field, 2017; Widrig and Field, 2022).

**Type:** the pterygoid condyle exhibits a flat and smooth articular surface with the pterygoid (Figure 5A). This condition applies to ^†^Lithornithiformes, Casuariiformes, Tinamiformes, ^†^*Asteriornis maastrichtensis* (in which the extent of the orbitopterygoid facet remains in question due to an incomplete orbital process; see the medial view of the *Asteriornis* quadrate in Plate 5), ^†^*Osteodontornis* sp., four Anseriformes (^†^*Conflicto antarcticus*, *Anhima cornuta*, *Biziura lobata*, and *Merganetta armata*), *Anhinga anhinga*, and *Pelecanus occidentalis* (Table 2 and Table S1). In Tinamiformes, the pterygoid condyle faces medially instead of rostrally with a flat articular surface, differing from that of other avian quadrates (see the medial view of the tinamou quadrate in Plate 4).

### Mandibular process (Processus mandibularis)

The mandibular process of the avian quadrate is the articular surface contacting the mandible (Figure 1C). In some avian quadrates, a clear groove, the intercondylar sulcus (*sulcus intercondylaris*), is located at the centre of the articular surface, separating the medial and lateral condyles. Based on the arrangement of the three condyles of the mandibular process (the medial, lateral, and caudal condyles), the mandibular process can be separated into three major groups:

**Type** and **Type**: three condyles (medial, lateral, and caudal condyles) present on the mandibular process (Figure 6A). Based on whether (1) the caudal condyle is confluent with the lateral condyle and (2) the medial condyle bifurcates into two articular surfaces (medial condyle and lateral trochlea), we identify four different types: **Type-A** and **Type-B** exhibit three clearly separated condyles, whereas the lateral condyle is confluent with the caudal condyle in **Type-A** and **Type-B** (Figure 6A). The medial condyle in **Type-B** and **Type-B** develop a concave articular surface, forming a bifurcated articular surface on the medial condyle (lateral trochlea) (Figure 6A).

**Fig. 6.**
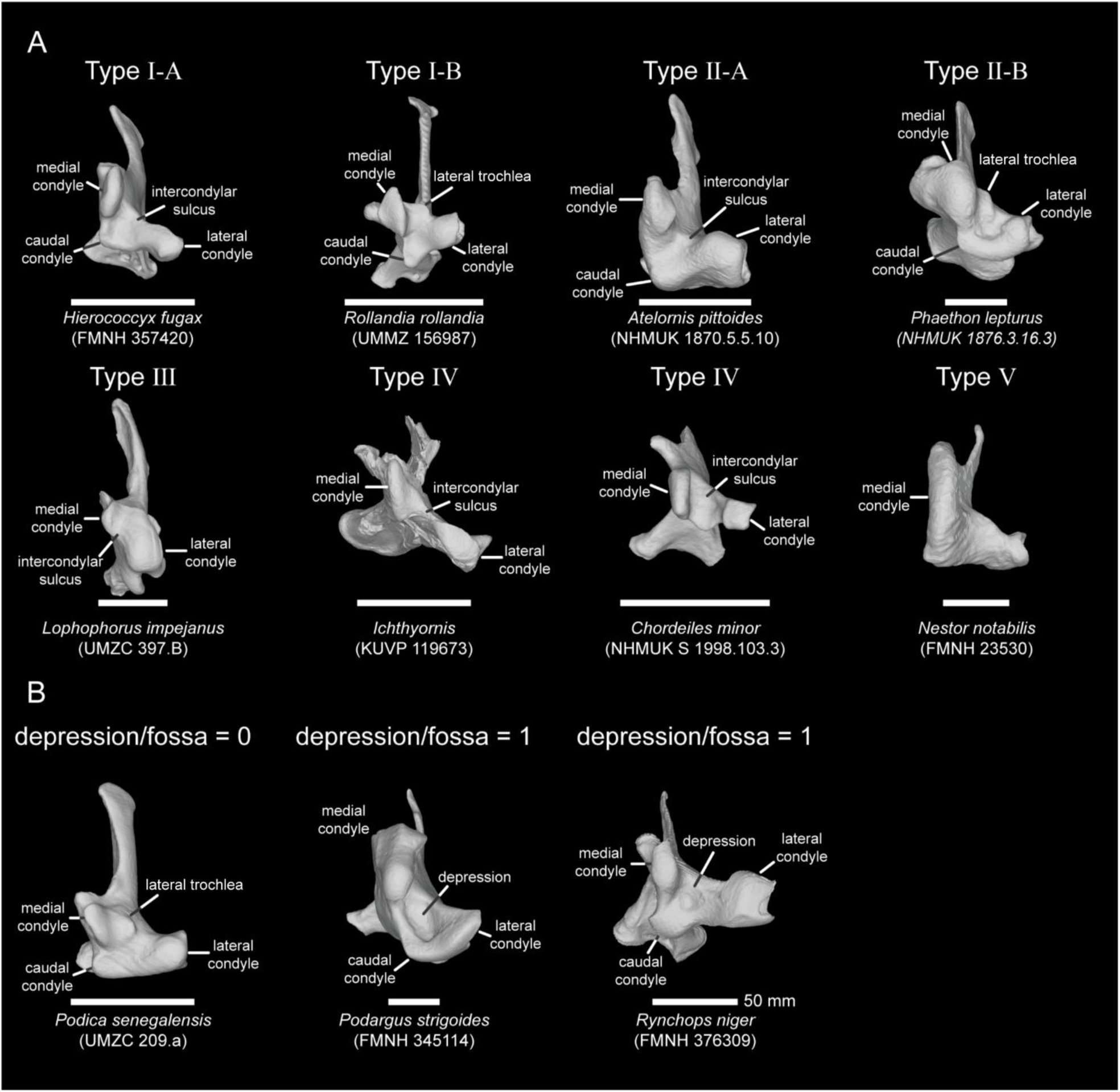
Classification for the types of the mandibular process (A) and the absence/presence of the depression/fossa (B) on the mandibular process in ventral view. Scale bar: 5 mm. (Except for *Rhynchops niger*.)

**Type-A** characterises *Rhea americana* (IMNH R 2585), *Hierococcyx fugax*, some Charadriiformes (*Limosa lapponica*, *Turnix varius*, *Stercorarius skua*, *Glareola pratincola*, *Uria*, *Chroicocephalus novaehollandiae*, and *Rynchops niger*), *Pelagodroma marina*, most examined Suliformes (except *Sula dactylatra* and *Morus bassanus*), some examined Pelecaniformes (*Tigrisoma lineatum*, *Ixobrychus minutus*, *Pelecanus occidentalis*), *Opisthocomus hoazin*, two examined Coraciiformes (*Coracias benghalensis* and *Momotus momota*), some examined Piciformes (*Galbula dea*, *Indicator exilis*, *Psilopogon chrysopogon*, and *Lybius dubius*), *Caracara cheriway*, and Acanthisittidae (Table 2 and Table S1). The intercondylar sulcus in **Type-A** is usually located at the centre of the mandibular process and takes the form of a shallow depression. In some extant birds (e.g., *Hierococcyx fugax*, *Fregata aquila*, *Leucocarbo atriceps*, *Ixobrychus minutus*, *Momotus momota*, and *Psilopogon chrysopogon*), a deep depression (or fossa) is clearly present on the mandibular process.

**Type-B** includes *Rhea americana* (FMNH B 339616), all extent Mirandornithes (Phoenicopteriformes and Podicipediformes), some Charadriiformes (*Recurvirostra avosetta*, *Scolopax rusticola*, and *Numenius phaeopus*), Eurypygiformes, Gaviiformes, ^†^*Madrynornis mirandus*, most Procellariiformes (except *Pelagodroma marina*, *Hydrobates leucorhous*, and *Pelecanoides urinatrix*), two Suliformes (*Sula dactylatra* and *Morus bassanus*), two Pelecaniformes (*Eudocimus ruber* and *Ardea alba*), *Cathartes burrovianus*, *Bucorvus abyssinicus*, and ^†^*Llallawavis scagliai* (Table 2 and Table S1). The intercondylar sulcus in **Type-B** usually takes the form of a shallow depression, but is deeper in some taxa such as *Podilymbus gigas*, *Rhynochetos jubatus*, some Procellariiformes (*Phoebetria palpebrata*, and *Ardenna tenuirostris*), some Suliformes (*Sula dactylatra* and *Morus bassanus*), *Ardea alba*, and *Cathartes burrovianus*.

**Type-A** characterises most Palaeognathae (except *Rhea americana*), some caprimulgiform representatives of Strisores (*Steatornis caripensis* and *Podargus strigoides*), most Otidimorphae (except *Hierococcyx fugax*), *Syrrhaptes paradoxus*, some Charadriiformes (*Pedionomus torquatus* and *Lymnocryptes minimus*), *Eudyptes chrysolophus*, *Balaeniceps rex*, most examined Accipitriformes (except *Cathartes burrovianus*), Strigiformes, Coliiformes, *Leptosomus discolor*, Trogoniformes, some Coraciiformes (*Merops orientalis*, *Atelornis pittoides* and *Todus mexicanus*), four Piciformes (*Bucco capensis*, *Jynx torquilla*, *Picus viridis*, and *Ramphastos ambiguous*), extent Cariamidae, most Falconiformes (except *Caracara cheriway*), Tyranni, and Passeri (Table 2 and Table S1). The intercondylar sulcus in **Type-A** is usually shallow, but it is significantly deeper in some taxa (Cariamiformes, *Ptilonorhynchus violaceus*, and *Falcunculus frontatus*). Finally, a deep depression is present on the mandibular process in some clades (e.g., *Struthio camelus*, *Megalapteryx didinus*, *Crypturellus tataupa*, *Podargus strigoides*, most Otidimorphae, *Balaeniceps rex,* and *Ninox novaeseelandiae*).

**Type-B** includes *Monias benschi*, Gruiformes, ^†^*Palaelodus ambiguus*, four examined Charadriiformes (*Burhinus senegalensis*, *Charadrius vociferus*, *Haematopus ostralegus* and *Jacana jacana*), *Phaethon lepturus*, *Spheniscus humboldti*, *Pelecanoides urinatrix*, Ciconiiformes, *Scopus umbretta*, two examined Bucerotiformes (*Upupa epops* and *Phoeniculus purpureus*), *Alcedo atthis*, and two ^†^Phorusrhacidae (^†^*Andalgalornis steulleti* and ^†^*Psilopterus lemoinei*) (Table 2 and Table S1). The intercondylar sulcus in **Type-B** is usually shallow and located at the centre of the mandibular process, though it takes the form of a deep depression on the mandibular process in *Merops orientalis*.

**Type** and **Type**: two condyles (medial and lateral) form the mandibular process, and the third condyle (caudal condyle) is either poorly developed or absent (Figure 6A). Based on the relative position of these two condyles, the shape of the mandibular process can be divided into two groups: **Type** (medial and lateral condyles are rostrocaudally positioned) and **Type** (medial and lateral condyles are lateromedially positioned).

**Type** includes ^†^*Asteriornis maastrichtensis*, ^†^Pelagornithidae, Megapodiidae, most Anseriformes (except *Mergus merganser*, *Mergellus albellus*, and *Sarkidiornis melanotos*), *Aegotheles cristatus*, most Apodiformes (except *Streptoprocne zonaris*), and *Ptilinopus leclancheri* (Table 2 and Table S1). The intercondylar sulcus is usually shallow but distinctly separates the medial and lateral condyles.

**Type** includes ^†^*Ichthyornis*, most Galliformes (except Megapodiidae), three examined Anseriformes (*Mergus merganser*, *Mergellus albellus*, and *Sarkidiornis melanotos*), most examined caprimulgiform Strisores (except *Steatornis caripensis*, *Podargus strigoides*, and *Aegotheles cristatus*), *Streptoprocne zonaris*, and most Columbidae (except *Ptilinopus leclancheri*) (Table 2 and Table S1). The shape of the intercondylar sulcus varies among taxa in **Type IV**. For instance, the intercondylar sulcus is mediolaterally broad in ^†^*Ichthyornis*, while it is very deep in some Anseriformes (e.g., *Mergus merganser*, *Mergellus albellus*, and *Sarkidiornis melanotos*) and shows a furrow-like shape in most caprimulgiform Strisores.

**Type**: only the medial condyle is prominently developed, forming a remarkably deep joint with the mandible (Figure 6A). The medial condyle is rostrocaudally elongate with an oval shape, and its articular surface extends caudodorsally. This type of mandibular process is unique to Psittaciformes and represents a distinctive osteological synapomorphy of the group (Table 2 and Table S1).

### Pneumatic structures of avian quadrates

Pneumatic foramina of avian quadrates connect with the quadrate diverticulum, which belongs to one of the major tympanic diverticula of the air sac system of bird crania (Witmer, 1990). Previous studies have suggested that the pneumatic diverticula impresses upon the surface of the quadrate to form an excavation or fossa (Witmer, 1990), and, these excavations or fossae may be thin enough to enable gas exchange across the bone surface (Müller, 1908). In some avian quadrates (e.g., *Tribonyx mortierii* and *Turnix varius*), several fossae appear at the position where pneumatic foramina appear in other bird species. Therefore, these fossae were taken into consideration as potential evidence of a capacity for gas exchange in our survey of avian quadrate pneumaticity (marked as ▲ in Table 3 and Table S2). Some quadrates do not show any pneumatic foramina (apneumatic) or fossae for air exchange, such as *Struthio camelus*, some Galliformes (*Odontophorus guttatus* and *Tragopan satyra*), *Treron capellei*, some Gruiformes (*Podica senegalensis*, *Sarothrura elegans*, and *Atlantisia rogersi*), Podicipediformes, *Uria*, Gaviiformes, Sphenisciformes, two Procellariiformes (*Puffinus puffinus* and *Pelecanoides urinatrix*), *Opisthocomus hoazin, Pandion haliaetus*, and *Jynx torquilla*. The apneumaticity of these quadrates may be related to (1) an early ontogenetic stage of some specimens (indeed, our ostrich specimen is recognized as a subadult, and the development of skeletal pneumaticity occurs throughout post-hatching ontogeny; see Plateau et al., 2023 for the pneumaticity of cranial elements and Burton et al., 2023 and Burton et al., 2025 for the pneumaticity of postcranial elements) or (2) adaptations to particular ecological habits, especially aquatic diving (Burton et al., 2023). The morphology of avian pneumatic foramina varies from taxon to taxon, with differing positions and sizes. Based on the position of pneumatic foramina, quadrates were divided into seven different types:

**Type A:** the post-squamosal capitulum foramen (*foramen pneumaticum postcapitulum squamosum*) is caudoventrally located at the squamosal capitulum (Figure 7A). This condition characterises *Apteryx owenii*, *Eudromia elegans*, *Aegotheles cristatus*, Apodiformes, most Cuculiformes (*Tapera naevia*, *Centropus goliath*, and *Hierococcyx fugax*), *Monias benschi*, Tooth-billed pigeon *Didunculus strigirostris*, some examined Charadriiformes (*Pedionomus torquatus* and *Scolopax rusticola*), *Rhynochetos jubatus*, *Necrosyrtes monachus*, Coliiformes, *Phoeniculus purpureus*, *Indicator exilis*, *Falco sparverius*, some Tyranni (*Neodrepanis coruscans* and *Hymenops perspicillatus*), and some Passeri (*Malurus melanocephalus*, *Parus major* (▲), *Bombycilla garrulus*, and *Regulus ignicapillus*) (Table 3 and Table S2).

**Fig. 7.**
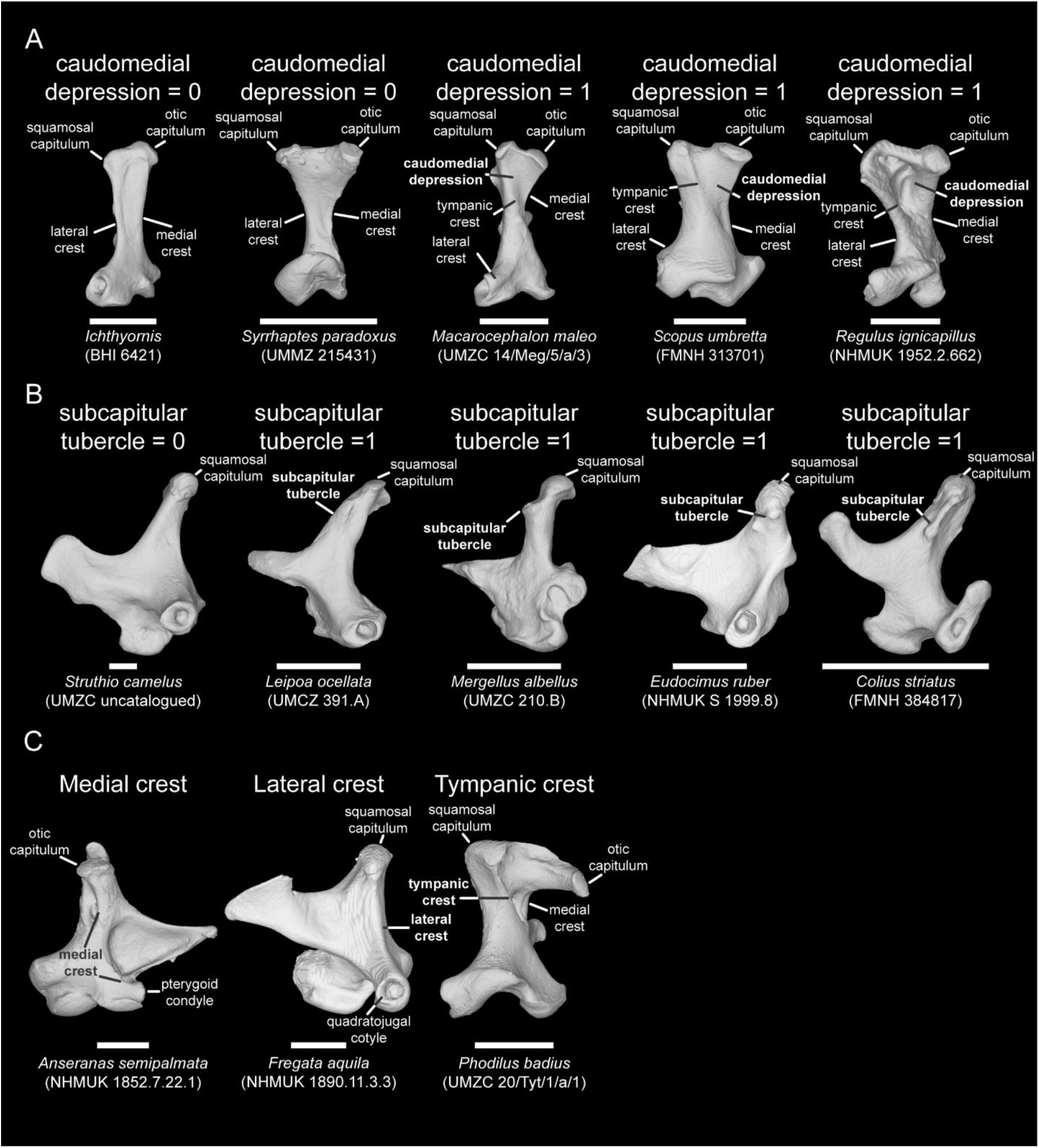
Identification for the absence/presence of the caudomedial depression (A) in caudal view, the absence/presence of the subcapitular tubercle (B) in lateral view, and the crests on the avian quadrate body in medial (left), lateral(middle), and caudal (right) view. Scale bar: 5 mm.

**Type B:** the postotic capitulum foramen (*foramen pneumaticum postcapitulum oticum*) is caudoventrally located at the otic capitulum (Figure 7B). This is observed in Apterygiformes, two examined Tinamiformes (*Eudromia elegans* and *Crypturellus tataupa*), *Aegotheles cristatus*, most examined Apodiformes (except *Chaetura brachyura*), *Tapera naevia*, *Stercorarius skua*, *Urocolius macrourus*, *Alcedo atthis*, *Galbula dea*, *Indicator exilis*, *Micrastur ruficollis*, *Hymenops perspicillatus*, and *Bombycilla garrulus* (Table 3 and Table S2).

**Type C**: the rostromedial foramen (*foramen pneumaticum rostromediale*) is mediodorsally located on the quadrate body, above the basiorbital fossa, and faces rostromedially (Figure 7C). This condition characterises Casuariiformes, ^†^*Megalapteryx didinus*, Tinamiformes (*Eudromia elegans* and *Crypturellus tataupa*), ^†^*Asteriornis maastrichtensis*, most Galliformes (except *Ortalis ruficauda*, *Odontophorus guttatus*, *Tragopan satyra*, and *Perdix cinerea*), Pelagornithidae (^†^*Dasornis toliapica* and ^†^*Osteodontornis sp*), ^†^*Conflicto antarcticus*, Strisores belonging to the subclade Vanescaves (that is, all examined Strisores except the caprimulgids *Caprimulgus europaeus* and *Chordeiles minor*; Chen and Field, 2020), three Apodiformes (*Colibri coruscans*, *Archilochus colubris*, and *Patagona gigas*), three Cuculiformes (*Crotophaga ani*, *Tapera naevia*, and *Hierococcyx fugax*), *Monias benschi*, *Syrrhaptes paradoxus*, two examined Columbiformes (^†^*Raphus cucullatus* and *Leptotila rufaxilla*), some examined Gruiformes (*Porphyrio hochstetteri*, *Rallus limnicola*, *Lewinia striata*, *Tribonyx mortierii* (▴), and *Gallirallus australis*), some examined Charadriiformes (*Recurvirostra avosetta*, *Pedionomus torquatus*, *Jacana jacana*, *Lymnocryptes minimus*, *Limosa lapponica*, *Numenius phaeopus*, *Turnix varius* (▲), and *Glareola pratincola*), Eurypygiformes, *Scopus umbretta*, two Accipitriformes (*Cathartes burrovianus* and *Necrosyrtes monachus*), *Leptosomus discolor*, *Bucorvus abyssinicus, Merops orientalis,* some examined Piciformes (*Galbula dea* and *Bucco capensis*), two ^†^Phorusrhacidae (^†^*Llallawavis scagliai* and ^†^*Psilopterus lemoinei*), *Strigops habroptilus*, and most examined Passeri (except *Bombycilla garrulus*) (Table 3 and Table S2).

**Type D:** the caudomedial foramen (*foramen pneumaticum caudomediale*) is caudally located on the quadrate body, and is restricted by the medial crest and the tympanic crest. This pneumatic foramen is usually accommodated within the caudomedial depression (detailed description in next section; Figure 7D). This condition applies to Lithornithiformes (▴), Casuariiformes, ^†^*Conflicto antarcticus*, Anseriformes, some caprimulgiform Strisores (*Eurostopodus mysticalis* (▲), *Caprimulgus europaeus* (▴), and *Podargus strigoides*), *Corythaeola cristata*, most Cuculiformes (except *Crotophaga ani*), *Monias benschi* (▴), *Balearica pavonina*, Phoenicopteriformes, most Charadriiformes (except *Haematopus ostralegus*, *Jacana jacana*, *Limosa lapponica*, *Scolopax rusticola*, *Stercorarius skua*, and *Uria*), *Eurypyga helias* (▲), *Anhinga anhinga*, some Pelecaniformes (*Eudocimus ruber* and Ardeidae), most Accipitriformes (▲; except *Pandion haliaetus*, *Bueto rufofuscus*, and *Necrosyrtes monachus*), Strigiformes, *Urocolius macrourus*, two Bucerotiformes (*Upupa epops* and *Phoeniculus purpureus* (▲)), two Piciformes (*Bucco capensis* (▴), and *Picus viridis*), two ^†^Phorusrhacidae (^†^*Llallawavis scagliai* and ^†^*Psilopterus lemoinei*), most Psittaciformes (except *Strigops habroptilus*), *Acanthisitta chloris*, and some Passeri (*Malurus melanocephalus* (▲), *Edolisoma tenuirostre* (▴), *Falcunculus frontatus* (▴), *Hirundo rustica* (▲), *Actinodura cyanouroptera* (▴), *Leiothrix lutea* (▴), *Bombycilla garrulus*, *Chlorodrepanis virens* (▲), and *Emberiza calandra*) (Table 3 and Table S2).

**Type E:** the basiorbital foramen (*foramen pneumaticum basiorbitale*) is medially positioned on the quadrate body and is close to the base of the orbital process and the pterygoid condyle (Figure 7E). This applies to ^†^*Ichthyornis*, ^†^*Asteriornis maastrichtensis*, ^†^*Osteodontornis sp.*, some Galliformes (Megapodiidae, Cracidae, *Numida meleagris*, *Colinus virginianus*, *Meleagris gallopavo*, *Perdix cinerea*, *Pavo cristatus*, and *Gallus gallus*), ^†^*Conflicto*, ^†^*Presbyornis*, caprimulgiform Strisores (Caprimulgidae, *Nyctibius griseus*, and *Aegotheles cristatus*), *Chaetura brachyura*, *Corythaeola cristata*, *Syrrhaptes paradoxus*, two Columbiformes in our dataset (*Ptilinopus leclancheri* and *Columba livia*), two sampled Gruiformes (*Tribonyx mortierii* (▲) and *Leucogeranus leucogeranus*), ^†^*Palaelodus ambiguus*, four sampled Charadriiformes (*Charadrius vociferus*, *Stercorarius skua*, *Glareola pratincola*, and *Chroicocephalus novaehollandiae*), *Phaethon lepturus*, most examined Procellariiformes (except the procellariids *Puffinus puffinus* and *Pelecanoides urinatrix*), Ciconiiformes, most examined Suliformes (except *Anhinga anhinga*), *Balaeniceps rex*, most examined Accipitriformes (except *Cathartes burrovianus* and *Pandion haliaetus*), *Leptosomus discolor*, Bucerotiformes (*Upupa epops* and *Phoeniculus purpureus*), two examined Coraciiformes (*Merops orientalis* and *Coracias benghalensis*), *Galbula dea*, two ^†^Phorusrhacidae (^†^*Llallawavis scagliai* and ^†^*Psilopterus lemoinei*), and most examined Falconiformes (except *Micrastur ruficollis*) (Table 3 and Table S2).

**Type F:** the postcapitular foramen (*foramen pneumaticum postcapitulare*) is caudally located on the otic process (Figure 7F). This pneumatic foramen appears in ^†^*Lithornis plebius*, Rheiformes, *Apteryx australis*, two examined Tinamiformes (*Nothoprocta ornata* and *Crypturellus tataupa*), two examined Anseriformes (*Malacorhynchus membranaceus* and *Melanitta nigra*), *Aegotheles cristatus*, four examined Apodiformes (*Hemiprocne comata*, *Streptoprocne zonaris*, *Chaetura brachyura*, and *Phaethornis superciliosus*), *Corythaeola cristata*, two examined Cuculiformes (*Tapera naevia*, and *Hierococcyx fugax*), two examined Columbiformes (^†^*Raphus cucullatus* and *Ptilinopus leclancheri*), four examined Gruiformes (*Psophia crepitans*, *Aramus guarauna*, *Balearica pavonina*, and *Leucogeranus leucogeranus*), some sampled Charadriiformes (*Burhinus senegalensis*, *Haematopus ostralegus*, *Limosa lapponica*, *Scolopax rusticola*, and *Glareola pratincola*), two examined Procellariiformes (*Fulmarus glacialis* and *Macronectes giganteus*), Ciconiiformes, *Pelecanus occidentalis*, two examined Accipitriformes (*Sagittarius serpentarius* and *Necrosyrtes monachus*), *Colius striatus*, Trogoniformes, *Upupa epops*, three examined Coraciiformes (*Atelornis pittoides*, *Todus mexicanus* and *Momotus momota*), most examined Piciformes (except *Bucco capensis*, *Jynx torquilla*, and *Picus viridis*), ^†^*Psilopterus lemoinei*, extent Cariamiformes, *Micrastur ruficollis*, Acanthisittidae, Tyranni, and four sampled Passeri (*Malurus melanocephalus*, *Parus major*, *Regulus ignicapillus* and *Emberiza calandra*) (Table 3 and Table S2).

**Type G:** the dorsal foramen (*foramen pneumaticum dorsale*) is dorsally present between two capitula on the otic process, and faces dorsally (Figure 7G). This condition is found in Columbidae (e.g., *Ptilinopus leclancheri*, *Columba livia*, and *Leptotila rufaxilla*) (Table 3 and Table S2).

### Depressions on avian quadrates

Avian quadrates exhibit several depressions at different positions on the quadrate body, such as the caudomedial depression, the laterocaudal depression (Degrange, 2021; Pascotto and Donatelli, 2003), the laterodorsal depression (Pascotto and Donatelli, 2003), and the lateroventral depression (Pascotto and Donatelli, 2003). Furthermore, a very large depression is located rostroventrally on the otic process of gaviiform quadrates (*Gavia arctica* and *Gavia stellata* sampled in this study). Here, only the caudomedial depression is addressed, as this is an easily recognizable and morphologically comparable character among higher-level bird clades. The caudomedial depression of avian quadrates is caudally located on the quadrate body and is surrounded by the medial crest and the tympanic crest (Figure 8A). The caudomedial depression is usually shallow and wide on most avian quadrates, but it is relatively deep in some clades, such as Accipitriformes.

**Fig. 8.**
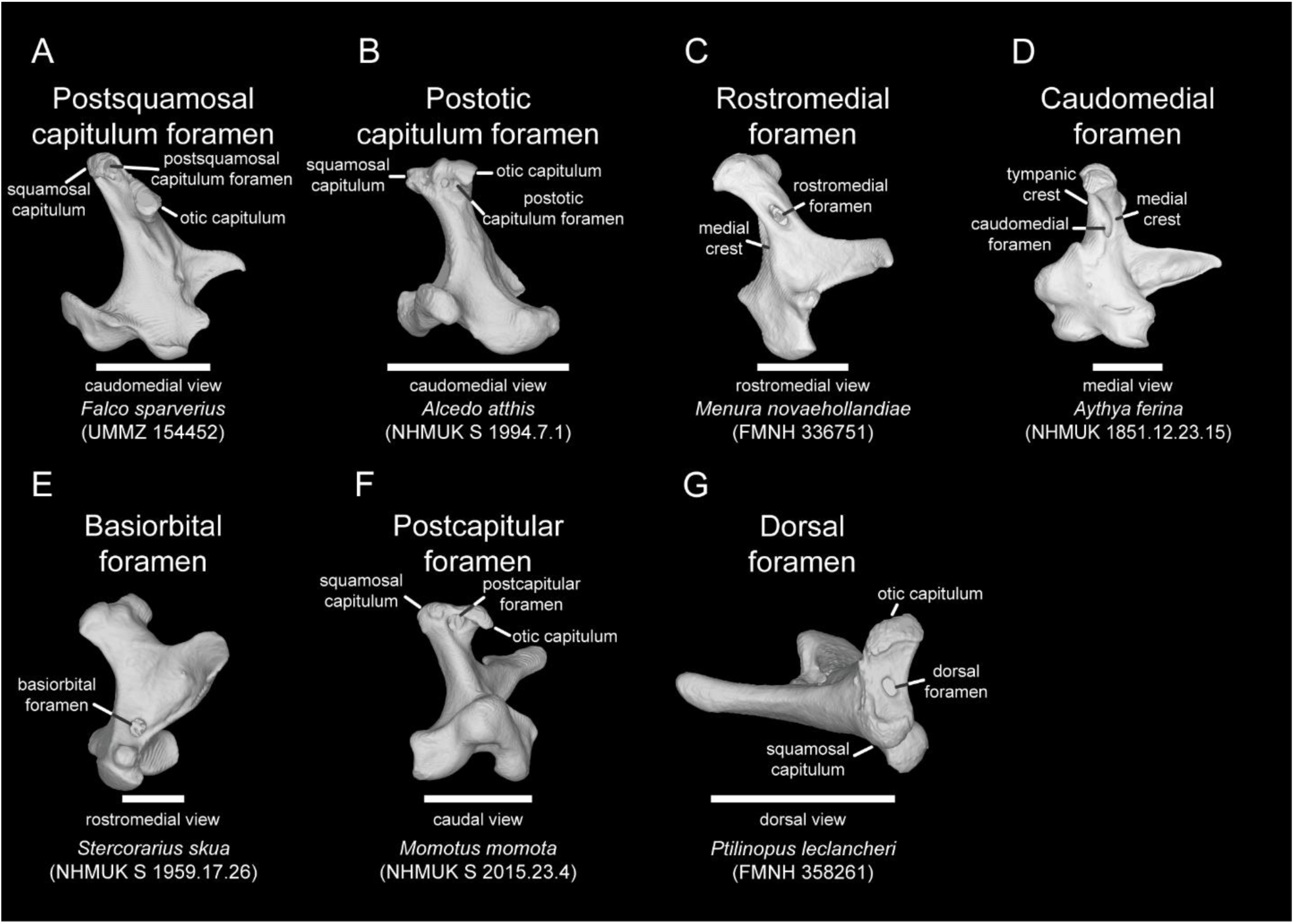
Classification for the types of the pneumatic foramen on avian quadrate in caudomedial view (A-B), rostromedial view (C, E), caudal view (F), and dorsal view (G). Scale bar: 5 mm.

### Crests on avian quadrates

There are three distinct ridges on avian quadrates: the medial crest, the lateral crest, and the tympanic crest (Figure 8C). The medial crest originates at the dorsal margin of the pterygoid condyle and ends at the medial margin of the otic capitulum, the lateral crest originates on the dorsal margin of the quadratojugal cotyle and ends at the lateral margin of the squamosal capitulum, and the tympanic crest extends from its interaction with the medial crest to the caudal margin of the squamosal capitulum. In some Coraciimorphae (e.g., *Upupa epops*, *Phoeniculus purpureus*, *Momotus momota*, *Alcedo atthis*, *Galbula dea*, and *Picus viridis*), a caudal crest is located on the caudal aspect of the quadrate body, and it expands from the tympanic crest to the caudal condyle (Fig. 1B_3_ of Elzanowski and Boles, 2015).

### Quadrate morphological variation within major avian subclades

We present detailed descriptions of neornithine quadrates from major clades in the supplementary, excluding Psittaciformes (parrots) and Passeriformes (passerines). All crown psittaciform quadrates, regardless of subclade, share a suite of distinctive morphological similarities. For passerines, we compare the shape variance across the three extant subordinal subclades (Acanthisitti, Tyranni, and Passeri), but do not undertake detailed comparisons within these subclades, the latter two of which are exceptionally diverse (Ricklefs, 2004; Derryberry et al., 2011; Moyle et al., 2016; Oliveros et al., 2019; Steell et al., 2023; Schmitt and Edwards, 2022).

Avian quadrates exhibit considerable morphological disparity and high phylogenetic signal (Kuo et al., 2024), indicating that major extant subclades often exhibit distinct quadrate geometries. Here, we examined quadrate morphological variance across seven higher clades of Neornithes, and compared shape variance between known stem group representatives of these clades with their extant members. Descriptions and comparisons for Eurypygiformes, Phaethontiformes, Gaviiformes, Sphenisciformes, Ciconiiformes, Coliiformes, Leptosomiformes and Trogoniformes are available in the supplementary.

**(A) Palaeognathae**

Palaeognathae includes the paraphyletic flightless ratites (e.g., ostriches, rheas, cassowaries, emu, and kiwi) and volant tinamous (relevant phylogenetic studies summarised in Widrig and Field, 2022), and each family-level palaeognath clade has its own distinctive quadrate features (see palaeognath quadrates in Plate 2-4 and Table 2). This observation is in line with the interpretation that palaeognaths exhibit lineage-specific biomechanical specialisations of the feeding apparatus (McDowell, 1948). Based on shape variance, crown palaeognath quadrates can be grouped into three types: Struthioniformes-Rheiformes-Casuariiformes, Apterygiformes, and Tinamiformes.

In Struthioniformes-Rheiformes-Casuariiformes, all quadrates exhibit a dorsoventrally broad orbital process (Type in Figure 3A) with a sturdy and deep quadratojugal cotyle (Type in Figure 4A). In Apterygiformes, the quadrate displays a relatively slender orbital process (Type in Figure 3A), a rostrally projecting pterygoid condyle with a rectangular shape (Type in Figure 5A), and a well-separated otic process (Type in Figure 2A). Tinamiform quadrates are characterised by a slender quadrate body, a boat-like orbital process (Type in Figure 3A), a saddle-like quadratojugal cotyle (Type in Figure 4A), and a medially facing pterygoid condyle (Type in Figure 5A).

Lithornithidae are thought to represent a group of Paleogene stem palaeognaths (Nesbitt and Clarke, 2016; Mayr, 2022b; Widrig and Field, 2022; only *Lithornis celetius* and *Lithornis plebius* are included in this study). The shape of Lithornithidae quadrates combines elements of the three abovementioned palaeognath types. For instance, the overall shape is similar to those of ostriches, rheas, cassowaries, and emu with respect to a thick quadrate body and a robust orbital process with a flat tip (Type in Figure 3A). Two capitula on the otic process are packed close to each other, similar to that of most Palaeognathae (Type in Figure 2A, except Apterygiformes and some Tinamiformes). The quadratojugal cotyles are similar to those of Casuariiformes and kiwi (*Apteryx owenii*), with a cup-like articular surface and a thick margin (Type in Figure 4A; see the lateral view of Casuariiformes and *Apteryx owenii* in Plate 3), while the pterygoid condyle of the *Lithornis* quadrate is similar to that of tinamous (Tinamiformes) in that it is ventromedially located with the orbital process facing medially (Type in Figure 5A). Finally, the configuration of the mandibular process in Lithornithidae closely resembles that of kiwi, given that the lateral condyle is confluent with the caudal condyle (Type in Figure 6A).

The quadrates of Dinornithidae (moa; an extinct group of flightless palaeognaths endemic to New Zealand) most closely resemble those of ostriches, with two capitula packed tightly on the otic process (Type in Figure 2A), a deep quadratojugal cotyle (Type in Figure 4A), a blunt pterygoid condyle with a flat articular surface (Type in Figure 5A), a three-condyle mandibular process with a confluent lateral-caudal condyle (Type-B in Figure 6A), and a horizontally oriented caudal condyle (see the caudal view of the quadrates of *Struthio* in Plate 2 and the moa *Megalapteryx didinus* in Plate 3). Although the orbital process of the moa quadrate is dorsoventrally robust with a flat tip, it does not closely resemble other palaeognath quadrates in the overall morphology of the orbital process.

The significant morphological variance observed in palaeognath quadrates confounds strong inferences of the ancestral condition of the palaeognath quadrate. As such, further research on the morphology of cranial elements of stem palaeognaths (e.g., Lithornithiformes) will be necessary to clarify our understanding of morphological evolution of the palaeognath palate (McDowell, 1948; Benito et al., 2022).

**(B) Galloanserae (Galliformes + Anseriformes) and close relatives**

The morphological characters of the quadrates of galloanseran birds (Neognathae: Galloanserae)—the major neornithine subclade uniting Galliformes (chicken-like birds) and Anseriformes (duck-like birds)—have been thoroughly examined both from the perspective of discrete anatomical characters (Livezey and Zusi, 2007; Elzanowski and Stidham, 2010) and with quantitative landmark-based approaches (Kuo et al., 2023). Livezey and Zusi (2007) listed a caudolaterally positioned quadratojugal cotyle relative to the lateral condyle and a pedunculate otic capitulum as two apomorphic galloanseran features (character 513 and 546). Elzanowski and Stidham (2010) identified several additional features, such as the shape of certain muscle attachments, the presence of caudomedial pneumatic foramina, and the shape of the mandibular process, as synapomorphies of Galloanserae, Galliformes, or Anseriformes. For instance, all galloanseran quadrates exhibit an orbital crest and a subcapitular tubercle that serve as muscle attachments, a compact and bicondylar mandibular process, and a wide orbitocondylar angle. Galliform quadrates exhibit a linear orbital crest, an obtuse orbital angle, and a wide intercondylar sulcus on the mandibular process. Anseriform quadrates exhibit a U- or V-shaped orbital crest and a distinct caudomedial foramen. A landmark-based quantitative framework (Kuo et al. 2023) identified a relatively wide quadrate body (mediolateral width >25% of dorsoventral depth) and a lateral condyle with a flat articular surface of the mandibular process as the ancestral condition of the galloanseran quadrate (see Discussion of Kuo et al., 2023). Macroevolutionary patterns of morphological evolution and development of the galloanseran, galliform and anseriform quadrates were investigated in detail by Kuo et al. (2023) and Arnaout et al. (2025) respectively; thus, only broadscale patterns are summarised here.

Galliform quadrates all share two articular surfaces for contact with the pterygoid (Type in Figure 5A, pterygoid condyle and orbitopterygoid facet) and bicondylar mandibular process (Type and in Figure 6A). The otic process and quadratojugal cotyle of galliform quadrates show evidence of major morphological transformations: the otic process appears to have evolved from a condition with well-separated capitula (Type in Figure 2A) to closely packed capitula (Type in Figure 2A) or a medially facing otic capitulum (Type in Figure 2 A); the quadratojugal cotyle evolved from a condition exhibiting a complete rounded margin (Type in Figure 4A) to a saddle-like shape (Type in Figure 4A). By contrast, the otic process of extant anseriform quadrates exhibits limited disparity, with the only notable transformations representing a shift from well-separated capitula in the last common ancestor of crown anseriforms (Type in Figure 2A) to closely packed capitula (Type in Figure 2A) and a distinct intercapitular incisure (Figure 2D) in derived groups, such as Anatidae. Similar to Galliformes, anseriform quadrates all exhibit a bicondylar mandibular process (Type and in Figure 6A), but the orbital process and the pterygoid condyle in Anseriformes encompass considerable morphological disparity (Table 2).

Two previously published stem galloanseran quadrates (Elzanowski and Stidham, 2011; Elzanowski and Boles, 2012; Field et al., 2020) and three recently published stem anseriform quadrates (Tambussi et al., 2019; Houde et al., 2023) provide an opportunity to draw inferences about the ancestral state of galloanseran quadrate geometry. In general, these extinct Galloanserae and Anseriformes share morphological affinities with Galliformes. This is particularly evident in the morphology of articulations between the quadrate and neighbouring bones, such as the shape of the otic process (Type, separated capitula), the quadratojugal cotyle (Type, a deep quadratojugal cotyle), the pterygoid condyle (rounded shape without protruding processes), and the mandibular process (Type, two rostrocaudally aligned condyles). The subcapitular tubercle of *Asteriornis* (Fig. S4 of Field et al., 2020), *Conflicto* (Fig. 6A of Tambussi et al., 2019), *Presbyornis* (Fig. 1 of Elzanowski and Stidham, 2010), the isolated Lance quadrate (Fig. 1B of Elzanowski and Stidham, 2011), the isolated Tingamarra quadrate (QMF 23019, Fig. 1B of Elzanowski and Boles, 2012), and *Anachronornis anhimops* (Fig. 1I of Houde et al., 2023) are positioned laterally relative to the squamosal capitulum, differing from the condition observed in crown Galloanserae. Although the phylogenetic position of *Presbyornis* remains unsettled (as either a crownward stem-anseriform [Elzanowski, 2013; Tambussi et al., 2019] or a crown anseriform outside of Anatidae [Ericson, 1997; Dyke, 2001; Clarke et al., 2005]), quadrate morphology is generally more congruent with the hypothesis that *Presbyornis* is a stem anseriform (see the Discussion of Kuo et al., 2023). Therefore, the ancestral state of the anseriform quadrate can be interpreted as generally similar to that of extant Galliformes.

In addition to well-established total-clade galloanseran fossils (*Asteriornis*, *Presbyornis*, *Conflicto*, *Anachronornis*, and *Danielsavis*), we investigated the quadrates of several additional extinct taxa such as *Brontornis burmeisteri* (Agnolín, 2021) and the extinct clades Pelagornithidae, Gastornithidae, and Dromornithidae, which have often been thought to be closely allied with, or positioned within, Galloanserae (Bourdon, 2005; Mayr, 2011; Worthy et al., 2017; McInerney et al., 2024). We also included recently described quadrates from the phylogenetically controversial *Vegavis* (Torres et al., 2025; Irazoqui et al., 2026).

The *Brontornis* quadrate differs substantially from all known galloanseran quadrates in exhibiting a three-condyle mandibular process (Fig. 6J of Agnolín, 2021), a distinctly acute pterygoid condyle (Fig. 6B and F of Agnolín, 2021), and a pyramidal quadratojugal cotyle with a flat articular surface (Fig. 6B of Agnolín, 2021). *Brontornis* has recently been alternately hypothesized as either a representative of Phorusrhacidae (Cariamiformes; Worthy et al., 2017) or as a stem-anseriform (Agnolín, 2021). If *Brontornis* is a crownward stem-anseriform, then several apparently apomorphic features of anseriform quadrates would need to be reevaluated; as such, quadrate morphology fails to support the hypothesis that *Brontornis* is closely related to Anseriformes.

Moreover, the quadrates of Pelagornithidae, Gastornithidae, and Dromornithidae also differ notably from those of any known total-clade galloanserans. This is especially evident in the geometry of the articulations with neighbouring bones (see detailed comparisons of pelagornithid quadrate shape in the supplementary). For example, an intercapitular incisure is absent from the otic process of most Pelagornithidae (see dorsal view of the Pelagornithidae quadrate, Plate 4), Gastornithidae (*Gastornis parisiensis* [MHNT.PAL.2013.15.2] in Fig. 2A_2_ of Bourdon et al., 2016), and Dromornithidae (*Genyornis newtoni* [NMV P256893] in Fig. 6c of McInerney et al., 2024). The orbital process of Pelagornithidae exhibits a triangular shape (see lateral view of the Pelagornithidae quadrate, Plate 4; Fig. 2F of Mayr and Rubilar-Rogers, 2010), while the orbital process of Gastornithidae is relatively elongate (Fig. 2A_1_ of Bourdon et al., 2016). The orbital process of *Dromornis stirtoni* (Dromornithidae; NTM P3202), on the other hand, shows a robust triangular outline (Fig. 4E of Worthy et al., 2016), similar to that of most anseriform quadrates (see lateral view of anseriform quadrate in Plate 11-14). The pterygoid condyle of Pelagornithidae and Gastornithidae quadrates is dorsally separated from the medial condyle (see rostral view of the Pelagornithidae quadrate, Plate 4), similar to that of all crown galloanseran quadrates. However, the pterygoid condyle of dromornithid quadrates is dorsally adjacent to the medial condyle (McInerney et al., 2024), differing with that of all crown galloanseran quadrates. In addition, Pelagornithidae quadrates lack a subcapitular tubercle and an orbital crest, but gastornithid and dromornithid quadrates exhibit a large and distinct subcapitular tubercle (e.g., *Gastornis* quadrates in Fig. 2A_1_ and Fig. 2B of Bourdon et al., 2016; ts of Dromornithidae quadrates in Fig. 4C, E and G of Worthy et al., 2016; t. sub of *Genyornis newtoni* quadrate in Fig. 6A of McInerney et al., 2024). Only the morphology of the quadratojugal cotyle and the mandibular process in Pelagornithidae, Gastornithidae, and Dromornithidae resembles that of extant Galloanserae in that they display a cup-like morphology with a deep or shallow fossa (similar to some Anseriformes, see lateral view of anseriform quadrates in Plate 11-14) and two condyles (medial and lateral condyles). Importantly, both of these features may represent crown bird symplesiomorphies rather than anseriform apomorphies; as such, these observations echo recent calls for the reassessment of pelagornithid and gastornithid phylogenetic affinities on the basis of other observations of the palate system (Benito et al. 2022; detailed discussion available in Benito et al., 2022 and Kuo et al., 2023; Field et al., 2025; Plateau et al., 2025; Benito et al., 2026). Indeed, a recent phylogenetic analysis of Dromornithidae (McInerney et al., 2024) confirms that further research is needed to decisively untangle the phylogenetic affinities of this bizarre extinct clade.

Though the phylogenetic position of *Vegavis* remains unsettled (Field et al., 2025; Crane et al., 2026), the recently published skull of *Vegavis geitononesos* (MLP-Pv 15-I-7-52) preserves a remarkable articulated palate system, providing an opportunity to compare its quadrate morphology with that of uncontroversial representatives of Galloanserae. The *Vegavis* quadrate exhibits several morphological similarities with those of crown-group Galloanserae, including a bicondylar mandibular process and the presence of the subcapitular tubercle (Elzanowski and Stidham, 2010). In addition, the *Vegavis* quadrate shares several features with fossil total-clade representatives of Galloanserae, such as the absence of a tympanic crest (Fig. 4e of Irazoqui et al., 2026; except in *Asteriornis*) and a circular quadratojugal cotyle with a deep fossa (Fig. 4b of Irazoqui et al., 2026), a condition similar to that observed in *Asteriornis*. Also, the *Vegavis* quadrate displays a combination of characters resembling the quadrates of both extant galliforms and anseriforms. For instance, the orbital process of the *Vegavis* quadrate is elongate with a high aspect ratio (Fig. 4b of Irazoqui et al., 2026), a galliform-like feature, while the orbital process is oriented strongly rostrally, an anseriform-like feature. The quadratojugal cotyle is positioned dorsally, adjacent to the lateral condyle (Fig. 4b of Irazoqui et al., 2026), as seen in most galliforms, and the two condyles of the mandibular process are aligned caudolaterally to rostromedially, as seen in megapodiid galliforms and most anseriforms, and not mediolaterally, as stated in Irazoqui et al., (2026). However, despite these morphological similarities, the *Vegavis* quadrate also possesses several distinctive features compared to the quadrates of uncontroversial representatives of Galloanserae. The otic capitulum is oriented dorsocaudally in *Vegavis* (Fig 4e of Irazoqui et al., 2026), whereas it is oriented dorsally or dorsomedially in most Galloanserae (see Plate 5-14). Moreover, the two capitula of the otic process are aligned rostrocaudally (Fig. 4f of Irazoqui et al., 2026), in contrast to the mediolaterally aligned condition of galloanseran quadrates. The intercondylar sulcus between the two condyles of the mandibular process is narrow and shallow in *V. geitononesos* (Fig. 4a of Irazoqui et al., 2026), while the articulation with the quadrate on the mandible of *V. iaai* (AMNH FARB 30899) suggests a much deeper intercondylar sulcus (Fig. 3a of Torres et al., 2025). This disparity indicates some notable variation in cranial morphology within *Vegavis*. Finally, unlike those of any other birds, the quadrate of *Vegavis* apparently exhibits three distinct articulations with the pterygoid: a pterygoid condyle, an orbitopterygoid facet, and an additional, otherwise undocumented articulation between the rostral portion of the orbital process and the basipterygoid process of the pterygoid (Fig. 5h-j of Irazoqui et al., 2026).

**(C) Strisores (‘Caprimulgiformes’ + Apodiformes)**

Most strisorean quadrates show morphological similarities in the otic process (separated capitula; Type in Figure 2A), and exhibit a mandibular process (bicondylar mandibular process; Type and in Figure 6A) with a furrow-like intercondylar sulcus (see ventral view of the Strisores quadrates in Plate 15-16). Meanwhile, the orbital process and the relative positions of the pneumatic foramen differ among the various major lineages within Strisores. For instance, most nocturnal members of Strisores (except *Steatornis caripensis* and *Podargus strigoides*) and the treeswift examined (*Hemiprocne comata*) exhibit a reduced/underdeveloped orbital process (see the lateral view of the Strisores quadrates in Plate 15-16), while most apodiform quadrates (that is, representatives of Apodidae and Trochilidae but not *H. comata*) show a dorsoventrally broad but relatively short orbital process (Type Ⅲ in Figure 3A). Most non-apodiform strisorean quadrates exhibit medial pneumatic foramina on the quadrate body, such as the rostromedial foramen and the basiorbital foramen (Figure 8; Table 2). By contrast, in Apodiformes, pneumatic foramina are usually present on the caudal side of the otic process, such as the postsquamosal capitulum foramen, the postotic capitulum foramen, and the postcapitular foramen (Mayr and Richter, 2025). Several features of strisorean quadrates have been put forward as diagnostic apomorphies of the clade, including a reduced orbital process (Mayr, 2010; Chen and Field, 2020; Chen and Field, 2024), and a reduced caudal condyle with an elongated medial condyle and a furrow-like intercondylar sulcus on the mandibular process (Character 14 of Mayr, 2010).

A number of Eocene Strisores quadrates have been described (Nesbitt et al., 2011; Mayr, 2015a; Mayr, 2021a; Mayr and Kitchener, 2024a; Mayr and Kitchener, 2024c), which help clarify patterns of evolutionary change in strisorean quadrate morphology. The quadrate of *Masillapodargus longipes* (SMF-ME 3405A) is not well preserved (Fig. 3 a and b of Mayr, 2015a), but its mandibular process exhibits a deep intercondylar sulcus and a strongly projecting medial condyle, similar to that of frogmouths (see caudal view of the *Podargus strigoides* quadrate in Plate 15). The quadrate of *Archaeodromus anglicus* (SMF Av 654) shows some morphological similarities with the quadrates of non-apodiform members of Strisores, such as two separated capitula on the otic process with a distinct intercapitular incisure (Type in Figure 2A; Fig. 2C and F of Mayr, 2021a), a dorsally facing quadratojugal cotyle (Type in Figure 4A; Fig. 2G of Mayr, 2021a), and a rostrodorsally elongate medial condyle with a prominent articular surface. However, it also exhibits some unusual features with respect to muscle attachment sites, its articulation with adjacent cranial elements, and pneumaticity, such as a rostrally protruding pterygoid condyle (Type in Figure 5A; Fig 2A-B of Mayr, 2021a), a three-condyle mandibular process (Type-A in Figure 6A; Fig. 2E of Mayr, 2021a), a well-developed orbital process (Fig. 2A of Mayr, 2021a), presence of a ventrolaterally positioned subcapitular tubercle, and a caudally located pneumatic foramen (Figure 8; Fig. 2D of Mayr, 2021a). The quadrates of *Fluvioviridavis platyrhamphus* (FMNH PA 607) and *F. michaeldanielsi* (NMS.Z.2021.40.168) also share some characteristics with ‘Caprimulgiformes’ (that is, the paraphyletic group of non-apodiform strisoreans), such as separated capitula on the otic process with a well-developed intercapitular incisure (Type in Figure 2A; *F. platyrhamphus* in Fig. 4 [left] of Nesbitt et al., 2011 and *F. michaeldanielsi* in Fig. 7d of Mayr and Kitchener, 2024c), a laterally protruding quadratojugal cotyle with a shallow fossa (Type in Figure 4A; *F. platyrhamphus* in Fig 4 [right] of Nesbitt et al, 2011; *F. michaeldanielsi* in Fig. 7a of Mayr and Kitchener, 2024c), and a rostrocaudally elongate medial condyle with a prominent articular surface (see *F. platyrhamphus* in Fig. 4 of Nesbitt et al., 2011; arf of *F. michaeldanielsi* in Fig. 7a of Mayr and Kitchener, 2024c). However, the *Fluvioviridavis* quadrate differs from most non-apodiform Strisores quadrates, given that it has a caudomedial foramen (‘f’ in Fig. 4 of Nesbitt et al., 2011, although this foramen is only found in *F. platyrhamphus*, not in *F. michaeldanielsi*) and a relatively elongate orbital process with a flat tip (Type in Figure 3A; Fig. 4 of Nesbitt et al., 2011), similar to that of the Oilbird (*Steatornis caripensis*) and hornbills (see lateral views of the *Steatornis* quadrate in Plate 15 and the *Bucorvus* quadrate in Plate 37). Finally, *Fluvioviridavis* preserves a large subcapitular tubercle, different to that of all crown Strisores. Due to its unsettled phylogenetic relationship with respect to extant strisoreans, further work aimed at elucidating the phylogenetic affinities of *Fluvioviridavis* is warranted (Nesbitt et al., 2011; Mayr et al., 2015; Chen et al., 2019), which could help elucidate the ancestral condition of the Strisores quadrate. The quadrate of the apodiform *Eocypselus* (NMS.Z.2021.40.120) exhibits several characters shared with the quadrates of Aegothelidae and Apodiformes, such as the shape of the otic process with a widely separated capitula and a clear Intercapitular incisure (Type Ⅱ in Figure 2A; Figure 2p of Mayr and Kitchener, 2024a), and a pneumatic foramen dorsally located on the otic process (pnf in Figure 2p of Mayr and Kitchener, 2024a). The shape of the quadratojugal cotyle in *Eocypselus* appears to be a flat articular surface with an underdeveloped rim (cdl in Figure 2o of Mayr and Kitchener, 2024a; Type in Figure 4A), similar to that seen in Apodi (i.e. Hemiprocnidae and Apodidae; Plate 16). However, the *Eocypselus* mandibular process shows a bicondylar configuration (Figure 2r of Mayr and Kitchener, 2024a; Type Ⅲ in Figure 6A), with a L-shaped arrangement and a dorsoventrally broadened orbital process (Figure 2o of Mayr and Kitchener, 2024a), which is similar to that of the Trochilidae quadrate (Plate 16-17). The pterygoid condyle of the *Eocypselus* quadrate is not rostrally protruding, similar to that of non-apodiform strisorean quadrates (Plate 15-16) and the hummingbird *Phaethornis* (Plate 17). Interpreting the complex patterns of similarity/dissimilarity of the *Eocypselus* quadrate morphology with respect to other major groups of extant strisoreans might provide some insight into the evolutionary acquisition of the distinctively elongate hummingbird bill (Mayr and Kitchener, 2024a) and its specialised quadrate-driven mechanical function (Rico-Guevara et al., 2024).

**(D) Columbaves** (bustards, cuckoos, turacos, mesites, sandgrouse, and pigeons)

**1. Otidimorphae** (Musophagiformes, Otidiformes, and Cuculiformes)

The three major clades comprising Otidimorphae, Musophagiformes (turacos), Otidiformes (bustards), and Cuculiformes (cuckoos), differ in their quadrate morphology with respect to the orbital process, the quadratojugal cotyle, the quadrate body, and details of pneumaticity. For instance, the orbital process is relatively elongate with a flat tip (Type in Figure 3A) in Musophagiformes and most Cuculiformes, while it is relatively short and robust (Type in Figure 3A) in Otidiformes. The quadratojugal cotyle exhibits notable clade-specific shape variance, with the quadrates of Musophagiformes and Otidiformes exhibiting a relatively shallow fossa (Type in Figure 4A), and the quadrates of Cuculiformes exhibiting either a deep fossa (Type in Figure 4A; see lateral view of *Crotophaga* quadrate in Plate 18) or a flat articular surface (Type in Figure 4A; see lateral view of *Coccyzus* quadrate in Plate 18). The quadrate body is lateroventrally wide in Musophagiformes, lateroventrally slender in Otidiformes, and distinctively twisted in Cuculiformes. The position of the pneumatic foramen varies among Otidimorphae: only a rostromedial foramen is present in Otidiformes, while the pneumatic foramen is located at a different position in the other two clades. In Musophagiformes, pneumatic foramina are located on the caudal aspect of the otic process (postcapitular foramen), the caudal part of the quadrate body (caudomedial foramen), and the medial part of the quadrate body (basiorbital foramen). By contrast, pneumatic foramina on cuculiform quadrates are usually present on the caudal aspect of the quadrate body (caudomedial foramen), the caudal side of the squamosal capitulum (postsquamosal capitulum foramen), or the rostral face of the quadrate (rostromedial foramen) (Table 2).

**2. Columbimorphae** (Mesitornithiformes, Pterocliformes, and Columbiformes)

The three major clades making up Columbimorphae, Mesitornithiformes (mesites), Pterocliformes (sandgrouse), and Columbiformes (pigeons), also exhibit differing morphologies of important quadrate structures including the orbital process, the quadratojugal cotyle, the pterygoid condyle, the mandibular process, the quadrate body, and aspects of pneumaticity. The orbital process of Mesitornithiformes and Pterocliformes is elongate and robust with a flat tip (Type in Figure 3A), while it is usually elongate with a rounded tip in Columbiformes (Type in Figure 3A). The quadratojugal cotyle is relatively deep in Mesitornithiformes and Pterocliformes (Type in Figure 4A), but exhibits a saddle-like shape in most Columbiformes (Type in Figure 4A). The pterygoid condyle sightly protrudes (Type in Figure 5A) in both Mesitornithiformes and Pterocliformes, but it displays two articular surfaces with the pterygoid in Columbiformes (Type in Figure 5A). The mandibular process in Columbimorphae also exhibits pronounced differences between Columbiformes and non-Columbiformes, in that it exhibits only two condyles in Columbiformes (Type and in Figure 6A) and three condyles (Type in Figure 6A) in Mesitornithiformes and Pterocliformes. The quadrate body is lateromedially wide in Mesitornithiformes, but it is relatively slender in Pterocliformes and Columbimorphae. The pneumaticity of the quadrate is highly variable in Columbimorphae, especially with respect to the otic process: for example, no foramen is present on the otic process in Pterocliformes, but one is present in both Mesitornithiformes and Columbimorphae. A dorsal foramen (i.e. the pneumatic foramen located between the two capitula of the otic process; Figure 8G) is only present in Columbimorphae and in the falcon *Micrastur ruficollis* among taxa sampled in this study.

**(E) Gruiformes (cranes, rails and kin)**

The quadrates of Gruiformes are generally distinctive, characterised by a distinct separation between the two capitula of the otic process with a clear intercapitular incisure (Type in Figure 2A), a broad and elongate orbital process (Type in Figure 3A), a deep quadratojugal cotyle (Type in Figure 4A), and two articular surfaces contacting the pterygoid (Type in Figure 5A). All gruiform quadrates have three distinct condyles on the mandibular process with a clearly developed lateral trochlea (Type-B in Figure 6A).

**(F) Aequorlitornithes** (Mirandornithes, Aequornithes, and Charadriiformes)

**1. Mirandornithes** (Phoenicopteriformes and Podicipediformes)

The two clades comprising Mirandornithes, Phoenicopteriformes (flamingos) and Podicipediformes (grebes), share many morphological features in common. For example, both clades exhibit well-separated capitula of the otic process (Type in Figure 2A), an elongate orbital process with a flat tip (Type in Figure 3A), a deep quadratojugal cotyle (Type in Figure 4A), a rostrally directed pterygoid condyle (Type in Figure 5A), and a distinct three-condyle mandibular process with a clearly developed lateral trochlea (Type-B in Figure 6A). However, the morphology of mirandornithine quadrates differs with respect to the arrangement of muscle attachments and the degree of pneumaticity between these two clades, as may be expected considering the divergent feeding modes and ecological habits of these two distinctive lineages. In terms of muscle attachment arrangements, all flamingo quadrates have an orbital crest and a subcapitular tubercle (Figures 3B and 7B), but in Podicipediformes, only the quadrate of *Podilymbus gigas* exhibits an orbital crest, and only *Podiceps taczanowskii* exhibits a subcapitular tubercle. Phoenicopteriform quadrates exhibit a caudomedial pneumatic foramen, whereas all podicipediform quadrates are apneumatic, in keeping with the generally apneumatic nature of their skeletons related to their diving habits.

A Mongolian pan-mirandornithine quadrate (IGM 100/1418) shares some morphological features with those of extant flamingos and grebes (Hood et al., 2019), especially with respect to its articulation with adjacent bones, such as the shape of the quadratojugal cotyle (with a laterally protruding articular surface and a ventrally expanded tip; Fig. 4A, K of Hood et al., 2019), the mandibular process (three distinct condyles with a lateral trochlea; Fig. 4P of Hood et al., 2019), and a distinct medial crest (Fig. 4F of Hood et al., 2019). The stem-phoenicopteriform *Palaelodus ambiguus* (Palaelodidae) (Mayr, 2015), exhibits an interesting mosaic of anatomical similarities of the quadrate shared uniquely with grebes and with flamingos. For instance, the orbital process of the *Palaelodus* quadrate is elongate with a horizontally twisted tip (Fig. 2A of Mayr, 2015b), and the pterygoid condyle of the *Palaelodus* quadrate is rostrally protruding (Fig. 2C of Mayr, 2015b) as in grebes but not flamingos. Moreover, the *Palaelodus* quadrate body is relatively slender with a distinct medial crest (Fig. 2C of Mayr, 2015b), similar to that of grebes. Meanwhile, several morphological characters of the *Palaelodus* quadrate resemble those of crown-flamingo quadrates, such as the presence of an orbital crest and a subcapitular tubercle, and the presence of a caudomedial pneumatic foramen (Fig. 2A and C of Mayr, 2015b). The quadratojugal cotyle of *Palaelodus* is laterally protruding (similar to Podicipediformes) with a rostrally directed tip at the ventral margin (as in Phoenicopteriformes; Fig. 2B of Mayr, 2015b). Finally, unlike any extant Mirandornithes quadrates, *Palaelodus* displays two articular surfaces with the pterygoid, an orbitopterygoid facet (labelled ‘fap’ in Fig. 2C of Mayr, 2015b) and a pterygoid condyle. Surprisingly, one known *Palaelodus* quadrate shows two foramina, the caudomedial pneumatic foramen and the basiorbital pneumatic foramen (see dorsal view of the *Palaelodus* quadrate in Plate 23), diverging from the condition in its extant relatives. Collectively, the morphological differences between the quadrates of *Palaelodus* and extant flamingos may be linked to pronounced biomechanical changes throughout the evolutionary history of total-group Phoenicopteriformes, as extant flamingos exhibit a unique feeding strategy among birds.

**2. Charadriiformes**

The quadrates of Charadriiformes exhibit a remarkable degree of disparity, particularly regarding the morphology of the orbital process (Type, and in Figure 3A) and pterygoid condyle (Type and in Figure 5A). Most charadriiform quadrates show separated capitula of the otic process (Type in Figure 2A) and a three-condyle mandibular process (Type-A in Figure 6A). Charadriiform quadrates also exhibit pronounced variation in patterns of pneumaticity: for instance, the rostromedial foramen and the caudomedial foramen/fossa are present in most charadriiform quadrates, but pneumatic openings or fossae also appear in other positions of the quadrate body, such as the caudal aspect of the otic process (postsquamosal capitulum foramen, postotic capitulum foramen, and postcapitular foramen) and the ventromedial portion of the basiorbital fossa (basiorbital foramen). Furthermore, the quadrates of alcids, represented in this dataset by the murre *Uria*, are apneumatic.

The fossil record of extinct Charadriiformes helps reveal evolutionary patterns in quadrate morphology (Eocene: Bertelli et al., 2010; Musser and Clarke, 2020; Heingård et al., 2021, and Miocene: Olson, 2010). The quadrate of *Morsoravis sedilis* (MGUH 28930; Bertelli et al., 2010) shares numerous morphological features with plovers (Charadriidae; see *Charadrius* quadrate in Plate 25). For instance, two capitula are separated on the otic process of *Morsoravis* (Type in Figure 2A) with a wide and shallow intercapitular incisure. The quadrate body of *Morsoravis* is wide with a deep caudomedial fossa and a ventrolaterally located subcapitular tubercle (Fig. 6B of Bertelli et al., 2010), similar to the condition Charadriidae (see the lateral and caudal view of *Charadrius* quadrate in Plate 25). On the other hand, the orbital process of *Morsoravis* is slightly elongate with a flat tip (Type in Figure 3A; Fig. 6A of Bertelli et al., 2010), which is shared with the quadrates of jacanas (Jacanidae) (Bertelli et al., 2010; see the lateral view of the *Jacana* quadrate in Plate 25). The quadrates of *Nahmavis grandei* (FMNH PA778) and *Scandiavis mikkelseni* (NHMD 625345) are poorly preserved, but some diagnostic features are discernible which are similar to those observed in extant Charadriiformes, such as a deep quadratojugal cotyle (Type in Figure 4 A), and prominent lateral and tympanic crests (Musser and Clarke, 2020; Heingård et al., 2021). The Miocene quadrate assigned to *Feducciavis loftini* (Laridae; USNM 23692) is similar to that of Alcidae (*Uria*) and Laridae (*Chroicocephalus novaehollandiae*; see Plate 26), as shown in the outline of its quadrate body (Fig. 3D of Olson, 2011; similar to the caudal view of the *Uria* quadrate in Plate 26), its distinct medial crest (Fig. 1D of Olson, 2011; similar to the medial view of the *Chroicocephalus* quadrate in Plate 26), the presence of caudomedial and basiorbital foramina (similar to the medial and rostral views of the *Chroicocephalus* quadrate in Plate 26), and the tilted caudal condyle (Fig. 3D of Olson, 2011; similar to the caudal views of both the *Uria* and *Chroicocephalus* quadrates in Plate 26).

**3. Phaethoquornithes** (Eurypygiformes, Phaethontiformes, Gaviiformes, Sphenisciformes, Procellariiformes, Ciconiiformes, Suliformes, and Pelecaniformes)

The quadrate of the Eocene stem-phaethontid *Prophaethon shrubsolei* (SMF Av 602) resembles that of crown Phaethontidae (Mayr, 2015c; see *Phaethon* quadrate in Plate 27), especially with regard to its laterally protruding quadratojugal cotyle with a deep fossa, the rostrally oriented pterygoid condyle, three condyles of the mandibular process with a lateral trochlea, and the wide quadrate body (Fig. 2b-g of Mayr, 2015c); nevertheless, some characters of the *Prophaethon* quadrate differ from those of extant *Phaethon*, presumably reflecting plesiomorphies. For instance, the intercapitular incisure of the *Prophaethon* quadrate is wider and deeper than that of *Phaethon* (Fig. 2g of Mayr, 2015c; see the rostral view of *Phaethon* quadrate in Plate 27). Its medial condyle is less ventrally prominent, and it is located more medially than that of *Phaethon* (Fig. 2c of Mayr, 2015c; see the rostral view of *Phaethon* quadrate in Plate 27). In *Prophaethon*, the caudal condyle is tilted dorsocaudally, resulting in a caudoventrally oriented articular surface, while it is horizontally oriented in *Phaethon* (Fig. 2d of Mayr, 2015c; see caudal view of *Phaethon* quadrate in Plate 27).

The Eocene total-group gaviiform *Nasidytes ypresianus* (NMS.Z.2021.40.24) is similar to extant Gaviiformes in overall quadrate geometry, as illustrated by the shape of its outline, the morphology of the quadratojugal cotyle, the mandibular process, and the absence of any pneumatic foramen (Fig. 2A-E of Mayr and Kitchener, 2022a; see the rostral view of *Gavia* quadrates in Plate 27), but some characters are conspicuously different. For instance, a pit-like fossa is caudally located on the otic process of the *Nasidytes* quadrate (Fig. 2C of Mayr and Kitchener, 2022a), while it is rostrally located on the otic process in *Gavia* (see rostral view of *Gavia* quadrates in Plate 27). The orbital process in *Nasidytes* (Fig. 2A of Mayr and Kitchener, 2022a) is dorsoventrally deeper than that of extant gaviiforms in which this structure is elongate and markedly slender (see the lateral view of *Gavia* quadrates in Plate 27; character 534c in Livezey and Zusi, 2007). The pterygoid condyle of *Nasidytes* is ball-like in shape (Fig. 2D of Mayr and Kitchener, 2022a), differing from *Gavia* which exhibits a slender ovoid shape (see the rostral view of *Gavia* quadrates in Plate 27). Unlike extant Gaviiformes, an orbitopterygoid facet is absent in *Nasidytes* (Fig. 2B of Mayr and Kitchener, 2022a; see the medial view of *Gavia* quadrates in Plate 27). The tympanic crest of the *Nasidytes* quadrate is distinct, forming a clear caudomedial depression (Fig. 2C of Mayr and Kitchener, 2022a), but it is not well-developed in extant Gaviiformes (see the caudal view of *Gavia* quadrates in Plate 27).

The fossil record of penguins is taxonomically and stratigraphically extensive, providing a great opportunity to examine the evolution of quadrate morphology in Sphenisciformes (Bertelli et al., 2006; Ksepka et al., 2008; Ksepka et al., 2012; Degrange et al., 2018). Known fossil sphenisciform quadrates share several anatomical affinities with extant penguins, such as a robust orbital process (character 534b in Livezey and Zusi, 2007), a deep quadratojugal cotyle (Type in Figure 4A), a three-condyle mandibular process, development of key muscle attachment sites (orbital crest and subcapitular tubercle), and the absence of any pneumatic foramina. Nevertheless, evolutionary transformations of three interesting anatomical features are revealed through comparisons of total-group sphenisciform fossils and extant penguins. First, the two capitula of the otic process are well separated (Type in Figure 2A) on the quadrates of the Eocene and Oligocene penguins *Icadyptes salasi* (MUSM 897) and *Kairuku waitaki* (OU 12652; see Fig. 2E of Ksepka et al., 2012), but these two capitula are packed tightly together (Type in Figure 2A) on the quadrates of Miocene and extant penguins (e.g., *Paraptenodytes antarcticus*, AMNH 3338; Fig.8 of Bertelli et al., 2006; *Madrynornis mirandus*, MEF-PV 100, see the dorsal view of the *Madrynornis* quadrate on Plate 27 and the dorsal view of extant sphenisciform quadrates on Plate 28). In addition, the pterygoid articular surface on the quadrate of the Eocene stem penguin *Icadyptes* exhibits only one projecting pterygoid condyle (Type in Figure 5A; Fig. 6 of Ksepka et al., 2008); by contrast, Miocene penguins, such as *Paraptenodytes* (cp and ft in Fig. 9 of Bertelli et al., 2006) and *Madrynornis* (Fig. 5F of Degrange et al., 2018) and extant Sphenisciformes (see the medial view of crown group Sphenisciformes in Plate 28) exhibit two articular surfaces connecting with the pterygoid (Type in Figure 5A). Finally, the subcapitular tubercle is located close to the squamosal capitulum in most fossil taxa, such as *Icadyptes* (am in Fig. 6B of Ksepka et al., 2008), *Kairuku* (am in Fig. 2F of Ksepka et al., 2012), and *Paraptenodytes* (pam in Fig. 8 of Bertelli et al., 2006), while it is located rostroventral to the squamosal capitulum from which it is distinctly separated in the Miocene *Madrynornis* (ts in Fig. 5A of Degrange et al., 2018) and in extant penguins (see the lateral view of extant sphenisciform quadrates in Plate 28). These observations likely correspond to differences in the organization of muscles acting upon the quadrate and may indicate evolutionary changes in feeding biomechanics between giant stemward fossil penguins and extant Sphenisciformes (Degrange et al., 2018).

The quadrates of Procellariiformes generally exhibit well-separated capitula on the otic process (Type in Figure 2A), a robust and elongate orbital process with a flat tip (Type in Figure 3A), a deep quadratojugal cotyle (Type in Figure 4A), a medially positioned pterygoid condyle with a rostrally oriented articular surface (Type in Figure 5A), a three-condyle mandibular process (Type-B in Figure 6A), the presence of a basiorbital pneumatic foramen (Figure 8E), and the absence of a distinct tympanic crest (Figure 7B). The quadrates of *Hydrobates* and *Pelecanoides* differ from most Procellariiformes in the shape of their tympanic crest and orbital process: in *Hydrobates*, the tympanic crest is distinct and forms a deep caudomedial depression (see the caudal view of the *Hydrobates* quadrate in Plate 28), whereas the quadrate of *Pelecanoides* exhibits a relatively robust orbital process with a pointed tip, similar to that of the passerine clade Acanthisittidae (see lateral view of Acanthisittidae quadrate on Plate 42). The quadrate of the Pliocene fossil total-clade albatross *Aldiomedes angustirostris* (NMNZ S.046313) shares numerous similarities with extant Diomedeidae, such as the shape of the caudal condyle, the otic process, and the quadrate body (Fig. 3D of Mayr and Tennyson, 2020), but since much of the quadrate is obscured by matrix, informative comparisons regarding structures such as the orbital process or the pterygoid condyle are not possible.

The quadrates of Suliformes are distinctive, exhibiting well-separated capitula on the otic process (Type in Figure 2A), a triangular orbital process with a pointed tip (Type in Figure 3A), a deep quadratojugal cotyle (Type in Figure 4A), a three-condyle mandibular process (Type-A in Figure 6A), a robust quadrate body with relatively straight medial and lateral crests, and the presence of a subcapitular tubercle and a basiorbital foramen. The shape of the pterygoid condyle on the quadrates of Suliformes is phylogenetically variable. For instance, the quadrate of Fregatidae exhibits two articular surfaces connecting with the pterygoid (Type in Figure 5A; p and po in the medial view of *Fregata aquila* quadrate in Plate 30), while the Sulidae quadrate shows a slightly rostrally directed pterygoid condyle (Type in Figure 5A). The pterygoid condyle of Phalacrocoracidae projects rostrally (Type in Figure 5A), but that of its sister taxon Anhingidae exhibits a flat articular surface (Type in Figure 5A). The quadrate of *Fregata* and *Anhinga* markedly differ in the shape of muscle attachment sites, articulations with neighbouring bony elements, and the arrangement of pneumatic foramina (Table S2). Unlike most Suliformes, the orbital process on the quadrate of *Fregata* is elongate and robust with a flat tip (Type in Figure 3A). In *Anhinga*, the two capitula of the otic process are rostrocaudally aligned, differing from other avian quadrates which are aligned mediolaterally—this arrangement may relate to the particularly narrow skulls of anhingids. In addition, only *Anhinga* exhibits a rostromedial foramen on the quadrate among Suliformes. The Miocene Suliformes *Ramphastosula aguirrei* preserve complete quadrates (Stucchi et al., 2015), and their shape generally resembles that of extant Sulidae: all share a mediolaterally wide quadrate body and a triangular orbital process with a pointed tip (Fig. 4 A5-A6 of Stucchi et al., 2015).

The quadrates of Pelecaniformes generally exhibit well-separated capitula on the otic process (Type in Figure 2A), an elongate and robust orbital process with a flat tip (Type in Figure 3A), a deep quadratojugal cotyle (Type in Figure 4A), and a three-condyle mandibular process (Type and in Figure 6A), but notable disparity is observed in the shape of the pterygoid condyle and the position of pneumatic foramina. For example, the *Eudocimus* quadrate displays a rostrally projecting pterygoid condyle (Type in Figure 5A), while the quadrates of *Tigrisoma*, *Ardea*, and *Scopus* exhibit a slightly rostrally projecting pterygoid condyle (Type in Figure 5A). In *Ixobrychus* and *Balaeniceps*, the quadrate exhibits two articular surfaces for the contact with the pterygoid (Type in Figure 5A). Finally, the quadrate of *Pelecanus* is distinctive in terms of the shape of its articulations with adjacent cranial elements and the arrangement of its pneumatic foramina. For instance, the otic capitulum of the *Pelecanus* quadrate faces medially, while its quadratojugal cotyle and pterygoid condyle exhibit flat articular surfaces. The *Pelecanus* quadrate exhibits a caudally positioned pneumatic foramen on the otic process (postcapitular foramen) (see the dorsal view of the *Pelecanus* quadrate in Plate 32).

The quadrate of the Eocene stem-threskiornithid *Rhynchaeites litoralis* (NMS.Z.2021.40.28) resembles that of crown group Threskiornithidae, in exhibiting closely packed capitula on the otic process with a deep intercapitular incisure (Type in Figure 2A; Fig. 5c of Mayr and Kitchener, 2023a), an elongate orbital process with a flat tip (Type in Figure 3A; Fig. 5a of Mayr and Kitchener, 2023a), a circular quadratojugal cotyle with a deep articular fossa (Type in Figure 4A; Fig. 5a of Mayr and Kitchener, 2023a), and a rostrally protruding pterygoid condyle (Type in Figure 5A; Fig. 5b of Mayr and Kitchener, 2023a). Unlike in crown group Threskiornithidae, the *Rhynchaeites* quadrate displays a three-condyle mandibular process, with a lateral trochlea and confluent lateral-caudal condyles (Type-B in Figure 6A; Fig. 5d-e of Mayr and Kitchener, 2023a). It also exhibits a wide and shallow caudomedial depression (Fig. 5c of Mayr and Kitchener, 2023a), different from the condition in crown group Threskiornithidae (see the caudal view of the *Eudocimus* quadrate in Plate 31). In addition, the *Rhynchaeites* quadrate does not exhibit any pneumatic foramina. The quadrate of the Miocene total-group Ardeidae *Matuku otagoense* (NMNZ S. 50852; HH4) resembles that of crown Ardeidae in the arrangement of articulations with surrounding cranial bones, such as well-separated capitula on the otic process with a deep intercapitular incisure (Type in Figure 2A; Fig. 2A-B of Scofield et al., 2010), a deep quadratojugal cotyle (Type in Figure 4A; Fig. 2C of Scofield et al., 2010), two articular surfaces contacting the pterygoid (Type in Figure 5A; pf and pt in Fig. 2D of Scofield et al., 2010), a medially located pterygoid condyle, a three-condyle mandibular process with a lateral trochlea (Type-B in Figure 6A; Fig. 2E of Scofield et al., 2010), a distinct fossa at the centre of the mandibular process (in in Fig. 2E of Scofield et al., 2010), and the presence of a caudomedial foramen (fm in Fig. 2D of Scofield et al., 2010). Despite these points of resemblance the *Matuku* quadrate body is much narrower mediolaterally than that of crown group Ardeidae (Fig. 2A of Scofield et al., 2010).

**(G) Telluraves** (core landbirds)

**1. Accipitriformes**

The quadrates of Accipitriformes show separated capitula on the otic process (Type in Figure 2A), either a slightly elongate orbital process (Type in Figure 3A) or a ventrally directed orbital process (Type in Figure 3A), a deep quadratojugal cotyle (Type in Figure 4A), either a blunt pterygoid condyle (Type in Figure 5A) or two articular surfaces with the pterygoid (Type in Figure 5A), and a three-condyle mandibular process with confluent lateral and caudal condyles (Type in Figure 6A). In terms of pneumatization, accipitriform quadrates generally exhibit a basiorbital foramen and a caudomedial fossa (Table 2). The quadrate of Osprey (*Pandion haliaetus*) is apneumatic, which is potentially related to its unique foraging and hunting style among Accipitriformes involving plunging into water in pursuit of fish (Bierregaard et al., 2020).

**2. Strigiformes**

The quadrates of Strigiformes (owls) exhibit distinct features with respect to all other birds, such as widely separated capitula on the otic process (Type in Figure 2A) with a caudomedially elongate ‘internal’ otic process (character 548 in Livezey and Zusi, 2007), and a remarkably large caudomedial pneumatic foremen. The quadrate of the Eocene stem strigiform *Ypresiglaux michaeldanielsi* (NMS.Z.2021.40.26; Mayr and Kitchener, 2023b), generally resembles that of living owls in exhibiting two articular surfaces for the pterygoid (Type in Figure 4A; Fig 3g of Mayr and Kitchener, 2023b) and a three-condyle mandibular process with confluent lateral and caudal condyles (Type in Figure 6A; Fig 3h of Mayr and Kitchener, 2023b). However, the quadrate of *Ypresiglaux* lacks two distinct features of extant Strigiformes: development of the internal otic process and a remarkably large caudomedial pneumatic foremen; instead, its capitula are well separated on the otic process (Type in Figure 2A; Fig 3f of Mayr and Kitchener, 2023b), and only a deep caudomedial depression is present on the *Ypresiglaux* quadrate (Fig 3f of Mayr and Kitchener, 2023b). These macroevolutionary transformations of the strigiform quadrate demand future attention to clarify the function of these distinctive features.

**3. Coraciimorphae** (Coliiformes, Leptosomiformes, Trogoniformes, Bucerotiformes, Coraciiformes, and Piciformes)

Despite the poorly preserved and only partially exposed state of most Eocene fossils belonging to the sandcoleid (stem-Coliiformes) taxon *Eoglaucidium* (SMF-ME 11110A and SMF-ME 11110B; Mayr, 2018), its quadrate shows some morphological differences with respect to the quadrates of extant Coliiformes. For instance, the intercapitular incisure of the otic process on the *Eoglaucidium* quadrate is much wider and deeper than that of crown Coliiformes (Fig. 3i-j of Mayr, 2018; see the caudal view of Coliiformes quadrate on Plate 36). Its caudomedial depression is significantly wider and shallower than that of crown Coliiformes (Fig. 3i-j of Mayr, 2018; see the caudal view of Coliiformes quadrate in Plate 36). Moreover, the dorsal margin of the *Eoglaucidium* quadratojugal cotyle is not dorsally expanded into a distinct “process” – one of the synapomorphic features of the coliiform quadrate (character 512b in Livezey and Zusi, 2007). This character has been recognized in other fossil Coliiformes (e.g., *Sandcoleus*, mm in Plate 2 of Houde and Olson, 1992). Further differing from the condition of crown group Coliiformes, no pneumatic foramen is present on the caudal surface of the otic process of the *Eoglaucidium* quadrate (Fig. 3i-j of Mayr, 2018; see caudal view of Coliiformes quadrate in Plate 36).

Two recently published stem group Leptosomiformes quadrates (Mayr and Kitchener, 2022b), *Plesiocathartes insolitipes* (NMS.Z.2021.40.36), *Waltonavis paraleptosomus* (NMS.Z.2021.40.16 and NMS.Z.2021.40.19), and *Waltonavis sp.* (NMS.Z.2021.40.21), share a large number of morphological similarities with extant Leptosomiformes, including well-separated capitula on the otic process with a shallow intercapitular incisure (Type in Figure 2A; Fig. 4c-d and 7d of Mayr and Kitchener, 2022b), a deep quadratojugal cotyle with a broad ventral rim (Type in Figure 4A; Fig. 4a and 7d of Mayr and Kitchener, 2022b), and the presence of a subcapitular tubercle. Our observation of *Waltonavis* differs from that of Mayr and Kitchener (2022b), who state that the quadratojugal cotyle of *Waltonavis* lacks a broad ventral rim. Despite the morphological similarities of the quadrate between these Eocene fossils and extant *Leptosomus*, the fossils differ from extant *Leptosomus* in the shape of the orbital process, and the nature of the mandibular process and pneumaticity. For instance, the orbital process of the *Waltonavis* quadrate is slightly elongate with a broad tip (Fig. 7b of Mayr and Kitchener, 2022b), while that of crown group Leptosomiformes is elongate with a pointed tip (see lateral view of the *Leptosomus* quadrate on Plate 36). The mandibular process of the *Plesiocathartes* quadrate exhibits a three-condyle mandibular process with confluent lateral-caudal condyles (Type-A in Figure 6A; Fig. 4e of Mayr and Kitchener, 2022b), similar to that of extant *Leptosomus*. However, *Waltonavis* displays a three-condyle mandibular process with a lateral trochlea and a confluent lateral-caudal condyle (Type-B in Figure 6A; Fig. 7c of Mayr and Kitchener, 2022b). Finally, the pneumatic foramen of the *Plesiocathartes* quadrate is located close to the basiorbital fossa and the pterygoid condyle (pnf in Fig. 4b of Mayr and Kitchener, 2022b), while it is rostromedially positioned on the quadrate of *Leptosomus* (see the medial view of the *Leptosomus* quadrate on Plate 36).

The quadrate of the Eocene total-clade trogoniform *Eotrogon stenorhynchus* (NMS.Z.2021.40.83) resembles that of crown group trogons in the shape of its rostrally protruding pterygoid condyle (Type in Figure 5A; Fig. 3I of Mayr et al., 2023), as well as its three-condyle mandibular process with confluent lateral-caudal condyles (Type-A in Figure 6A; Fig. 3L of Mayr et al., 2023), and its distinct intercondylar sulcus. Nevertheless, the *Eotrogon* quadrate differs from that of crown group trogons in lacking a medially expanded squamosal capitulum with a distinct intercapitular incisure (Fig. 3K of Mayr et al., 2023), a dorsoventrally deeper orbital process (Fig. 3I-J of Mayr et al., 2023), a deep quadratojugal cotyle (Type in Figure 4A; Fig. 3J of Mayr et al., 2023), and a lateromedially wider quadrate body (Fig. 3K of Mayr et al., 2023).

The quadrates of Bucerotiformes all share similarly deep quadratojugal cotyles (Type in Figure 4A) and exhibit a distinct lateral crest, but differences exist with regard to articulations with neighbouring bones, the morphology of the orbital process, and pneumaticity. Upupiformes such as *Upupa* and *Phoeniculus* exhibit well-separated capitula of the otic process (Type in Figure 2A), an elongate and delicate orbital process, a rostrally projecting pterygoid condyle (Type in Figure 5A), a three-condyle mandibular process with a confluent lateral-caudal condyle and a lateral trochlea (Type-B in Figure 6A), and a caudally located pneumatic foramen on the otic process (postcapitular foramen) (see Plate 37). Hornbills (Bucerotidae) exhibit strikingly different quadrate morphologies including a distinct intercapitular incisure on the otic process, an elongate but dorsoventrally deep orbital process, two articular surfaces connecting to the pterygoid (Type in Figure 5A), a three-condyle mandibular process with a lateral trochlea (Type-B in Figure 6A), and a rostromedially located pneumatic foramen on the quadrate body (rostromedial foramen), as illustrated by the quadrate of *Bucorvus abyssinicus* (see Plate 37).

The quadrates of extant Coraciiformes mostly exhibit closely packed capitula of the otic process (Type in Figure 2A), a rostrally projecting pterygoid condyle (Type in Figure 5A), a three-condyle mandibular process with confluent lateral-caudal condyles (Type-A in Figure 6A), and the presence of a subcapitular tubercle, a basiorbital foramen, and a postcapitular foramen. However, coraciiform quadrates also vary substantially in the shape of the orbital process and quadratojugal cotyle. For instance, the orbital processes of *Merops*, *Todus*, and *Alcedo* exhibit a triangular outline with a pointed tip (Type in Figure 3A), while those of *Coracias*, *Atelornis*, and *Momotus* are elongated with either a flat tip (Type in Figure 3A) or a pointed tip (Type in Figure 3A; see the lateral view of Coraciiformes quadrates on Plates 37-38). In addition, most extant Coraciiformes clades exhibit a deep quadratojugal cotyle (Type in Figure 4A), but *Todus* and *Alcedo* quadrates show a quadratojugal cotyle with incomplete margins. Unlike that of other extant Coraciiformes groups, the quadratojugal cotyle of *Merops* displays a flat articular surface without a clear margin (Type in Figure 4 A; see the lateral view of Coraciiformes quadrates in Plates 37-38).

The quadrates of recently reported of Eocene fossil Coraciiformes differ from extant coraciiforms in the shape of their articulations with neighbouring bones and the extent of pneumaticity (Elzanowski and Boles, 2015; Mayr and Walsh, 2018; Mayr, 2022a; Mayr and Kitchener, 2024b). For instance, two capitula are markedly separated with a wide intercapitular incisure on the otic process (Type in Figure 2A) of the Tingamarra coraciiform quadrate (QMF 22781) (Fig. 1 A1-A3 of Elzanowski and Boles, 2015), *Septencoracias morsensis* (SMF Av 655, Fig. 2 c and e of Mayr, 2022a; NMS.Z.2021.40.143 and NMS.Z.2021.40.144, Fig. 4 e-g of Mayr and Kitchener, 2024b), and *S. simillimus* (NMS.Z.2021.40.146) (Fig. 4 j of Mayr and Kitchener, 2024b). The orbital process of the *Septencoracias* quadrate shows a triangular outline (Fig. 4 e of Mayr and Kitchener, 2024b), similar to that of the Puerto Rican Tody (*Todus mexicanus*) (Plate 38), while its quadratojugal cotyle displays a shallow fossa, similar to that of *Merops* (Type in Figure 4A; Fig. 2 a-b of Mayr, 2022a; Fig 4 e-g, j of Mayr and Kitchener, 2024b). Unlike that of most Coraciiformes and *Septencoracias*, the Tingamarra coraciiform quadrate exhibits two articular surfaces connecting to the pterygoid (Type in Figure 5 A; *po* and *pt* in Fig. 2 A1 of Elzanowski and Boles, 2015). The mandibular process of the quadrates of an unnamed London Clay member of Coracii (NMS G.2014.54.1) (Mayr and Walsh, 2018) and *Septencoracias* show a similar morphology to that observed in most extant Coraciiformes (Type-A in Figure 6A), and are especially similar to the quadrate of *Coracias benghalensis* (see the caudal and ventral views of the *Coracias* quadrate on Plate 37). However, the caudal condyle of the quadrate belonging to the unnamed fossil representative of Coracii flattens lateromedially (cdc in Fig. 3h-j of Mayr and Walsh, 2018), which resembles that of some Bucerotiformes quadrates more closely than Coraciiformes (see the caudal and medial view of *Upupa* and *Phoeniculus* quadrates in Plate 37). Meanwhile, the medial condyle of the *Septencoracias* quadrate shows a deep articular surface (cdm in Fig. 2c-f of Mayr, 2022a), which corresponds morphologically with that of *Leptosomus* (see the rostral and ventral view of the *Leptosomus* quadrate on Plate 36). Unlike that of other extinct Coraciiformes, the mandibular process of the Tingamarra coraciiform quadrate exhibits a lateral trochlea (mt in Fig. 2 A_2_ of Elzanowski and Boles, 2015), which is only found on the quadrate of some Alcedinidae (e.g., *Alcedo*) among Coraciiformes (see the ventral view of the *Alcedo* quadrate on Plate 38); however, its caudal condyle is dissimilar to that of *Alcedo* in its outline shape and the direction of its orientation: the caudal condyle of the Tingamarra quadrate is flat on its dorsal surface, but the caudal condyle of the *Alcedo* quadrate shows a bulb-like structure facing caudally (see the lateral and medial views of the *Alcedo* quadrate on Plate 38). The quadrates of all Eocene stem-Coraciiformes display pneumatic foramina but differ in their position. For example, a large and significant caudomedial foramen is present on the Tingamarra quadrate (fm in Fig. 2A_1_ of Elzanowski and Boles, 2015), while a postcapitular foramen and a basiorbital foramen is present on the unnamed London Clay Coracii quadrate (pot in Fig.3 h-j of Mayr and Walsh, 2018) and the *Septencoracias* quadrate (fpm in Fig. 2d of Mayr, 2022a), respectively. Our comparisons of quadrate morphology among extant and fossil Coraciiformes indicate that the last common ancestor of crown-Coraciiformes likely exhibited features distributed more broadly across Coraciimorphae (e.g., Bucerotiformes and Leptosomidae), and therefore, further scrutiny will be required to clarify the evolutionary history of quadrate morphology in this diverse and morphologically variable clade.

The quadrates of Piciformes mostly exhibit well separated capitula of the otic process (Type in Figure 2A) with an indistinct intercapitular incisure, an elongate orbital process with a flat tip (Type in Figure 3A), a deep quadratojugal cotyle (Type in Figure 4A), a rostrally projecting pterygoid condyle (Type in Figure 5A), a three-condyle mandibular process (Type-A and Type-B in Figure 6A), a distinct tympanic crest, a broad caudomedial depression, and the presence of a postcapitular pneumatic foramen. Despite general morphological similarities among Piciformes quadrates, notable disparity exists in the shape of the quadratojugal cotyle, the medial condyle, and the positions of pneumatic foramina. For example, the quadratojugal cotyle of the quadrates of *Indicator* and *Psilopogon* display an incomplete margin (Type in Figure 4A), while *Jynx* and *Lybius* exhibit a flat articular surface connecting with the jugal (Type in Figure 4A). Additionally, the medial condyle of the quadrates of Ramphastides (the clade uniting barbets and toucans, including *Psilopogon*, *Lybius*, and *Ramphastos*) all show a prominent articular surface with a distinct lateral margin (see the lateral view of *Psilopogon*, *Lybius*, and *Ramphastos* quadrates in Plate 39). Finally, pneumatic foramina are caudally located on the otic process of *Galbula* and *Indicator*, while the *Jynx* quadrate is apneumatic.

**4. Cariamiformes**

Phorusrhacidae are a group of giant, flightless stem-cariamiforms, and the quadrates of phorusrhacids primarily differ from those of extant cariamiforms in the morphology of their articulations with adjacent bones (Agnolín and Chafrat, 2015; Degrange et al., 2015; Mayr, 2016; Degrange, 2020). For instance, the two capitula on the otic process of *Llallawavis scagliai* (MMP 5050) and *Psilopterus lemoinei* (YPM-PU15402) are widely separated (Type in Figure 2 A) with a lateromedially broad intercapitular incisure, unlike the condition in crown Cariamiformes where the separation is narrower (Type in Figure 2A). All Phorusrhacidae quadrates exhibit an elongate and robust orbital process with a pointed or a flat tip (see the lateral view of Phorusrhacidae quadrates in Plate 40), while those of all crown group Cariamiformes are elongate with a flat tip and a second, diminutive, dorsally projecting tip (see the lateral view of Cariamidae quadrates in Plate 40). The pterygoid condyle projects rostrally on the quadrate of the *Llallawavis* quadrate (Type in Figure 5A), unlike the condition observed in most Phorusrhacidae and extant Cariamidae. In addition, most Phorusrhacidae quadrates exhibit a three-condyle mandibular process with confluent lateral-caudal condyles, a lateral trochlea, and a deep intercondylar sulcus (Type-B in Figure 6A), as seen in *Llallawavis*, *Andalgalornis steulleti*, *Psilopterus*, and *Patagorhacos marshi* (NHMUK-A516; Fig. 9D of Degrange, 2020); nevertheless, the mandibular process of *Bathornis grallator* (CM 9377) and *Patagorhacos terrificus* (MPCN-PV-377) lack the lateral trochlea (Fig. 3-7 of Mayr, 2016 for *Bathornis*; Fig. 4A of Agnolin and Chafrat, 2015 for *Patagorhacos*). Additionally, the mandibular process of most Phorusrhacidae quadrates is lateromedially broader and sturdier than that of extant Cariamidae due to a strongly reduced caudal condyle (see the ventral view of Cariamidae quadrates in Plate 40). Furthermore, the caudal condyle is usually caudally projecting among Phorusrhacidae, as seen in *Patagorhacos* (cc in Fig. 4C of Agnolín and Chafrat, 2015; Fig. 9D of Degrange, 2020), *Andalgalornis* (Fig. 9E of Degrange, 2020), *Llallawavis* (ventral view of the *Llallawavis* quadrate in Plate 40), and *Psilopterus lemoinei* (ventral view of *Psilopterus* quadrate in Plate 40). The Phorusrhacidae quadrate body is robust and lateromedially wide with a deep caudomedial depression and a distinct tympanic crest, differing in all these respects from that of extant Cariamidae. Finally, the extant cariamiform quadrate shows a postcapitular foramen on the otic process, while the extinct groups show two pneumatic foramina on the quadrate body (the rostromedial foramen and basiorbital foramen). The presence of a caudomedial foramen on the quadrate of *Llallawavis* is uncertain (it is labeled as absent in Degrange et al., 2015, yet it is labelled as a small foramen in Degrange, 2020), whereas it is large on the quadrate of *Psilopterus* (fcm in Fig. 9A of Degrange, 2020; the medial view of *Psilopterus* quadrate in Plate 40).

**5. Falconiformes**

Quadrates of the Eocene group Masillaraptoridae have recently been described (Mayr and Kitchener, 2021), and despite their hypothesised phylogenetic position as stem falconiforms, the quadrates of Masillaraptoridae differ from those of crown group Falconiformes in several respects, including the shape of the otic process, the mandibular process, and the position of pneumatic foramina. For instance, though the two capitula are well-separated on the quadrate of the masillaraptorid *Danielsraptor phorusrhacoides* (NMS.Z.2021.40.12) (Type in Figure 2A), the squamosal capitulum of *Danielsraptor* projects as far dorsally as the otic process, unlike the quadrates of crown group Falconiformes which exhibit a significantly elevated squamosal capitulum. Furthermore, a posteriorly directed protuberance is present on the caudodorsal margin of the quadratojugal cotyle, confluent with the caudal condyle in all surveyed crown-falconids; however, this protuberance is absent on the quadrate of *Danielsraptor*. In addition, the arrangement of the mandibular process on the quadrate of *Danielsraptor* resembles that of crown group Cariamiformes with its confluent lateral-caudal condyles and the absence of an intercondylar sulcus (Fig. 1L of Mayr and Kitchener, 2021; see the ventral view of the Cariamiformes quadrate on Plate 40), unlike the quadrate of crown group Falconiformes. Finally, a pneumatic foramen is located on the caudal aspect of the *Danielsraptor* quadrate, within the caudomedial depression (caudomedial foramen; fpn in Fig. 1I-J of Mayr and Kitchener, 2021).

**6. Psittaciformes**

In Psittaciformes (parrots), the skeletal elements involved in cranial kinesis, such as the palatines, pterygoids and quadrates, are greatly modified from the condition seen in other birds (Livezey and Zusi, 2007) as a result of specialisations favouring exaggerated prokinesis of the upper jaw (Burton, 1974). As such, parrot quadrates are easily identified by a suite of distinctive features such as a rostrally projecting pterygoid condyle (Type in Figure 5A), a deep quadratojugal cotyle (Type in Figure 4A), the presence of a caudomedial foramen, and a single-condyle mandibular process that is otherwise unseen among extant birds (Type in Figure 6A). Additionally, parrots exhibit an especially deep medial condyle on the mandibular process, corresponding with the presence of the caudal fossa on the mandible (character 703b in Livezey and Zusi, 2007). A few fossil parrot quadrates have been described, such as the Pliocene *Nandayus vorohuensis* (MLP 94-IV-1-1) (Carril et al 2014) and the Pleistocene *Agapornis roseicollis* (TM 70525) (Stidham, 2010), although unsurprisingly these relatively recent crown psittaciforms exhibit quadrates resembling those of extant parrots, with a deep quadratojugal cotyle facing rostrolaterally (Type in Figure 4A), a rostrally projecting pterygoid condyle (Type in Figure 5A), and a single-condyle mandibular process (Type in Figure 6A). Understanding the evolutionary origins of the striking specialisations of the cranial kinetic system of extant parrots demands future research attention, and we hope that new fossil discoveries will cast light on this fascinating biomechanical system.

**7. Passeriformes**

The major groups of passerines generally exhibit morphologically distinctive quadrate morphologies, and therefore improvements in our understanding of passerine quadrate morphology may prove helpful for identifying the subordinal affinities of isolated fossil passerine quadrates. For instance, the two capitula of the otic process are usually well-separated in Acanthisitti and Passeri, two of the major passerine subclades (Type in Figure 2A), but quadrates belonging to Tyranni usually exhibit closely packed capitula (Type in Figure 2A) with a shallow intercapitular incisure. The orbital process of passerine quadrates is usually elongate with a flat tip (Type in Figure 3A), but in Acanthisitti, the orbital process exhibits a triangular outline with a pointed tip (Type in Figure 3A). The quadratojugal cotyle of Acanthisitti quadrates is shallow (Type in Figure 4A), while in Tyranni the quadratojugal cotyle protrudes laterally and exhibits a saddle-like margin with a shallow articular fossa (Type in Figure 4A), and the quadratojugal cotyle in Passeri is usually deep (Type in Figure 4A). The pterygoid condyle projects rostrally (Type in Figure 5A) in all Passeriformes. Acanthisitti quadrates exhibit a three-condyle mandibular process (Type-A in Figure 6A), while Tyranni and Passeri exhibit a three-condyle mandibular process with confluent lateral-caudal condyles (Type-B in Figure 6A). Although most passerine quadrates lack a subcapitular tubercle, the lateral crest of most Passeri quadrates protrudes dorsally. This dorsal projection likely performs a similar anatomical function to the subcapitular tubercle in other avian groups. Regarding pneumaticity of the quadrate, both Acanthisittidae and Tyranni generally exhibit a postcapitular foramen; by contrast, in Passeri, a rostromedial foramen is usually present on the quadrate body.

The quadrates of several Eocene fossil representatives of Psittacopasseres (the clade uniting crown group Psittacidae and Passeriformes) have recently been described (Svendsen, 2015; Mayr, 2020b; Mayr, 2021b; Mayr and Kitchener, 2022c; Mayr and Kitchener, 2023c), and these fossils exhibit a remarkable degree of morphological disparity, with features reminiscent of those seen in other groups of Australaves as well as crown group Passeriformes. For instance, the two capitula of the otic process are well separated (Type in Figure 2A) with a shallow intercapitular incisure in *Tynskya waltonensis* (SMF Av 652), *Parapsittacopes bergdahli* (SMF Av 653), *Psittacomimus eos* (NMS.Z.2021.40.38), *Primozygodactylus* cf. *danielsi* (NMS.2021.40.47), Zygodactylidae (NMS.2021.40.61), *Primozygodactylus major* (PMO 212.659), ?*Pulchrapollia eximia* (NMS.Z.2021.40.64), and *?Pulchrapollia* sp. (NMS.Z.2021.40.66), similar to those of most extant passerine quadrates. However, the squamosal capitulum of the quadrates belonging to *Tynskya, Primozygodactylus* (PMO 212.659) and ?*Pulchrapollia* project significantly further dorsally than the otic capitulum does (Fig. 2a and g of Mayr, 2021b; Fig. 11D of Svendsen, 2015; Fig. 8F and J of Mayr and Kitchener, 2023c), similar to that of most crown Falconiformes (see the rostral view of the falconiform quadrates on Plate 40-41). The quadratojugal cotyles of most Eocene fossil Psittacopasseres quadrates are laterally protruding, similar to that of crown passerines, except *Primozygodactylus major* (Fig. 11D of Svendsen, 2015). The quadratojugal cotyle of *Tynskya* and ?*Pulchrapollia* is deep with a circular margin (Type in Figure 4A; Fig. 2d and j of Mayr, 2021b; Fig. 8D and I of Mayr and Kitchener, 2023c), similar to that of most Australaves (i.e. Cariamiformes, Falconiformes, Psittaciformes, and Passeriformes), while both *Psittacomimus* and Zygodactylidae (NMS.2021.40.61) show a shallow fossa on the quadratojugal cotyle, similar to that of Acanthisittidae. Furthermore, the quadratojugal cotyle displays a saddle-like shape with a shallow fossa in *Primozygodactylus* cf. *danielsi* (Type in Figure 4A; Fig. 7E and L of Mayr and Kitchener, 2022c), like that of Tyranni. Similar to the condition in crown group Passeriformes generally, the pterygoid condyle projects rostrally (Type in Figure 5A) in all known fossil total-clade Passeriformes. Eocene fossil Psittacopasseres quadrates mostly exhibit a three-condyle mandibular process with confluent lateral-caudal condyles (Type-A in Figure 6A), but the quadrates of both *Tynskya*, *Parapsittacopes* and ?*Pulchrapollia* show a three-condyle mandibular process with confluent lateral-caudal condyles and a lateral trochlea (Type-B) (*Tynskya*: arf in Fig. 2c and i of Mayr, 2021b; *Parapsittacopes*: arf in Fig. 2F and G of Mayr, 2020b; ?*Pulchrapollia*: arf in Fig. 8D and I of of Mayr and Kitchener, 2023c). The quadrate of *Primozygodactylus major* displays a three-condyle mandibular process (Type in Figure 6A; Fig. 11F of Svendsen, 2015), similar to that of Acanthisittidae. All Eocene fossil Psittacopasseres quadrates show a deep caudomedial depression, corresponding to the morphology seen in most Passeri (oscine) quadrates. The lateral crest of the quadrates of *Psittacomimus* and Zygodactylidae (NMS.2021.40.61) is distinct and rostrally pointed, similar to that of some Passeri quadrates, while only the *Primozygodactylus* cf. *danielsi* quadrate displays a subcapitular tubercle, differing from most passerine quadrates. Unlike other Eocene stem Psittacopasseres and most Passeri quadrates, *Tynskya*, *Primozygodactylus major* and ?*Pulchrapollia* quadrates lack a caudally positioned pneumatic foramen on the otic process, such as the postsquamosal capitulum foramen, postotic capitulum foramen, or postcapitular foramen. Instead, the quadrate of *Tynskya* shows a basiorbital foramen (fpm in Fig. 2b and h of Mayr, 2021b) and *Primozygodactylus major* may exhibit a rostromedial foramen (fb? in Fig. 11B of Svendsen, 2015). The quadrate of *?Pulchrapollia* sp. displays a caudomedial foramen (pnf in Fig. 8K of Mayr and Kitchener, 2023c). On the basis of these features, as well as features of the quadratojugal cotyle and mandibular process, it appears likely that homoplasy among fossil psittacopasserans and crown Passeriformes may at present confound efforts to establish clear macroevolutionary patterns in quadrate morphology among total-group passeriforms, especially in light of continuing uncertainty regarding the phylogenetic relationships of early Psittacopasseres (Mayr, 2022b; Mayr and Kitchener, 2022c). This situation echoes the ongoing challenge of comprehensively characterising morphological variation across the rest of the skeleton of extant and fossil passeriforms (Steell et al., 2023; Steell et al., 2025; Somogyi et al., in review). Further work elucidating the morphology and phylogenetic interrelationships of Eocene fossil Psittacopasseres will be necessary to clarify the evolutionary history of the passerine quadrate.

## Conclusions

This investigation proposes a comprehensive nomenclature system for avian quadrates and presents descriptions and comparisons among major extant clades and relevant fossil representatives. This work has highlighted several previously identified diagnostic features for major crown clades from earlier literature, such as the elongation of the otic capitulum of owls (character 548 in Livezey and Zusi, 2007), the shape of the orbital process of Gaviiformes and Spheniscidae (character 534 in Livezey and Zusi, 2007), and the distinctive medial condyle of the mandibular process of parrots (corresponding to character 703b in Livezey and Zusi, 2007). Some clades illustrate clear evolutionary patterns with respect to distinct features of the quadrate, such as the shape of the otic process of Galliformes and the relative position of the subcapitular tubercle in stem and crown group Sphenisciformes. These comparisons suggest that assumptions regarding the ancestral condition of several avian crown clades may need to be reevaluated, such as Anseriformes (ducks and kin), Phoenicopteriformes (flamingos), Strigiformes (owls), and Cariamiformes (seriemas). Thoroughly understanding the shape variance of the bird quadrate provides an opportunity to evaluate hypotheses of the phylogenetic placement of fossils in light of recent phylogenomic investigations (e.g., Prum et al., 2015; Kimball et al., 2019; Oliveros et al., 2019; Braun and Kimball, 2021; Kuhl et al., 2021; Sangster et al., 2022; Stiller et al., 2024; Wu et al., 2024; Zhao et al., 2025). Additionally, this work highlights interpretive challenges raised by the as-yet unsettled phylogenetic relationships of certain fossil groups (e.g., Pelagornithidae, Gastornithiformes, certain fossil Strisores and Psittacopasseres), which precludes the confident assessment of macroevolutionary patterns of morphological features of the quadrate between these groups and their nearest living relatives. We hope the improved objectivity and consistency of avian quadrate nomenclature proposed here will facilitate future efforts to extract the maximum amount of useful phylogenetic information from fossil bird quadrates, and enable homologous morphological comparisons between crown birds and non-neornithine avialans (e.g., *Archaeopteryx*: Kundrát et al., 2019; O’Connor et al., 2025; Enantiornithes: Wang et al., 2021; Wang et al., 2022; Chiappe and Navalón et al., 2024; Jeholornithiformes: Hu et al., 2023) and non-avian theropod dinosaurs (Hendrickx et al., 2015). Moreover, since the avian quadrate provides several key muscle attachment sites related to avian cranial kinesis (e.g., *M. pesudotemporalis profundus* on the orbital process and *M. protractor pterygoidei et quadrati* medially attached on the quadrate body [Baumel and Witmer, 1993]), a revised understanding of quadrate morphology will help clarify how the quadrate drives the motion of neighbouring bones in the context of an integrated cranial kinetic system. Finally, the comparisons presented herein will help provide a robust foundation for further investigations into the evolution of avian quadrate pneumaticity across avian phylogeny and ontogeny (e.g., Plateau et al., 2023).

## ACKNOWLEDGMENTS

We thank K. Smithson (Cambridge Biotomography Centre) for access to micro-CT scanning facilities. We thank Dr. T. Tsuihiji (National Museum of Nature and Science, Japan) for providing the CT scans and figures of *Osteodontornis sp.* (NSM PV-18696). We thank Dr. F.J. Degrange (Centro de Investigaciones en Ciencias de la Tierra [CICTERRA], Argentina.) for providing CT scans of *Madrynornis mirandus* (MEF-PV 100) and 3D models of *Llallawavis scagliai* (MMP 5050), *Andalgalornis steulleti* (FMNH P14357), and *Psilopterus lemoinei* (YPM PU 15402). If anyone is interested in the 3D models of avian quadrates abovementioned, please contact them directly. We also thank Prof. Dr. Kirsten Grimm (the Institute of Geosciences of Johannes-Gutenberg-University Mainz, Germany) for access to the specimen *Palaelodus ambiguus* (GPIM Op 395).

## DATA AVAILABILITY STATEMENT

The data that support the findings of this study are openly available in MorphoSource (https://www.morphosource.org/projects/000810101), reference no. 000810101. Three-dimensional data will be made openly accessible from MorphoSource upon manuscript acceptance.

---

**Plate 1.**
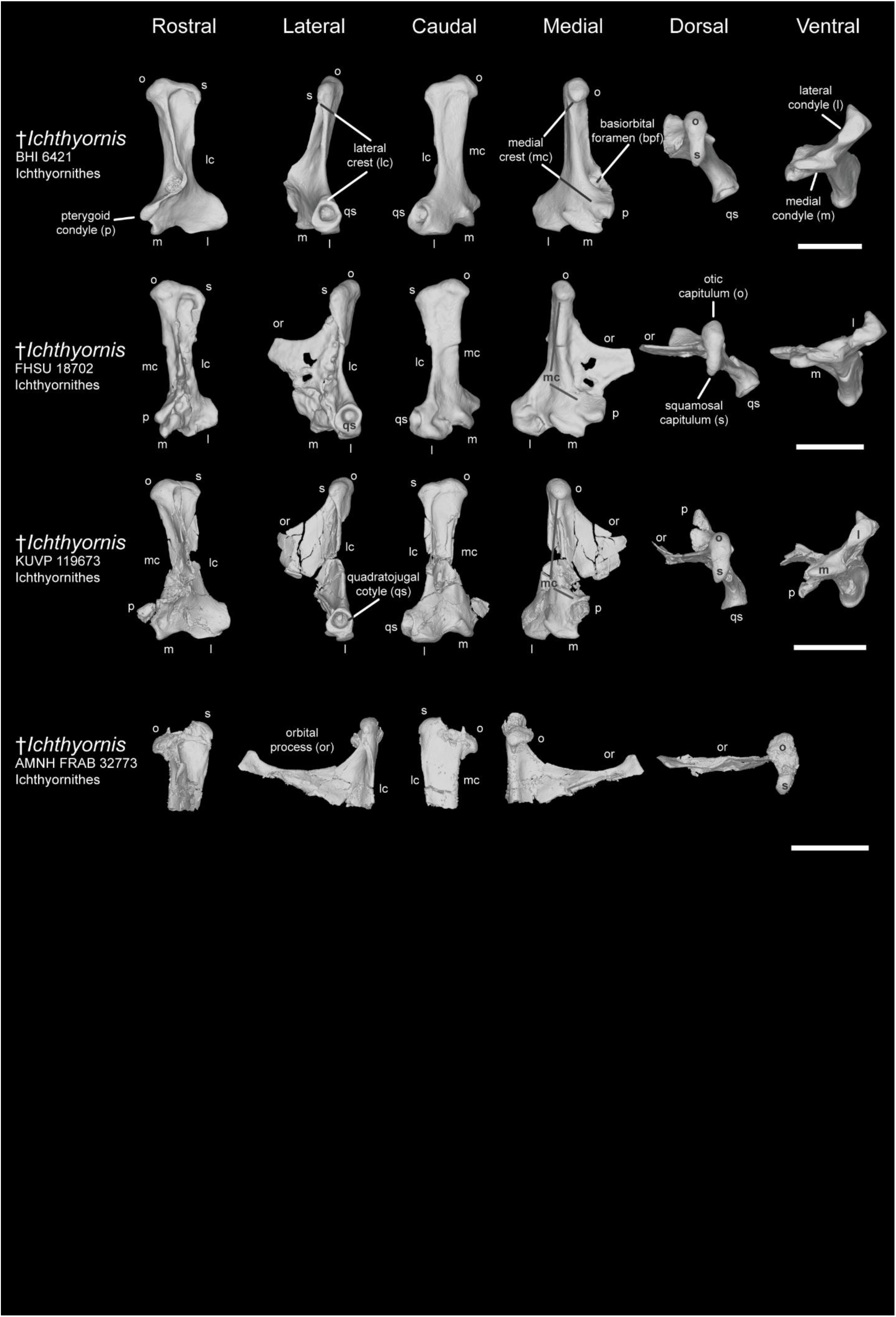
Comparison of avian quadrates among Ichthyornithes. Comparison of *Ichthyornis dispar* (BHI 621), *I. dispar* (FHSU 18702), *I. dispar* (KUVP 119673), and *I. dispar* (AMNH FRAB 32773) in rostral, lateral caudal, medial, dorsal, and ventral view. Scale bar: 5 mm.

**Plate 2.**
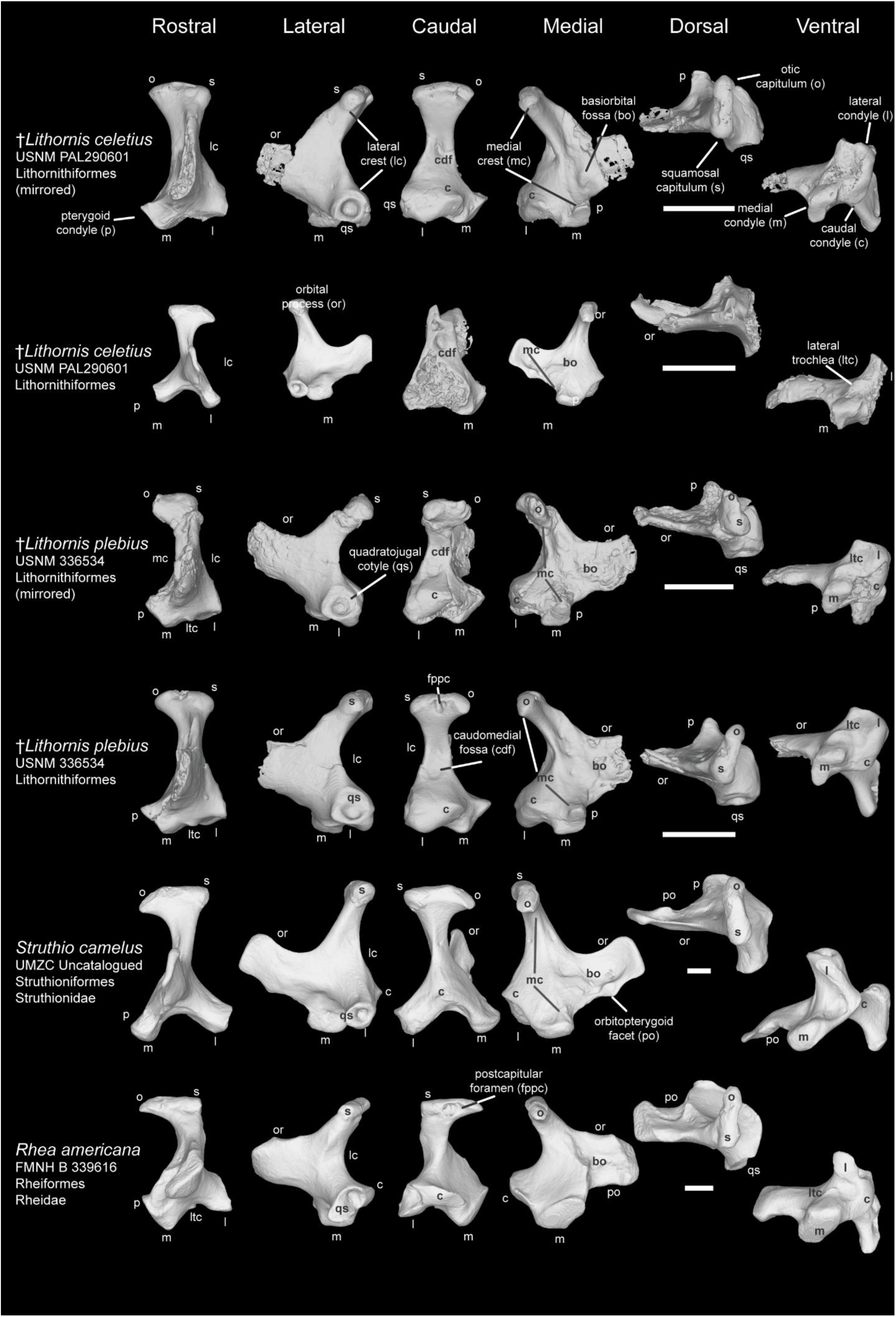
Comparison of avian quadrate among Palaeognathae Ⅰ (Lithornithiformes, Struthioniformes, and Rheiformes). Comparison of *Lithornis celetius* (USNM PAL290601), *L. plebius* (USNM 336534), *Struthio camelus* (UMZC uncatalogued), and *Rhea americana* (FMNH B 339616) in rostral, lateral caudal, medial, dorsal, and ventral view. Scale bar: 5 mm.

**Plate 3.**
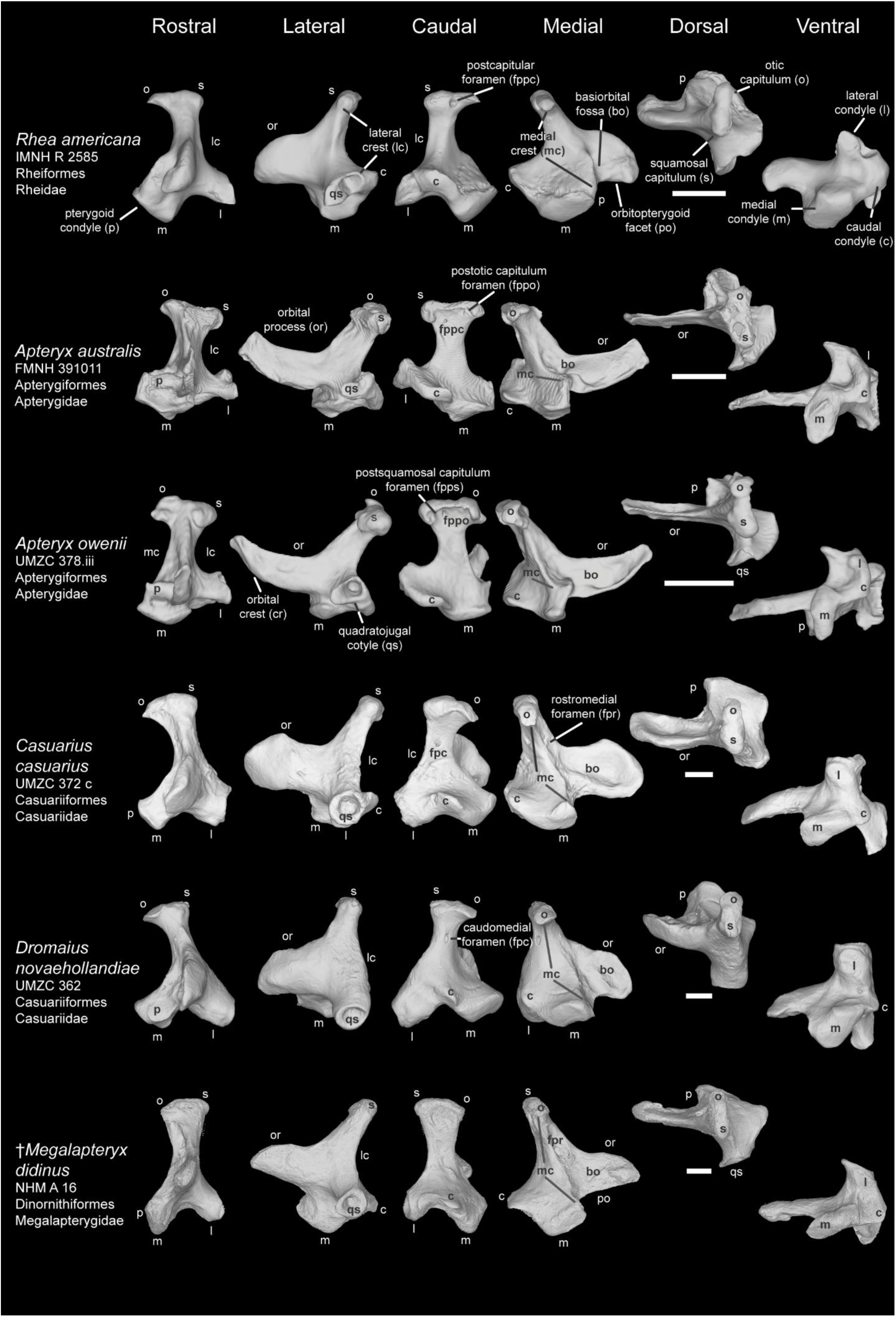
Comparison of avian quadrate among Palaeognathae Ⅱ. Comparison of *Rhea americana* (IMNH R 2585), *Apteryx australis* (FMNH 391011), *A. owenii* (UMZC 378.iii), *Casuarius casuarius* (UMZC 372.c), *Dromaius novaehollandiae* (UMZC 362), *Megalapteryx didinus* (NHM A 16) in rostral, lateral caudal, medial, dorsal, and ventral view. Scale bar: 5 mm.

**Plate 4.**
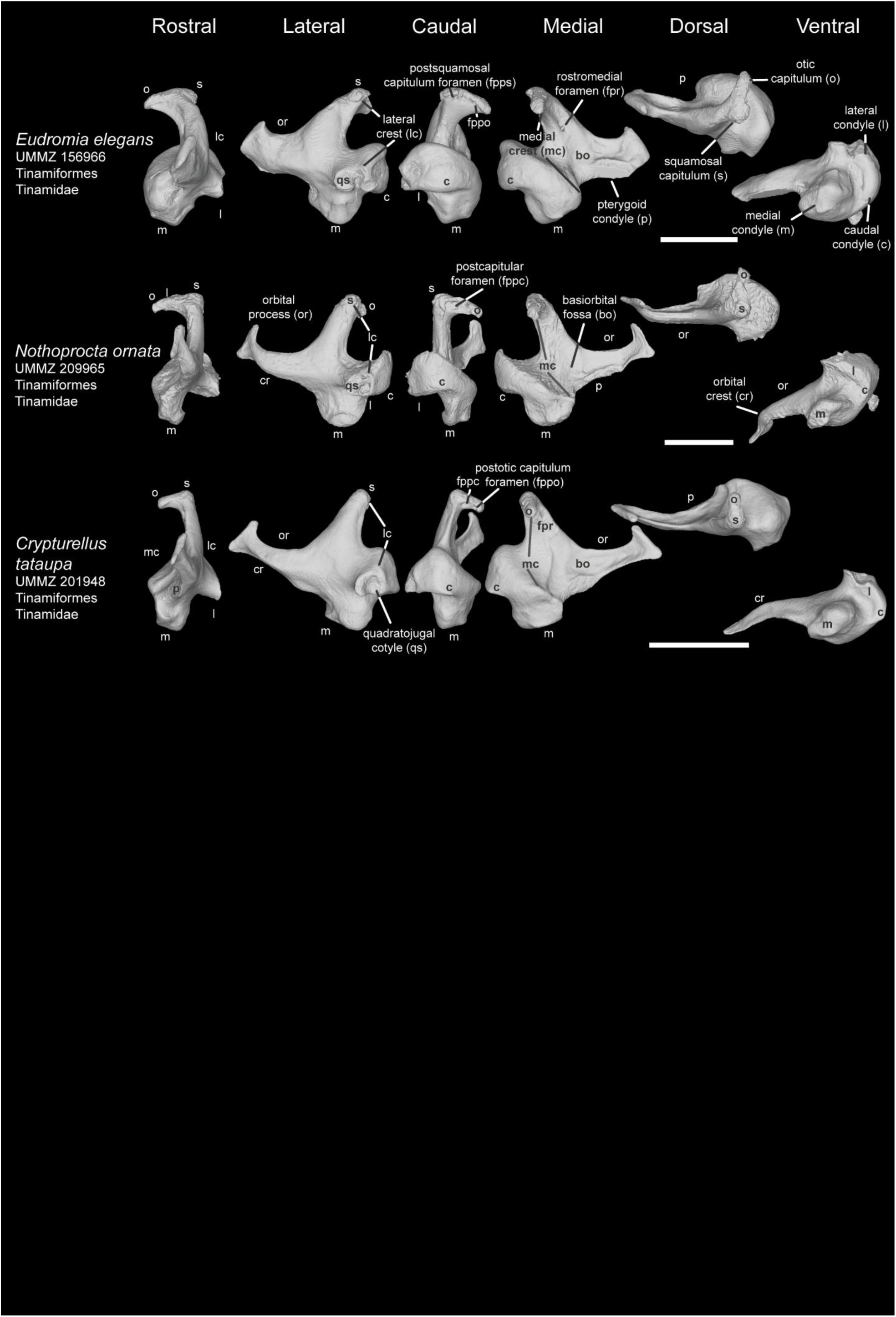
Comparison of avian quadrate among Palaeognathae Ⅲ (Tinamiformes). Comparison of *Eudromia elegans* (UMMZ 156966), *Nothoprocta ornata* (UMMZ 209965), and *Crypturellus tataupa* (UMMZ 201948) in rostral, lateral caudal, medial, dorsal, and ventral view. Scale bar: 5 mm.

**Plate 5.**
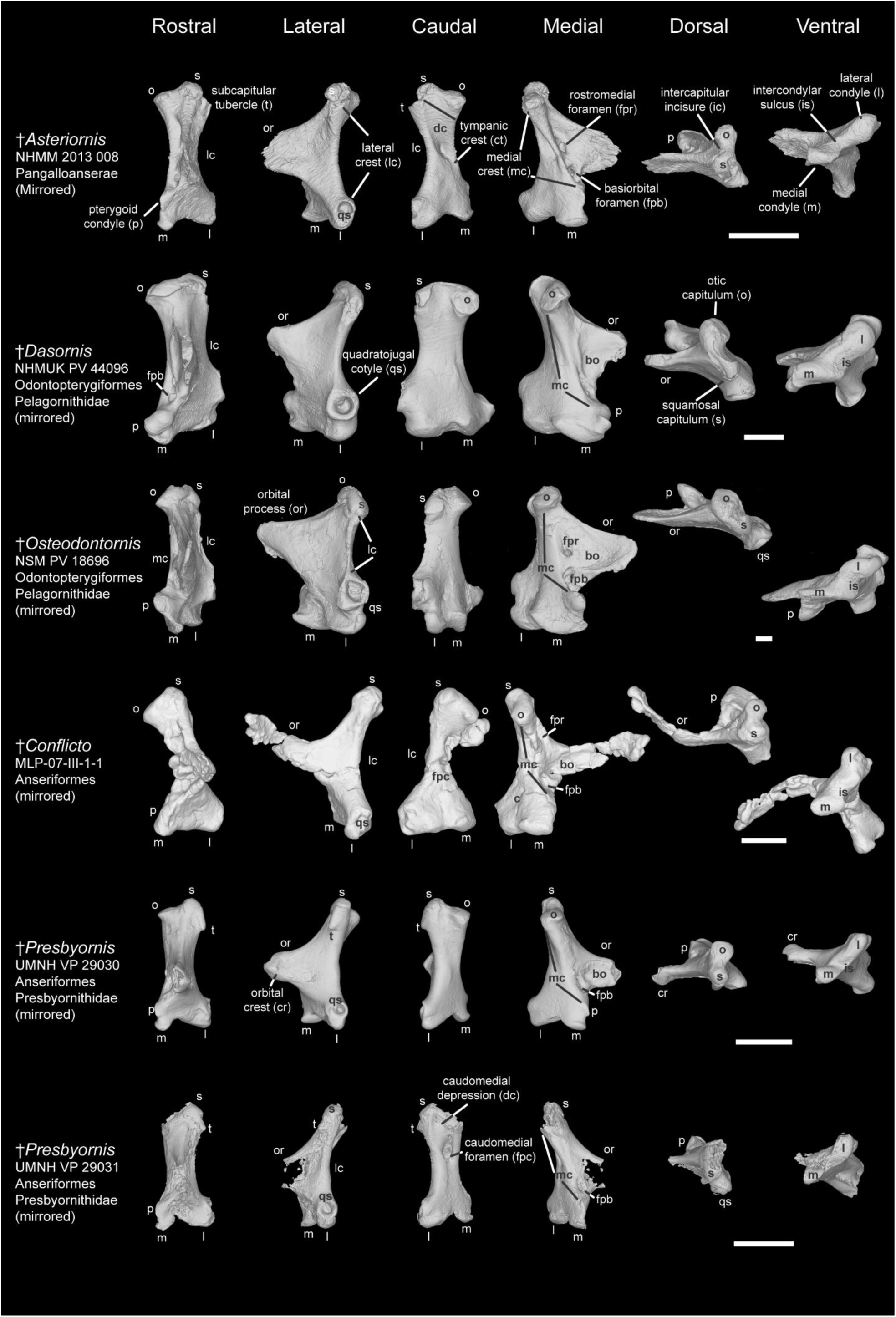
Comparison of avian quadrate in extinct Galloanserae relatives and Pelagornithidae. Comparison of *Asteriornis maastrichtensis* (NHMM 2013 008), *Dasornis toliapica* (NHMUK PV 44096), *Osteodontornis sp.* (NSM PV 18696), *Conflicto antarcticus* (MLP-07-III-1-1), *Presbyornis pervetus* (UMNH VP 29030), and *P. pervetus* (UMNH VP 29031) in rostral, lateral caudal, medial, dorsal, and ventral view. Scale bar: 5 mm.

**Plate 6.**
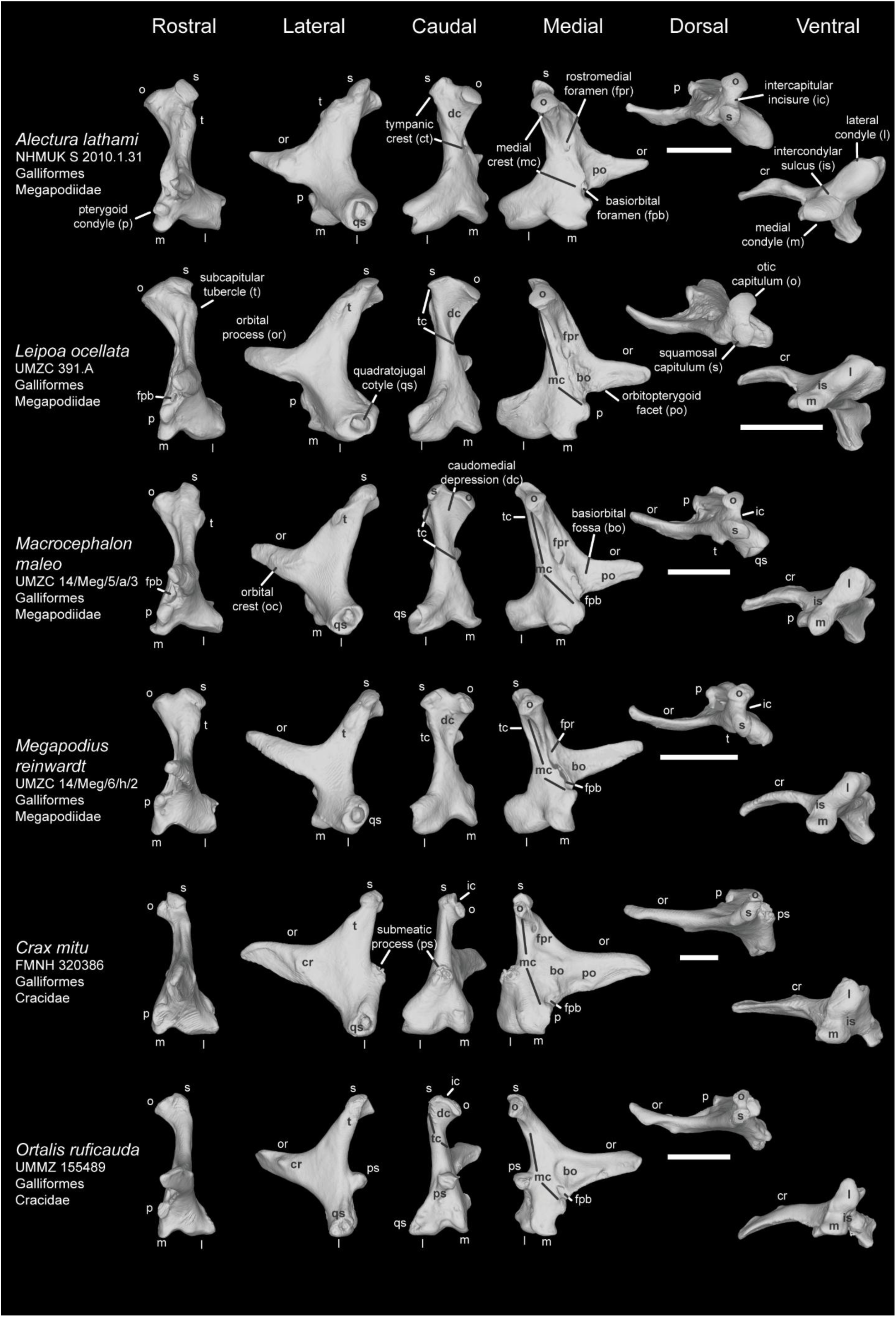
Comparison of avian quadrate among Galloanserae Ⅰ (Galliformes). Comparison of *Alectura lathami* (NHMUK S 2010.1.31), *Leipoa ocellata* (UMZC 391.A), *Macrocephalon maleo* (UMZC 14/Meg/5/a/3), *Megapodius reinwardt* (UMZC 14/Meg/6/h/2), *Crax mitu* (FMNH 320386), and *Ortalis ruficauda* (UMMZ 155489) in rostral, lateral caudal, medial, dorsal, and ventral view. Scale bar: 5 mm.

**Plate 7.**
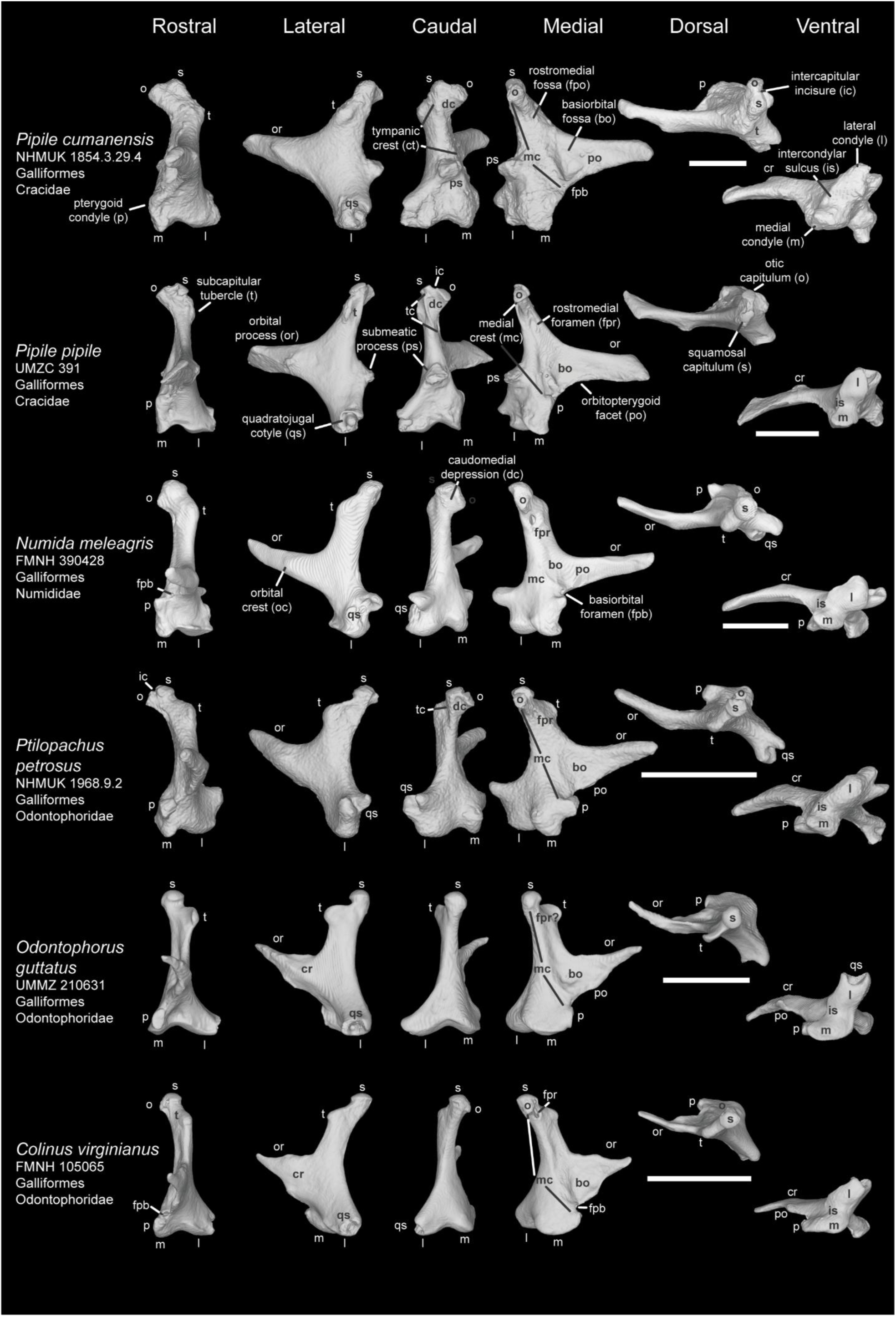
Comparison of avian quadrate among Galloanserae Ⅱ (Galliformes). Comparison of *Pipile cumanensis* (NHMUK 1854.3.29.4), *P. pipile* (UMZC 391), *Numida meleagris* (FMNH 390426), *Ptilopachus petrosus* (NHMUK 1968.9.2), *Odontophorus guttatus* (UMMZ 210631), and *Colinus virginianus* (FMNH 105065) in rostral, lateral caudal, medial, dorsal, and ventral view. Scale bar: 5 mm.

**Plate 8.**
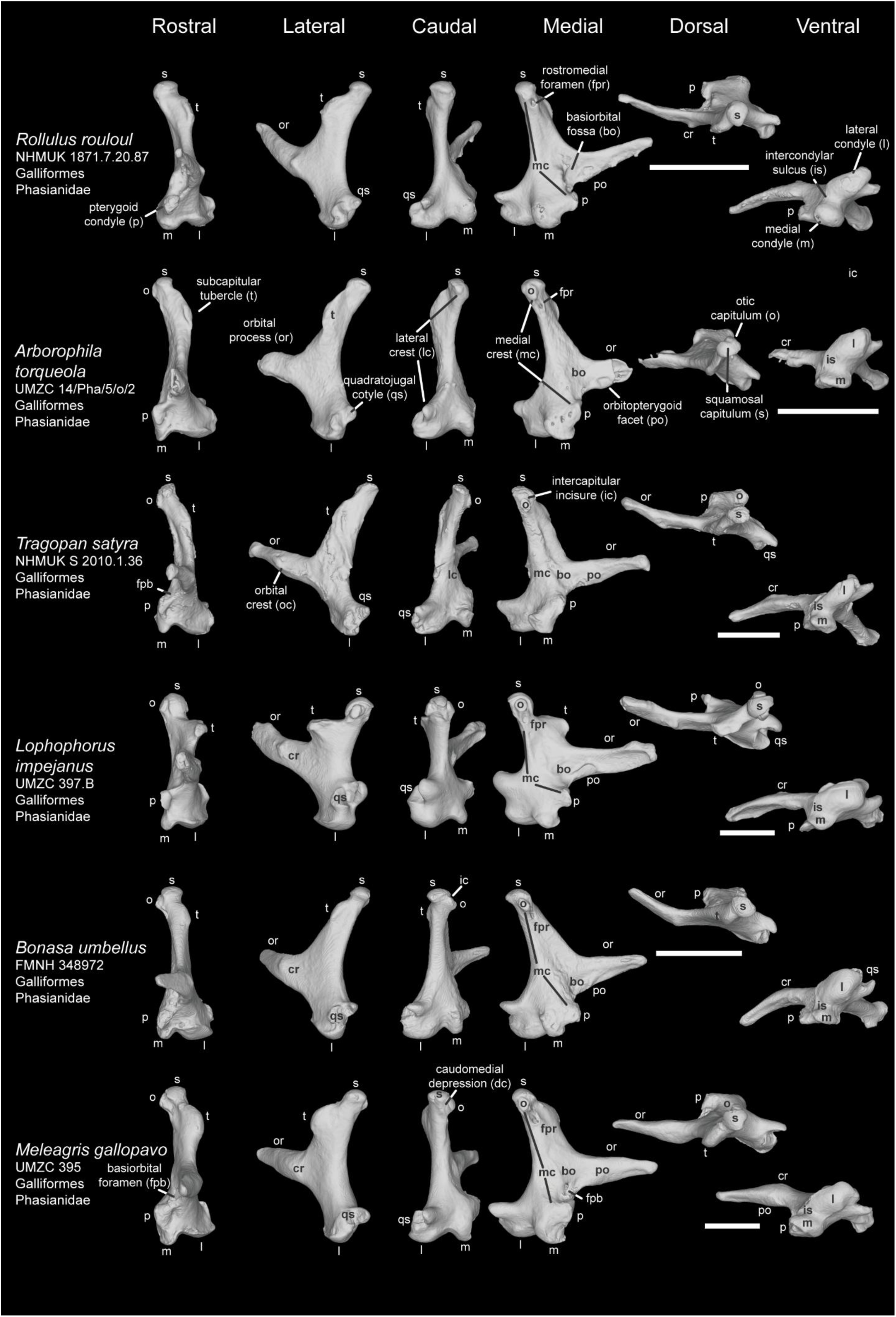
Comparison of avian quadrate among Galloanserae Ⅲ (Galliformes). Comparison of *Rollulus rouloul* (NHMUK 1871.7.20.87), *Arborophila torqueola* (UMZC 14/Pha/5/o/2), *Tragopan satyra* (NHMUK S 2010.1.36), *Lophophorus impejanus* (UMZC 397.B), *Bonasa umbellus* (FMNH 348972), and *Meleagris gallopavo* (UMZC 395) in rostral, lateral caudal, medial, dorsal, and ventral view. Scale bar: 5 mm.

**Plate 9.**
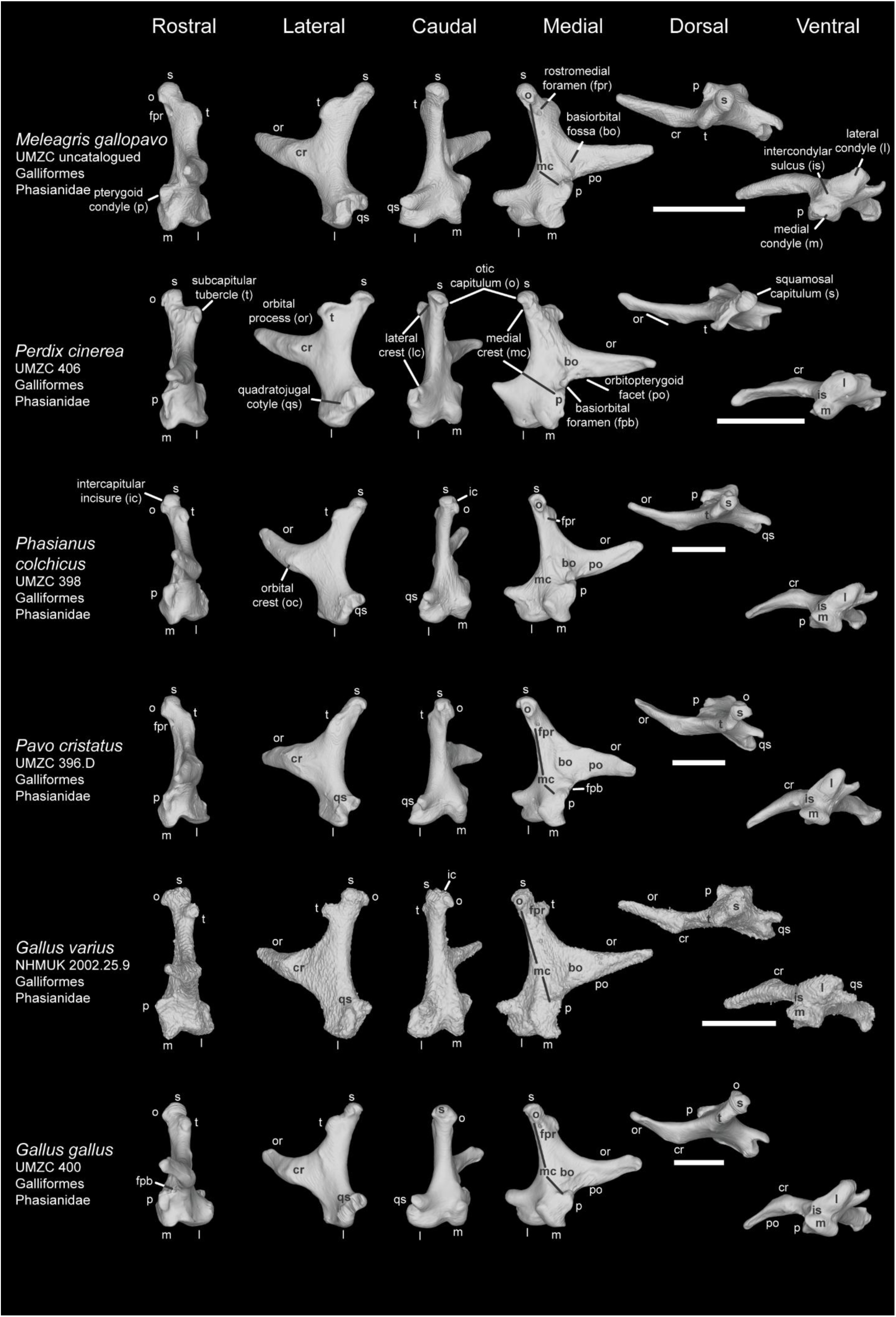
Comparison of avian quadrate among Galloanserae Ⅳ (Galliformes). Comparison of *Meleagris gallopavo* (UMZC uncatalogued), *Perdix cinerea* (UMZC 406), *Phasianus colchicus* (UMZC 398), *Pavo cristatus* (UMZC 396.D), *Gallus varius* (NHMUK 2002.25.9), and *G. gallus* (UMZC 400) in rostral, lateral caudal, medial, dorsal, and ventral view. Scale bar: 5 mm.

**Plate 10.**
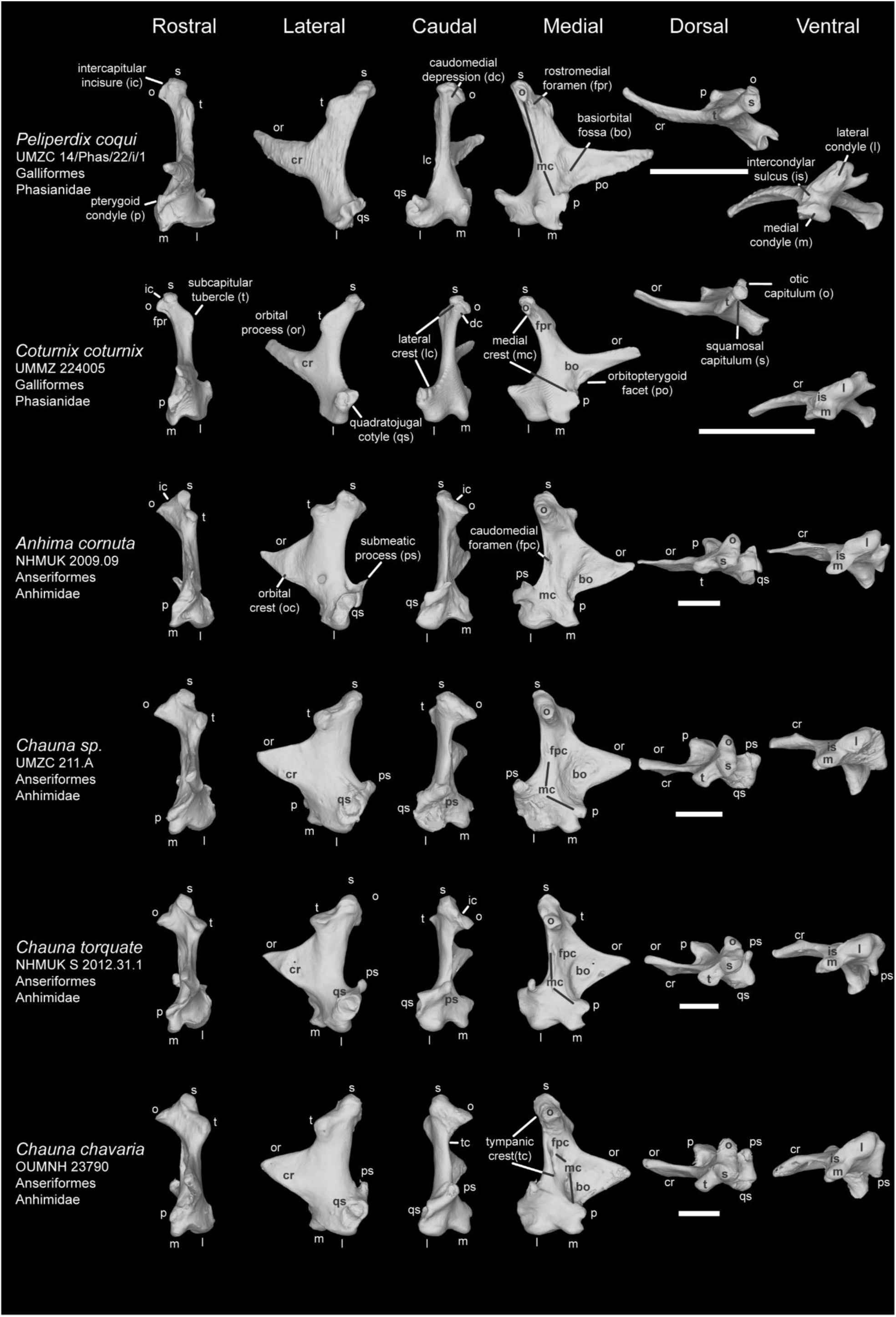
Comparison of avian quadrate among Galloanserae Ⅴ (Galliformes and Anseriformes). Comparison of *Peliperdix coqui* (UMZC 14/Phas/22/i/1), *Coturnix coturnix* (UMMZ 224005), *Anhima cornuta* (NHMUK 2009.2.2), *Chauna sp.* (UMZC 211.A), *Chauna torquata* (NHMUK S 2012.31.1), *and Chauna chavaria* (OUMNH 23790) in rostral, lateral caudal, medial, dorsal, and ventral view. Scale bar: 5 mm.

**Plate 11.**
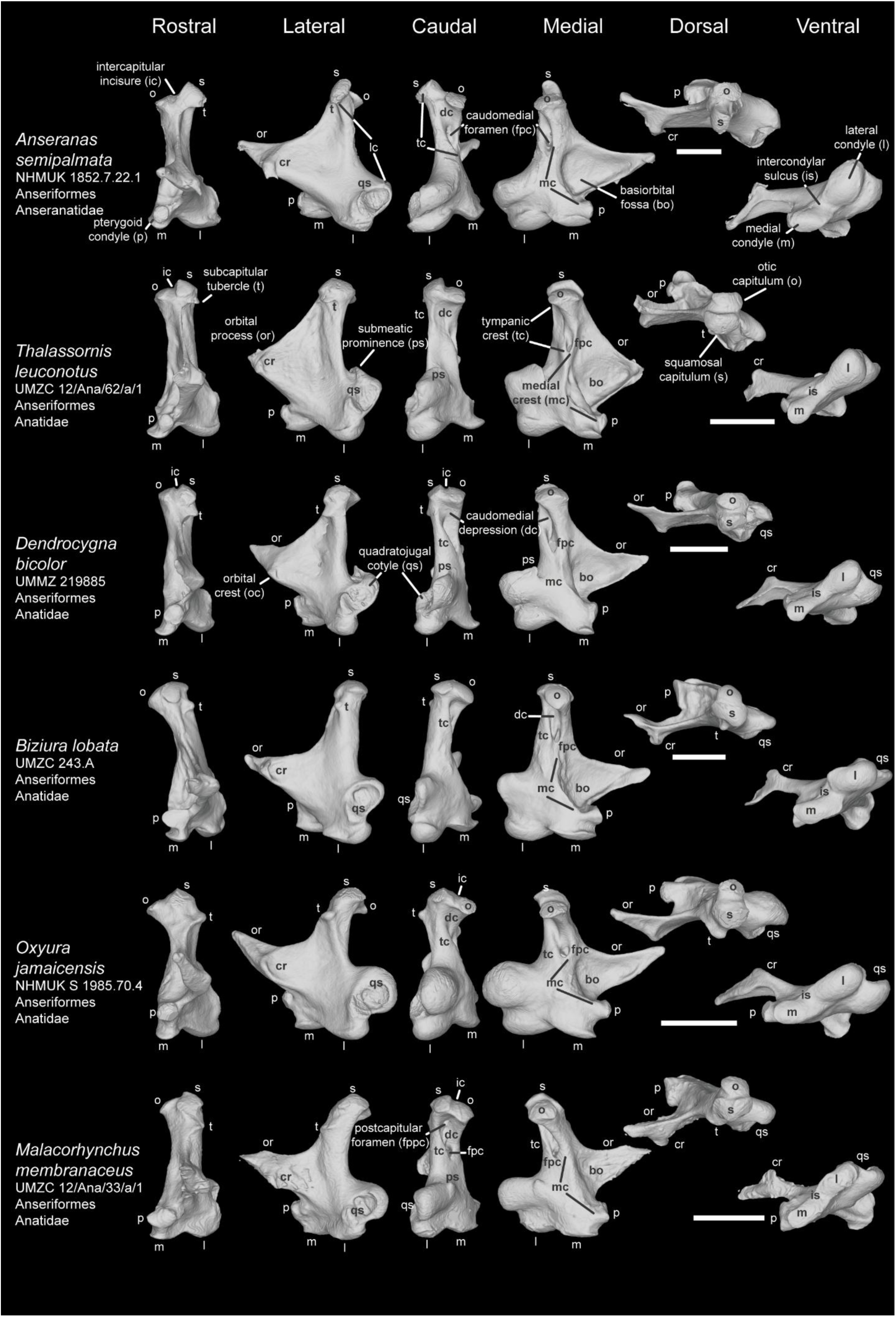
Comparison of avian quadrate among Galloanserae Ⅵ (Anseriformes). Comparison of *Anseranas semipalmata* (NHMUK 1852.7.22.1), *Thalassornis leuconotus* (UMZC 12/Ana/62/a/1), *Dendrocygna bicolor* (UMMZ 219885), *Biziura lobata* (UMZC 243.A), *Oxyura jamaicensis* (NHMUK S 1985.70.4), and *Malacorhynchus membranaceus* (UMZC 12/Ana/33/a/1) in rostral, lateral caudal, medial, dorsal, and ventral view. Scale bar: 5 mm.

**Plate 12.**
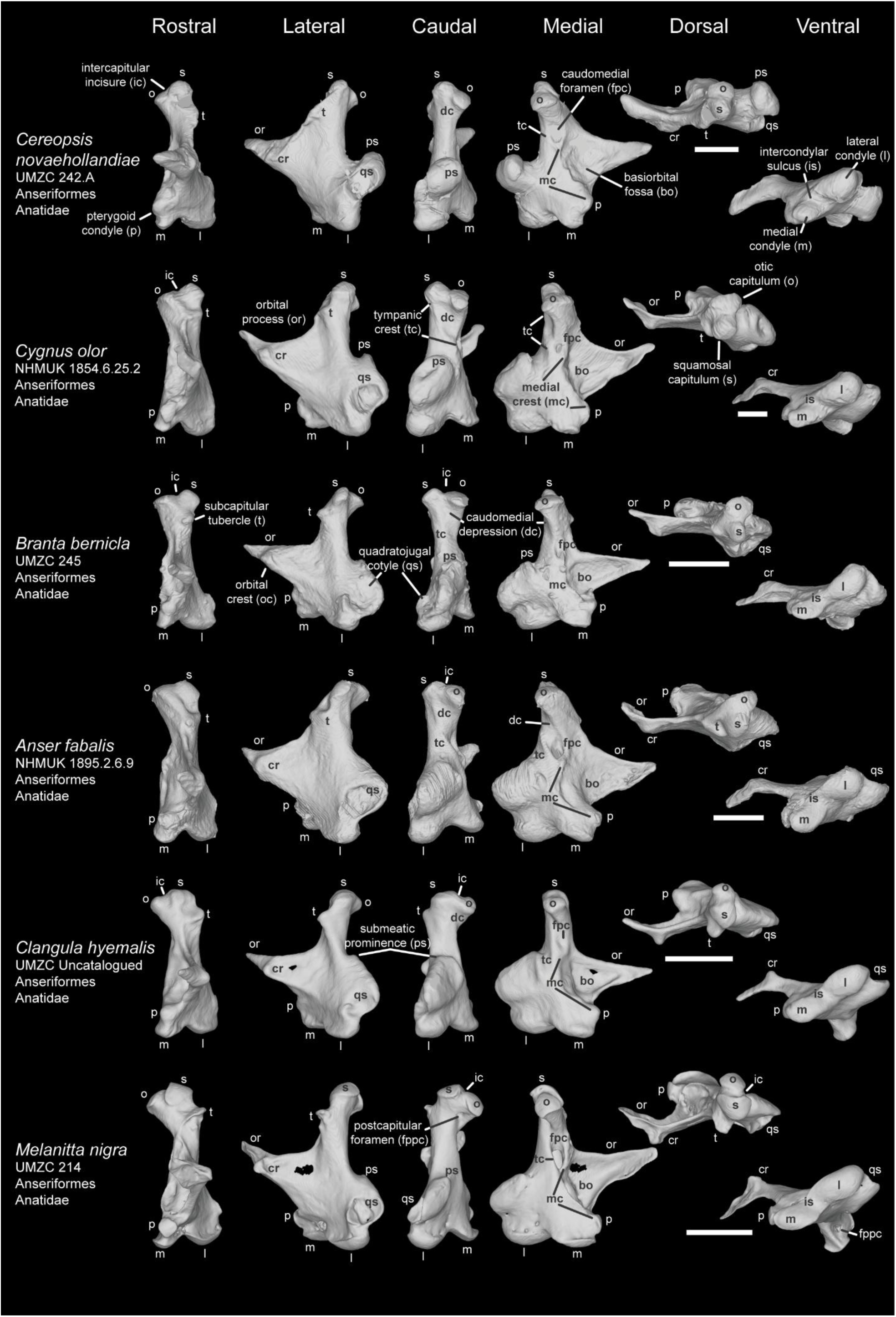
Comparison of avian quadrate among Galloanserae Ⅶ (Anseriformes). Comparison of *Cereopsis novaehollandiae* (UMZC 242), *Cygnus olor* (NHMUK 1854.6.25.2), *Branta bernicla* (UMZC 245), *Anser fabalis* (NHMUK 1895.2.6.9), *Clangula hyemalis* (UMZC uncatalogued), and *Melanitta nigra* (UMZC 214) in rostral, lateral caudal, medial, dorsal, and ventral view. Scale bar: 5 mm.

**Plate 13.**
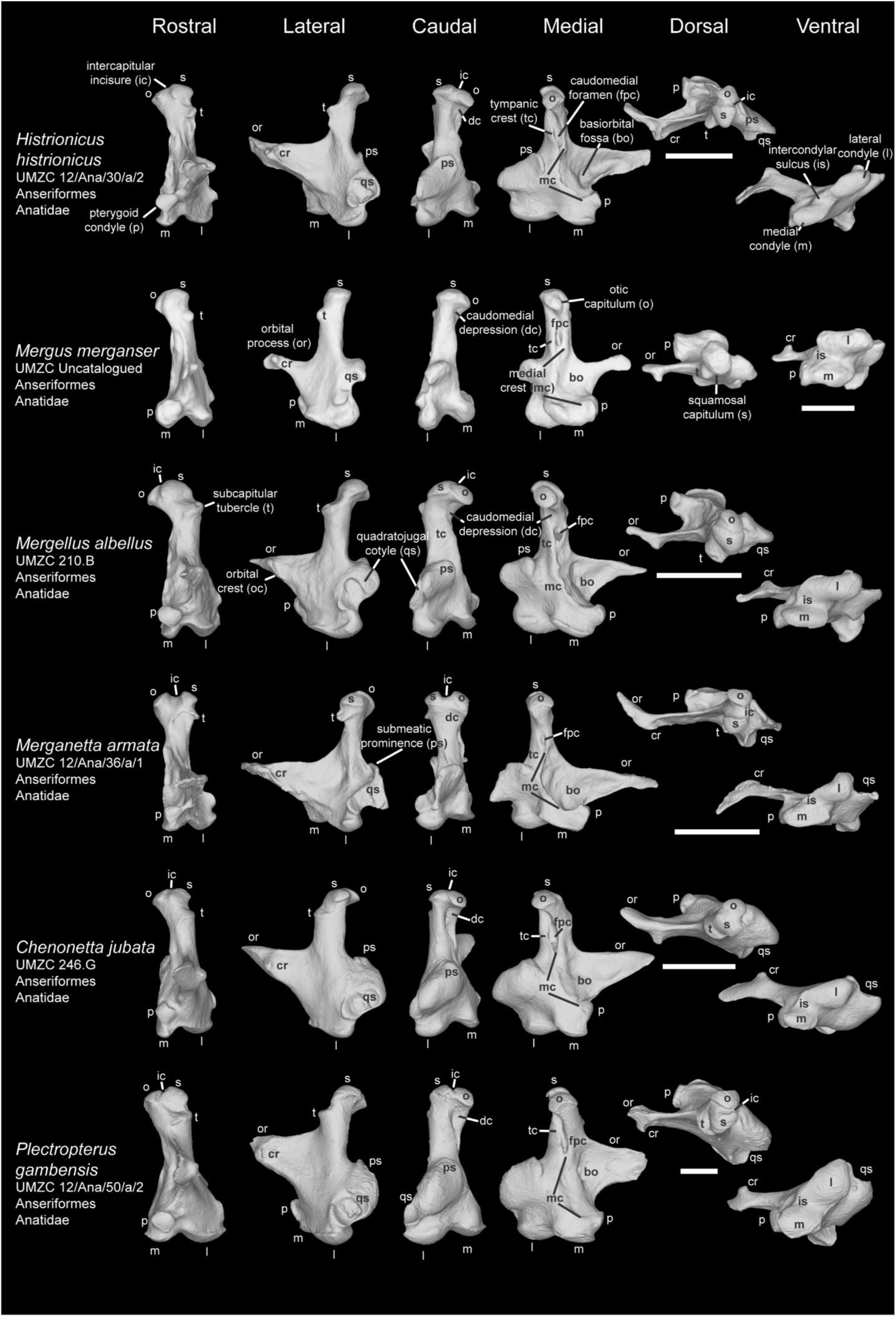
Comparison of avian quadrate among Galloanserae Ⅷ (Anseriformes). Comparison of *Histrionicus histrionicus* (UMZC 12/Ana/30/a/2), *Mergus merganser* (UMZC uncatalogued), *Mergellus albellus* (UMZC 210.B), *Merganetta armata* (UMZC 12/Ana/36/a/1), *Chenonetta jubata* (UMZC 246.G), and *Plectropterus gaubensis* (UMZC 12/Ana/50/a/2) in rostral, lateral caudal, medial, dorsal, and ventral view. Scale bar: 5 mm.

**Plate 14.**
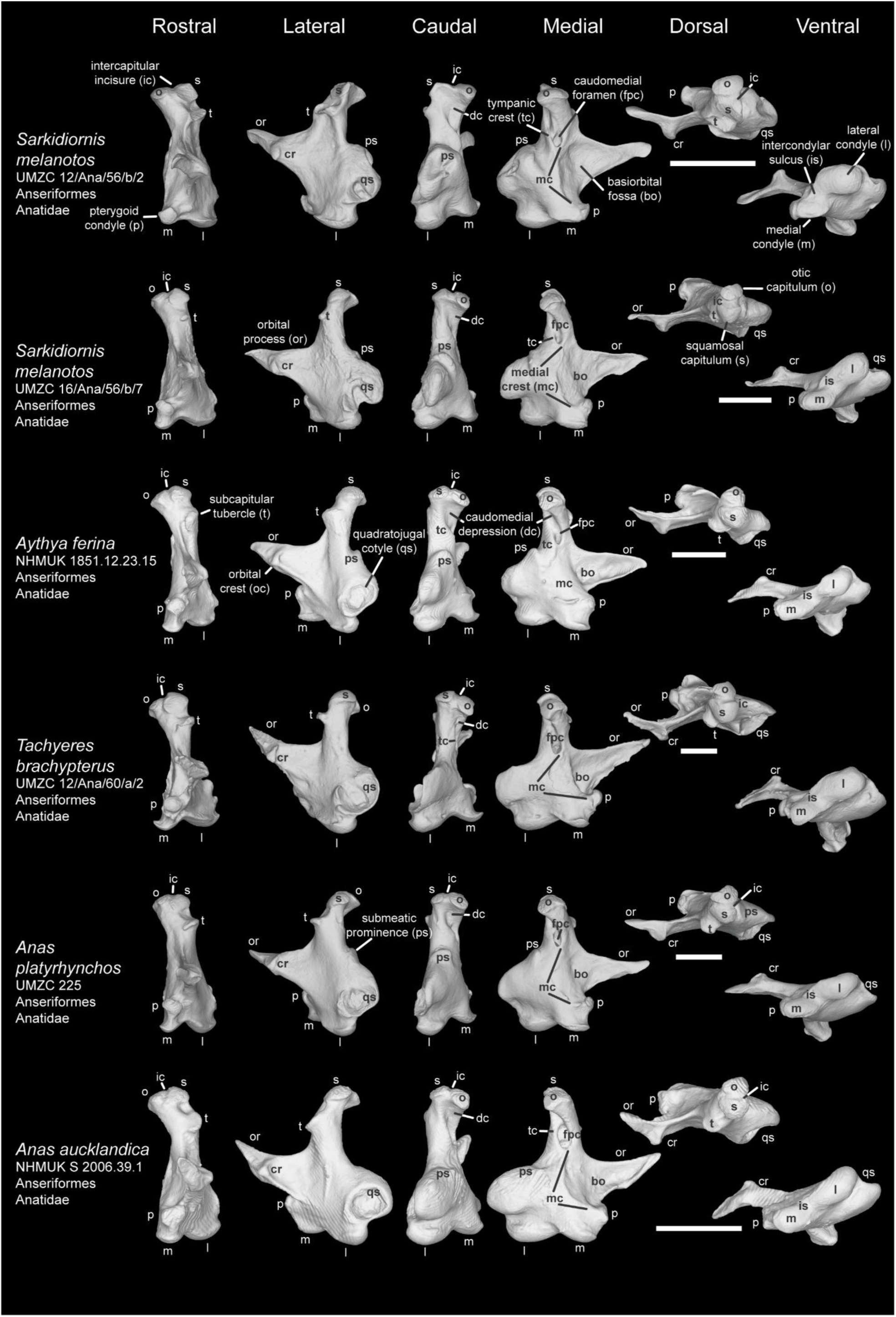
Comparison of avian quadrate among Galloanserae Ⅸ (Anseriformes). Comparison of *Sarkidiornis melanotos* (UMZC 12/Ana/56/b/2), *S. melanotos* (UMZC 16/Ana/56/b/7), *Aythya ferina* (NHMUK 1851.12.23.15), *Tachyeres brachypterus* (UMZC 12/Ana/60/a/2), *Anas platyrhynchos* (UMZC 225), and *A. aucklandica* (NHMUK S 2006.39.1) in rostral, lateral caudal, medial, dorsal, and ventral view. Scale bar: 5 mm.

**Plate 15.**
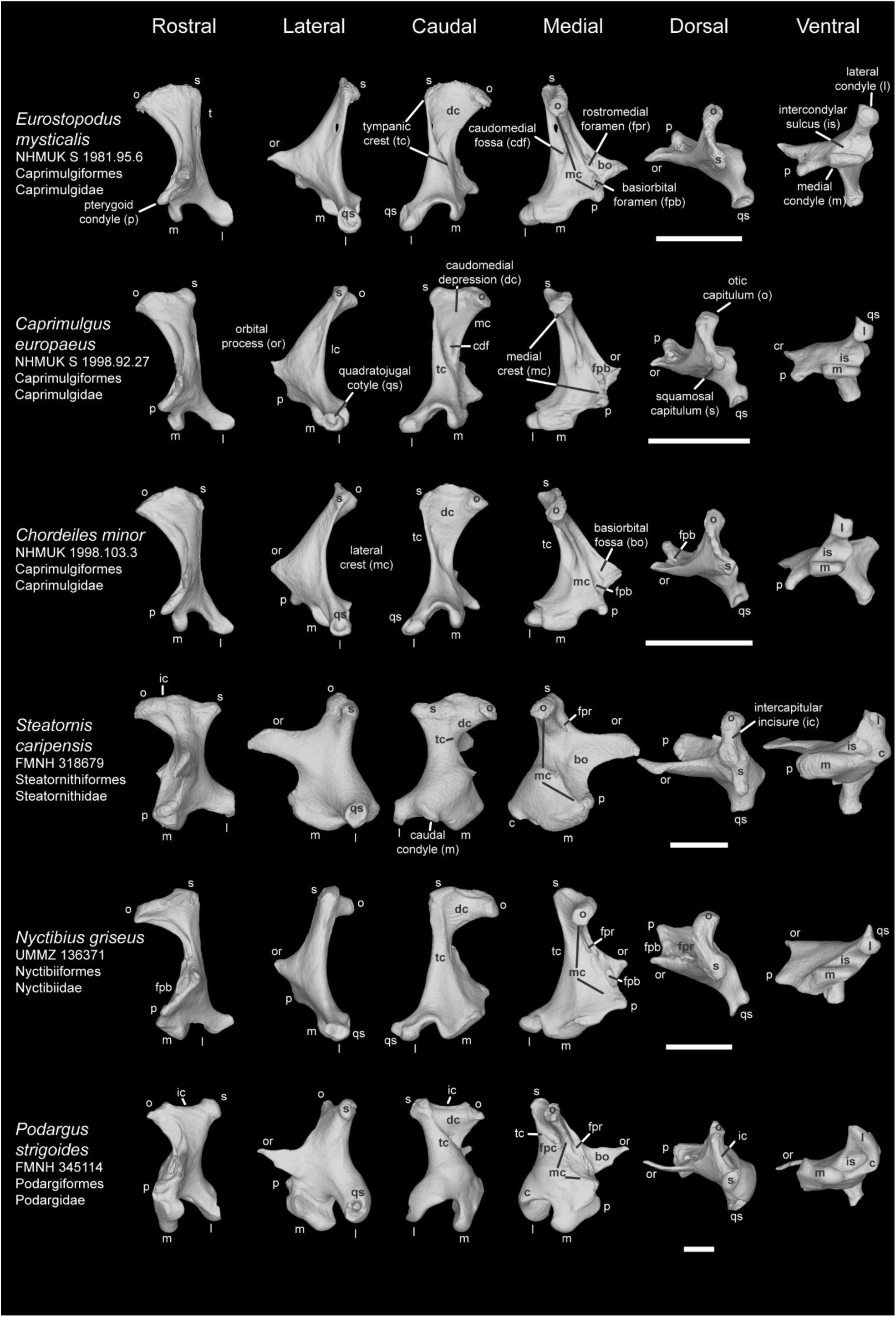
Comparison of avian quadrate among StrisoresⅠ. Comparison of *Eurostopodus mysticalis* (NHMUK S 1981.95.6), *Caprimulgus europaeus* (NHMUK S 1998.92.27), *Chordeiles minor* (NHMUK 1998.103.3), *Steatornis caripensis* (FMNH 318679), *Nyctibius griseus* (UMMZ 136371), and *Podargus strigoides* (FMNH 345114) in rostral, lateral caudal, medial, dorsal, and ventral view. Scale bar: 5 mm.

**Plate 16.**
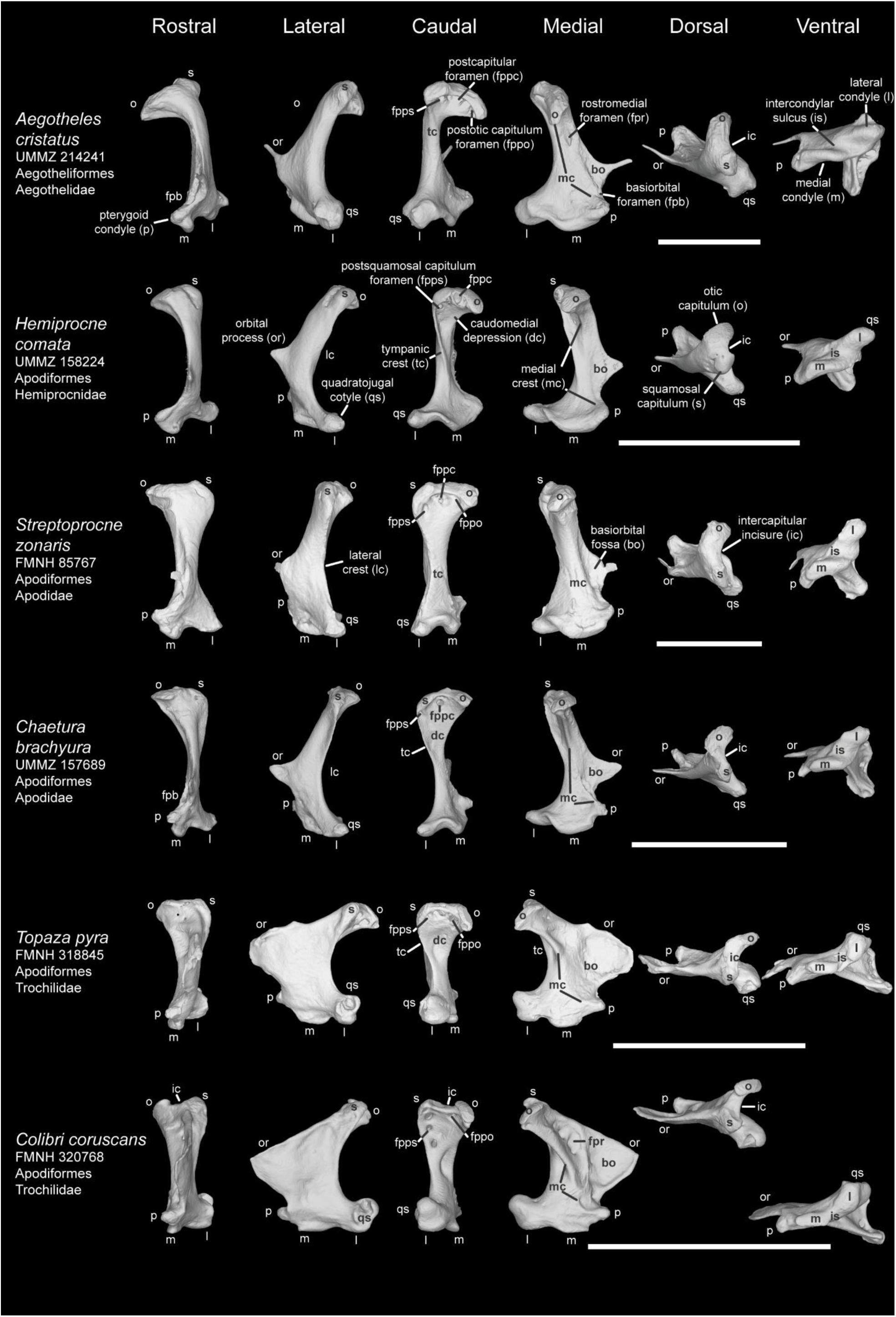
Comparison of avian quadrate among Strisores Ⅱ. Comparison of *Aegotheles cristatus* (UMMZ 214241), *Hemiprocne comata* (UMMZ 158224), *Streptoprocne zonaris* (FMNH 85767), *Chaetura brachyura* (UMMZ 157689), *Topaza pyra* (FMNH 318845), and *Colibri coruscans* (FMNH 320768) in rostral, lateral caudal, medial, dorsal, and ventral view. Scale bar: 5 mm.

**Plate 17.**
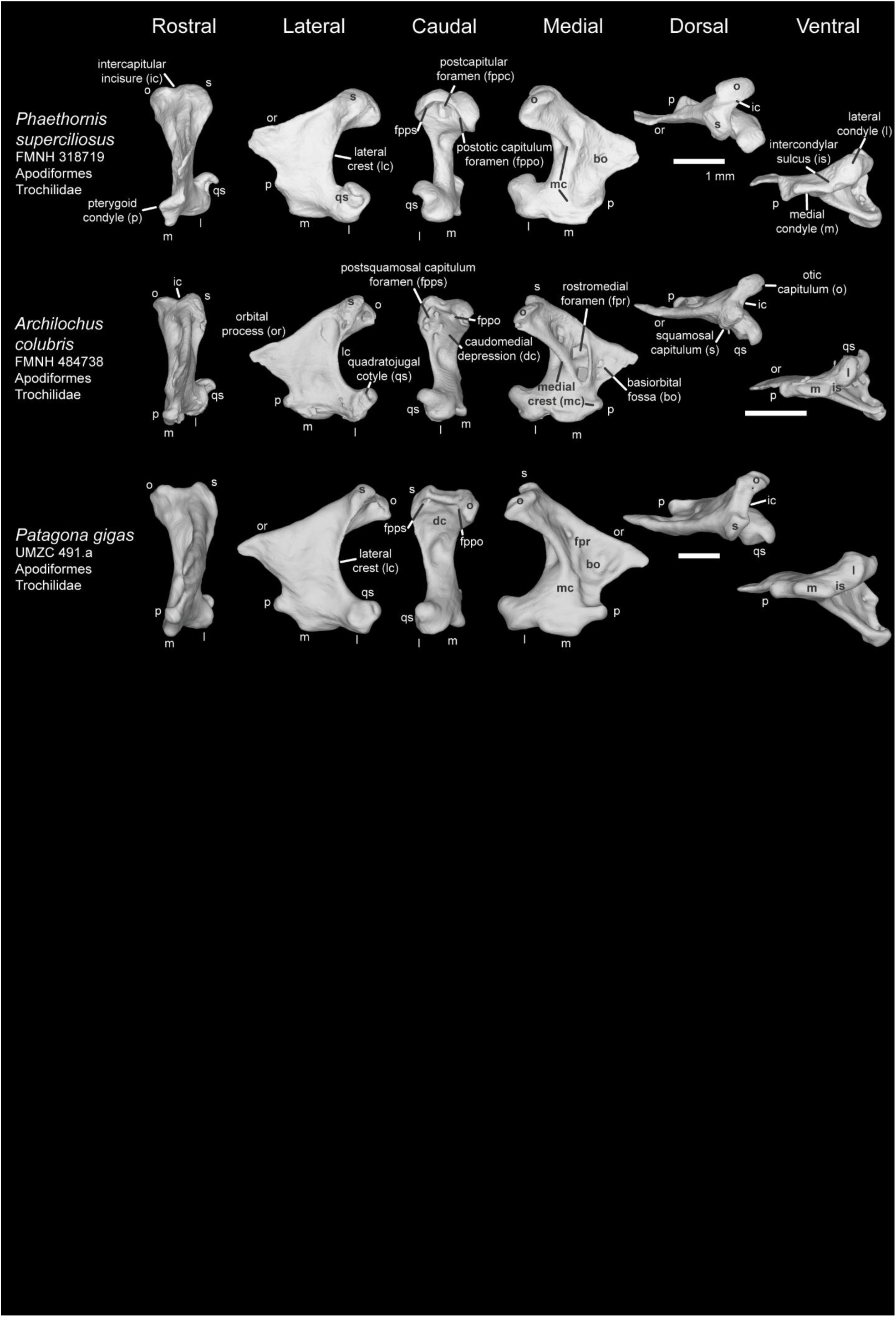
Comparison of avian quadrate among Strisores Ⅲ. Comparison of *Phaethornis superciliosus* (FMNH 318719), *Archilochus colubris* (FMNH 484738), and *Patagona gigas* (UMZC 491.a) in rostral, lateral caudal, medial, dorsal, and ventral view. Scale bar: 5 mm. (Except for *Phaethornis superciliosus*)

**Plate 18.**
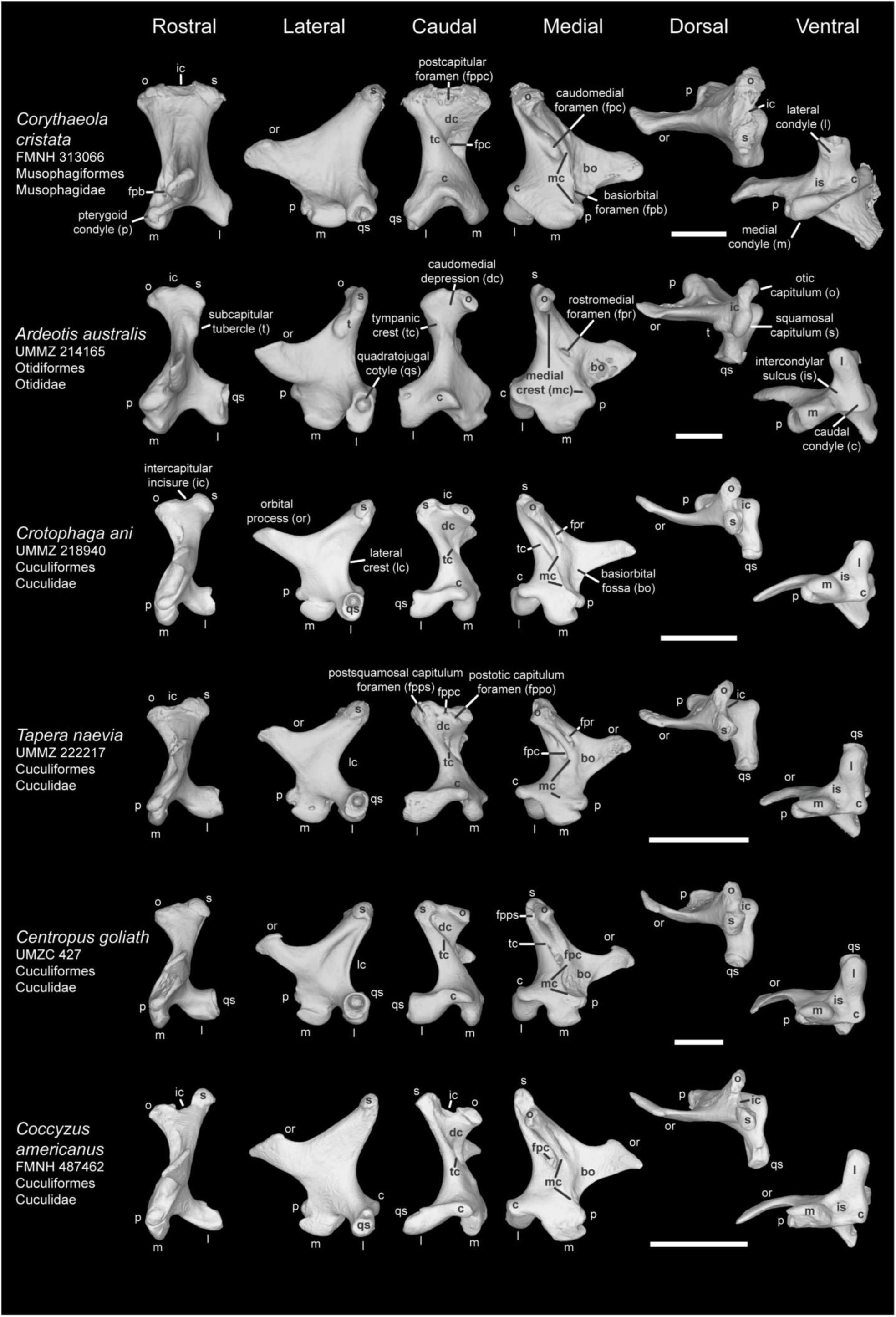
Comparison of avian quadrate among Columbaves Ⅰ. Comparison of *Corythaeola cristata* (FMNH 347273), *Ardeotis australis* (UMMZ 214165), *Crotophaga ani* (UMMZ 218940), *Tapera naevia* (UMMZ 214165), *Centropus goliath* (UMZC 427), and *Coccyzus americanus* (FMNH 487462) in rostral, lateral caudal, medial, dorsal, and ventral view. Scale bar: 5 mm.

**Plate 19.**
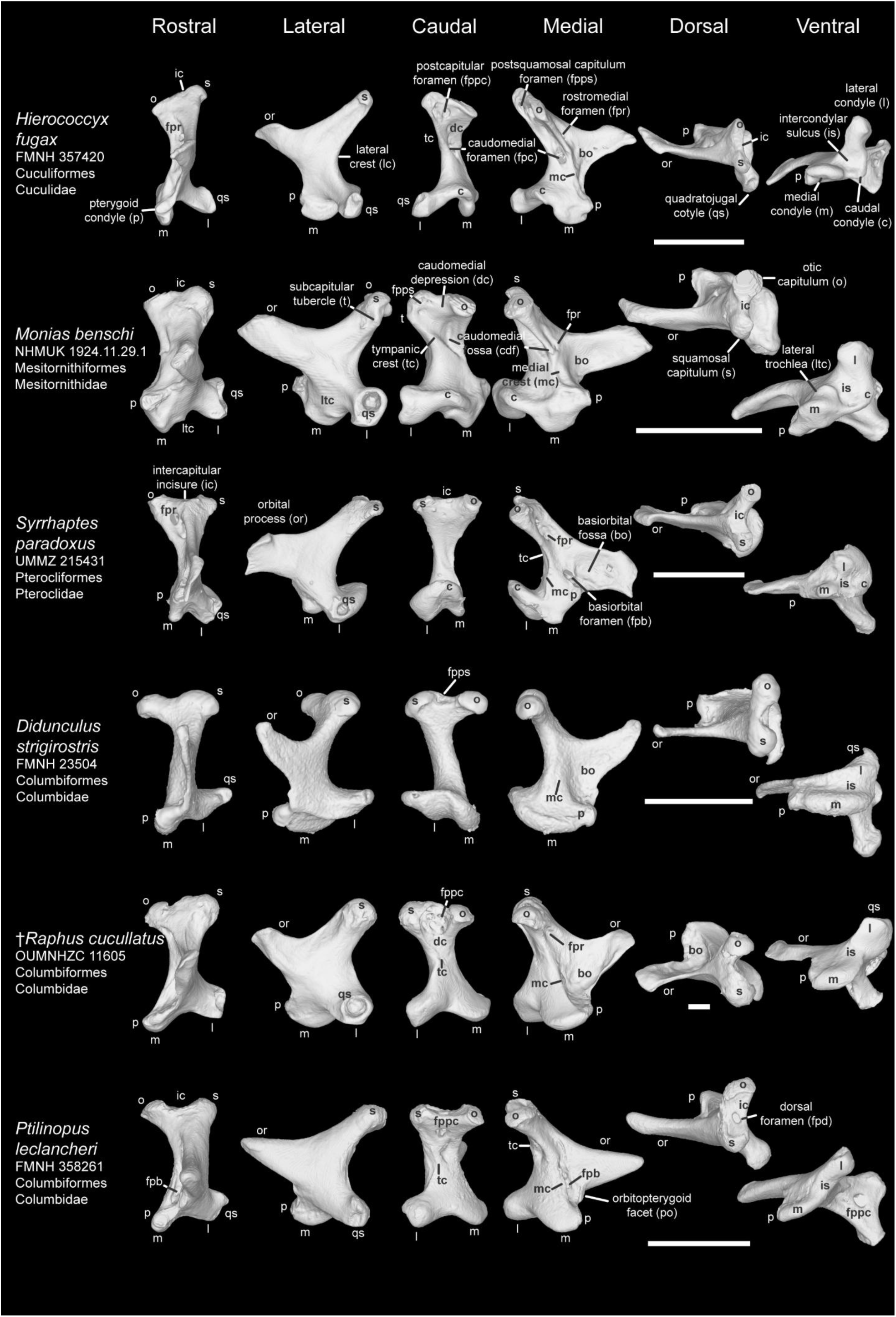
Comparison of avian quadrate among Columbaves Ⅱ. Comparison of *Hierococcyx fugax* (FMNH 357420), *Monias benschi* (NHMUK 1924.11.29.1), *Syrrhaptes paradoxus* (UMMZ 215431), *Raphus cucullatus* (OUMNHZC 11605), *Ptilinopus leclancheri* (FMNH 358261), and *Treron capellei* (UMMZ 220454) in rostral, lateral caudal, medial, dorsal, and ventral view. Scale bar: 5 mm.

**Plate 20.**
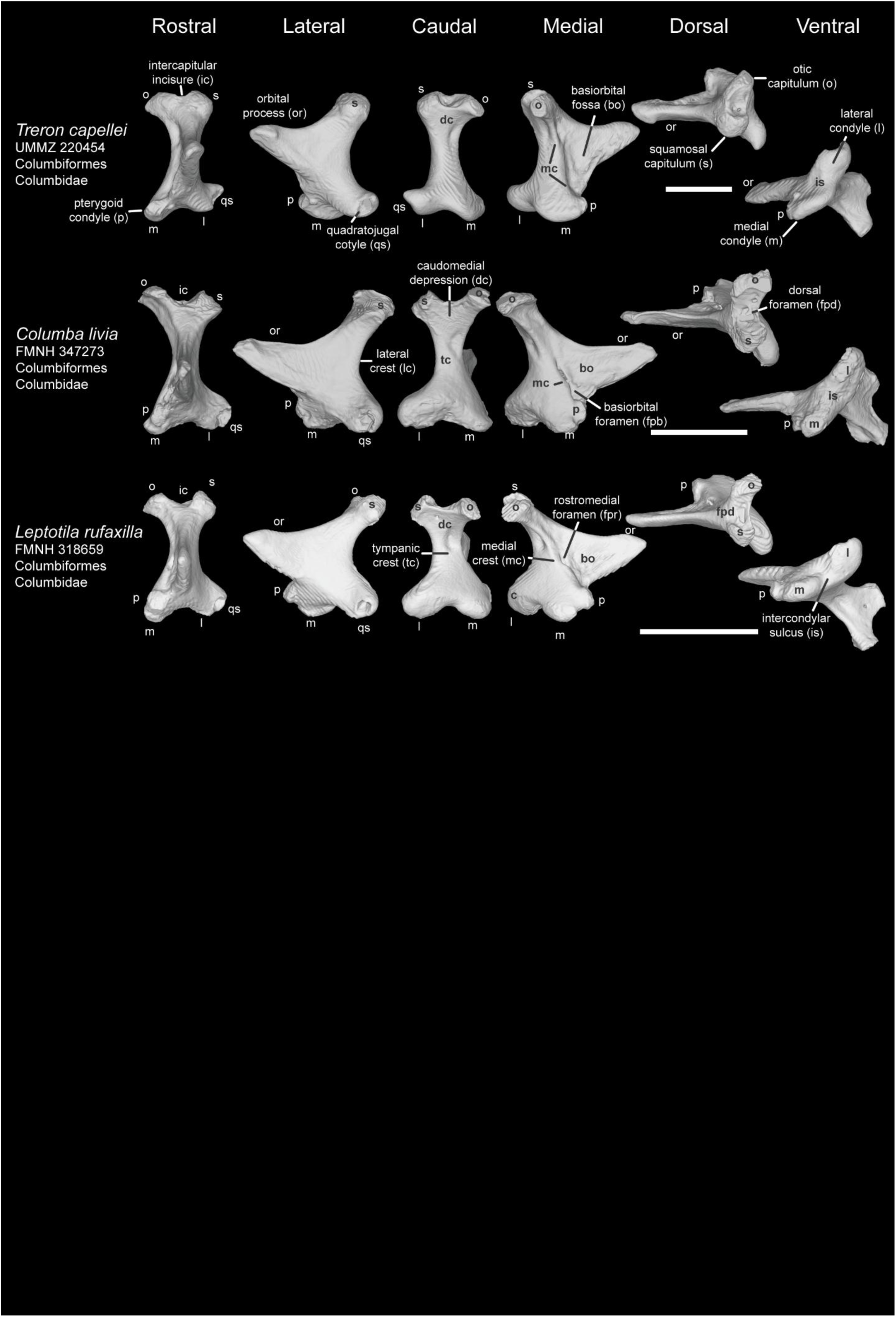
Comparison of avian quadrate among Columbaves Ⅲ. Comparison of *Columba livia* (FMNH 347273) and *Leptotila rufaxilla* (FMNH 318659) in rostral, lateral caudal, medial, dorsal, and ventral view. Scale bar: 5 mm.

**Plate 21.**
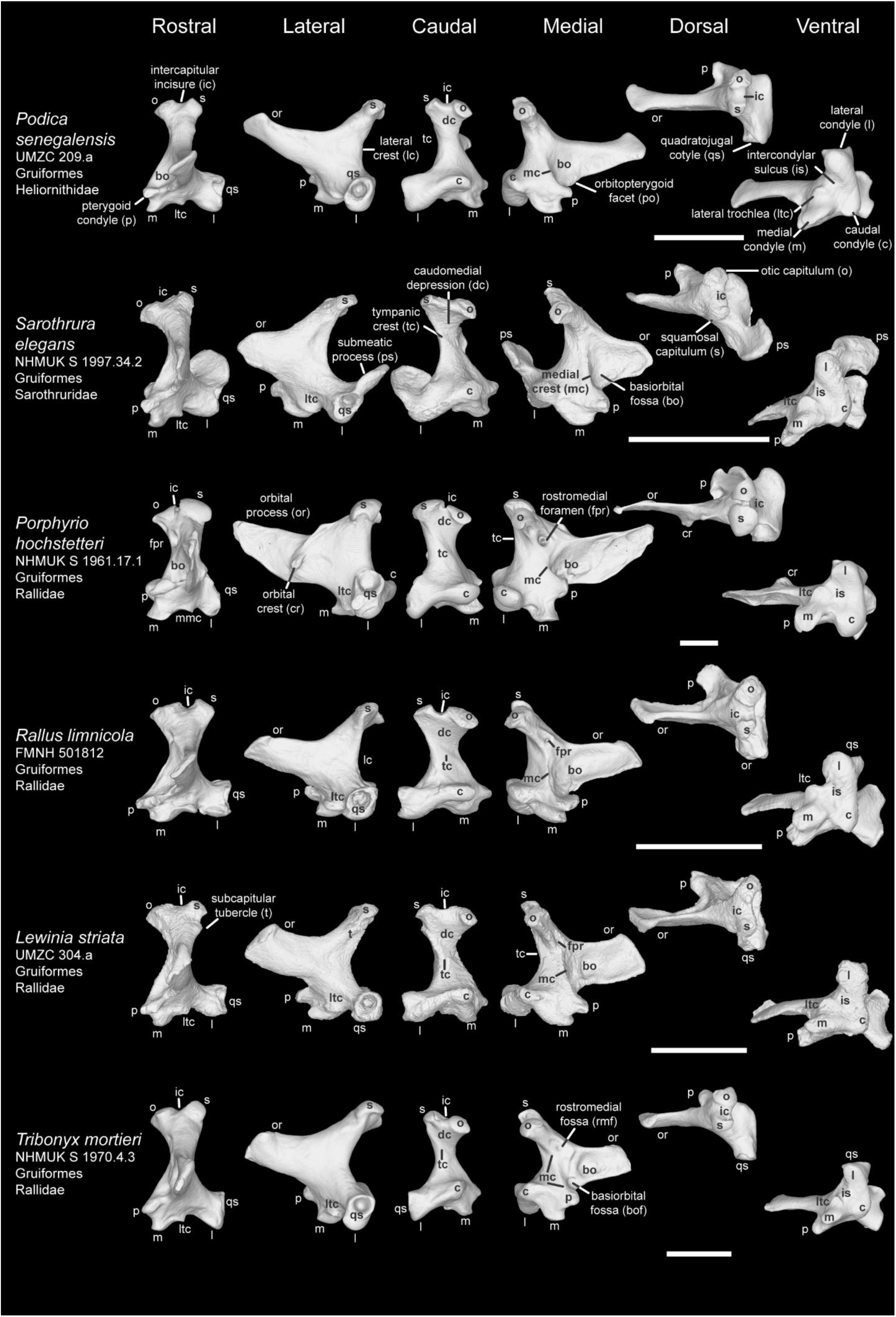
Comparison of avian quadrate among Gruiformes Ⅰ. Comparison of *Podica senegalensis* (UMZC 209.a), *Sarothrura elegans* (NHMUK S 1997.34.2), *Porphyrio hochstetteri* (NHMUK S 1961.17.1), *Rallus limnicola* (FMNH 501812), *Lewinia striata* (UMZC 304.a), and *Tribonyx mortierii* (NHMUK S 1970.4.3) in rostral, lateral caudal, medial, dorsal, and ventral view. Scale bar: 5 mm.

**Plate 22.**
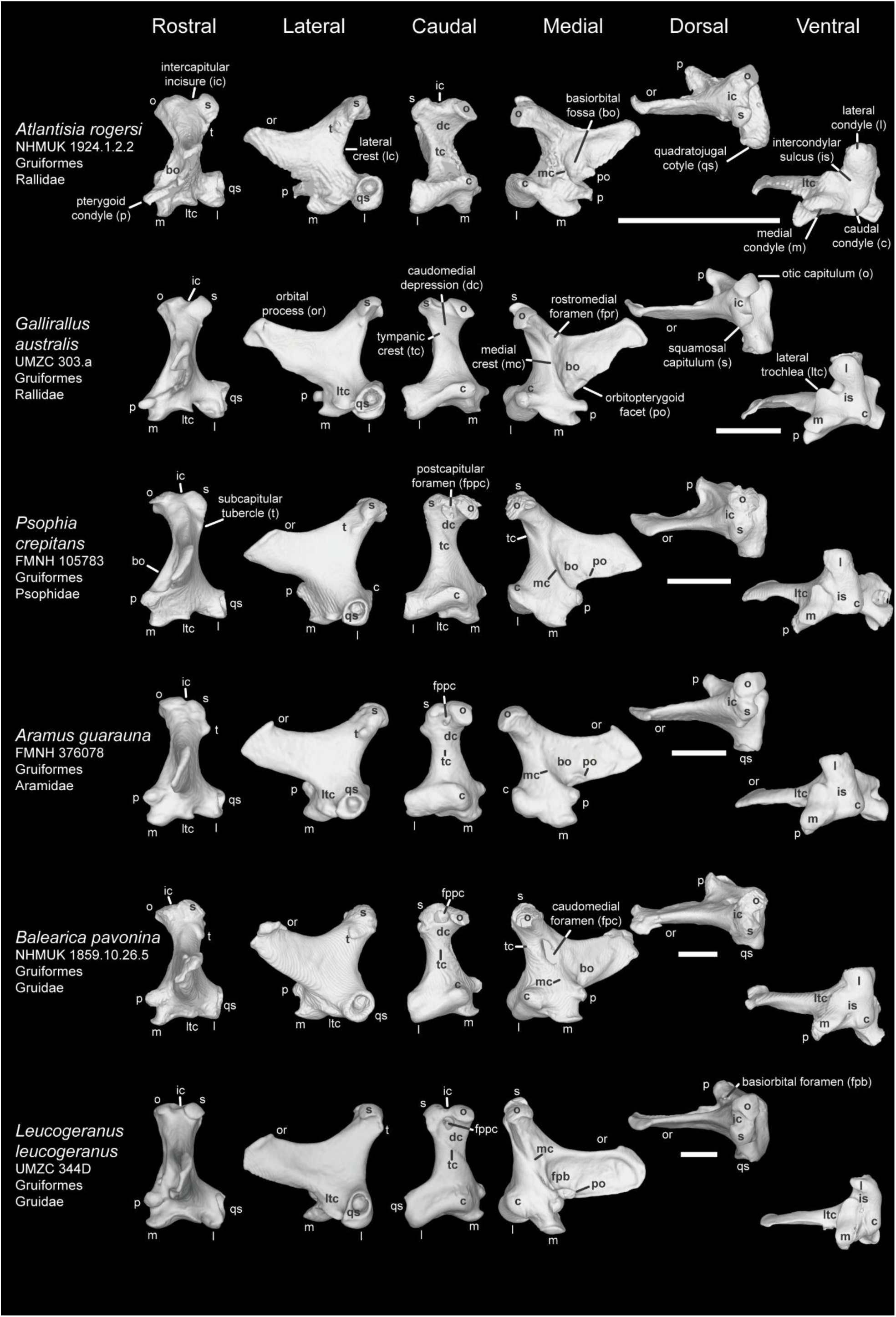
Comparison of avian quadrate among Gruiformes Ⅱ. Comparison of *Atlantisia rogersi* (NHMUK 1924.1.2.2), *Gallirallus australis* (UMZC 303.a), *Psophia crepitans* (FMNH 105783), *Aramus guarauna* (FMNH 376078), *Balearica pavonina* (NHMUK 1859.10.26.5), and *Leucogeranus leucogeranus* (UMZC 344D) in rostral, lateral caudal, medial, dorsal, and ventral view. Scale bar: 5 mm.

**Plate 23.**
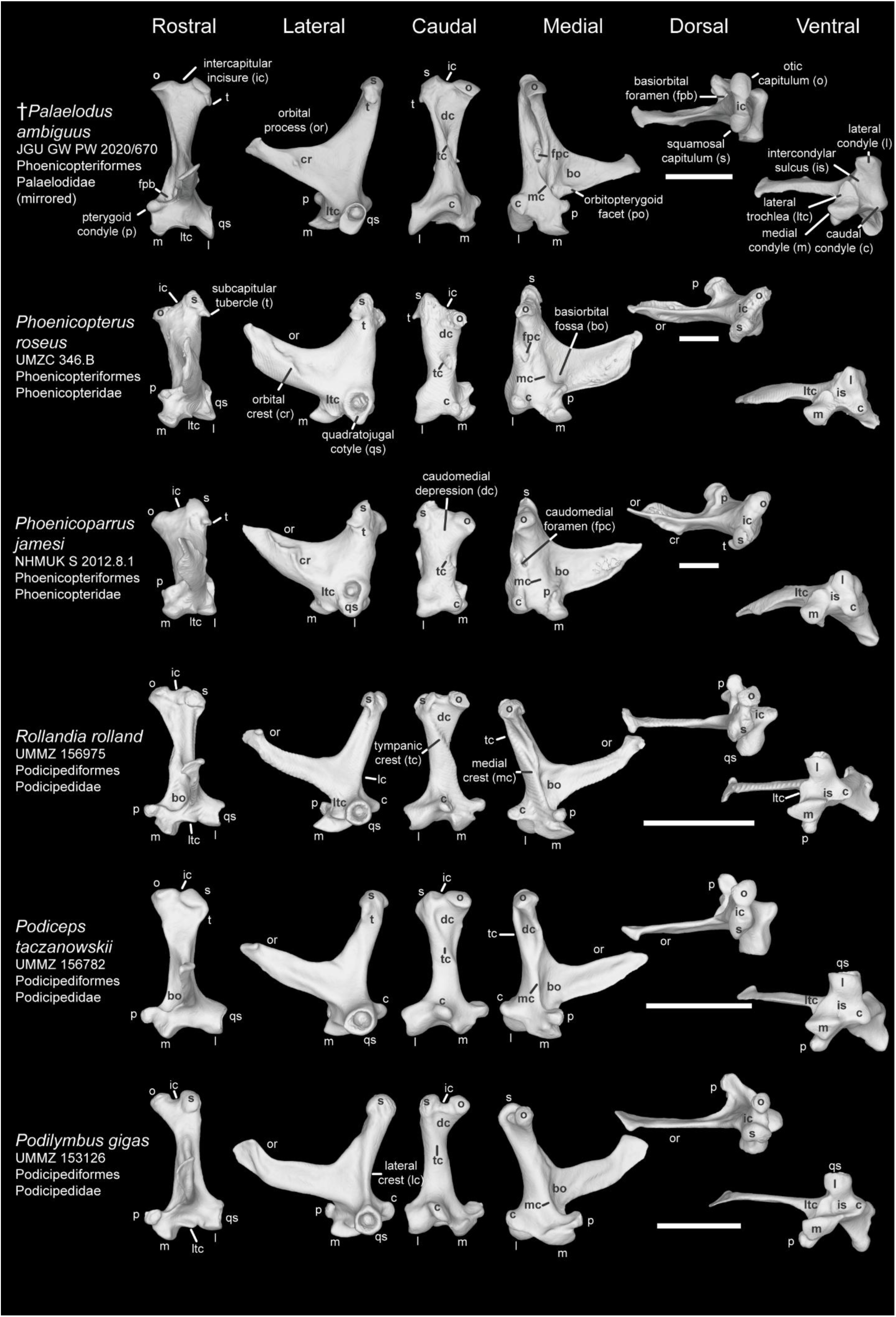
Comparison of avian quadrate among Aequorlitornithes Ⅰ (Phoenicopteriformes and Podicipediformes). Comparison of *Palaelodus ambiguus* (JGU GW PW 2020/670), *Phoenicopterus roseus* (UMZC 346.B), *Phoenicoparrus jamesi* (NHMUK S 2012.8.1), *Rollandia rolland* (UMMZ 156975), *Podiceps taczanowskii* (UMMZ 156782), and *Podilymbus gigas* (UMMZ 153126) in rostral, lateral caudal, medial, dorsal, and ventral view. Scale bar: 5 mm.

**Plate 24.**
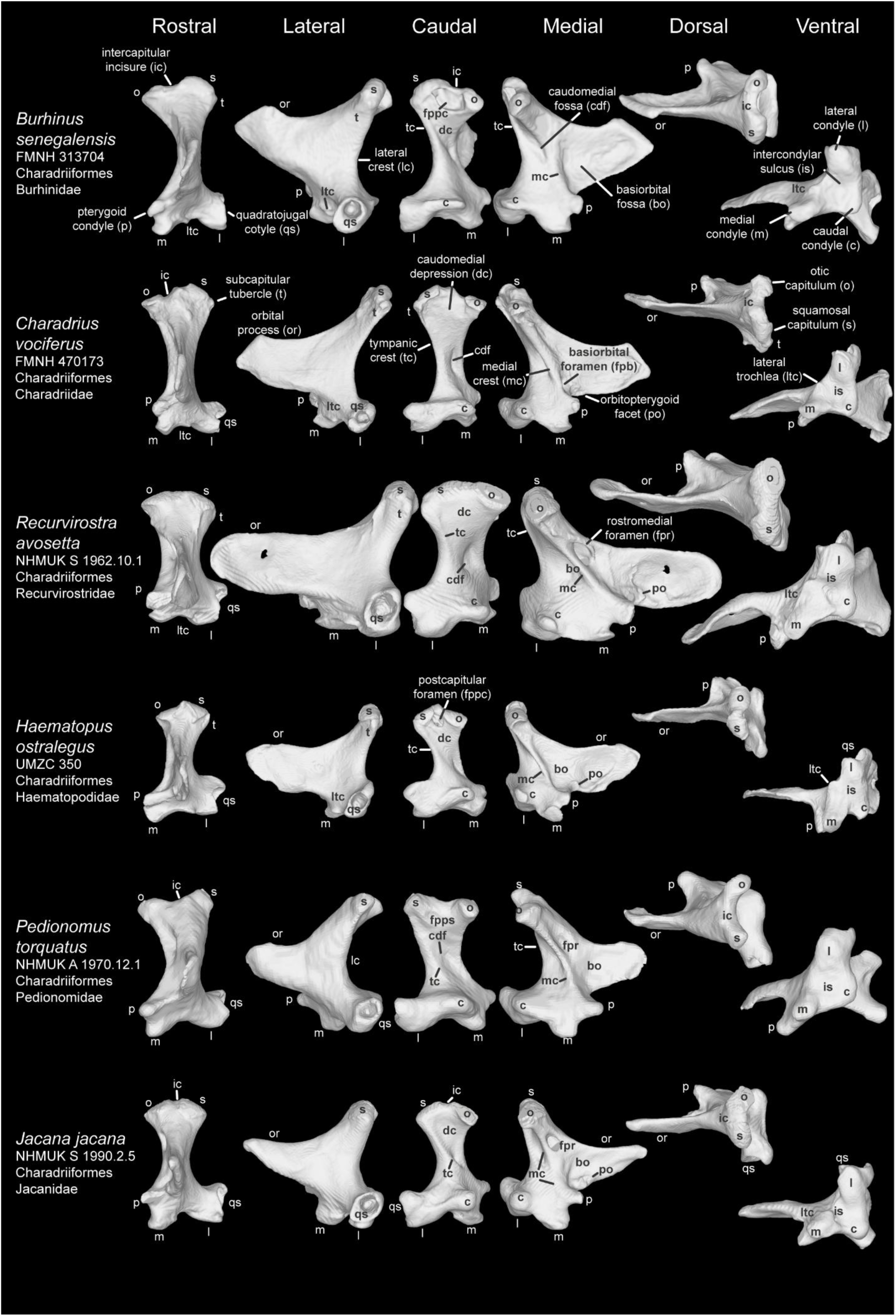
Comparison of avian quadrate among Aequorlitornithes Ⅱ (Charadriiformes). Comparison of *Burhinus senegalensis* (FMNH 313704), *Charadrius vociferus* (FMNH 470173), *Recurvirostra avosetta* (NHMUK S 1962.10.1), *Haematopus ostralegus* (UMZC 350), *Pedionomus torquatus* (NHMUK A 1970.12.1), and *Jacana jacana* (NHMUK S 1990.2.5) in rostral, lateral caudal, medial, dorsal, and ventral view. Scale bar: 5 mm.

**Plate 25.**
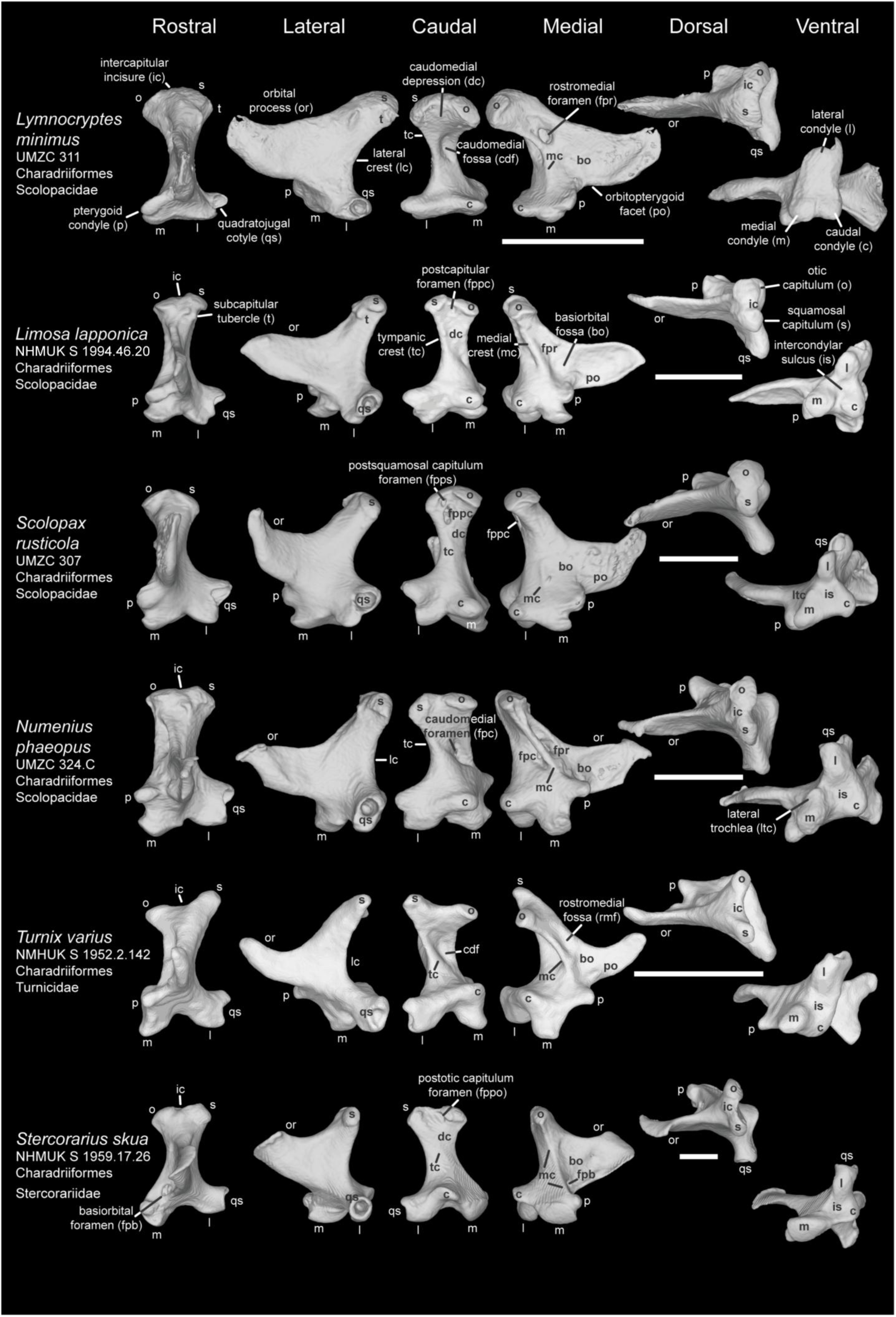
Comparison of avian quadrate among Aequorlitornithes Ⅲ (Charadriiformes). Comparison of *Lymnocryptes minimus* (UMZC 311), *Limosa lapponica* (NHMUK S 1994.46.20), *Scolopax rusticola* (UMZC 307), *Numenius phaeopus* (UMZC 324.C), *Turnix varius* (NMHUK S 1952.2.142), and *Stercorarius skua* (NHMUK S 1959.17.26) in rostral, lateral caudal, medial, dorsal, and ventral view. Scale bar: 5 mm.

**Plate 26.**
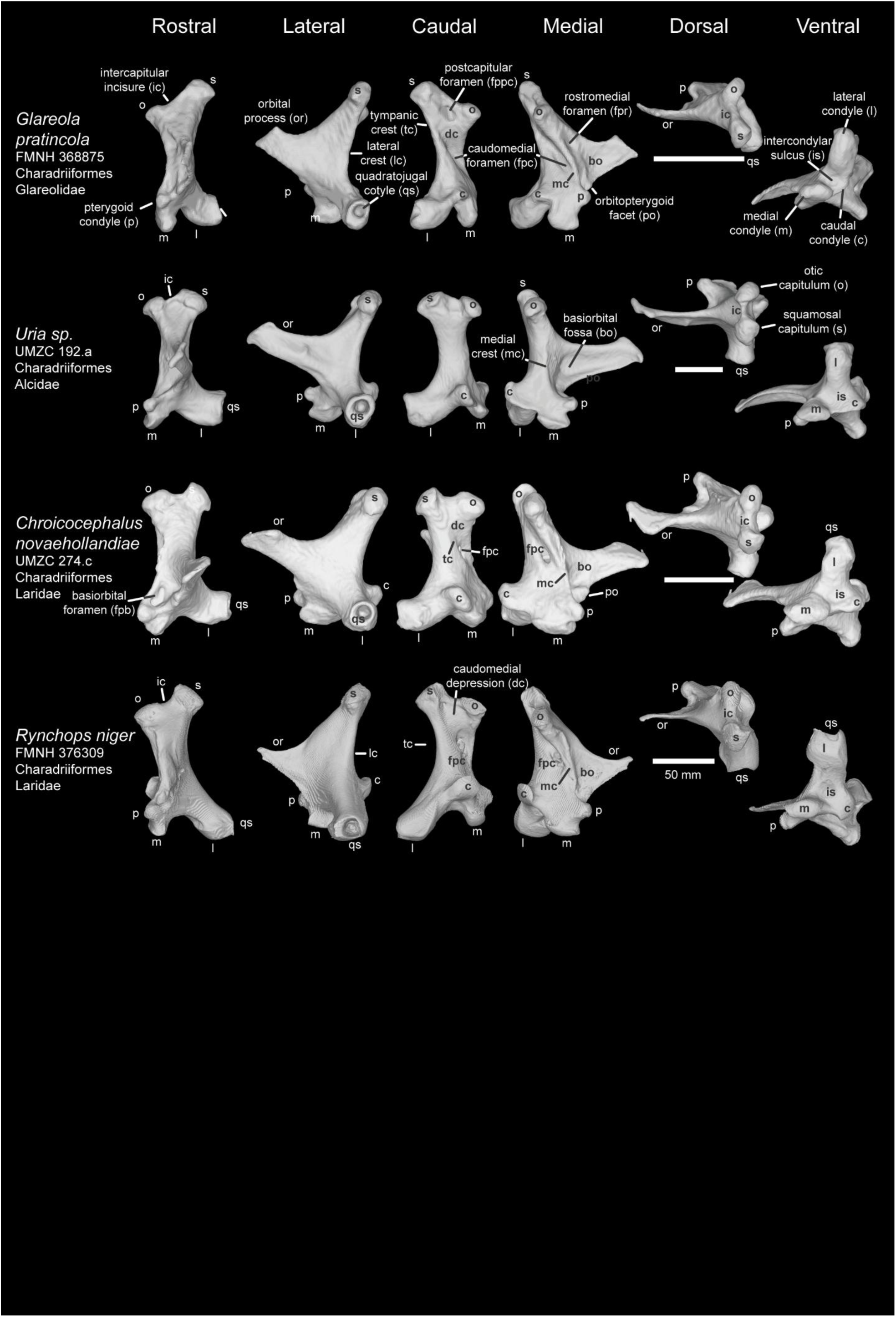
Comparison of avian quadrate among Aequorlitornithes Ⅳ (Charadriiformes). Comparison of *Glareola pratincola* (FMNH 368875), *Uria sp.* (UMZC 192.a), *Chroicocephalus novaehollandiae* (UMZC 274.c), and *Rynchops niger* (FMNH 376309) in rostral, lateral caudal, medial, dorsal, and ventral view. Scale bar: 5 mm. (Except for *Rynchops niger*).

**Plate 27.**
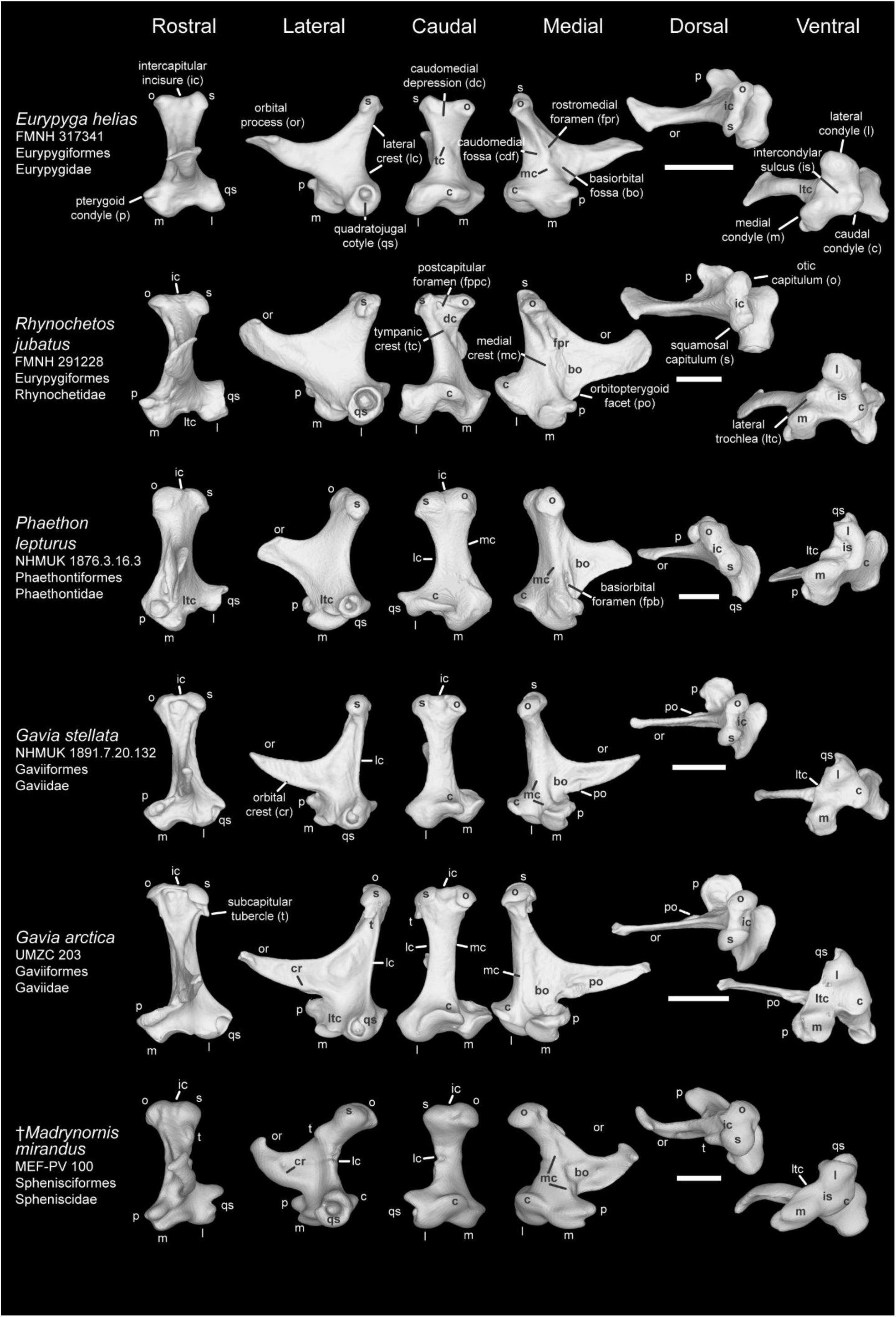
Comparison of avian quadrate among Aequorlitornithes Ⅴ. Comparison of *Eurypyga helias* (FMNH 317341), *Rhynochetos jubatus* (FMNH 291228), *Phaethon lepturus* (NHMUK 1876.3.16.3), *Gavia stellata* (NHMUK 1891.7.20.132), *G. arctica* (UMZC 203), and *Madrynornis mirandus* (MEF-PV 100) in rostral, lateral caudal, medial, dorsal, and ventral view. Scale bar: 5 mm.

**Plate 28.**
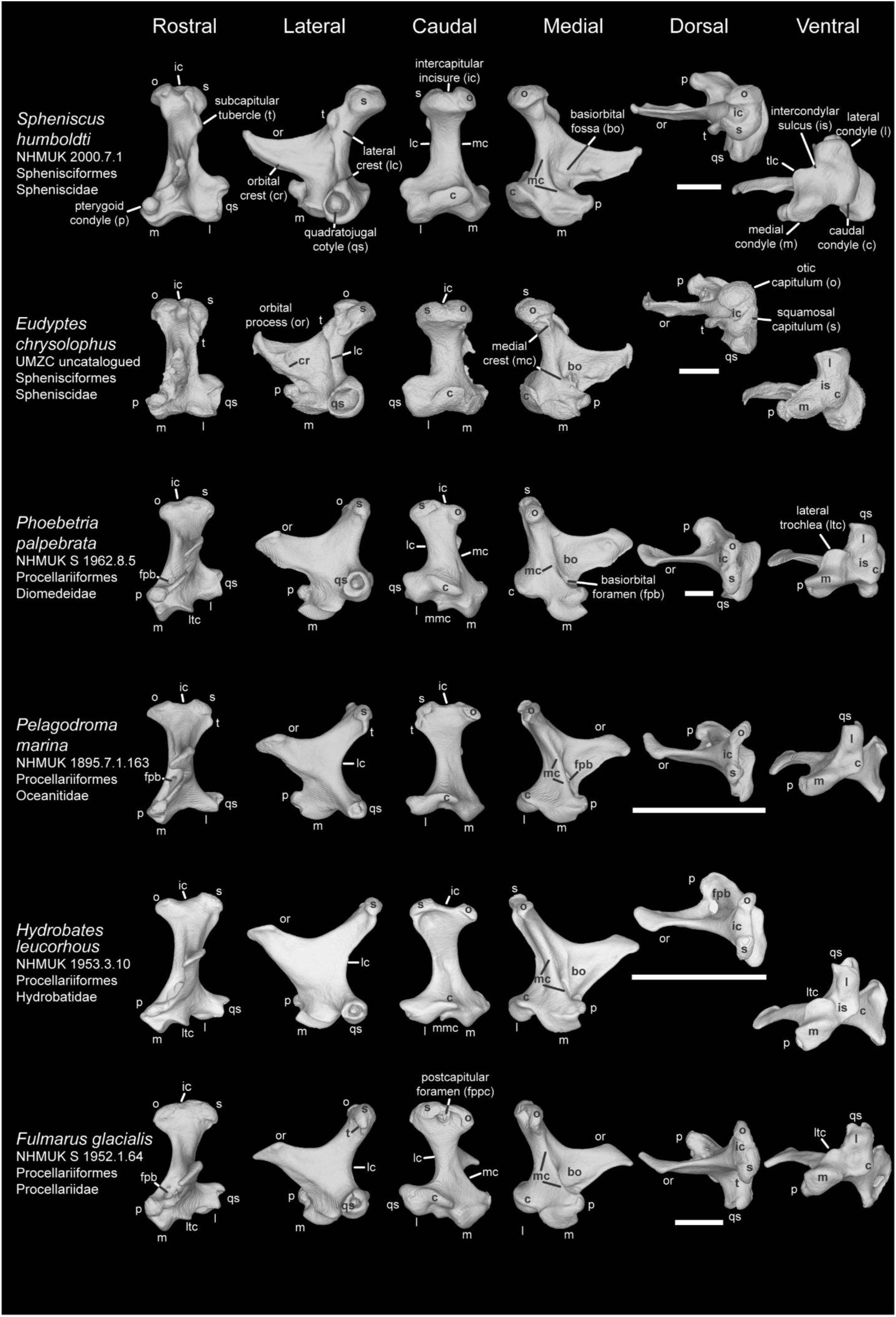
Comparison of avian quadrate among Aequorlitornithes Ⅵ (Sphenisciformes and Procellariiformes). Comparison of *Spheniscus humboldti* (NHMUK 2000.7.1), *Eudyptes chrysolophus* (UMZC uncatalogued), *Phoebetria palpebrata* (NHMUK S 1962.8.5), *Pelagodroma marina* (NHMUK 1895.7.1.163), *Hydrobates leucorhous* (NHMUK 1953.3.10), and *Fulmarus glacialis* (NHMUK S 1952.1.64) in rostral, lateral caudal, medial, dorsal, and ventral view. Scale bar: 5 mm.

**Plate 29.**
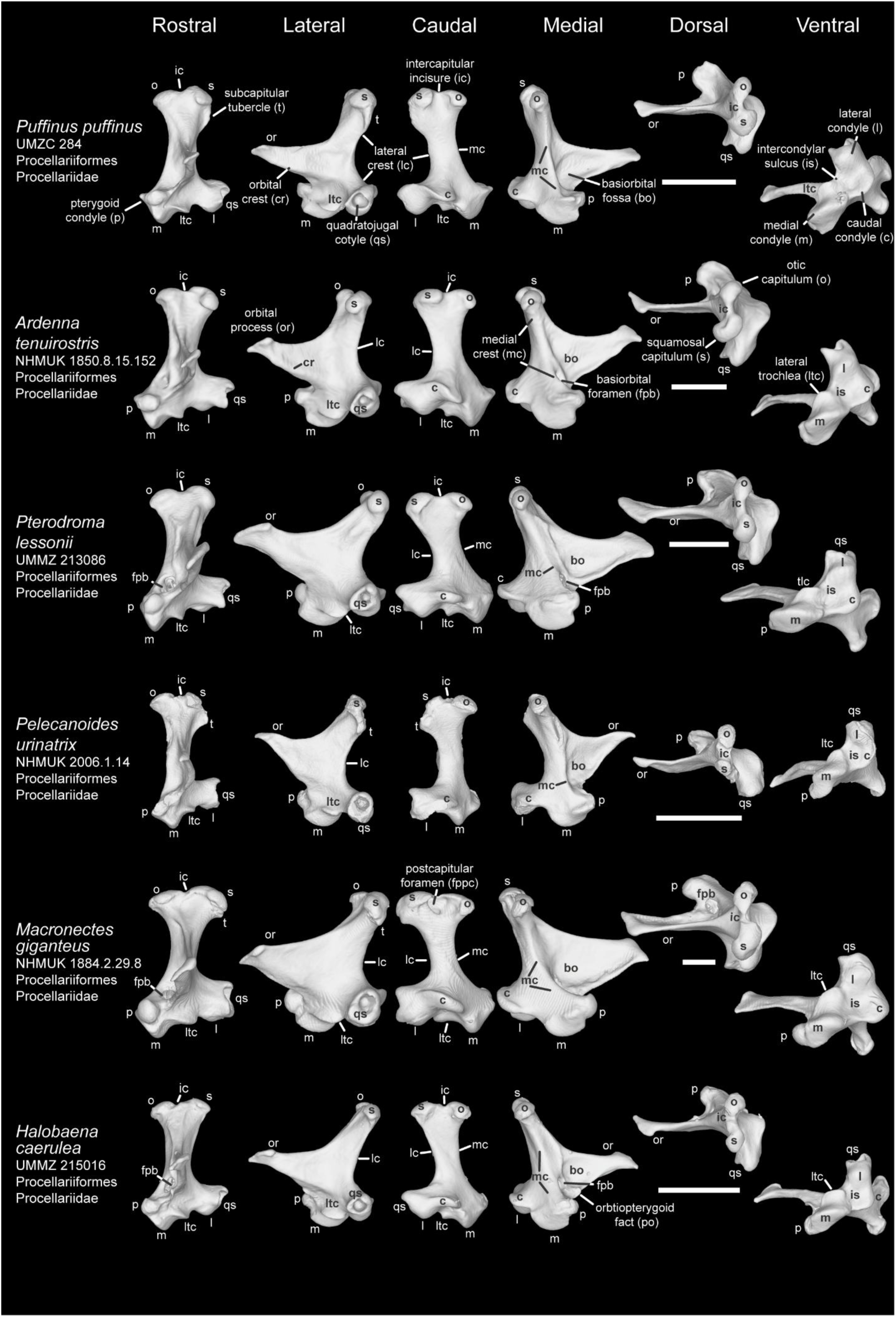
Comparison of avian quadrate among Aequorlitornithes Ⅶ (Procellariiformes). Comparison of *Puffinus puffinus* (UMZC 284), *Ardenna tenuirostris* (NHMUK 1850.8.15.152), *Pterodroma lessonii* (UMMZ 213086), *Pelecanoides urinatrix* (NHMUK 2006.1.14), *Macronectes giganteus* (NHMUK 1884.2.29.8), and *Halobaena caerulea* (UMMZ 215016) in rostral, lateral caudal, medial, dorsal, and ventral view. Scale bar: 5 mm.

**Plate 30.**
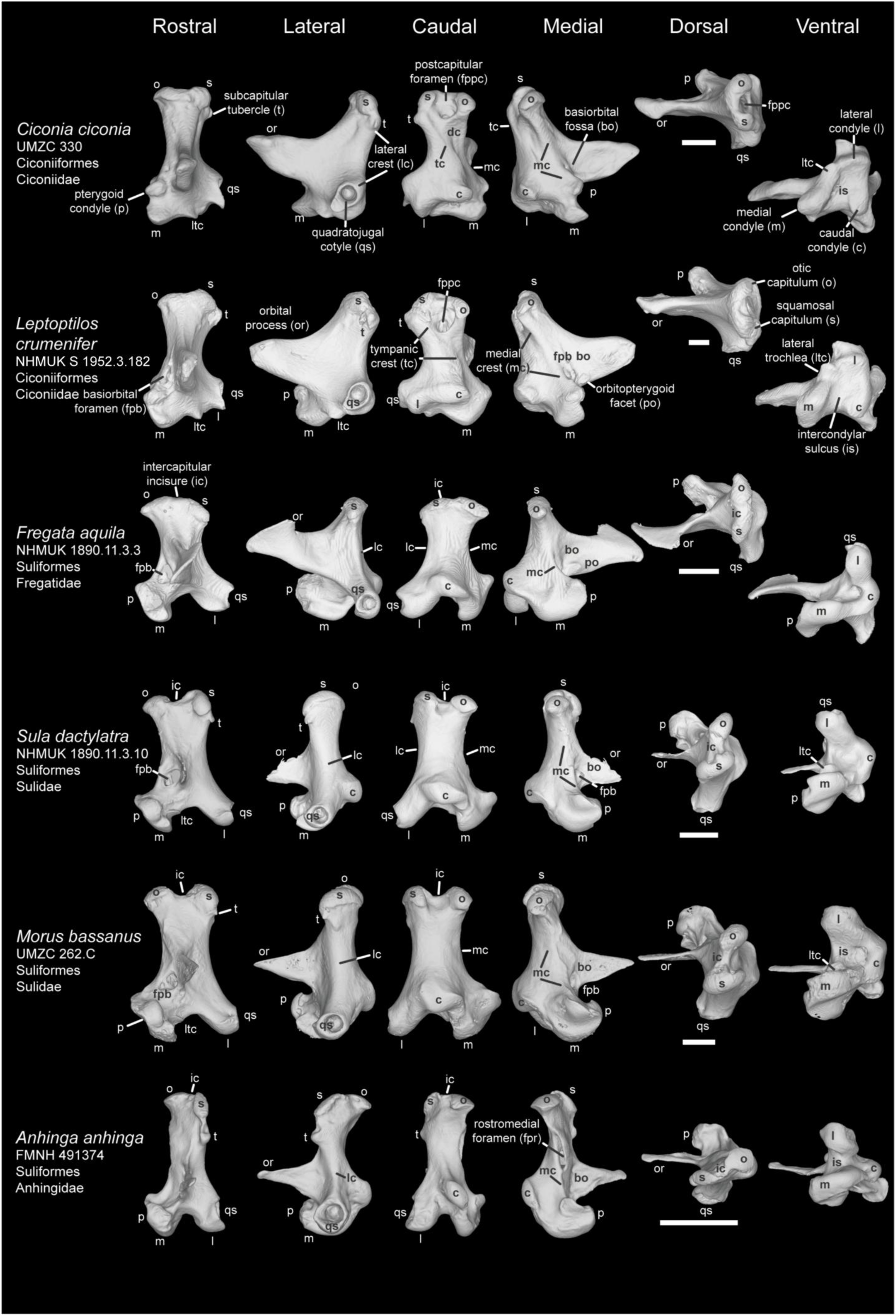
Comparison of avian quadrate among Aequorlitornithes Ⅷ (Ciconiiformes and Suliformes). Comparison of *Ciconia ciconia* (UMZC 330), *Leptoptilos crumenifer* (NHMUK S 1952.3.182), *Fregata aquila* (NHMUK 1890.11.3.3), *Sula dactylatra* (NHMUK 1890.11.3.10), *Morus bassanus* (UMZC 262.C), and *Anhinga anhinga* (FMNH 491374) in rostral, lateral caudal, medial, dorsal, and ventral view. Scale bar: 5 mm.

**Plate 31.**
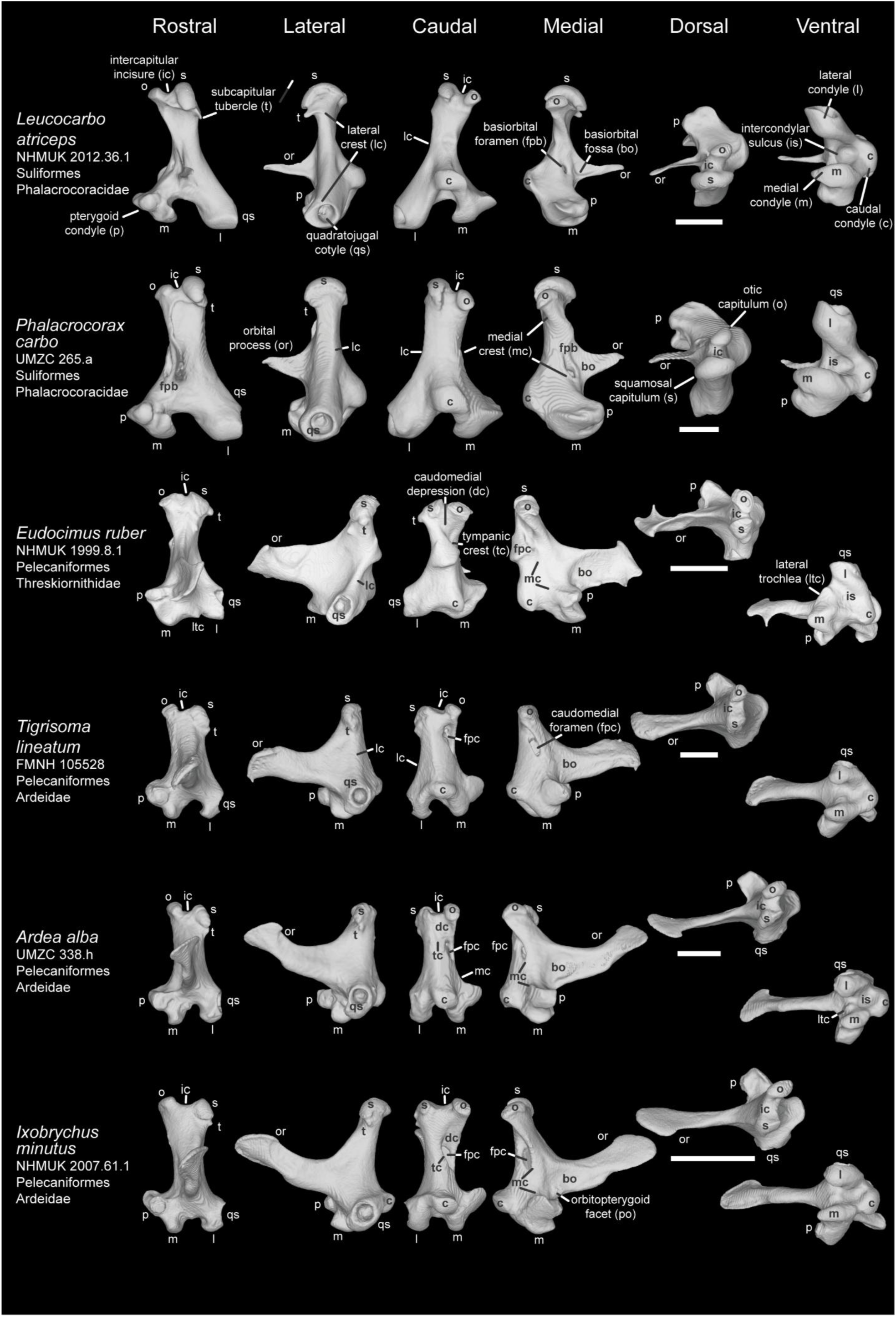
Comparison of avian quadrate among Aequorlitornithes Ⅸ (Suliformes and Pelecaniformes). Comparison of *Leucocarbo atriceps* (NHMUK 2012.36.1), *Phalacrocorax carbo* (UMZC 265.a), *Eudocimus ruber* (NHMUK 1999.8.1), *Tigrisoma lineatum* (FMNH 105528), *Ardea alba* (UMZC 338.h), and *Ixobrychus minutus* (NHMUK 2007.61.1) in rostral, lateral caudal, medial, dorsal, and ventral view. Scale bar: 5 mm.

**Plate 32.**
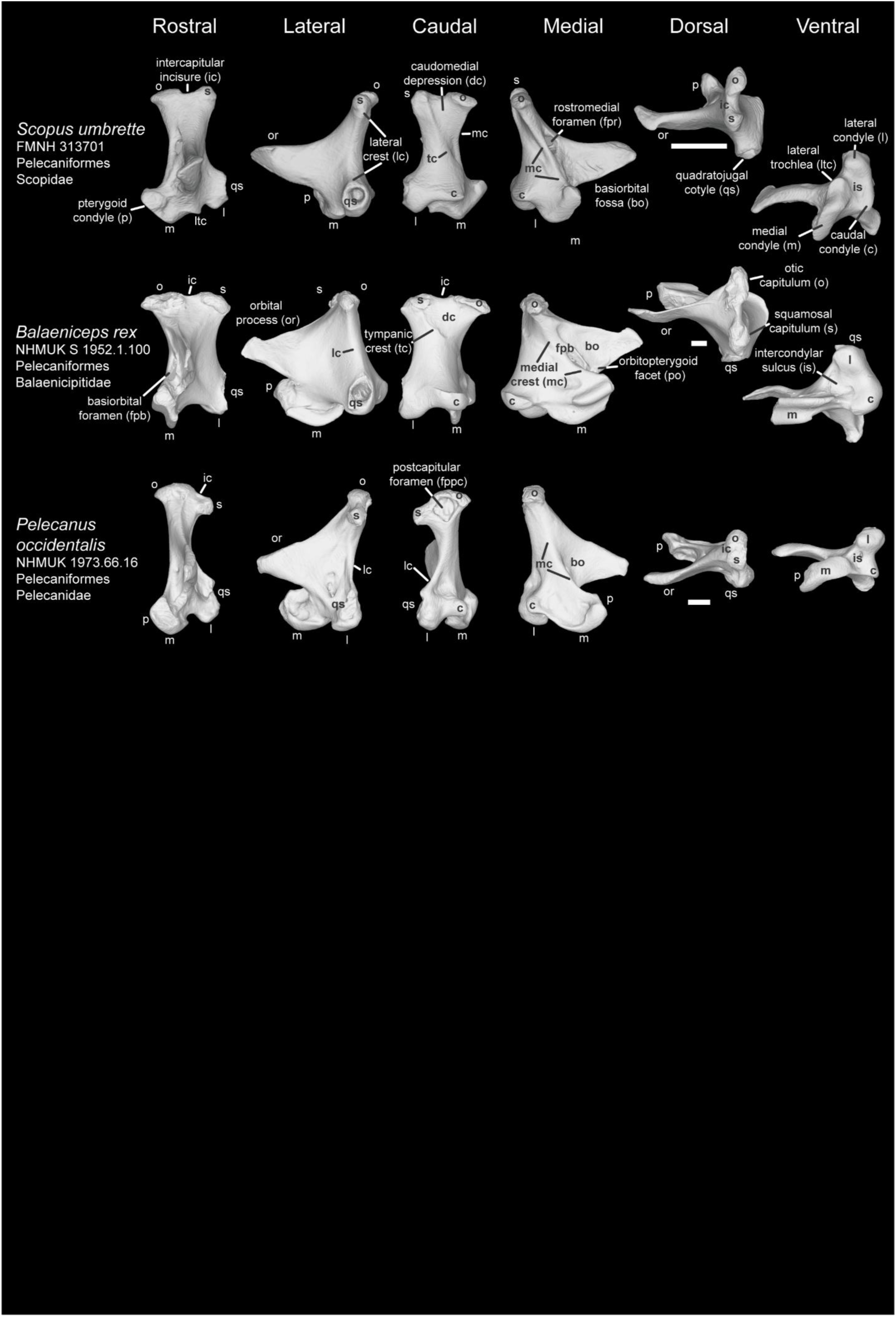
Comparison of avian quadrate among Aequorlitornithes Ⅹ (Pelecaniformes). Comparison of *Scopus umbretta* (FMNH 313701), *Balaeniceps rex* (NHMUK S 1952.1.100), and *Pelecanus occidentalis* (NHMUK 1973.66.16) in rostral, lateral caudal, medial, dorsal, and ventral view. Scale bar: 5 mm.

**Plate 33.**
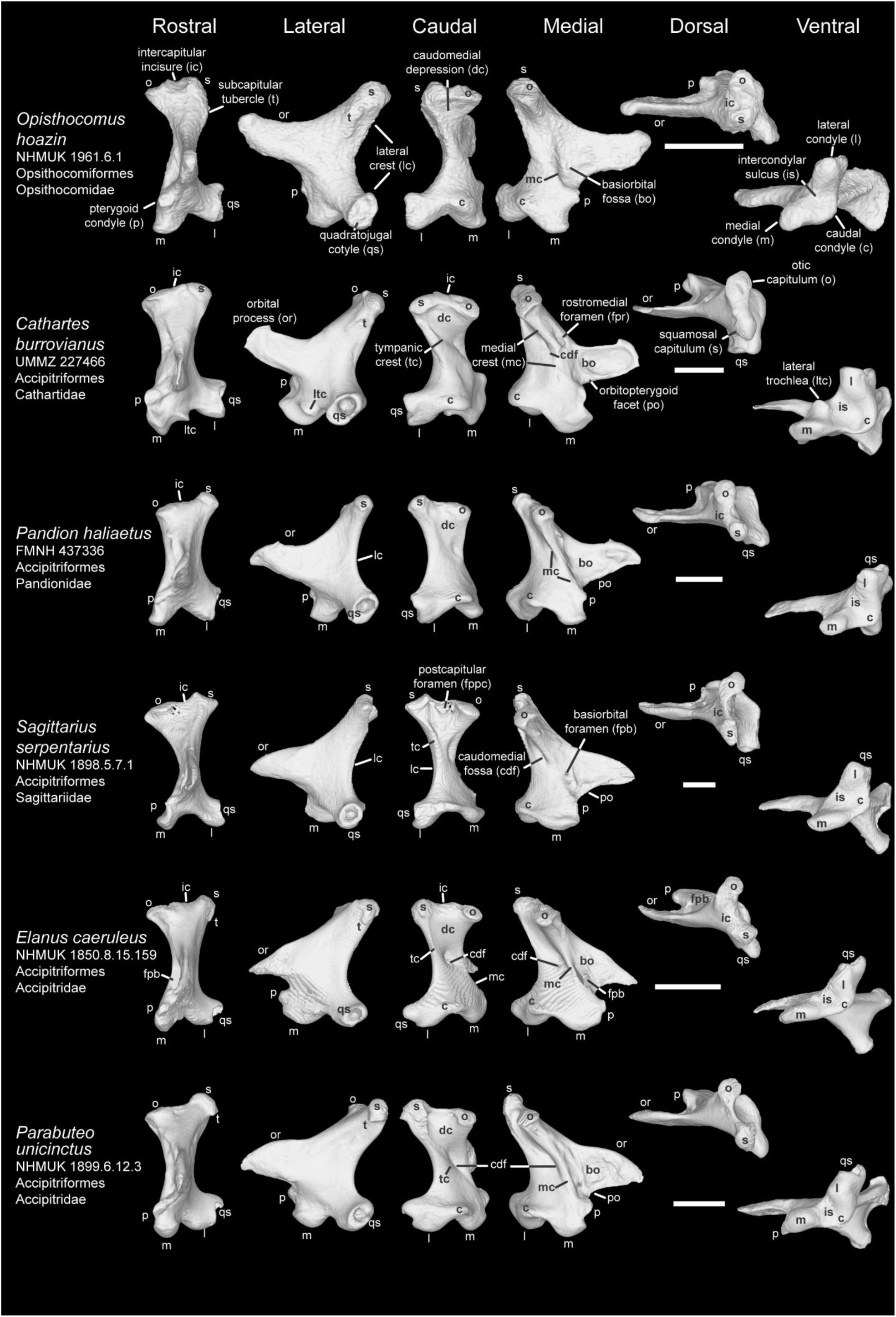
Comparison of avian quadrate among Opisthocomiformes and Accipitriformes Ⅰ. Comparison of *Opisthocomus hoazin* (NHMUK 1961.6.1), *Cathartes burrovianus* (UMMZ 227466), *Pandion haliaetus* (FMNH 437336), *Sagittarius serpentarius* (NHMUK 1898.5.7.1), *Elanus caeruleus* (NHMUK 1850.8.15.159), and *Parabuteo unicinctus* (NHMUK 1899.6.12.3) in rostral, lateral caudal, medial, dorsal, and ventral view. Scale bar: 5 mm.

**Plate 34.**
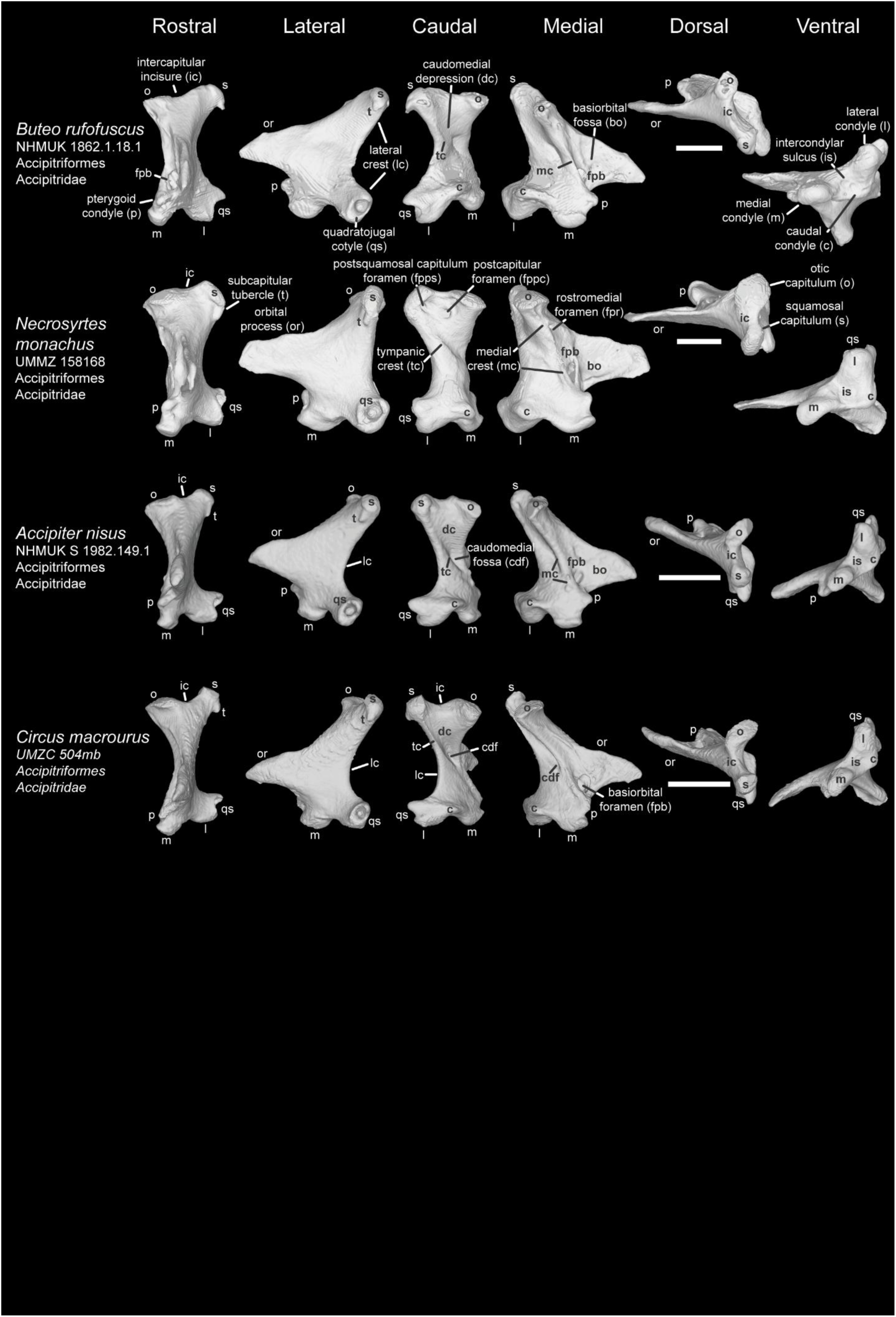
Comparison of avian quadrate among Accipitriformes Ⅱ. Comparison of *Buteo rufofuscus* (NHMUK 1862.1.18.1), *Necrosyrtes monachus* (UMMZ 158168), *Accipiter nisus* (NHMUK S 1982.149.1), and *Circus macrourus* (UMZC 504mb) in rostral, lateral caudal, medial, dorsal, and ventral view. Scale bar: 5 mm.

**Plate 35.**
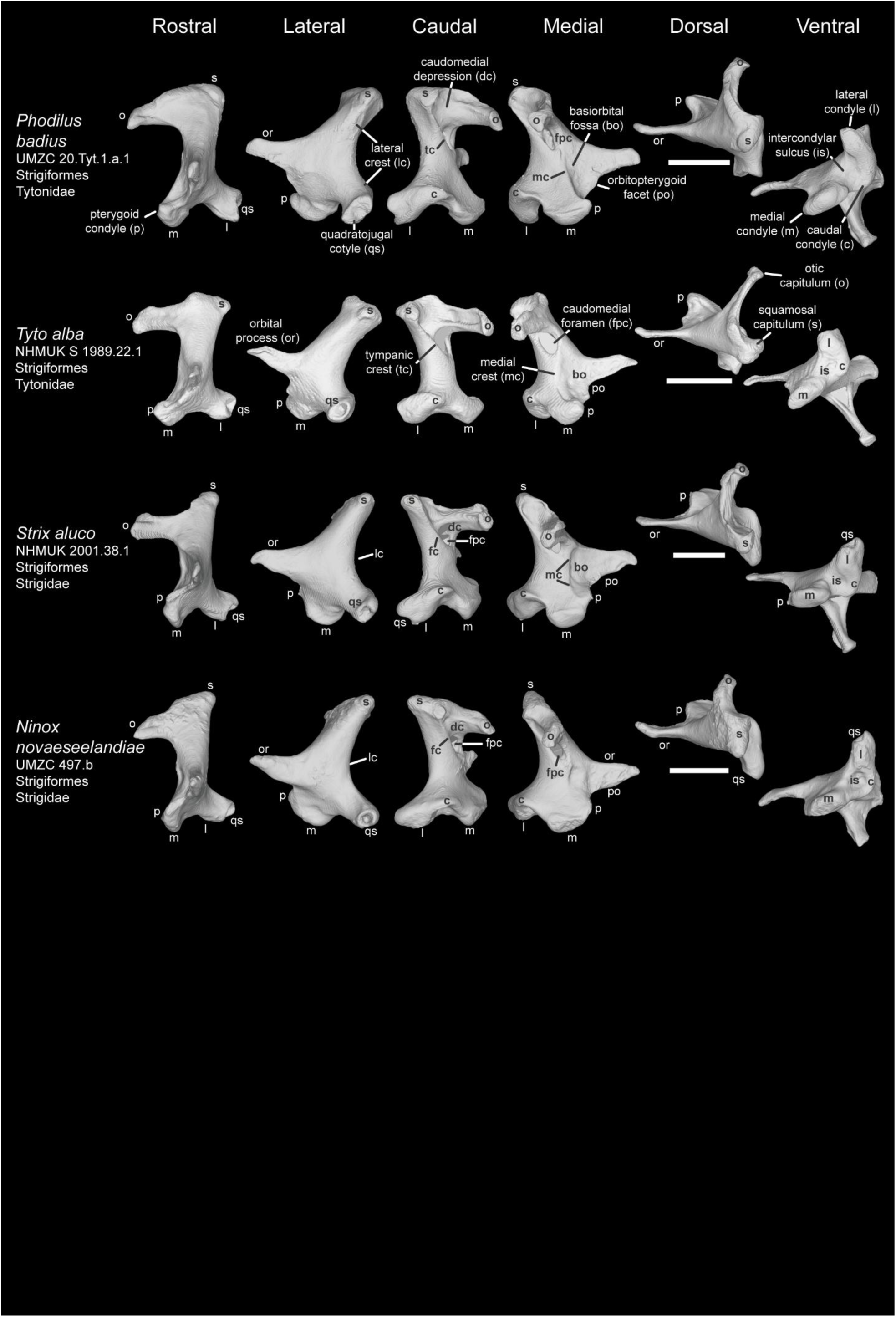
Comparison of avian quadrate among Strigiformes. Comparison of *Phodilus badius* (UMZC 20.Tyt.1.a.1), *Tyto alba* (NHMUK S 1989.22.1), *Strix aluco* (NHMUK 2001.38.1), and *Ninox novaeseelandiae* (UMZC 497.b) in rostral, lateral caudal, medial, dorsal, and ventral view. Scale bar: 5 mm.

**Plate 36.**
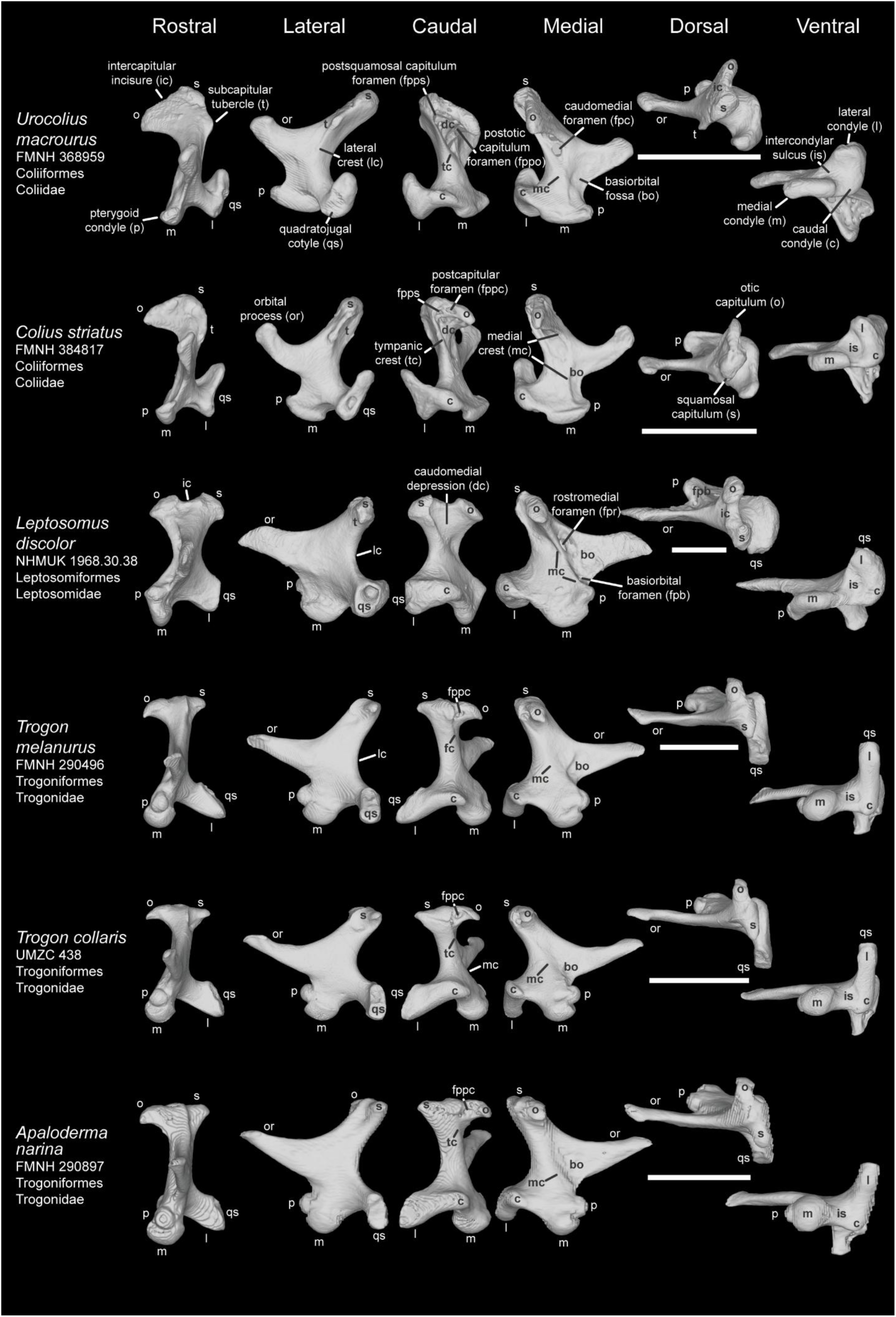
Comparison of avian quadrate among Coraciimorphae Ⅰ (Coliiformes, Leptosomiformes, and Trogoniformes). Comparison of *Urocolius macrourus* (FMNH 368959), *Colius striatus* (FMNH 384817), *Leptosomus discolor* (NHMUK 1968.30.38), *Trogon melanurus* (FMNH 290496), *T. collaris* (UMZC 438), and *Apaloderma narina* (FMNH 290897) in rostral, lateral caudal, medial, dorsal, and ventral view. Scale bar: 5 mm.

**Plate 37.**
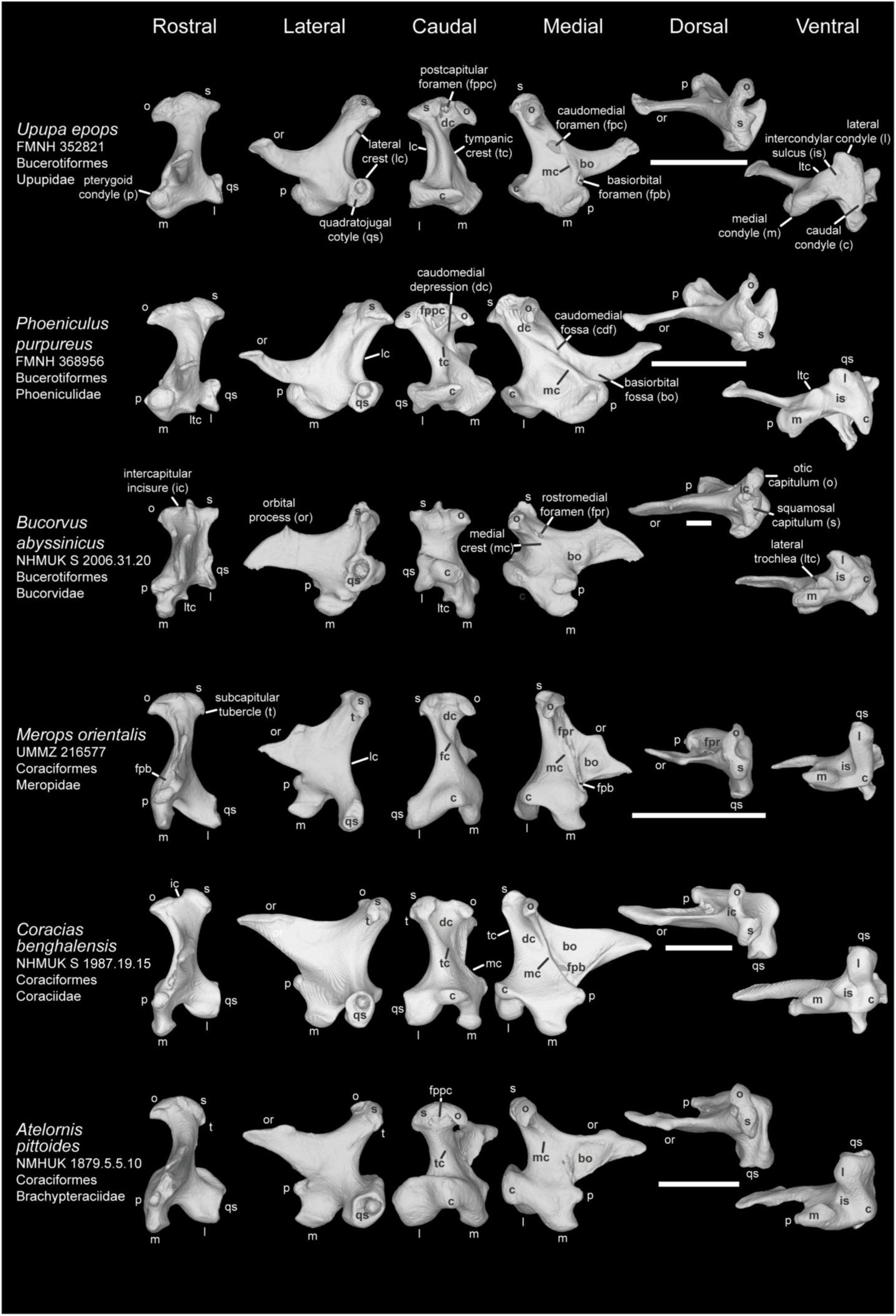
Comparison of avian quadrate among Coraciimorphae Ⅱ (Bucerotiformes and Coraciiformes). Comparison of *Upupa epops* (FMNH 352821), *Phoeniculus purpureus* (FMNH 368956), *Bucorvus abyssinicus* (NHMUK S 2006.31.20), *Merops orientalis* (UMMZ 216577), *Coracias benghalensis* (NHMUK S 1987.19.15), and *Atelornis pittoides* (NMHUK 1879.5.5.10) in rostral, lateral caudal, medial, dorsal, and ventral view. Scale bar: 5 mm.

**Plate 38.**
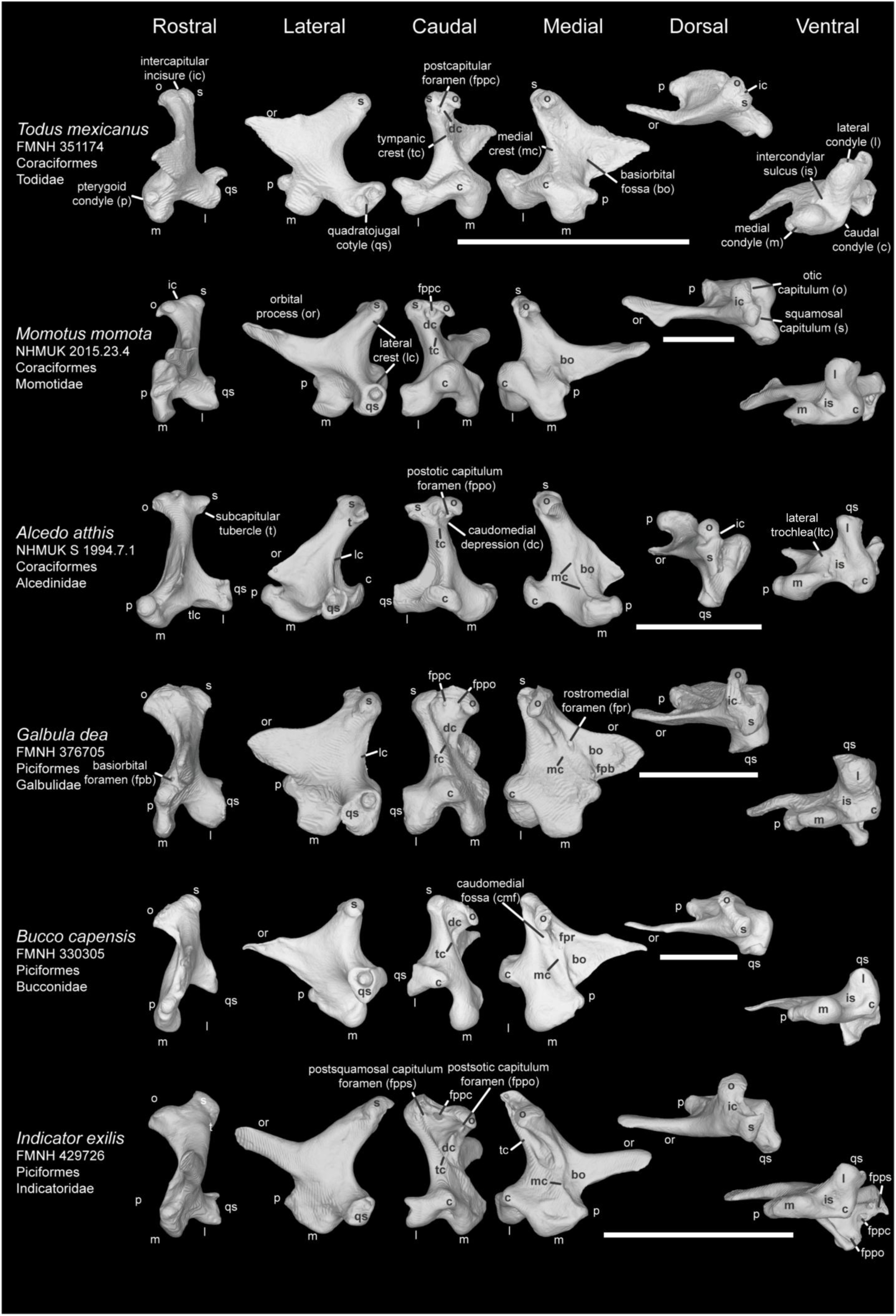
Comparison of avian quadrate among Coraciimorphae Ⅲ (Coraciiformes and Piciformes). Comparison of *Todus mexicanus* (FMNH 351174), *Momotus momota* (NHMUK 2015.23.4), *Alcedo atthis* (NHMUK S 1994.7.1), *Galbula dea* (FMNH 376705), *Bucco capensis* (FMNH 330305), and *Indicator exilis* (FMNH 429726) in rostral, lateral caudal, medial, dorsal, and ventral view. Scale bar: 5 mm.

**Plate 39.**
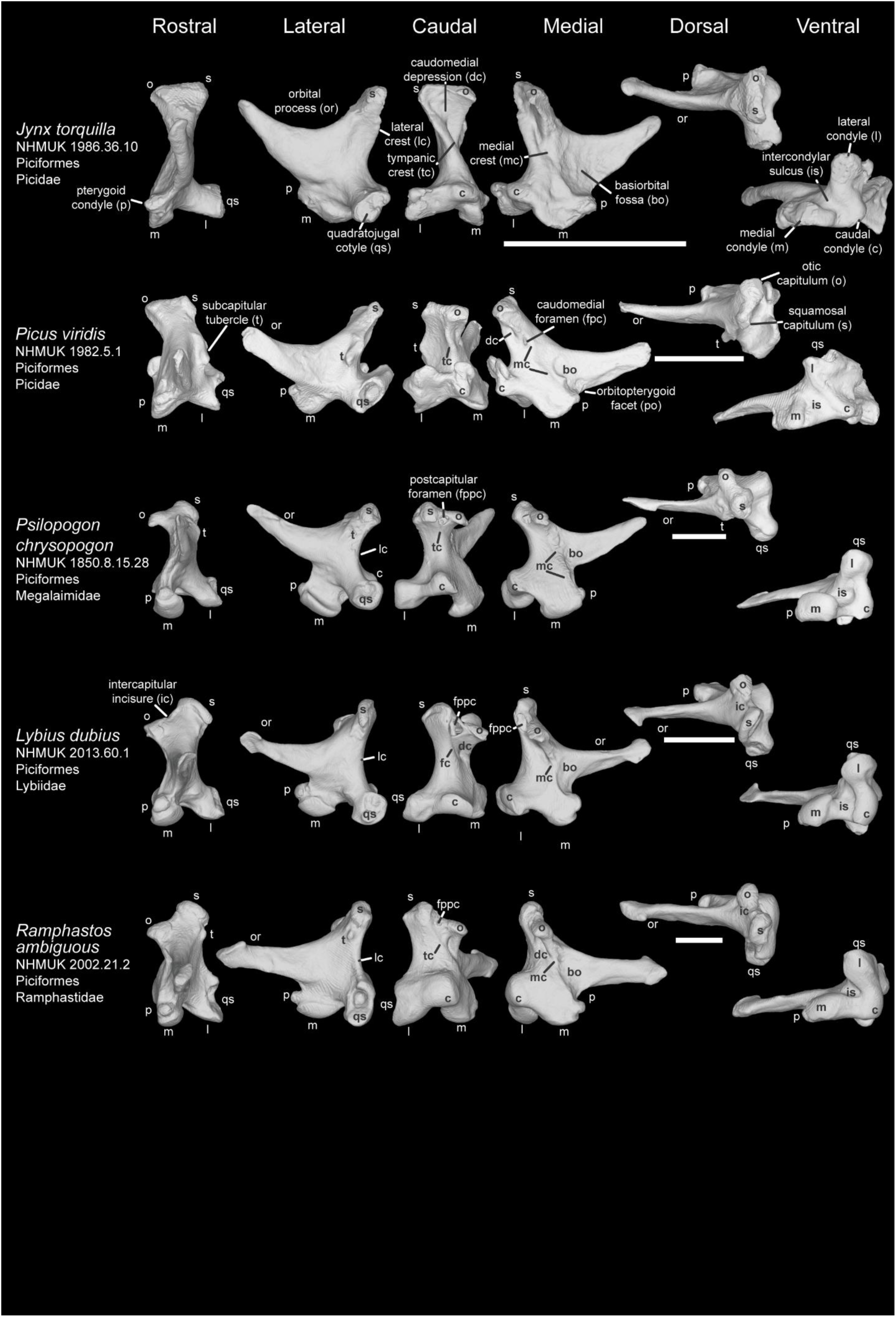
Comparison of avian quadrate among Coraciimorphae Ⅳ (Piciformes). Comparison of *Jynx torquilla* (NHMUK 1986.36.10), *Picus viridis* (NHMUK 1982.5.1), *Psilopogon chrysopogon* (NHMUK 1850.8.15.28), *Lybius dubius* (NHMUK 2013.60.1), and *Ramphastos ambiguus* (NHMUK 2002.21.2) in rostral, lateral caudal, medial, dorsal, and ventral view. Scale bar: 5 mm.

**Plate 40.**
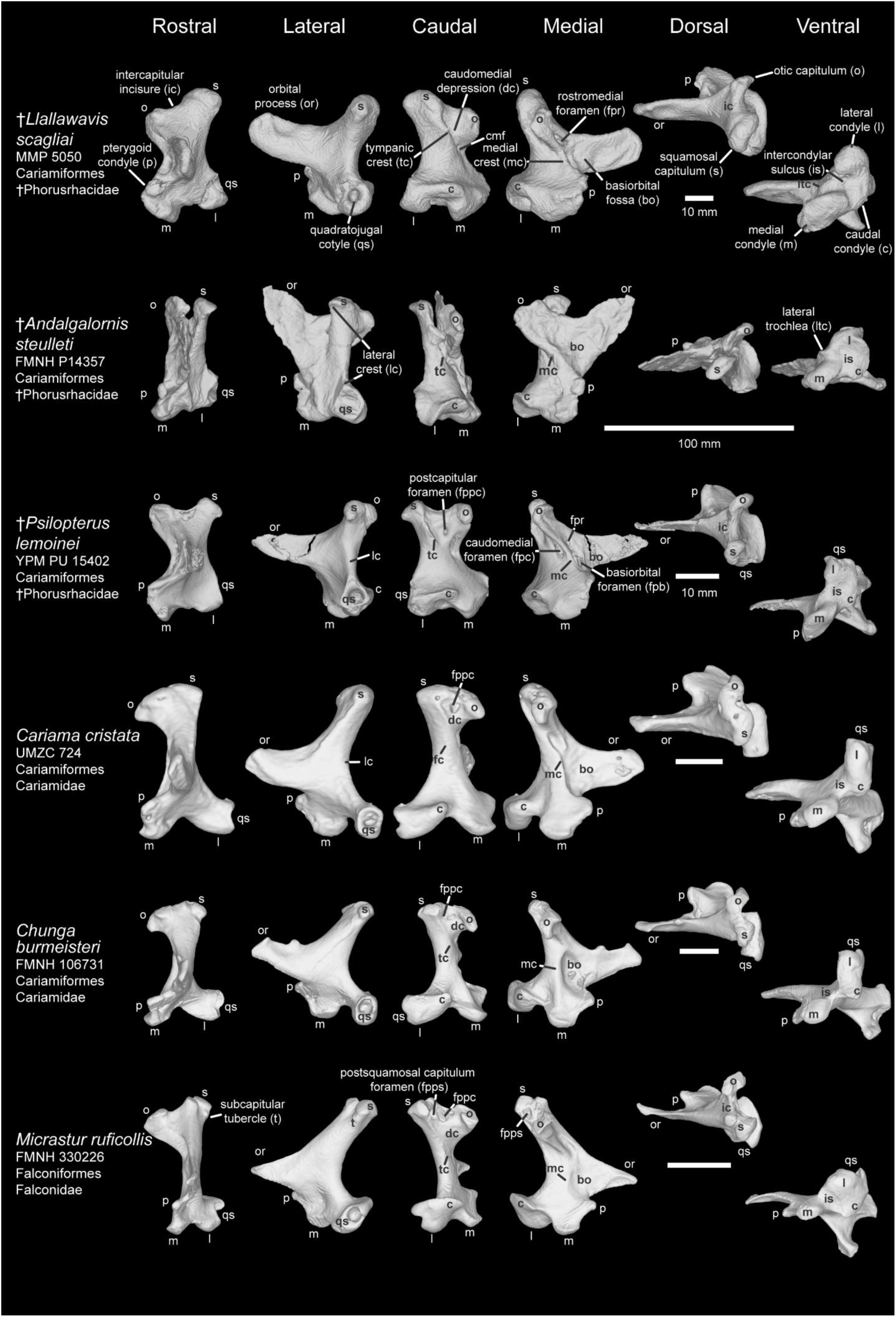
Comparison of avian quadrate among Australaves Ⅰ (Cariamiformes and Falconiformes). Comparison of *Llallawavis scagliai* (MMP 5050), *Andalgalornis steulleti* (FMNH P14357), *Psilopterus lemoinei* (YPM PU 15402), *Cariama cristata* (UMZC 724), *Chunga burmeisteri* (FMNH 106731), and *Micrastur ruficollis* (FMNH 330226) in rostral, lateral caudal, medial, dorsal, and ventral view. Scale bar: 5 mm. (Except for *Llallawavis scagliai*, *Andalgalornis steulleti*, and *Psilopterus lemoinei*).

**Plate 41.**
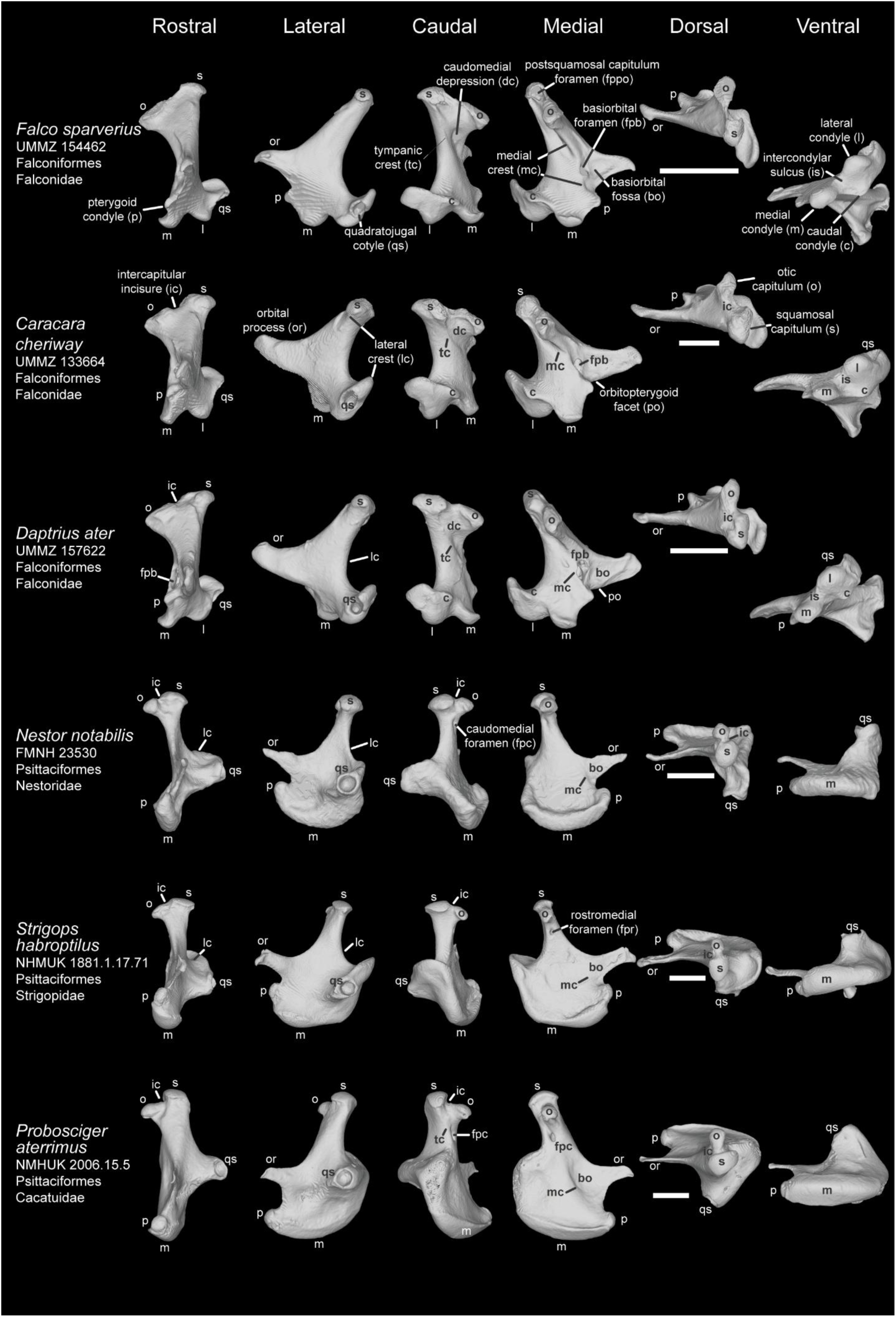
Comparison of avian quadrate among Australaves Ⅱ (Falconiformes and Psittaciformes). Comparison of *Falco sparverius* (UMMZ 154462), *Caracara cheriway* (UMMZ 133664), *Daptrius ater* (UMMZ 157622), *Nestor notabilis* (FMNH 23530), *Strigops habroptilus* (NHMUK 1881.1.17.71), and *Probosciger aterrimus* (NHMUK 2006.15.5) in rostral, lateral caudal, medial, dorsal, and ventral view. Scale bar: 5 mm.

**Plate 42.**
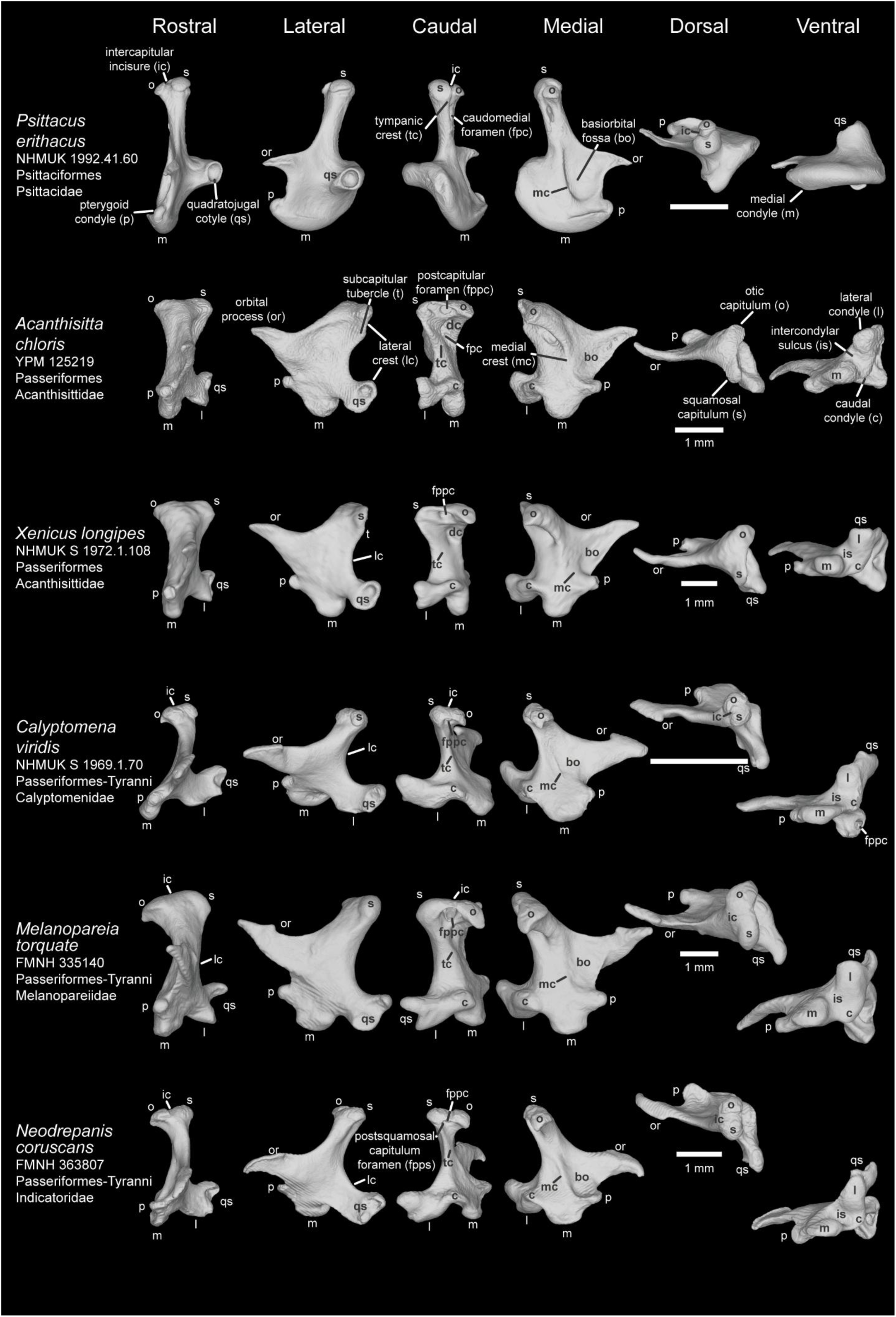
Comparison of avian quadrate among Australaves Ⅲ (Psittaciformes and Passeriformes). Comparison of *Psittacus erithacus* (NHMUK 1992.41.60), *Acanthisitta chloris* (YPM 125219), *Xenicus longipes* (NHMUK S 1972.1.108), *Calyptomena viridis* (NHMUK S 1969.1.70), *Melanopareia torquata* (FMNH 335140), and *Neodrepanis coruscans* (FMNH 363807) in rostral, lateral caudal, medial, dorsal, and ventral view. Scale bar: 5 mm. (Except for *Acanthisitta chloris*, *Xenicus longipes*, *Melanopareia torquata*, and *Neodrepanis coruscans*).

**Plate 43.**
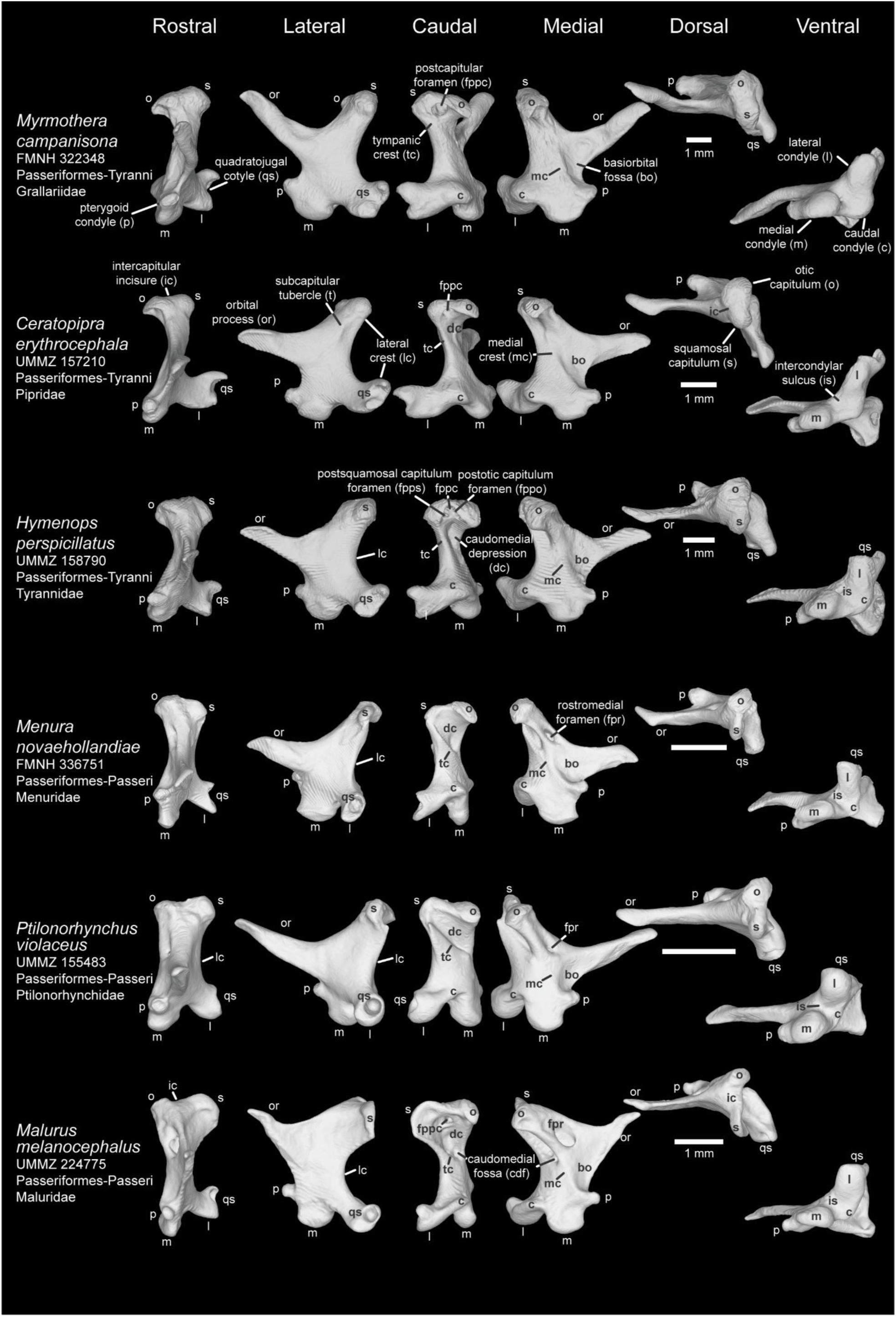
Comparison of avian quadrate among Australaves Ⅳ (Passeriformes). Comparison of *Myrmothera campanisona* (FMNH 322348), *Ceratopipra erythrocephala* (UMMZ 157210), *Hymenops perspicillatus* (UMMZ 158790), *Menura novaehollandiae* (FMNH 336751), *Ptilonorhynchus violaceus* (UMMZ 155483), and *Malurus melanocephalus* (UMMZ 224775) in rostral, lateral caudal, medial, dorsal, and ventral view. Scale bar: 5 mm. (Except for *Myrmothera campanisona*, *Ceratopipra erythrocephala*, *Hymenops perspicillatus*, and *Malurus melanocephalus*).

**Plate 44.**
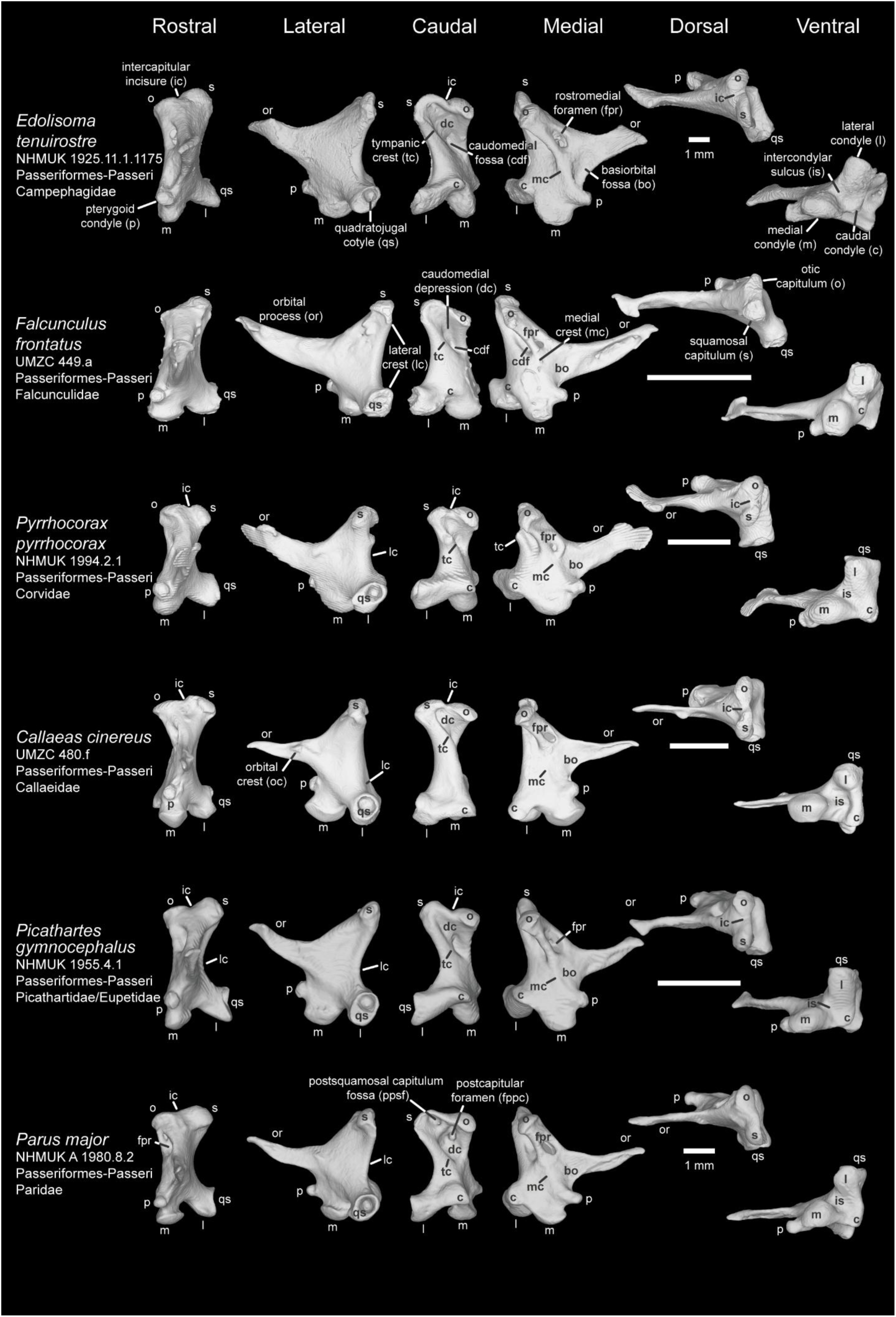
Comparison of avian quadrate among Australaves Ⅴ (Passeriformes). Comparison of *Edolisoma tenuirostre* (NHMUK 1925.11.1.1175), *Falcunculus frontatus* (UMZC 449.a), *Pyrrhocorax pyrrhocorax* (NHMUK 1994.2.1), *Callaeas cinereus* (UMZC 480.f), *Picathartes gymnocephalus* (NHMUK 1955.4.1), and *Parus major* (NHMUK A 1980.8.2) in rostral, lateral caudal, medial, dorsal, and ventral view. Scale bar: 5 mm. (Except for *Edolisoma tenuirostre* and *Parus major*).

**Plate 45.**
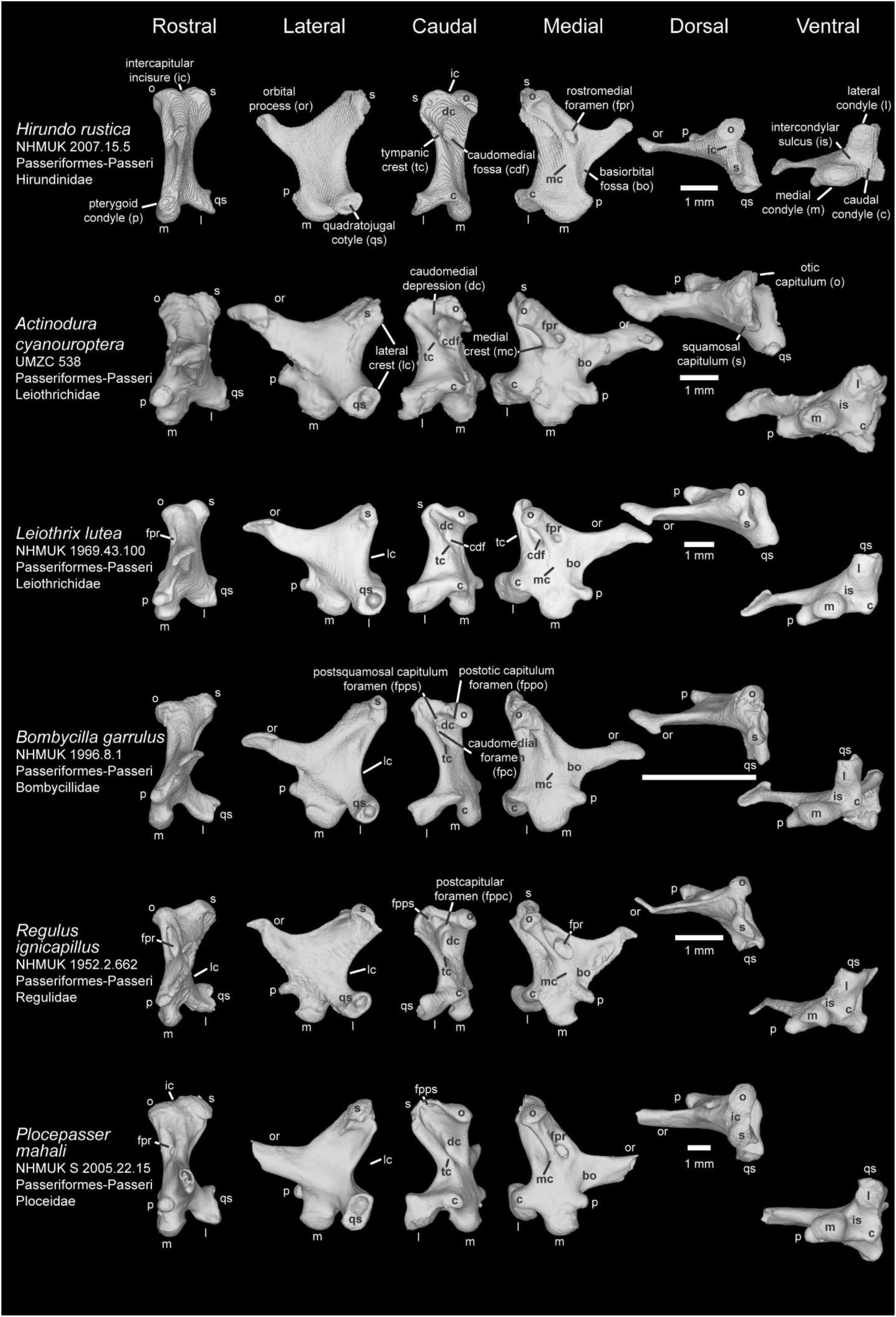
Comparison of avian quadrate among Australaves Ⅵ (Passeriformes). Comparison of *Hirundo rustica* (NHMUK 2007.15.5), *Actinodura cyanouroptera* (UMZC 538), *Leiothrix lutea* (NHMUK 1969.43.100), *Bombycilla garrulus* (NHMUK 1996.8.1), *Regulus ignicapillus* (NHMUK 1952.2.662), and *Plocepasser mahali* (NHMUK S 2005.22.15) in rostral, lateral caudal, medial, dorsal, and ventral view. Scale bar: 5 mm. (Except for *Hirundo rustica*, *Actinodura cyanouroptera*, *Leiothrix lutea*, *Regulus ignicapillus*, and *Plocepasser mahali*).

**Plate 46.**
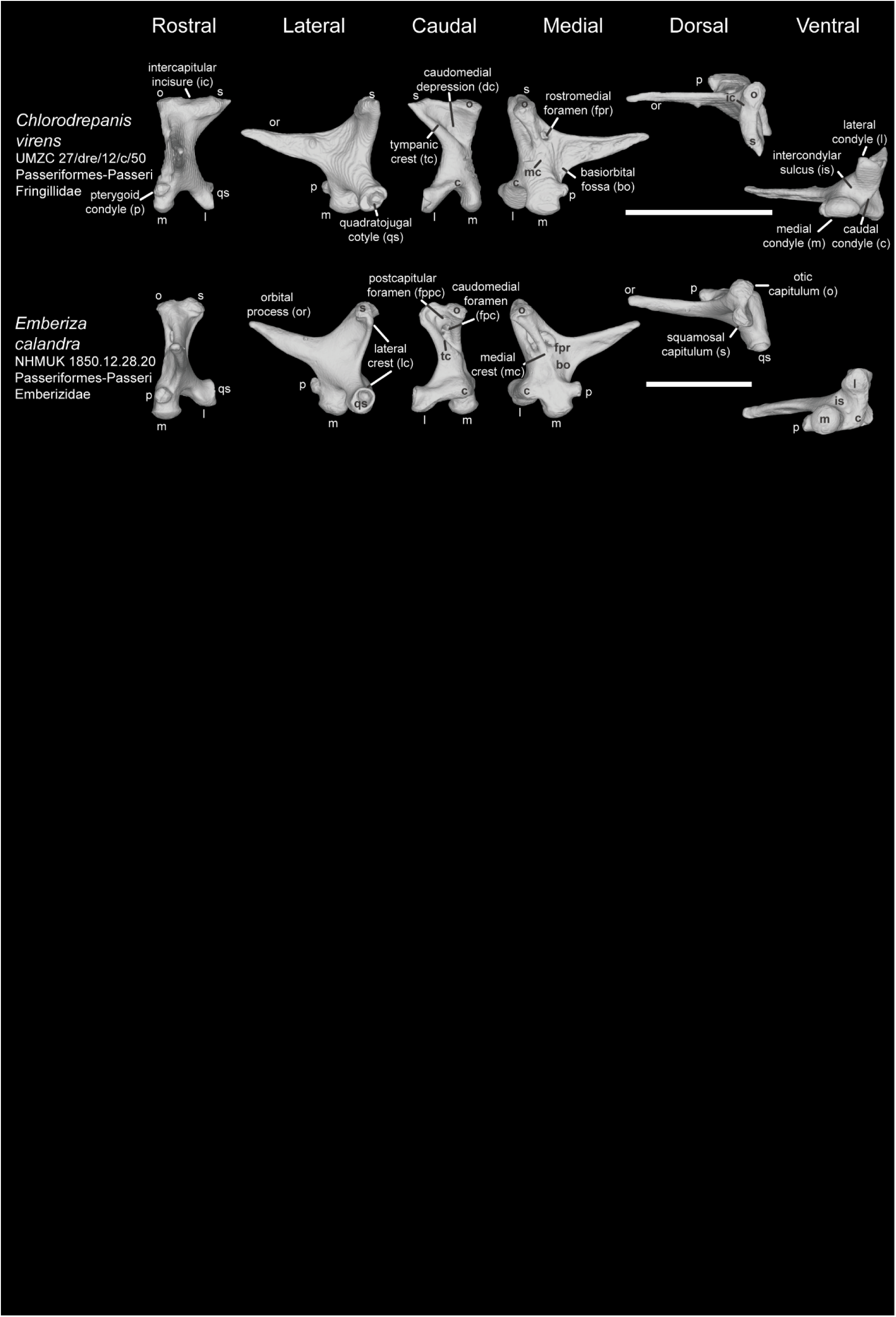
Comparison of avian quadrate among Australaves Ⅶ (Passeriformes). Comparison of *Chlorodrepanis virens* (UMZC 27/dre/12/c/50) and *Emberiza calandra* (NHMUK 1850.12.28.20) in rostral, lateral caudal, medial, dorsal, and ventral view. Scale bar: 5 mm.

