## Supplementary for "Revised nomenclature of avian quadrate morphology and a detailed survey of clade-specific anatomical features"

**Ichthyornithes:** ^†^*Ichthyornis* (Plate 1)

The otic process in most *Ichthyornis* shows that the otic capitulum is much higher than squamosal capitulum in either rostral or caudal view, different with most neornithine quadrates. All *Ichthyornis* quadrates show an oval shape of squamosal capitulum with a lateromedially elongate articular surface, while their otic capitula are diamond-shaped with flat articular surfaces. The squamosal capitulum of KUVP 119673 and AMNH FRAB 32773 is rostrocaudally wider than that of other two *Ichthyornis* specimens. By comparing size with each capitulum of *Ichthyornis* quadrates, the squamosal capitulum is smaller than otic capitulum. Between these two capitula, a shallow but distinct intercapitular incisure present (Clarke, 2004), indicating that a bicondylar morphology likely is plesiomorphic for Aves (Field et al., 2018).

*Ichthyornis* quadrates exhibit some morphological variance on the orbital process among different specimens. For instance, the orbital process is much curved on specimen FHSU 18702 than specimen KUVP 119673 and specimen AMNH FRAB 32773, which are relatively straight at the base of the orbital process. Surprisedly, the orbital process is elongate with a boot-like or hammer-like tip on AMNH FRAB 32773 (Torres et al., 2021), similar as the tip on Tinamiformes (e.g., *Crypturellus tataupa* and *Nothoprocta ornate*) and ^†^*Madrynornis mirandus* (Degrange et al., 2018) whose orbital process also has a boot-like tip. Comparing the shape variance of the orbital process tip with the abovementioned specimens, the tip of tinamiform orbital process is much slender and the tip of *Madrynornis* orbital process is much robust.

The quadratojugal cotyle of the *Ichthyornis* quadrates is a shallow fossa with a thick complete circle margin, and it is adjacent with the lateral condyle (Clarke, 2004). The pterygoid condyle of the *Ichthyornis* quadrates is rostrally blunt with an elliptical shape. Like ^†^*Asteriornis*, ^†^*Prebsyornis*, ^†^Pelagornithidae (^†^*Dasornis toliapica* and ^†^*Osteodontornis sp* in this study), extant Galloanserae, and Columbidae, *Ichthyornis* quadrate has a bicondylar mandibular process, lacking the caudal condyle. These two condyles (medial condyle and lateral condyle) are more lateromedially arrayed. Nevertheless, the Intercondylar sulcus of the *Ichthyornis* quadrates is much wider than any groups abovementioned. The medial condyle is prominently elongate, dorsocaudalward extending, forming a shape like Statue of Liberty's right feet, while the lateral condyle exhibits a flat and smooth articular surface.

The quadrate body of *Ichthyornis* is relatively straight due to both straight lateral and medial crest. The caudomedial depression and tympanic crest is absent on quadrate body. A single and large pneumatic foramen (basiorbital foramen) is medially located at the quadrate body (e.g., specimen BHI 6421 and KUVP 119673), close to the base of the orbital process and pterygoid condyle (Clarke, 2004).

**Palaeognathae**: ^†^Lithornithiformes (Plate 2)

The otic process shows a huge morphological disparity in Lithornithiformes. In *Lithornis celetius*, two capitulua are continuous, similar as that of most extant Palaeognathae. However, it shows a clear intercapitular incisure between two capitula in *L. plebius* quadrate. Both squamosal and otic capitulum are oval shaped with equal size. The squamosal capitulum is rostrocaudally elongate and faces more rostrolaterally instead of dorsally. On the other hand, the otic capitulum is lateromedially elongate with a flat articular surface and faces more dorsocaudally.

The orbital process of Lithornithiformes quadrate has a relatively high aspect ratio with a blunt, robust, and rounded tip (Leonard et al., 2005; Nesbitt and Clarke, 2016). Its tip points more medially in *L. celetius*, while it points more rostrally in *L. plebius*. This difference might result from the fact that the orbital process of *L. celetius* is slightly broken and compressed.

The quadratojugal cotyle shows a cup shape with a complete and thick circular margin, adjacent to the medial condyle. The rostral margin of the quadratojugal cotyle is distinct, similar as rhea (Rheiformes), kiwi (Apterygiformes) and some tinamou (Tinamiformes) (Houde, 1988). Unlike most avian quadrates, the pterygoid condyle of Lithornithiformes quadrate does not exhibit a ball-shaped process ; instead, it is a flat articular surface below the orbital process (Houde, 1988), facing medially. Therefore, their pterygoid condyles are obviously separated from the mandibular process by a distinct gap. This unique scenario is only similar as tinamou, and perhaps, suggesting that the palatine system of Lithornithiformes skull shows a similar biomechanical function as that of tinamou.

The mandibular process of Lithornithiformes quadrate has three condyles (medial, lateral, and caudal condyle), and the lateral condyle is continuous with caudal condyle (Houde, 1988). The medial condyle shows a shallow articular surface with a concave articular surface (lateral trochlea) to connect with the lower jawbone. The lateral condyle has a flat articular surface, and it is located caudally with the quadratojugal cotyle. The caudal condyle tilts medially.

The lithornithiform quadrate body is relatively medially curved due to a distinctly curved medial crest. Apparently, there is a clear fossa/depression at the caudal side of quadrate body, far below the otic process. Besides, a pneumatic foramen (postcapitular foramen) is caudally located at the middle position between two capitulua on *L. plebius*. This might indicate that the pneumaticity is variant among different species in Lithornithiformes.

**Palaeognathae:** Struthioniformes (Plate 2)

On the otic process of ostrich (Struthioniformes), the intercapitular incisure is absent or under-developed between two capitula. Both capitula show distinct shapes: the squamosal capitulum has a ball-like shape with a larger size, facing more laterodorsally. The otic capitulum has an oval shape with a flat articular surface, and faces more mediodorsally.

The orbital process is massive (Saiff, 1981) and has a relatively high aspect ratio with a blunt and flat tip, pointing rostrodorsally. There is an obvious bump located at the dorsal margin of the orbital process, and it is part of the pterygoid articular surface (orbitopterygoid facet).

The quadratojugal cotyle of ostrich quadrate exhibits a cup shape with a complete circular margin, adjacent to the lateral condyle. The ostrich quadrate has two articular facets with the pterygoid: one (pterygoid condyle) is just above the medial condyle with an extremely flat articular surface, and the other (orbitopterygoid facet) is below the orbital process with a distinct attachment.

The mandibular process on ostrich quadrate has three condyles, but the third condyle (caudal condyle) is not well developed, as shown in a tiny tilt. The medial condyle has an oval shape with a smooth articular surface. The lateral condyle has a flat articular surface and is continuous with the caudal condyle (Saiff, 1981). A clear and huge fossa/depression is located between the medial and lateral condyle, close to the centre of the mandibular process.

The quadrate body of ostrich looks relatively thin with wider otic process and wider mandibular process, and there is not any pneumatic foramen on its quadrate. The ostrich quadrate in this study is sub-adult, and therefore, some of anatomical observation abovementioned (e.g., pneumatic foramen) might be strongly influenced by the ontogenetic development (Plateau et al., 2024).

**Palaeognathae**: Rheiformes (Plate 2 and 3)

The rhea quadrate share many characters with ostrich quadrate, such as the shape of the otic process, orbital process, and the articular facet with the pterygoid. The squamosal capitulum of rhea quadrate exhibits a square-shaped with a bulbed articular surface, while its otic capitulum is oval, facing dorsally.

The orbital process is thick and has relatively smaller aspect ratio compared with ostrich quadrate. The tip of the orbital process is blunt and flat, and points more medially. On the ventral side of the orbital process tip, a slant surface is present, perhaps, for the muscle attachment.

The quadratojugal cotyle has a cup shape with a thin margin, adjacent to the lateral condyle. Unlike Struthioniformes, the quadratojugal cotyle has a flat articular surface at the rostral margin for attaching jugal bar. The articular surface with pterygoid on rhea quadrate shows two facets, similar as ostrich: one facet (pterygoid condyle) is just above the medial condyle with a distinct gap with a flat articular surface, and the other (orbitopterygoid facet) is below the orbital process. However, unlike ostrich, the facet below the orbital process doesn’t show a clear attachment on rhea quadrate.

The mandibular process on Rheiformes quadrate has three condyles, and the lateral condyle is not continuous with caudal condyle. The medial condyle is slightly deep and only specimen FMNH B 339616 shows a shallow lateral trochlea contacting with the lower jaw. The lateral condyle has a small and flat articular surface below the quadratojugal cotyle. The caudal condyle is large and well-developed and tilts caudally.

The quadrate body of Rheiformes is relatively medially curved with a clear medial crest, while the lateral crest is not well-developed. A relatively large pneumatic foramen (postcapitular foramen) is caudally located below the otic process.

**Palaeognathae:** Apterygiformes (Plate 3)

The apterygiform quadrate shows a significantly variant in morphology of several characters. In *Apteryx australis*, the squamosal and otic capitula are oval and equal in size. The squamosal capitulum is mediolaterally convex and faces more caudolaterlly, whereas the otic capitulum is rostrocaudally elongate and oriented more dorsally. In contrast, in *A. owenii*, the squamosal capitulum is apparently larger than the otic capitulum. Both the squamosal and otic capitulum in *A. owenii* are oval and mediolaterally elongate, and they faces more rostrolaterally and caudodorsally, respectively. A protrusion is present below the otic capitulum in *A. owenii*, but is absent in *A. australis*. The intercapitular incisure is absent in *A. australis* (Saiff, 1982) but is distinct in *A. owenii*.

The orbital process has a relatively high aspect ratio (Saiff, 1982) and exhibits species-specific variation in the shape of its tip. In *A. australis*, the tip of the orbital process is blunt, robust and flat, and is oriented more rostrodorsally; in *A. owenii*, the tip is flat and slant, facing rostrodorsally. In addition, the orbital process of *A. australis* is more robust and straighter than that of *A. owenii*, whose orbital process is relatively elongate and slender.

The quadratojugal cotyle is adjacent to the lateral condyle but its shape is more diverse than in other palaeognath groups. In *A. australis*, the quadratojual cotyle is cashew-like shape, with a very shallow fossa for articulation with jugal bar, whereas in *A. owenii* it is oval and bears a deep articular surface. The rostral margin of the quadratojugal cotyle in Apterygiformes is distinct for jugal bar attachment (Saif, 1982). Unlike most birds, the pterygoid condyle in Apterygiformes is rectangular in shape, with a relatively smooth articular surface and a distinct gap separating it from the medial condyle.

The mandibular process of the apterygiform quadrate has three condyles, and the lateral condyle is continuous with the caudal condyle, forming a S-shaped outline. In *A. australis*, the medial condyle forms a distinct ridge along the middle margin. The lateral condyle has a flat articular surface, and the caudal condyle is well-developed and slightly caudally inclined. The medial condyle is separated from the lateral and caudal condyle with a shallow intercondylar sulcus (Saif, 1982). A larege fossa is present on the mandibular process, positioned rostral to the caudal condyle (Saif, 1982).

The quadrate body of Apterygiformes is relatively laterally curved with a distinct medial and lateral crest. Pneumaticity varies within the group, with foramina occurring in two different positions. In *A. australis*, pneumatic foramina are restricted to positions below the otic capitulum (postotic capitulum foramen) and below the otic process (postcapitular foramen). In contrast, in *A. owenii*, the otic process is highly pneumatic, including the postsquamosal capitulum foramen and postotic capitulum foramen.

**Palaeognathae:** Casuariiformes (Plate 3)

Like most palaeognath quadrates, the two capitula of casuariiform quadrates are continuous without a distinct intercapitular incisure between them. Both the squamosal and otic capitula are oval in shape. The squamosal capitulum is much larger than the otic capitulum in *Casuarius casuarius*, whereas they are equal in size in *Dromaius novaehollandiae*. The squamosal capitulum is mediolaterally convex. The otic capitulum has relative a smooth articular surface, and points more dorsomedially.

The orbital process is significantly massive and has a relatively high aspect ratio. In *C. casuarius*, the tip of the orbital process looks similar to an arrow and faces rostrally. In contrast, the orbital process tips in *D. novaehollandiae* is oriented more ventrally.

The quadratojugal cotyle has a deep fossa with a complete and very thick circular margin, adjacent to the lateral condyle. Unlike most avian groups, the quadratojugal cotyle is oriented more lateroventrally, instead of laterally. Similar to that of most palaeognath quadrates, the articular contact with the pterygoid on Casuariiformes quadrate has two facets: one (pterygoid condyle) is just above the medial condyle with a flat articular surface; the other (orbitopterygoid facet) is below the orbital process with a distinct groove for bone attachment, especially in *D. novaehollandiae*.

The mandibular process of the Casuariiformes quadrate has three condyles arranging a V-shape, and the lateral condyle is continuous with the caudal condyle. The medial condyle bears a prominent articular surface, whereas the lateral condyle shows an oval shape with a flat articular surface. The caudal condyle is well-developed in *C. casuarius* but shows a tiny tilt in *D. novaehollandiae*. A clear fossa/depression is located between medial and lateral condyles.

The quadrate body is relatively curved and bears a curved medial crest. In *D. novaehollandiae*, the lateral crest is relatively straight. In Casuariiformes, pneumatic foramina occur in two positions: both *C. casuarius* and *D. novaehollandiae* possess a pneumatic foramen (caudomedial foramen) on the caudal side of the quadrate body. Only *C. casuarius* has an additional pneumatic foramen (rostromedial foramen) on the rostromedial side of the quadrate body, below the otic process.

**Palaeognathae:** ^†^Dinornithiformes (^†^*Megalapteryx didinus*) (Plate 3)

Similar as most palaeognath quadrates, the intercapitular incisure between two capitula of the otic process is not distinct on *Megalapteryx didinus* quadrate (NHM A 16; Owen, 1883). Both squamosal and otic capitula have an oval outline with an equal in size. The squamosal capitulum faces dorsolaterally with a convex articular surface, while the otic capitulum is oriented dorsomedially with a flat articular surface.

The orbital process is massive with a high aspect ratio and bears a blunt and pointed tip, and it faces rostrally. The shape of the orbital process is similar as that of most palaeognath quadrates except for kiwi and tinamou.

The quadratojugal cotyle shows a deep fossa with a droplet-like margin, adjacent to the lateral condyle. Similar as most palaeognath quadrates (including Lithornithiformes, rhea, kiwi and some tinamou), the rostral margin of the quadratojugal cotyle is distinctly protruding for the jugal attachment in *Megalapteryx*is. In addition, the *Megalapteryx* quadrate has two articular facets with pterygoid: one (pterygoid condyle) is just above the medial condyle with a flat and smooth surface, facing laterally; the other (orbitopterygoid facet) is below the orbital process with a distinct but smooth surface for bone attachment. The articular with pterygoid on *Megalapteryx* quadrate shares associated feature with Casuariiformes and Tinamiformes: the orbitopterygoid facet attaches along the orbital process and faces medially (tinamiform-like feature), while the pterygoid condyle is located dorsally to the medial condyle with a smooth articular surface (casuariiform-like feature).

The mandibular process of the *Megalapteryx* quadrate has three condyles arranging a V or L shape and the lateral condyle is continuous with the caudal condyle. The medial condyle bears a rostrocaudally elongate oval shape with a prominent articular surface. The lateral condyle is relatively small with a smooth surface. The caudal condyle is well-developed, caudally projecting with a flat articular surface. A clear and huge fossa/depression is located between medial and lateral-caudal condyles.

The quadrate body is relatively straight due to a slightly curved medial and lateral crest. A huge pneumatic foramen (rostromedial foramen) is located in the rostromedial side of the quadrate body, below the otic process.

**Palaeognathae:** Tinamiformes (Plate 4)

On the otic process of most tinamou (Tinamiformes), there is not clear intercapitular incisure between two capitula, except in *Nothoprocta ornate* (Saiff, 1988). Both squamosal and otic capitula shows oval outline, but the squamosal capitulum is slightly larger in size than otic capitulum. The squamosal capitulum is rostrocaudally elongate, and it is oriented dorsally. The otic capitulum is lateromedially elongate with a flat articular surface, and it is oriented much mediodorsally.

The orbital process is highly variant in Tinamiformes: *Crypturellus tataupa* and *N. ornata* exhibit a slender orbital process with a high aspect ratio and a blunt, boot-like tip which points rostrodorsally (Bertelli et al., 2014). In contrast, *Eudromia elegans* exhibits a robust orbital process with a relatively lower aspect ratio and an arrow-like tip which points rostrally. The orbital crest, a muscle attachment laterally located at the orbital process, is present in *C. tataupa* and *N. ornata* with a linear ridge, but it is absent in *E. elegans*.

The quadratojugal cotyle bears a saddle shape (the dorsal and ventral margin of the contact has lost) with a shallow fossa, dorsally adjacent to the lateral condyle. At the dorsocaudal side of the quadratojugal cotyle, a significant protruding ridge extends to the medial side of the quadrate bodies. Unlike most avian quadrates, the pterygoid condyle in tinamou does not bear a ball-like process on the quadrate; instead, it is an extremely flat articular surface below the orbital process and faces medially.

The mandibular process on Tinamiformes quadrate has three condyles aligned in a U shape, and the lateral condyle is continuous with the caudal condyle. There is a clear intercondylar sulcus between medial condyle and lateral-caudal condyles on Tinamiformes quadrate (Saiff, 1988), especially in *E. elegans*. The medial condyle is prominent and much deepr than the other two condyles. In *C. tataupa* and *N. ornata*, the medial condyle is slender and rostrocaudally elongate with a diamond-shape outline, while in *E. elegans*. The medial condyle is mediolaterally elongate with a rectangular or oval in the outline. The lateral condyle is not well-developed and shows a tiny articular surface only. The caudal condyle shows a flat articular surface below the ridge abovementioned.

The quadrate body is relatively curved and slender. The position of the pneumatic foramen is variant among Tinamiformes (Saiff, 1988). In *C. tataupa*, one pneumatic foramen (postcapitular foramen) is caudally positioned between two capitula on the otic process, one (postotic capitulum foramen) is located below the otic capitulum, and the other foramen (rostromedial foramen) is rostromedially located below the otic process. In *E. elegans*, the otic process is highly pneumatic with pneumatic foramens caudally located at the squamosal, otic capitula and the position between them. In *N. ornata*, only postcapitular foramen is present.

**Pangalloanserae**: ^†^*Asteriornis maastrichtensis* (Plate 5)

The otic process of *Asteriornis* quadrate has a shallow but wide intercapitular incisure between two capitula (Field et al., 2020). The facet of both capitula is square- or diamond-shaped, and the facet of the squamosal capitulum is slightly larger than that of the otic capitulum. The squamosal capitulum has a lateraomedially convex surface, and thus, it is divided into rostral and lateral region. The rostral region of the squamosal capitulum is much larger than the lateral region, and rostrally slopes onto the squamosal capitulum. The otic capitulum has a flat and wide surface.

The orbital process remains broad up to the broken rostral end with dorsal slant, indicating that its orbital process might taper dorsally, not rostrally. The orbital crest, the mound on the lateral or ventral surface of orbital process, is at the ventral margin of the orbital process with an elongate shape.

Unlike most galloanseraens, the quadratojugal cotyle has a cup shape with a complete and thin circular margin and a deep articular fossa in *Asteriornis* (Field et al., 2020), though the ventrocaudal margin is slightly broken. The quadratojugal cotyle is separated from the lateral condyle with a small gap. The pterygoid condyle of *Asteriornis* quadrate is medially located at the ventral margin of the base of the orbital process, and it is also separated from the medial condyle by a distinct gap (Field et al., 2020). It is not rostrally protruding with a triangular shape.

Like other galloanseraen quadrates (Elzanowski and Stidham, 2010), the mandibular process in *Asteriornis* has two condyles, separated by a wide but distinct intercondylar sulcus. The medial condyle has a rostrocaudally elongate oval shape with a prominent concave surface, while the lateral condyle is round with a flat surface. The size of these two condyles is similar.

The quadrate body of *Asteriornis* is relatively straight and wide due to a straight medial and lateral crest. The subcapitular tubercle has a liner mound and is lateroventrally located on the margin of the squamosal capitulum, different with any extant galloanseraen quadrates. In caudal view, it seemingly merges with the dorsal part of the lateral crest. Two pneumatic foramina are present on the quadrate body: one (rostromedial foramen) is rostromedially located below the otic process, and the other (basiorbital foramen) is dorsally located at the pterygoid condyle (Field et al., 2020). It is worth to mention that the pneumatic foramen (caudomedial foramen) located at the caudomedial depression would be showed in specific range of setting in software VGStudio, and it might result from the similar density between avian quadrate and the sedimentary matrix.

^†^**Odontopterygiformes**: ^†^Pelagornithidae (^†^*Dasornis toliapica* and ^†^*Osteodontornis sp.*) (Plate 5)

The quadrate of Pelagornithidae is similar to avian quadrates with four different connections to neighbouring bones, but their shapes are significantly different with any other avian, even with their closest clade (Galloanserae).

The otic process does not show a clear intercapitular incisure between two capitula (*Lutetodontopteryx tethyensis* in Mayr and Zvonok, 2012 and *Osteodontornis sp.* in Ono, 1989). In contrast, in *Pelagornis chilensis* (Mayr and Rubilar-Roger, 2010) and *P. mauretanicus* (Mourer-Chauviré and Geraads, 2005), the intercapitular incisure is shallow but distinct. Below the otic process, a distinct notch shows between otic process and quadrate body in some Pelagornithidae, such as *Dasornis*, *Lutetodontopteryx*, and *Osteodontornis*. The morphological difference of the otic process between *Pelagornis* and other pelagornithid quadrates likely suggests a diversity of biomechanical function. The otic process without an intercapitular incisure might be effective in rostrocaudal movement, while the separated capitula with distinct intercapitular incisure is restricted in the gliding but much robust.

The squamosal capitulum of *Dasornis* quadrate is partially preserved, missing some lateral part of the facet, but it appears to exhibit an oval/rounded outline with a prominently lateromedially elongate articular surface, similar to that of other Pelagornithidae. The otic capitulum is generally rounded or oval with a flat articular surface; however, in several Pelagornithidae (e.g., *Dasornis*, *Lutetodontopteryx*, *Osteodontornis*, and *P. mauretanicus*), it is rostrocaudally bent. In these taxa, this articular surface can be subdivided into rostral and caudal portions, with the caudal portion being markedly larger than the rostral portion. In *P. chilensis*, the otic capitulum is slightly concave; however, whether it is subdivided into rostral and caudal portions remains uncertain due to poor preservation (Mayr and Rubilar-Roger, 2010).

The orbital process is broken in most of Pelagornithidae, and only *Osteodontornis* and *P. chilensis* persevered a complete orbital process. However, these taxa do not share a similar morphology. In *Osteodontornis*, the orbital process is triangular in lateral bview, relatively robust and tapers into a blunt tip, pointing mediorostrally (Ono, 1989). In *P. chilensis*, the orbital process is also triangular shape in lateral view but is proportionally longer and oriented more rostrally than in *Osteodontornis* (Mayr and Rubilar-Roger, 2010). The orbital crest, a mound located on the lateral or ventral surface of the orbital process in crown Galloanserae, is absent in all Pelagornithidae.

The quadratojugal cotyle is a cup-shaped, with a complete, thick circular margin and a deep fossa, and is distinctly separated from the the lateral condyle (Ono, 1989; Mourer-Chauviré and Geraads, 2008; Mayr and Rubilar-Roger, 2010). The pterygoid condyle is located at the ventral margin of the base of the orbital process and is well separated from the medial condyle. This articulation exhibits considerable morphological disparity among pelagornithid quadrates. In *Dasornis*, the pterygoid condyle has an oval outline with a minor ridge along its medial margin, forming a convex articular surface that closely fits the quadrate fossa on the pterygoid (Fig. S1). This feature has not been described in other pelagornithid quadrates and might be associated with the distinct lower jawbone. In *Osteodontornis*, the pterygoid condyle is not rostrally projected and bears a smooth articular surface, while in *P. chilensis* it is strongly projected rostrally (Mayr and Rubilar-Roger, 2010).

Similar to other galloanseran quadrates (Elzanowski and Stidham, 2010), the mandibular process of pelagornithid quadrate is bicondylar, consisting of rostrocaudally aligned medial and lateral condyles, separated by an intercondylar sulcus (Ono, 1989; Mourer-Chauviré and Geraads, 2008; Mayr and Rubilar-Roger, 2010). The medial condyle is elongated, while the lateral condyle is rounded or circular outline. In *Osteodontornis*, the medial condyle is remarkably slender and rostrocaudally elongate, and the lateral condyle is oval. This unusual shape likely is caused by taphonomy process, affecting the mandibular process of the quadrate.

The quadrate body in Pelagornithidae is relatively wide and robust and bears a straight lateral crest (Ono, 1989). Unlike in Galloanseres, the subcapitular tubercle – one of synapomorphic feature in this clade (Mayr and Clarke, 2003) – is absent in palgornithid quadrates. Two pneumatic foramina, rostromedial foramen and basiorbital foramen, are present in most Pelagornithidae quadrates (Ono, 1989; Mourer-Chauviré and Geraads, 2008; Mayr and Rubilar-Roger, 2010; Mayr and Zvonok, 2012) but not in *Dasornis* which appears to possess only the basiorbital foramen.

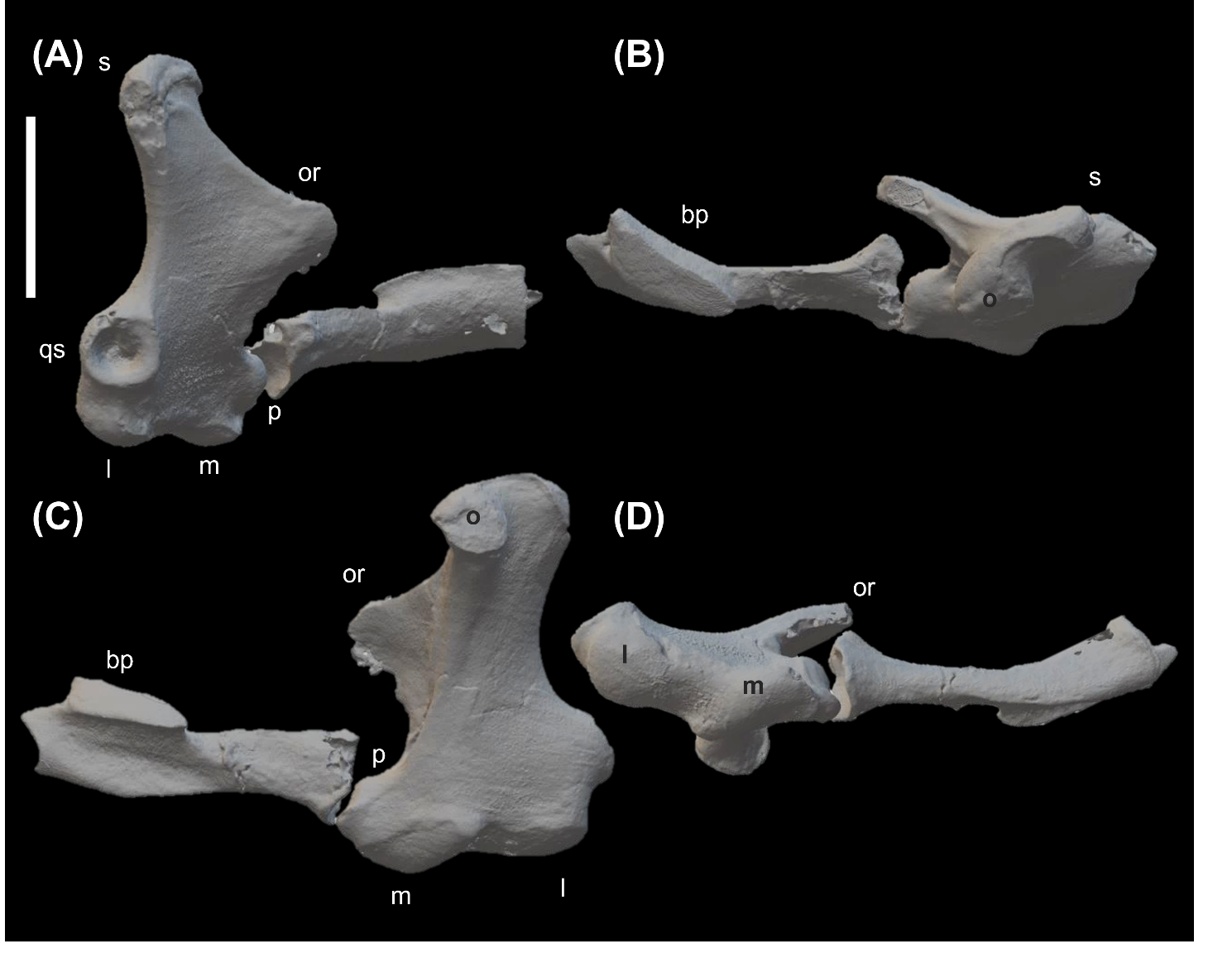

**Fig. S1** *Dasornis toliapica* right quadrate NHMUK PV 44096 with pterygoid (mirror) in (A) lateral, (B) dorsal, (C) medial, and (D) ventral view. Abbreviations: bp: basiopterygoid, l: lateral condyle, m: medial condyle, o: otic capitulum, or: orbital process, p: pterygoid condyle, qs: quadratojugal cotyle, s: squamosal capitulum. Scale bar: 10 mm.

**Galliformes**: Megapodiidae (Plate 6)

The quadrate of Megapodiidae has two capitula with a shallow and wide intercapitular incisure between them. The squamosal capitulum is obviously higher than the otic capitulum in rostral view but with equal size to each other. In most Megapodiidae, the squamosal capitulum has a diamond or square-like flat facet facing rostrally or except *Macarocephalon maleo*. In *Macarocephalon maleo*, the facet of squamosal capitulum is convex, separating into rostral and caudal parts: the rostral part of the facet faces rostrally and is much larger than the caudal part, while the caudal part faces dorsally. The otic capitulum has a flat square-like or rectangle with round corners shape, pointing dorsomedially.

Like other Galliformes, the orbital process in Megapodiidae is elongated and slender with a round/pointed tip, pointing rostrally or rostroventrally. The orbital crest, which is located at the ventral or lateral side of the orbital process, has a linear mound-like shape.

The quadratojugal cotyle of Megapodiidae quadrate is a cup shape with a deep articular fossa and thick circular margin (Elzanowski and Stidham, 2010), adjacent to the lateral condyle. The dorsal and ventral margins of the quadratojugal cotyle are not slightly well-developed. Like other Galliformes, the pterygoid condyle in Megapodiidae has two articulation contact: one (orbitopterygoid facet) locates the base of the orbital process with a flat articular surface facing medially; the other (pterygoid condyle) is above the medial condyle with a distinct gap (Elzanowski and Stidham, 2010). The pterygoid condyle in megapodiids is protruding and has two different types of shape: *Alectura lathami* and *Megapodius reinwardt* have a round or ball-like facet, facing rostrally. On the other hand, *Leipoa ocellata* and *M. maleo* has an oval shape of facet with a concave articular surface, facing rostrolaterally.

The mandibular process on Megapodiidae quadrate has two condyles, rostrocaudally lining up. The medial condyle is much smaller than the lateral condyle, and there is a clear but narrow intercondylar sulcus between them (Elzanowski and Stidham, 2010). Both medial condyle and lateral condyle have relatively smooth articular surfaces.

The quadrate body in Megapodiidae is relatively curved and slender, and it has an obvious medial and tympanic crest. Apparently, a mound-like subcapitular tubercle is located rostroventrally at the squamosal capitulum with an oval shape. In the caudal view, there is a shallow caudomedial depression between two capitula, compassed by medial and tympanic crests. In Megapodiidae, there are two pneumatic foramens on the quadrate body, medially located at the quadrate body (Elzanowski and Stidham, 2010). One (rostromedial foramen) is below the otic capitulum, while the other (basiorbital foramen) is above the pterygoid condyle, just between the base of the orbital process and the pterygoid condyle.

**Galliformes**: Cracidae (Plate 6 and 7)

On the otic process in Cracidae, a clear intercapitular incisure is present between squamosal capitulum and otic capitulum. In *Crax mitu* (*Mitu mitu*), *Ortalis ruficauda*, and *Pipile cumanensis*, the intercapitular incisure is narrow, while the intercapitular incisure in *Pipile pipile* is much wider. The squamosal capitulum is obviously higher and larger than the otic capitulum, especially *Crax mitu*. The squamosal capitulum is oval and rostrocaudally convex, dividing into rostral and caudal part of the facet. The rostral part of the facet faces rostrally, while the caudal part faces dorsally. The otic capitulum has a flat articular surface, pointing dorsomedially except *Crax mitu* whose oitc capitulum points more medially.

The orbital process in most Cracidea is elongate with a flat and slant tip, pointing rostrally. In *Pipile cumanensis*, its tip is relatively sharp, but in other Cracidae, their tips are round or blunt. The orbital crest is also a linear mound-like shape, similar as other Galliformes.

Unlike Megapodiidae, the quadratojugal cotyle of Cracidea quadrate becomes saddle-like shape due to noticeably inward ventral and dorsal margin of quadratojugal cotyle, especially the ventral margin of the quadratojugal cotyle (Elzanowski and Stidham, 2010). It has a thick margin, adjacent to the lateral condyle. Like other Galliformes, the pterygoid condyle in Cracidea has two articulation contact: the orbitopterygoid facet is located at the base of the orbital process with a flat articular surface, facing medially; the pterygoid condyle is above the medial condyle with a distinct and shallow gap. Their pterygoid condyle is not as protruding as Megapodiidae and it has a slightly dorsoventrally convex articular surface, facing laterally in *Pipile cumanensis* or medially in other Cracidea.

The mandibular process on Cracidea quadrate has two condyles, lining up lateromedially. The medial condyle is much smaller than the lateral condyle, and there is a clear and deep intercondylar sulcus between them. Also, unlike Megapodiidae, the medial condyle expands caudodorsally in Cracidea, making it tilted. On the other hand, the lateral condyle is slightly convex.

In most Cracidea, the quadrate body is relative curved and slender, and it has an obvious medial crest, except in *Pipile cumanensis* which has a much wider quadrate body. The mound-like subcapitular tubercle is located below the otic process, and it is slender in *Crax mitu* and *Ortalis ruficauda* but not in *Pipile pipile* nor *Pipile cumanensis* which of that is much wider. Like other Galliformes, a shallow caudomedial depression caudally presents between two capitula, compassed by medial and tympanic crests, but it moves more medially by the curved tympanic crest. Unlike any other Galliformes, there is a bulge caudally attached to the quadrate body, called submeatic process. It is only found in cracid, some Anseriformes quadrate (mentioned below), *Sarothrura elegans*, *Picus viridis*, and Falconiformes, and it might be apormorphic to Cracidae. The basiorbital foramen presents on all Cracidae quadrates in this study. The rostromedial foramen shows on the quadrates of *Crax mitu* and *Pipile pipile* only, and is medially located at the quadrate body, encompassed by the medial crest and the orbital process. On the *Pipile cumanensis* quadrate, the rostromedial fossa is located at the same position of the rostromedial foramen on the two species abovementioned, and thus, it is likely able to exchange air between cranial air sac and the internal quadrate body.

**Galliformes**: Numididae (Plate 7)

In Numididae, a shallow and narrow intercapitular incisure presents between two capitula on the otic process. The squamosal capitulum is apparently higher than the otic capitulum in rostral view, and the otic capitulum becomes relatively smaller. The squamosal capitulum is round and convex and faces dorsally, while the otic capitulum is oval with a flat articular surface, pointing medially.

The orbital process in Numididae is elongate with high aspect ratio and pointed tip, facing mediorostrally. Like other Galliformes, the orbital crest also has a mound-like shape.

The quadratojugal cotyle of Numididae quadrate becomes saddle shape due to noticeably inward ventral and dorsal margin of quadratojugal cotyle. This joint connection has a thick margin and is close to the lateral condyle with a small gap. It is worth to mention that the caudal margin of the quadratojugal cotyle become caudally elongate. Like other galliform quadrates, the articular connection with pterygoid in Numididae could be divided into two articulation contact: the orbitopterygoid facet is medially located close to the base of the orbital process, and it has a distinct articular surface facing medially; the pterygoid condyle is dorsally located at the medial condyle. Its pterygoid condyle is blunt and has an an up-side down triangle shape with a flat articular surface, facing more rostrolaterally.

The mandibular process on Numididae quadrate has two condyles, rostrocaudally lining up. Like Cracidea, the medial condyle is much smaller than the lateral condyle, and there is a clear and deep intercondylar sulcus between them. Also, the medial condyle expands caudodorsally, making it tilted. The lateral condyle is gently convex.

The quadrate body of Numididae is relative curved and slender and it has a distinct medial crest. The mound-like subcapitular tubercle is ventrally located below the otic process, but more rostral position than Cracidae or Megapodiidae. Also, comparing to Megapodiidae and Cracidae, its shape is much wider with a concave articular surface. Like other Galliformes, a shallow caudomedial depression caudally presents between two capitula. In Numididae, two pneumatic foramina are medially located at the quadrate body: one (rostromedial foramen) is located below the otic capitulum; the other (basiorbital foramen) is just above the pterygoid condyle, close to the base of the orbital process.

**Galliformes**: Odontophoridae (Plate 7)

The morphology of quadrates in Odontophoridae have two different types of shape, strongly driven by phylogeny. In Odontophoridae, only *Ptilopachus petrosus* (Ptilopachinae) shares morphological similarity with Numididae and Phasianidae, while *Colinus virginianus* and *Odontophorus guttatus* (Odontophorinae) evolves their unique features (apomorphy). On the otic process of *Ptilopachus petrosus*, a clear but shallow intercapitular incisure presents between two capitula. On the other hand, the intercapitular incisure in Odontophorinae is either unclear (*Colinus virginianus*) or absent (*Odontophorus guttatus*) due to the fact that the otic capitulum in *Odontophorus guttatus* disappear (Elzanowski and Stidham, 2010). Like other Galliformes, the squamosal capitulum is apparently higher than the otic capitulum in rostral view, and also it is much larger than otic capitulum. The squamosal capitulum has a slightly mediolaterally elongate oval shape in *Ptilopachus petrosus* and *Colinus virginianus* with a convex articular surface, but it is rounded with a convex articular surface in *Odontophorus guttatus*. The otic capitulum in *Ptilopachus petrosus* and *Colinus virginianus* is round with a flat articular surface, pointing in mediodorsal direction.

The orbital process in Odontophoridae shows a huge range of morphological disparity. In *Ptilopachus petrosus*, the orbital process is slender and elongate with a pointed tip, similar as that of Phasianidae (see next section). In *Colinus virginianus*, the orbital process is robust at the base but turns to be slender at the rostral portion with a pointed tip, facing rostrally. In *Odontophorus guttatus*, the orbital process is robust with a clear ridge (orbital crest) at the lateral side and a sharp tip, pointing dorsally. The orbital crest, a mound-like muscle attachment, is also located at the ventral margin or lateral surface of the orbital process.

The quadratojugal cotyle of *Ptilopachus petrosus* looks closer to that of Numididae or Phasianidae, which has a saddle-like joint connection with a deep fossa. For other Odontophoridae quadrates (*Colinus virginianus* and *Odontophorus guttatus*), their shapes of the quadratojugal cotyle look much similar to Megapodiidae, which still have a relative complete ellipse margin with a slightly inward dorsal or ventral margin (Elzanowski and Stidham, 2010). This joint connection has a thick margin and deep fossa, adjacent to the lateral condyle.

Like other Galliformes, the connection to pterygoid on quadrates in Odontophoridae also has two articulations: the orbitopterygoid facet in *Ptilopachus petrosus* is medially located at the ventral margin of the orbital process with an oval shape and a smooth articular surface. For *Colinus virginianus* and *Odontophorus guttatus*, it is also medially located at the ventral margin of the orbital process, but its position is more rostral than *Ptilopachus petrosus*, far from the base of the orbital process. The pterygoid condyle is rostrally protruding with an upside-down triangle shape outline and a convex articular surface in *Ptilopachus petrosus*. Its pterygoid condyle is also dorsally located at the medial condyle with a significant gap. In Odontophorinae, the pterygoid condyle is not as standing as its sister group (*Ptilopachus petrosus*), and it has an oval shape, facing rostrally. It is dorsally adjacent to the medial condyle in *Colinus virginianus*, but still with a short gap in *Odontophorus guttatus*.

The mandibular process on Odontophoridae quadrate has two condyles, rostrocaudally lining up in *Ptilopachus petrosus* and *Colinus virginianus* but lateromedially lining up in *Odontophorus guttatus*. The medial condyle is significantly smaller than the lateral condyle in *Ptilopachus petrosus*, but it only slightly smaller in Odontophorinae. Also, a clear and wide intercondylar sulcus between medial and lateral condyle. The medial condyle is convex and rostrocaudally elongate, but it caudodrsally expands in *Ptilopachus petrosus* only. For the lateral condyle, Odontophorinae has a flat articular surface, and *Ptilopachus petrosus* has a slightly convex articular surface.

The quadrate body of Odontophoridae is relatively straight and slender, and it has an obvious medial crest. The mound-like subcapitular tubercle is located below the otic process (below the squamosal capitulum), similar as other Galliformes. However, comparing with other non-Phasianidae galliform, it moves more rostrally and is more protruding, especially on *Odontophorus guttatus* quadrate. A caudomedial depression is unclear on Odontophorinae quadrate, but it is a small but deep caudomedial depression on *Ptilopachus petrosus* quadrate. The pneumaticity in Odontophoridae quadrate varies taxon to taxon. In *Odontophorus guttatus*, there is no pneumatic foramen present on the quadrate body. In *Ptilopachus petrosus*, only one pneumatic foramen (rostromedial foramen) is medially located below the otic process. In *Colinus virginianus*, two pneumatic foramina are medially located at the quadrate body: one (rostromedial foramen) is medially located below the otic process, and the other (basiorbital foramen) is close to the base of the orbital process and pterygoid condyle.

**Galliformes**: Phasianidae (Plate 8-10)

Generally speaking, the morphology of Phasianidae quadrates look similar to Numididae and *Ptilopachus petrosus*; however, it still shows a great morphological diversity on different characters, such as otic process, pterygoid condyle, and subcapitular tubercle. For example, on the otic process, all Phasianidae has two capitula except for *Rollulus rouloul* whose otic capitulum is underdeveloped, same as *Odontophorus guttatus*. In most phasianids, an unclear and shallow intercapitular incisure between squamosal and otic capitulum, such as *Arborophila torqueola*, *Phasianus colchicus*, *Gallus varius*, *Gallus gallus*, *Peliperdix coqui* (*Campocolinus coqui*), and *Coturnix coturnix*. In some species (e.g., *Tragopan satyra*, *Meleagris gallopavo*, *Perdix cinerea*, and *Pavo cristatus*), the intercapitular incisure is absent between two capitula (Elzanowski and Stidham, 2010), while in *Lophophorus impejanus*, *Bonasa umbellus*, and *Meleagris gallopavo*, there is a clear intercapitular incisure between two capitula. Like other clades in Galliformes, the squamosal capitulum is apparently higher than the otic capitulum in rostral view, and much larger than otic capitulum. The squamosal capitulum has a rostrocaudally elongate oval shape with a convex articular surface and faces dorsally in most Phasianidae, but in *Rollulus rouloul* and *Coturnix coturnix*, their squamosal capitula have a round shape with a convex articular surface. The otid capitulum in Phasianidae is round with a flat articular surface, pointing medially or mediodorsally. In *Lophophorus impejanus*, one eclipse flat facet with a flat articular surface is laterally located at the squamosal capitulum. This facet does not articulate with the cranium, and as a result, it might be the muscle attachment.

The orbital process in most Phasianidae is slender and elongate with a pointed tip (Elzanowski and Stidham, 2010), facing either rostrodorsally, rostrally, or ventrally (*Pavo cristatus*), except *Lophophorus impejanus*. In *Lophophorus impejanus*, the orbital process is also elongate, but its tip is blunt. The orbital crest is located at different positions taxa by taxa in Phasianidae: in most Phasianidae, it is located at either close to the base of the orbital process (*Rollulus rouloul*, *Coturnix coturnix*, *Tragopan satyra*, and *Meleagris gallopavo*) or the centre position of the orbital process (*Pavo cristatus*, *Gallus varius*, *Gallus gallus*, *Lophophorus impejanus*, *Bonasa umbellus*, *Perdix cinerea*, and *Phasianus colchicus*). In *Arborophila torqueola* and *Peliperdix coqui*, the orbital crest is at the ventral margin of the orbital process.

Similar to Numididae and *Ptilopachus petrosus*, the quadratojugal cotyle of Phasianidae quadrate has a saddle-like shape with a deep fossa (Elzanowski and Stidham, 2010), and its caudodorsal margin caudally expands, except *Coturnix coturnix* whose quadratojugal cotyle has a shallow fossa without the elongate caudal margin. This articular connection in most Phasianidae is distinct to the lateral condyle with a small gap, but this gap is much larger in several species, such as *Coturnix coturnix*, *Lophophorus impejanus*, and *Phasianus colchicus*. Like other Galliformes, the articular connection with pterygoid on quadrate in Phasianidae also has two portions: one (orbitopterygoid facet) is medially located at the ventral margin of the orbital process with a distinct contact (e.g., *Arborophila torqueola*, *Rollulus rouloul*, *Pavo cristatus*, *Gallus gallus*, *Coturnix coturnix*, *Lophophorus impejanus*, *Meleagris gallopavo*, *Bonasa umbellus*, and *Phasianus colchicus*), or with a smooth articular surface (e.g., *Gallus varius*, *Peliperdix coqui*, *Tragopan satyra*, and *Perdix cinerea*); the other (pterygoid condyle) is not as protruding as Megapodiidae (except *Lophophorus impejanus*) and has an oval shape (e.g., *Arborophila torqueola*, *Rollulus rouloul*, *Pavo cristatus*, *Gallus gallus*, *Coturnix coturnix*, *Lophophorus impejanus*, and *Meleagris gallopavo*) or an up-side-down triangle shape (e.g., *Gallus varius*, *Peliperdix coqui*, *Tragopan satyra*, *Meleagris gallopavo*, *Perdix cinerea*, and *Phasianus colchicus*), with a distinct gap to the medial condyle (Elzanowski and Stidham, 2010).

The mandibular process in Phasianidae has two condyles, mostly lining up rostrocaudally except *Arborophila torqueola* whose two condyles line up more lateromedially.

The medial condyle is significantly smaller than the lateral condyle, and there is a clear and wide intercondylar sulcus between them. The medial condyle has a curved and convex articular surface, and it is caudodorsally elongate. The lateral condyle, on the other hand, also has a convex articular surface.

The quadrate body in most Phasianidae is relative slender, but it has a much wider body in *Lophophorus impejanus* only. The mound-like subcapitular tubercle is ventrally located at the otic process, but based on its shape, it can be grouped into three different types in Phasianidae: in most Phasianidae, it is relatively thin, elongate and standing, such as *Arborophila torqueola*, *Rollulus rouloul*, *Pavo cristatus, Peliperdix coqui*, *Coturnix coturnix*, *Tragopan satyra*, and *Bonasa umbellus*. In some taxon (e.g., *Gallus gallus, Gallus varius, Meleagris gallopavo, Phasianus colchicus*), its shape is oval or round, and finally, in *Lophophorus impejanus* and *Perdix cinerea*, the subcapitular tubercle looks more like that of Anseriformes whose subcapitular tubercles is protruding and platform-like with a bulbed surface. In most Phasianidae, there is a clear and shallow caudomedial depression, but it is either absent or not well-developed in some species, like *Arborophila torqueola*, *Rollulus rouloul*, *Tragopan satyra*, *Lophophorus impejanus*, and *Gallus gallus*. In Phasianidae, one or two pneumatic foramina on the quadrate body but not in *Tragopan satyra* whose quadrate does not have any pneumatic foramina. The rostromedial foramen is medially located at the quadrate body, encompassed by the medial crest and the orbital process on most Phasianidae quadrates except *Perdix cinerea* quadrate, while the basiorbital foramen is close to the base of the orbital process and pterygoid condyle in *Meleagris gallopavo*, *Perdix cinerea*, *Pavo cristatus*, and *Gallus gallus*.

**Anseriformes**: Anhimidae (Plate 10)

On the otic process in Anhimidae, both two capitula are far from each other without a clear and deep intercapitular incisure between them. The squamosal capitulum is obviously higher than otic capitulum in rostral view, and it also has larger size than the otic capitulum. In most Anhimidae (except *Anhima cornuta*), the squamosal capitulum has a lateromedial elongate oval or rectangle shape with round corners and it has a convex articular surface, diving into rostral and caudal portion. The rostral portion is larger than the caudal portion. In *Anhima cornuta*, the squamosal capitulum does not divide into two portions, and it faces rostrally, instead of dorsolaterally. The otic capitulum has a square/diamond shape with a flat articular surface, pointing dorsomedially.

The orbital process in Anhimidae has a triangle shape in lateral view with different types of tips: in *Anhima cornuta*, the orbital process is relatively thin, and its tip is pointed, facing rostrally. In *Chauna sp.* (UMZC 291.A, *Chauna chavaria*) and *Chauna chavaria* (OUMNH 23790), their tip is blunt, facing rostrally, while in *Chauna torquata*, it has a flat tip. The orbital crest, which is located at the ventral side of the orbital process in Anhimidae, is a linear ridge-like shape, especially in *Chauna torquata*.

The quadratojugal cotyle of Anhimidae quadrate has a saddle-like shape without clear dorsal and ventral margin of the articular connection, close to the lateral condyle. In *Anhima cornuta*, the margin of the quadratojugal cotyle is thin, but in other Anhimidae (e.g., *Chauna*), it is relatively thicker. Unlike their sister group (Galliformes), the pterygoid condyle in Anhimidae has only one articular contact (pterygoid condyle) above the medial condyle with a distinct distance (Elzanowski and Stidham, 2010). The pterygoid condyle in *Chauna* is protruding with an oval shape. However, in *Anhima cornuta*, it is blunt with a slightly concave articular surface.

The mandibular process on Anhimidae quadrate has two condyles, rostrocaudally lining up. The medial condyle is much smaller than the lateral condyle, and there is a clear but narrow intercondylar sulcus between them. The medial condyle has a slightly flat articular surface in *Anhima cornuta*, but it has a convex articular surface in *Chauna*. On the other hand, the lateral condyle has a convex articular surface in all Anhimidae.

The quadrate body of Anhimidae is relatively slender in *Anhima cornuta* only, and others are obviously much wider. The subcapitular tubercle in Anhimidae has a standing platform-like shape, ventrally located at the otic process. Nevertheless, compared with Galliformes, the subcapitular tubercle is located much rostrally in Anhimidae. Similar to Cracidae, the submeatic process has either a flat protrusion in *Anhima cornuta*, or a bulge standing protrusion in *Chauna* caudodorsally attached with the quadratojugal cotyle (Elzanowski and Stidham, 2010). This unique character might provide a muscle attachment. In Anhimidae quadrate, only one elongate and slender pneumatic foramen (caudomedial foramen) is medially located at the quadrate body (Elzanowski and Stidham, 2010), far from the otic process and it faces medially.

**Anseriformes**: Anseranatidae (Plate 11)

The otic process in Anseranatidae (*Anseranas semipalmata*) has a clear and deep intercapitular incisure between squamosal and otic capitulum. Its squamosal capitulum is obviously higher than the otic capitulum in rostral view, but they have equal size. The squamosal capitulum has a square or diamond shape with a smooth convex articular surface, dividing into rostral and caudal parts: the rostral part is much larger than caudal part. The otic capitulum has an rostrocaudally elongate oval shape with a smooth articular surface, facing dorsomedially.

The orbital process in *Anseranas semipalmata* has a triangle shape in lateral view with a pointed tip, pointing rostromedially. The orbital crest is a small ridge with a V-like or U-like shape, expanding dorsally and ventrally, and it is laterally located at the surface of the orbital process. In the medial side of the orbital process, there is a deep basiorbital fossa in *Anseranas semipalmata*, different with Galliformes and Anhimidae.

The quadratojugal cotyle of *Anseranas semipalmata* quadrate has an oval shape with a deep fossa, adjacent to the lateral condyle. Different with other Anseriformes (Anhimidae and most Anatidae, mentioned below), the submeatic process is absent in *Anseranas semipalmata*. The pterygoid condyle has a standing rounded shape with a smooth articular surface, just above the medial condyle with a small gap, and it faces rostrally.

The mandibular process on *Anseranas semipalmata* quadrate has two condyles, rostrocaudally lining up. The medial condyle is much smaller than the lateral condyle, and there is a clear but narrow intercondylar sulcus between them. The medial condyle is rostrocaudally elongate and has a concave articular surface, while the lateral condyle has a slightly concave articular surface and a laterally distinct margin.

Like most Anseriformes (*Chauna* and most Anatidae), the quadrate body of Anseranatidae is much wider with a straight medial and lateral crest. However, unlike other clades in Anseriformes, the subcapitular tubercle in *Anseranas semipalmata* is either absent, extremely tiny close to squamosal capitulum, or merged with the squamosal capitulum. In *Anseranas semipalmata*, there is a clear shallow and deep caudomedial depression on the quadrate body. One pneumatic foramen (caudomedial foramen) medially displays on the quadrate body, and it faces more caudomedially than that does on Anhimidae quadrates.

**Anseriformes**: Anatidae (Plate 11-14)

The quadrates of Anatidae mostly share similar characters with each other, except some specific adaptors, like *Mergus merganser*. The otic process in all Anatidae has a distinct and shallow intercapitular incisure between squamosal and otic capitulum, but in some taxon (e.g., *Dendrocygna bicolor* and *Merganetta armata*), it has a deep intercapitular incisure. The squamosal capitulum is apparently higher than the otic capitulum in rostral view and mostly has an equal or slightly larger size than otic capitulum, except *Mergus merganser*. In *Mergus merganser*, the otic capitulum is significantly smaller than squamosal capitulum, and faces dorsally instead of medially or mediodorsally. The squamosal capitulum has a rostrocaudally elongate oval shape with a convex articular surface in most Anatidae, but no in *Sarkidiornis melanota* which has a concave surface. The shape of the otic capitulum shows a great variance in Anatidae. For instance, the otic capitulum has a rostrocaudally elongate oval shape in most Anatidae, such as *Thalassornis leuconotus*, *Dendrocygna bicolor*, *Oxyura jamaicensis*, *Ceropsis novaehollandiae*, *Cygnus olor*, *Branta bernicla*, *Anser fabalis*, *Merganetta armata*, *Plectropterus gaubensis*, *S. melanotos*, *Tachyeres brachptera*, *Anas platyrhynchos*, and *Anas aucklandica*. It has a rounded shape in some species (e.g., *Biziura lobata*, *Clangula hyemalis*, *Mergus merganser*, and *Mergellus albellus*), or a square/diamond shape in *Melanitta nigra*, *Histrionicus histrionicus*, *Chenonetta jubata*, and *Aythya farina*. The otic capitulum faces either dorsally or mediodorsally in most Anatidae.

Like their sister group, Anseranatidae (*Anseranas semipalmata*), the orbital process in most Anatidae has a triangle shape in lateral view with a robust base, except *Mergus merganser*. The orbital process in *Mergus merganser* is much slender and elongate but with a robust base and a rounded tip. The tip of the orbital process in other Anatidae is either pointed or blunt facing different directions. For instance, it points rostrally in most of Anatidae, such as *Thalassornis leuconotus*, *Malacorhynchus membranaceus*, *Branta bernicla*, *Mergellus albellus*, *Plectropterus gaubensis*, *Sarkidiornis africana*, *Aythya ferina*, and *Anas platyrhynchos*. In other Anatidae, like *Biziura lobata*, *Ceropsis novaehollandiae*, *Anser fabalis*, *Clangula hyemalis*, *Histrionicus histrionicus*, *Chenonetta jubata*, and *Sarkidiornis melanotos*, the tip of the orbital process points rostromedially, while in some Anatidae (e.g., *Dendrocygna bicolor*, *Oxyura jamaicensis*, *Cygnus olor*, *Tachyeres brachptera*, and *Anas aucklandica*), it points rostrodorsally. Surprisedly, the tip of orbital process in *Melanitta nigra* points dorsomedially, and its points medially in *Merganetta armata*, completely different with other Anatidae. Unlike Galliformes, the orbital crest in most Anatidae has a distinct ridge with a triangle shape on the lateral surface of the orbital process and it dorsally and ventrally extends, making it a V- or U-shape (Elzanowski and Stidham, 2010). However, in some Anatidae, it has its unique shape of the orbital crest, such as *Oxyura jamaicensis* (a thick and robust linear ridge), *Ceropsis novaehollandiae* (a thin linear ridge), *Anser fabalis* (a bulbed-like ridge), *Clangula hyemalis* (a bulbed-like ridge), *Mergus merganser* (a bulbed-like ridge), *Merganetta armata* (a thick and robust linear ridge), *Sarkidiornis melanotos* (a rounded ridge), and *Sarkidiornis africana* (an oval ridge). Usually, the basiorbital fossa, which is a depression at the medial side of the orbital process, is relatively deeper in Anatidae than in Galliformes, except *Mergus merganser*.

The quadratojugal cotyle in most Anatidae quadrates is similar with each other and *Anseranas semipalmata*: they all have an oval outline with a deep fossa. In other Anatidae, it has an incomplete outline: the dorsal margin is absent in *Branta bernicla*, and the ventral margin is not well-developed in *Ceropsis novaehollandiae*, *Mergellus albellus*, and *Sarkidiornis africana*. In *Mergus merganser* and *Merganetta armata*, its dorsal and ventral margin of the quadratojugal cotyle is absent. The relative position between quadratojugal cotyle and lateral condyle is variant in Anatidae. In most Anatidae, the quadratojugal cotyle is close to the lateral condyle with a small gap. The quadratojugal cotyle is adjacent to the lateral condyle in *Thalassornis leuconotus* and *Cygnus olor*, while it is far from the lateral condyle in *Dendrocygna bicolor*, *Ceropsis novaehollandiae*, and *Sarkidiornis melanotos*.

The pterygoid condyle in Anatidae shows a great morphological variance in shape and its relative position to medial condyle. In most Anatidae, it has an oval shape with a convex or slightly convex articular surface, and it is not protruding and dorsally located at the medial condyle with a distinct or small gap (Elzanowski and Stidham, 2010), such as *Biziura lobata*, *Malacorhynchus membranaceus*, *Ceropsis novaehollandiae*, *Cygnus olor*, *Anser fabalis*, *Mergellus albellus*, *Sarkidiornis africana*, *Tachyeres brachptera*, *Anas platyrhynchos*, and *Anas aucklandica*. In *Histrionicus histrionicus*, it has an oval shape with a flat articular surface, and though it is rostrally blunt, it is far from the medial condyle with a clear gap. In *Chenonetta jubata*, it has an oval shape with a convex articular surface and a lateromedial ridge, separating into dorsal and rostral parts. Its pterygoid condyle is not protruding and is only above the medial condyle with a small gap. In other Anatidae, the pterygoid condyle is rounded with a convex or slight convex articular surface, and it is not rostrally protruding. It is dorsally located at the medial condyle with a distinct or small gap, such as *Dendrocygna bicolor*, *Branta bernicla*, *Clangula hyemalis*, *Plectropterus gaubensis*, and *Aythya farina*. In *Thalassornis leuconotus* and *Mergus merganser*, their pterygoid condyles are rounded with convex articular surfaces and adjacent to the medial condyle. In other dataset, three Anatidae have distinct morphological features on the pterygoid condyle, and they are totally different with others: In *Oxyura jamaicensis*, its pterygoid condyle has a rounded shape with a convex articular surface. It is much rostrally protruding than any other Anatidae and is far from the medial condyle with a clear distance. In *Merganetta armata* and *Melanitta nigra*, their pterygoid condyles are close to the medial condyle with a small gap, but they have a distinguishable shape: in *Merganetta armata*, it has an up-side-down triangle with a slightly convex articular surface, while it has a diamond shape with a flat articular surface in *Melanitta nigra*.

The mandibular process on Anatidae quadrate has two condyles, rostrocaudally lining up, except *Mergus merganser* and *Mergellus albellus* whose condyles lateromedially line up and are parallel to each other with a deep and wide intercondylar sulcus between them. The medial condyle is much smaller than the lateral condyle in most Anatidae, but not in *Malacorhynchus membranaceus*, *Anser fabalis*, *Clangula hyemalis*, *Melanitta nigra*, *Histrionicus histrionicus*, *Mergus merganser*, *Merganetta armata*, and *Aythya farina* whose medial condyle has equal or larger size (*Merganetta armata* only) with lateral condyle. The intercondylar sulcus is much wider in Anatidae. The medial condyle is elongate and has a rounded or oval outline with either a concave aritcular surface in most Anatidae, or slightly concave aritcular surface in other taxon, such as *Dendrocygna bicolor*, *Malacorhynchus membranaceus*, *Clangula hyemalis*, and *Sarkidiornis*. In *Merganetta armata*, the medial condyle has a much flat aritcular surface, different with other Anatidae. The lateral condyle in Anatidae is also elongate and has an oval outline with a concave articular surface. Two condyles have distinct ridges with the quadrate body.

Like most Anseriformes (*Chauna* and *Anseranas semipalmata*), the quadrate body in Anatidae is much wider and has a relatively straight medial and lateral crest. The subcapitular tubercle in most Anatidae is located more rostrally and has a platform-like surface for muscle attachment, but not in *Thalassornis leuconotus*, *Dendrocygna bicolor*, *Merganetta armata*, *Ceropsis novaehollandiae*, *Biziura lobata*, *Mergus merganser*, and *Clangula hyemalis*. In *Thalassornis leuconotus*, the subcapitular tubercle is morphologically similar as *Anseranas semipalmata* whose subcapitular tubercle is located laterally at the squamosal capitulum. In *Dendrocygna bicolor* and *Merganetta armata*, the subcapitular tubercles is close to the otic process and is much laterally located. In *Ceropsis novaehollandiae*, however, its subcapitular tubercles is much similar as most Galliformes with a mound-like shape. In *Biziura lobata*, and *Mergus merganser*, its subcapitular tubercle has a small, tiny and bulb shape. Finally, in *Clangula hyemalis*, the subcapitular tubercle is located laterally. Like their sister group, *Anseranas semipalmata*, a clear and shallow caudomedial depression is caudally or medially located at the quadrate body in all Anatidae. Similar as Anhimidae, the submeatic process is caudodorsally attached to the quadratojugal cotyle with a clear ridge in most Anatidae (Elzanowski and Stidham, 2010). In some Anatidae (e.g., *Biziura lobata*, *Malacorhynchus membranaceus*, *Sarkidiornis melanotos*, and *Tachyeres brachptera*), it is not well-developed with a tiny but clear crest only. The submeatic process in *Mergus merganser* is absent. Like all other Anseriformes, a single and large pneumatic foramen (caudomedial foramen) is caudomedially located at the quadrate body (Elzanowski and Stidham, 2010) and it is between medial and tympanic crest, facing much medially than *Anseranas semipalmata* on most Anatidae quadrates, but not on *Malacorhynchus membranaceus* (facing caudally), *Merganetta armata* (relatively small pneumatic foramen), and *Plectropterus gaubensis* (facing caudomedially) quadrates. The postcapitular foramen, caudally located at the otic process, is only found on *Malacorhynchus membranaceus* and *Melanitta nigra* quadrates.

**Capmarginulgiformes**: Capmarginulgidae (Plate 15)

The otic process in Capmarginulgidae does not have a clear clear intercapitular incisure between squamosal and otic capitulum, and each capitulum is far from each other. The squamosal capitulum also has either the same height as its otic capitulum or is slightly higher only in *Capmarginulgus europaeus*. It has a much smaller size of the squamosal capitulum than its otic capitulum. The squamosal capitulum in Capmarginulgidae has an oval shape with a convex articular surface and faces dorsolaterally, while the otic capitulum also has an oval shape with a slight convex articular surface but faces dorsomedially.

The orbital process in Capmarginulgidae is strongly reduced and short with a sharp tip (Cracraft, 1981; Mayr, 2002), pointing rostrally (*Eurostopodus mysticalis* and *Chordeiles minor*) or rostrodorsally (*Capmarginulgus europaeus*).

The quadratojugal cotyle of Capmarginulgidae quadrate is much laterally protruding, and it has a cup shape with a clear margin and a shallow articular fossa, adjacent to the lateral condyle. This shallow fossa faces more laterodorsally instead of laterally, similar as Aegothelidae (*Aegotheles cristatus*), and Trogonidae (*Trogon melanurus* and *Trogon collaris*). The pterygoid condyle in Capmarginulgidae is continuous with base of the orbital process. It has an oval shape with a convex articular surface, and it far from the medial condyle in Capmarginulgidae.

Similar as most Capmarginulgiformes (except Steatornithidae and Podargidae) and some Apodiformes (Hemiprocnidae and Apodidae), the mandibular process on Capmarginulgidae quadrate has two condyles (medial and lateral condyles) only, lateromedially lining up with a clear and deep intercondylar sulcus (furrow-like). It is uncertain whether the third condyle, caudal condyle, exists or not, because there is still a tiny protrusion on Capmarginulgidae quadrate in the caudal view. If this is considered as a caudal condyle, it means that caudal condyle in most Capmarginulgiformes is strongly reduced (Bϋhler, 1970; Mayr, 2002). The medial condyle has an oval shape and is much rostrocaudally elongate a standing convex articular surface. The lateral condyle has a rounded or squared shape with a flat articular surface.

The quadrate body of Capmarginulgidae is medially curved with an obvious curved medial crest and straight lateral crest. The caudomedial depression between two capitula is wide but shows a deep fossa surrounded by tympanic and medial crest. The pneumatic foramen (basiorbital foramen), medially located at quadrate body, exhibits on all Capmarginulgidae quadrates in this study. It is larger on *Eurostopodus mysticalis* and *Capmarginulgus europaeus* quadrates, but relatively smaller on *Chordeiles minor* quadrate. In *Eurostopodus mysticalis*, another pneumatic foramen (rostromedial foramen) is rostromedially present at the quadrate body, and one fossa (caudomedial fossa) caudally present at quadrate body. In *Capmarginulgus europaeus*, one fossa (caudomedial fossa) caudally present at quadrate body, compressed by medial and tympanic crest.

**Steatornithiformes:** Steatornithidae (Plate 15)

In *Steatornis caripensis*, the otic process has a clead and wide intercapitular incisure between squamosal and otic capitulum. Like most avian quadrates, the squamosal capitulum is slightly higher than the otic capitulum. However, unlike other avian quadrates, the squamosal capitulum in *Steatornis caripensis* is bifurcated, meaning that the squamosal capitulum has two articular surfaces with the cranium: lateral and medial portion. These two parts all have the lateromedial elongate oval shape with equal size. The lateral portion of the squamosal capitulum has a convex articular surface, while the medial portion has a flat articular surface. The otic capitulum also a lateromedial elongate oval shape with a convex articular surface, pointing dorsomedially.

Unlike most Capmarginulgidae, the orbital process in Steatornithidae is well-developed and elegant with a slant and flat tip (Mayr, 2002), pointing rostrally. Besides, *Steatornis caripensis* has a broad and shallow basiorbital fossa.

The quadratojugal cotyle of *Steatornis caripensis* quadrate is much laterally protruding and has a cup shape with a clear margin and shallow joint connection. It faces more laterally and is adjacent to the lateral condyle. Different with Capmarginulgidae, the pterygoid condyle in *Steatornis caripensis* is far from the base of the orbital process. It is not protruding but has an oval shape and close to the medial condyle with a small gap.

Unlike most Capmarginulgiformes, *Steatornis caripensis* quadrate has three condyles on its mandibular process, with a L or tick shape. The medial condyle is the largest and standing condyle, while the lateral condyle is the smallest and flat condyles. The medial condyle has a rostrocaudally elongate oval shape with a standing convex articular surface. The lateral condyle has an oval shape with a flat articular surface, and the caudal condyle has a bulged- like shape caudally attached with the quadrate body.

The quadrate body of *Steatornis caripensis* is curved with an obviously curved medial and lateral crest. In Steatornithidae, only one pneumatic foramen (rostromedial foramen) is medially located at the quadrate body, and it is just below the otic capitulum.

**Nyctibiiformes:** Nyctibiidae (Plate 15)

The quadrate of Nyctibiidae (*Nyctibius griseus*) looks morphologically similar as Capmarginulgidae. For instance, the otic process does not have a well-developed intercapitular incisure between two capitula, and each capitulum is far from each other. The squamosal capitulum is higher than the otic capitulum, but it is much smaller than the otic capitulum, different with Capmarginulgidae. The squamosal capitulum has an oval shape with a convex articular surface, facing dorsally, while the otic capitulum also has an oval shape with a standing convex articular surface, facing dorsomedially.

Like some Capmarginulgiformes (e.g., Capmarginulgidae and *Aegotheles cristatus*), the orbital process in *Nyctibius griseus* is strongly reduced and short with a sharp tip (Cracraft, 1981; Mayr, 2002; Zusi, 2013), pointing rostrally.

The quadratojugal cotyle of Nyctibiidae quadrate is much caudolaterally protruding, and it has a saddle shape (without a clear dorsal and ventral margin) with a shallow joint connection. It is adjacent to the lateral condyle and faces more caudolaterally, instead of laterally or laterodorsally. The pterygoid condyle in *Nyctibius griseus* is continuous with base of the orbital process, and it has an oval shape with a convex aritcular surface. Its pterygoid condyle is also far from the medial condyle, same as nightjar (Capmarginulgidae).

The mandibular process on Nyctibiidae quadrate has two condyles (medial and lateral condyles), rostrocaudally lining up with a clear and deep groove (intercondylar sulcus) between them. Similar as Capmarginulgidae, the caudal condyle is either absent or strongly reduced on *Nyctibius griseus* quadrate (Mayr, 2002). The medial condyle has a rostrocaudally elongate oval shape with a standing convex articular surface. The lateral condyle has a squared shape with a flat articular surface.

The quadrate body of Nyctibiidae is medially curved with an obvious curved medial crest and straight lateral crest. In the caudal view, there is a distinct shallow caudomedial depression between two capitula. In *Nyctibius griseus*, two pneumatic foramina are medially located at the quadrate body – one (basiorbital foramen) has a relatively huge opening and is above the pterygoid condyle, like all Capmarginulgidae. The other (rostromedial foramen) is just above the aforesaid pneumatic foramen.

**Podargiformes:** Podargidae (Plate 15)

The otic process in Podargidae has a clear and wide intercapitular incisure between squamosal and otic capitulum, and each capitulum is far from each other. The squamosal capitulum is slightly higher than otic capitulum in *Podargus strigoides* in rostral view, and also has a much larger size than otic capitulum. The squamosal capitulum has a kidney shape (curved in the lateral margin of the facet) with a convex articular surface, facing dorsolaterally, while the otic capitulum has an oval shape with a flat articular surface, facing dorsomedially.

The orbital process in Podargidae has an elongate and triangle shape in lateral view with a sharp tip (Mayr, 2002), pointing rostrally. Besides, *Podargus strigoides* has a broad and shallow basioorbtail fossa.

The quadratojugal cotyle of Podargidae quadrate is much laterally protruding, and it has a cup shape with a clear margin and a deep but small joint connection. It is adjacent to the lateral condyle and faces more caudolaterally. The pterygoid condyle in Podargidae is distinct with the orbital process, and it has an oval outline and a slightly convex articular surface. Unlike other Capmarginulgiformes, the pterygoid condyle is close to the medial condyle with a small gap in Podargidae.

Similar as *Steatornis caripensis*, the mandibular process on Podargidae quadrate has three condyles with a hook or J shape in ventral view. Unlike other Capmarginulgiformes, there is a deep but wide intercondylar sulcus between medial and lateral condyle and a clear depression at the centre of the mandibular process. The medial condyle has a rostrocaudally elongate rectangle shape with a standing convex articular surface. The lateral condyle is continuous with the caudal condyle, and it has an oval shape with flat articular surface. The caudal condyle is only a tiny protrusion caudally attached with quadrate body.

The quadrate body of Podargidae is curved with an obvious curved medial crest. In the caudal view, there is a shallow and wide caudomedial depression between two capitula. In Podargidae, there are two pneumatic foramens on the quadrate body – one (rostromedial foramen) is medially located at the quadrate body, and the other (caudomedial foramen) is located at the caudomedial depression.

**Aegotheliformes:** Aegothelidae (Plate 16)

Generally speaking, the quadrate of Aegothelidae (*Aegotheles cristatus*) shares a great similarity with nightjar (Capmarginulgidae) quadrate in shape of the orbital process and quadrate body. However, there are still several differences between them. For instance, in Aegothelidae, the otic process has a shallow intercapitular incisure between squamosal and otic capitulum, and each capitulum is far from each other. The squamosal capitulum is higher than otic capitulum in rostral view, but it is much smaller than otic capitulum. Different with Capmarginulgidae, the squamosal capitulum in *Aegotheles cristatus* has a rounded shape with a rostrocaudally convex articular surface, separating into rostral and caudal portion. The otic capitulum has a slender and elongate oval shape with a flat articular surface, similar as *Podargus strigoides*.

Similar as Capmarginulgidae, the orbital process in Aegothelidae is strongly reduce (Cracraft, 1981; Mayr, 2002; Zusi, 2013) and short with a sharp tip, pointing rostrodorsally. Besides, the basiorbital fossa is much shallower in *Aegotheles cristatus* than *Steatornis caripensis* or *Podargus strigoides*.

The quadratojugal cotyle in Aegothelidae is not as laterally protruding as Capmarginulgidae, and it has a saddle shape (the dorsal and ventral margin is not well-developed) with an extremely shallow joint connection. It is adjacent to the lateral condyle, similar as *Nyctibius griseus*. This shallow fossa faces more laterodorsally instead of laterally, like Capmarginulgidae or Trogonidae (*Trogon melanurus* and *Trogon collaris*). The pterygoid condyle is far from the base of the orbital process, and it has an oval shape with a convex articular surface (Zusi, 2013). Comparing with other Capmarginulgiformes, the pterygoid condyle is adjacent to the medial condyle in Aegothelidae quadrate.

The mandibular process on Aegothelidae quadrate has two condyles, rostrocaudally lining up with a clear and deep intercondylar sulcus between them. Like Capmarginulgidae and Nyctibiidae, the caudal condyle might be either absent or reduced (Bϋhler, 1970). The medial and lateral condyle have equal size with similar shape – a rostrocaudally elongate and slender shape with a standing convex articular surface.

The quadrate body in Aegothelidae is medially curved with an obvious curved medial crest and straight lateral crest. There is a distinct caudomedial depression between two capitula. The quadrate of Aegothelidae is highly pneumatic: several pneumatic foramina (e.g., postsquamosal capitulum foramen, postotic capitulum foramen, and postcapitular foramen) caudally present at the otic process (Mayr, 2002). Also, another two pneumatic foramina are medially located at the quadrate body. One (basiorbital foramen) is medially close to the base of the orbital process, and the other (rostromedial foramen) is below the otic process.

**Apodiformes:** Hemiprocnidae (Plate 16)

In Hemiprocnidae (*Hemiprocne comata*), the otic process has a shallow intercapitular incisure between two capitula (squamosal and otic capitulum). Comparing with Capmarginulgiformes, each capitulum is relatively close to each other. The squamosal capitulum is with the same height as the otic capitulum but is much smaller than the otic capitulum. The squamosal capitulum in Hemiprocnidae has a rounded shape with a convex articular surface, facing dorsolaterally, while the otic capitulum also has a rostocaudally elongate oval shape with a flat articular surface, facing dorsomedially.

Like some Capmarginulgiformes (e.g., Capmarginulgidae and Aegothelidae), the orbital process in Hemiprocnidae is strongly reduced and short with a blunt tip (Cracraft, 1981; Mayr, 2002; Zusi, 2013), pointing rostrally.

The quadratojugal cotyle in Hemiprocnidae is much caudolaterally protruding. It has a rounded shape with a flat articular surface without a distinct margin, and it is adjacent to the lateral condyle. This flat joint faces more laterodorsally, instead of laterally. The pterygoid condyle in Hemiprocnidae has an elongate oval shape with a convex articular surface, and it is close to the medial condyle with a small gap.

Similar as most Capmarginulgiformes (except Steatornithidae and Podargidae) and some Apodiformes (Apodidae, mentioned below), the mandibular process on Hemiprocnidae quadrate has two condyles (medial and lateral condyles), lateromedially lining up with a clear and deep furrow-like intercondylar sulcus. The caudal condyle, however, is either absent or underdeveloped (Zusi, 2013). Like all Capmarginulgiformes, the medial condyle is much larger than the lateral condyle. The medial condyle and the lateral condyle all have an oval shape and is much rostrocaudally elongate with a standing convex articular surface (Zusi, 2013).

The quadrate body of Hemiprocnidae is medially curved with an obvious curved medial crest and straight lateral crest. There is a distinct caudomedial depression between two capitula. In Hemiprocnidae, there are two pneumatic foramina: one (postsquamosal capitulum foramen) is caudally located at the squamosal capitulum, and the other (postcapitular foramen) is caudally located at the midpoint of the otic process.

**Apodiformes:** Apodidae (Plate 16)

The quadrate of Apodidae is morphologically similar as that of Hemiprocnidae (*Hemiprocne comata*) on its overall shape, orbital process, quadratojugal cotyle, pterygoid condyle, and mandibular process. However, their quadrates still have many characters different from that of Hemiprocnidae. For instance, the intercapitular incisure in Apodidae only shows in *Streptoprocne zonaris* (shallow but relatively wide), not in *Chaetura brachyura*. Different with the otic process in Hemiprocnidae (*Hemiprocne comata*), the otic capitulum in swift (Apodidae) is much larger than the squamosal capitulum. The squamosal capitulum is with the same height as the otic capitulum in *Chaetura brachyura* but it is much higher than otic capitulum in *Streptoprocne zonaris*. The squamosal capitulum in Apodidae has a lateromedial elongate oval articular surface with a standing convex articular surface. The otic capitulum, on the other hand, has a rostrocaudally elongate oval shape with a flat or slightly convex articular surface, facing dorsally (*Streptoprocne zonaris*) or dorsomedially (*Chaetura brachyura*). A clear notch is rostroventrally located at the otic capitulum in Apodidae (it is much clearer in *Chaetura brachyura*).

Like some Capmarginulgiformes (e.g., Capmarginulgidae and Aegothelidae) and Hemiprocnidae, the orbital process in Apodidae is strongly reduced and short with a blunt tip (Cracraft, 1981; Mayr, 2002; Zusi, 2013), pointing rostrally. Besides, Apodidae has a clear but shallow basiorbital fossa.

The quadratojugal cotyle of Apodidae quadrate is caudolaterally protruding and is adjacent to the lateral condyle. Similar as *Hemiprocne comata*, it has a round shape with a flat articular surface and without a clear margin. This flat articulation faces more laterodorsally, instead of laterally. The pterygoid condyle in Apodidae is elongate and standing with a convex articular surface and it is close to medial condyle with a small gap.

The mandibular process on Apodidae quadrate has two condyles, rostrocaudally lining up with a clear and wide furrow-like intercondylar sulcus. Like Capmarginulgiformes, the medial condyle is much larger than the lateral condyle. The medial condyle has a rostrocaudally elongate oval shape with a standing convex articular surface, and it has a clear edge with the medial side of the quadrate body. The lateral condyle also has a rostrocaudally elongate oval shape with a slightly convex articular surface.

The quadrate body of Apodidae is medially curved due to an obvious curved medial crest and straight lateral crest. In the caudal view, only *Chaetura brachyura* has a distinct caudomedial depression between two capitula. In Apodidae, the pneumatic foramina are caudally located at the otic process (Cracraft, 1981; Mayr, 2002; Zusi, 2013): postsquamosal capitulum foramen and postcapitular foramen are present in *Streptoprocne zonaris* and *Chaetura brachyura*, while the postotic capitulum foramen is present in *Streptoprocne zonaris* only. In *Chaetura brachyura* only, the pneumatic foramina (basiorbital foramen) are medially located at the quadrate body, close to the pterygoid condyle with two openings.

**Apodiformes:** Trochilidae (Plate 16 and 17)

Hummingbirds (Trochilidae) have distinguishable quadrates with any other Apodiformes or Capmarginulgiformes, and it might result from their unique ecology and feeding behaviour.

On the oitc process in Trochilidae, the intercapitular incisure between squamosal and otic capitulum is either shallow (*Phaethornis superciliosus* and *Patagona gigas*), deep (*Colibri coruscans* and *Archilochus colubris*) or absent in *Topaza pyra*. The otic capitulum usually has an equal size with the squamosal capitulum in most hummingbirds, but it is much larger in *Topaza pyra* and *Colibri coruscans*. The squamosal capitulum has the same height as otic capitulum in Trochilidae quadrates. The shape of the squamosal capitulum in Trochilidae show a huge morphological diversity: it has a lateromedially elongate oval shape with a convex articular surface, facing dorsolaterally in *Topaza pyra*. It has a rostrocaudally elongate oval shape with a flat articular surface, facing rostrodorsally in *Colibri coruscans*. In *Phaethornis superciliosus*, the squamosal capitulum has a round shape with a flat articular surface, facing rostrodorsally. Finally, in *Archilochus colubris* and *Patagona gigas*, it has a dorsoventrally elongate oval shape with a slightly convex articular surface and faces dorsolaterally. The otic capitulum in Trochilidae has a rostrocaudally elongate oval shape (*Topaza pyra* has a much slender facet than other hummingbirds) with a flat articular surface and faces dorsomedially. Similar as Apodidae (e.g., *Chaetura brachyura*), a clear notch is rostroventrally located at the otic capitulum.

Unlike their sister group (Apodidae), the orbital process is robust with a lower aspect ratio and blunt tip, point rostrally in most hummingbirds (Zusi, 2013), but not in *Patagona gigas*, which points more rostroventrally instead. The basiorbital fossa is much smaller in *Colibri coruscans* and *Archilochus colubris* than in other hummingbirds.

The quadratojugal cotyle in Trochilidae quadrate is slightly caudolaterally protruding and it has a distinct articulation: the margin of the quadratojugal cotyle is not complete, losing the dorsal and ventral margin and the caudal margin extends laterally, making its shape like a notch, especially in *Phaethornis superciliosus* and *Archilochus colubris*. The quadratojugal cotyle is relatively deep and is close to the lateral condyle. The pterygoid condyle in hummingbirds is round and rostrally standing with a convex articular surface (Zusi, 2013), and it is close to the medial condyle with a small gap.

The mandibular process on Trochilidae quadrate has two condyles, rostrocaudally lining up with a clear and wide furrow-like intercondylar sulcus between them. Same most Striosres, the caudal condyle is absent or does not well-developed (Mayr, 2002; Zusi, 2013). Unlike Capmarginulgiformes or other Apodiformes, the medial condyle is equal or slightly smaller than the lateral condyle, and it is only larger than the lateral condyle in *Archilochus colubris* and *Patagona gigas*. The medial condyle has a rostrocaudally elongate oval shape with a standing convex articular surface, while the lateral condyle has a lateromedially elongate oval shape with a slightly convex articular surface.

The quadrate body in Trochilidae is not as curved as other Strisores due to a straight medial and lateral crest. The caudomedial depression is clearly located at the caudal side of the otic process. Like Apodidae, the Trochilidae quadrate is highly pneumatic, especially on its otic process (Cracraft, 1981; Mayr, 2002; Zusi, 2013). The postsquamosal capitulum foramen and postotic capitulum foramen are present in all hummingbird quadrates. The postcapitular foramen which is caudally located at the middle position between two capitula is only present in *Phaethornis superciliosus*. Also, the rostromedial foramen is medially located at some hummingbird quadrate body, such as *Colibri coruscans*, *Archilochus colubris*, and *Patagona gigas*.

**Musophagiformes:** Musophagidae (Plate 18)

In Musophagidae (*Corythaeola cristata*), the otic process has a shallow and wide intercapitular incisure, and each capitulum is far from each other. The squamosal capitulum is as high as the otic capitulum in rostral view and it has an equal size with the otic capitulum. The squamosal capitulum in Musophagidae has a lateromedially elongate oval shape with a convex articular surface, facing laterodorsally. The otic capitulum has a water-droplet outline with a convex articular surface, facing mediodorsally.

The orbital process in *Corythaeola cristata* is robust with a slight high aspect ratio and has a blunt, slant ant flat tip, pointing mediorostrally. Besides, Musophagidae has a shallow but distinct basiorbital fossa.

The quadratojugal cotyle of Musophagidae quadrate is ventrolaterally protruding, and it has a cup-like shape with a thick margin and deep joint. It is adjacent to the lateral condyle, and unlike most avian quadrate, this shallow fossa faces more ventrolaterally, instead of laterally. The pterygoid condyle in *Corythaeola cristata* is dorsally located at the medial condyle with a small but distinct gap, and it has an oval shape with a flat articular surface.

The mandibular process on Musophagidae quadrate has three condyles arranging in a L shape in ventral view. It has a clear fossa at the centre of the mandibular process. The medial condyle has a rostrocaudally elongate oval shape with a standing convex articular surface. The lateral condyle is not distinct with the caudal condyle, and it has a flat articular surface. The caudal condyle is not well-developed, and it is only a tiny tip caudally attached to the quadrate body.

The quadrate body in Musophagidae is relatively straight with a straight medial and lateral crest. The caudomedial depression is caudally located at the quadrate body. In Musophagidae, three pneumatic foramina are present on the quadrate – one (caudomedial foramen) is located at caudomedial depression. Another (basiorbital foramen) is medially located at the quadrate body, close to the pterygoid condyle. The other (postcapitular foramen) is caudally located at the otic process, at the middle position between two capitula.

**Otidiformes:** Otididae (Plate 18)

In Otididae (*Ardeotis australis*), the otic process also has a shallow but narrow intercapitular incisure, and compared with Musophagidae (*Corythaeola cristata*), each capitulum is close to each other. The squamosal capitulum is as high as the otic capitulum in rostral view and it has an equal size with the otic capitulum. Both capitula have a lateromedially elongate oval shape with a convex articular surface, facing laterodorsally and mediodorsally, respectively.

The orbital process in *Ardeotis australis* is robust with a slight high aspect ratio and has a slant ant flat tip, pointing rostrodorsally. A distinct ridge is located at the lateral surface of the orbital process, extending form the tip of the orbital process to the base of this process. Besides, Musophagidae has a shallow but distinct and broad basiorbital fossa.

The quadratojugal cotyle in Otididae quadrate is slightly laterally protruding and it is separated from the lateral condyle with a small gap. This articulation has an oval shape with a thin margin and a deep fossa. On the rostral or rostroventral margin of the quadratojugal cotyle, there is a flat articular surface for jugal attachment. The pterygoid condyle in *Ardeotis australis* is not standing but it is above the medial condyle with a clear gap. Also, it has a relatively large oval shape with a slightly convex articular surface.

Like Musophagidae, the mandibular process on Otididae quadrate has three condyles with a L shape in ventral view; however, unlike Musophagidae, there is a clear furrow-like intercondylar vallecula between medial and lateral-caudal condyle. The medial condyle has a rostrocuadally elongate oval shape with a standing convex articular surface, and it also has a clear margin with the quadrate body. The lateral condyle is continuous with the caudal condyle, and it has an oval shape with a slightly convex articular surface. The caudal condyle is a distinct bulge-like structure caudally protruding and attached with quadrate body.

The quadrate body in Otididae is relative slender and medially curved with a curved medial crest. A shallow caudomedial depression is caudally located at the otic process. Below the squamosal capitulum, the subcapitular tubercle is rostrally present with an oval shape and a flat articular surface. In Otididae, only one pneumatic foramen (caudomedial foramen) is medially located at the caudomedial depression.

**Cuculiformes:** Cuculidae (Plate 18 and 19)

In Cuculidae, the otic process has a shallow and clear intercapitular incisure between two capitula but not in *Coccyzus americanus* whose intercapitular incisure is much deeper than any other Cuculidae. The squamosal capitulum is obviously higher than the otic capitulum in rostral view, especially in *Coccyzus americanus* and *Hierococcyx fugax*. The squamosal capitulum has a larger shape than the otic capitulum in most Cuculidae, but it is much smaller than the otic capitulum in *Hierococcyx fugax*. The squamosal capitulum in Cuculidae shows a great morphological diversity: it has a lateromedially elongate oval shape with a slightly convex articular surface in *Crotophaga ani* and faces more rostrodorsally. In *Tapera naevia* and *Centropus goliath*, the squamosal capitulum has a rounded shape with a flat (*Tapera naevia*) or convex articular surface (*Centropus goliath*), facing more rostrodorsally, but a significant notch is located between the squamosal capitulum and the quadrate body only on *Centropus goliath*. In *Coccyzus americanus* and *Hierococcyx fugax*, the squamosal capitulum has a lateromedially elongate oval shape with a convex surface, facing more dorsolaterally. Also, a clear notch is ventrally located at the squamosal capitulum. The otic capitulum in Cuculidae usually has an oval shape with a flat articular surface, facing mediodorsally, but not in *Centropus goliath*. In *Centropus goliath*, the otic capitulum has an oval shape with a convex articular surface and it faces more caudodorsally.

The orbital process in Cuculidae is much thick and slender with a high aspect ratio, and its tip is a flat, blunt and slant, pointing mediorostrally. However, its tip has a standing hammer-like or boots-like shape in *Centropus goliath*. Besides, there is a shallow basiorbital fossa in Cuculidae.

Based on the shape of the articular connection, the quadratojugal cotyle in Cuculidae could be divided into two types: in *Crotophaga ani*, *Tapera naevia*, and *Centropus goliath*, it has a complete and distinct circular margin with a deep fossa, facing laterally. It is close to the lateral condyle with a small gap. In *Coccyzus americanus* and *Hierococcyx fugax*, the quadratojugal cotyle has an oval margin with a flat articular surface, facing laterally. It is adjacent to the lateral condyle. The pterygoid condyle in Cuculidae is rostrally standing but not in *Coccyzus americanus* and *Hierococcyx fugax*. It has an oval shape with a convex articular surface, dorsally located at the medial condyle with a deep but small gap.

The mandibular process on Cuculidae quadrate has three condyles, arranging in a L shape in ventral view. A shallow fossa is located at the centre of the mandibular process. The medial condyle has an equal or smaller size than the lateral condyle (Posso and Donatelli, 2001). The medial condyle has a rostrocaudally elongate oval shape with a standing convex articular surface, and it has a distinct margin with the quadrate body. The lateral condyle has an elongate oval shape with a smooth articular surface (Posso and Donatelli, 2001), and it is continuous with caudal condyle in most Cuculidae but not in *Hierococcyx fugax*. The caudal condyle is a bulge-like protrusion with a flat articular surface caudally attached with the quadrate body.

The quadrate body in Cuculidae is extremely curved with a curved medial, lateral and tympanic crest. In the caudal view, a distinct caudomedial depression is present between two capitula with a clear caudal fossa facing medially. In Cuculidae, the pneumatic foramen shows on three different positions on quadrate body. One is caudally located at the otic process, either at squamosal (postsquamosal capitulum foramen), otic capitulum (postotic capitulum foramen), or the middle between two capitula (postcapitular foramen) in *Tapera naevia*, *Centropus goliath*, and *Hierococcyx fugax*. Another pneumatic foramen (rostromedial foramen) is medially located at the quadrate body surrounded by the orbital process and medial crest on the quadrate of *Crotophaga ani*, *Tapera naevia*, and *Hierococcyx fugax*. The other (caudomedial foramen) medially exhibits at caudomedial depression on most Cuculidae quadrates.

**Mesitornithiformes:** Mesitornithidae (Plate 19)

On the quadrate of Mesitornithidae (*Monias benschi*), its otic process has a shallow and narrow intercapitular incisure between two capitula. The squamosal capitulum has the same height and an equal size as the otic capitulum. The squamosal capitulum in Mesitornithidae has a rounded shape with a convex articular surface, facing rostrodorsally, while the otic capitulum has an oval shape with a flat articular surface, facing mediodorsally.

The orbital process in *Monias benschi* is thick and robust with a high aspect ratio, and its tip is flat and slant, poiting rostrally or mediorostrally. Besides, the basiorbital fossa in Mesitornithidae is relatively deep and broad.

The quadratojugal cotyle in Mesitornithidae is slightly caudolaterally protruding and adjacent to the lateral condyle. It has a cup-like shape with a thick margin and deep articular connection. The pterygoid condyle in *Monias benschi* is above the medial condyle with a distinct gap, and it is rostrally protruding with an oval shape and a flat articular surface.

The mandibular process on Mesitornithidae quadrate has three condyles arranging a L shape in ventral view, and a shallow fossa is located at the centre of the articulation. The medial condyle has a lateromedially elongate oval shape with a standing convex articular surface (lateral trochlea). The lateral condyle is continuous with the caudal condyle, and it does not have a clear shape with a smooth articular surface. The caudal condyle is thick bulge-like structure caudally attached with the quadrate body.

In Mesitornithidae, the quadrate body is wide and relatively straight with a straight medial and lateral crest. The caudomedial depression is wide and deep, and a clear caudomedial fossa, which might be related to avian pneumaticity, is accommodated in the caudomedial depression. In Mesitornithidae, two pneumatic foramina are located at the quadrate body – one (postsquamosal capitulum foramen) is caudally located at the squamosal capitulum, and the other (rostromedial foramen) is medially located at the quadrate body and is above the basiorbital fossa.

**Pterocliformes:** Pteroclidae (Plate 19)

In Pteroclidae (*Syrrhaptes paradoxus*), the otic process has a shallow but wide intercapitular incisure between squamosal and otic capitulum, and each capitulum is far from each other. The squamosal capitulum has the same height and an equal size as the otic capitulum. The squamosal capitulum has a rounded shape with a slight convex articular surface, facing rostrodorsally, while the otic capitulum has a rounded shape with a flat articular surface, facing mediodorsally.

The orbital process in *Syrrhaptes paradoxus* is robust with a high aspect ratio, facing rostrally, and it has a broad tip. The dorsal margin of the tip expands laterally, and a ventral margin of the tip expands rostroventrally, forming a sting-like structure. In *Syrrhaptes paradoxus*, its basiorbital fossa is broad and shallow, rostrally located at the orbital process.

The quadratojugal cotyle in Pteroclidae has a cup-like shape with a thick and margin and deep articulation, adjacent to the lateral condyle. Unlike most avian quadrates, the quadratojugal cotyle faces more rostrolaterally, instead of laterally. A thick and lateromedially elongate bulge is caudodorsally attached to the quadratojugal cotyle. The pterygoid condyle in *Syrrhaptes paradoxus* is above the medial condyle with a distinct gap. It is not rostrally protruding or standing and has an oval shape with a flat articular surface.

The mandibular process on Pteroclidae quadrate has three condyles, arranging in a U shape. Also, there is a shallow fossa between lateral and medial condyle. The medial condyle has a rostrocaudally elongate oval shape with a standing convex articular surface. The lateral condyle also has a rostrocaudally elongate oval shape, but it only has a flat articular surface. The caudal condyle is not well-developed and has a flat articular surface.

The quadrate body in Pteroclidae is wide at the otic process but slender at the rest of the body due to a curved medial and lateral crest. In Pteroclidae, two pneumatic foramina are medially located at the quadrate body – one (rostromedial foramen) is ventrally located at the otic process, and the other (basiorbital foramen) is ventromedially located at the base of the orbital process, close to the pterygoid condyle.

**Columbiformes:** Columbidae (Plate 19 and 20)

On the otic process, Columbidae quadrates mostly shows a wide and relatively deep intercapitular incisure between squamosal and otic capitulum, but not on dodo quadrate (*Raphus cucullatus*). On the otic process of dodo quadrate, the intercapitular incisure is shallow or absent between two capitula. The squamosal capitulum has the same height and an equal size as the otic capitulum in most Columbidae, but it is much larger than otic capitulum in dodo and Large green pigeon (*Treron capellei*). Also, the otic capitulum of the Rock dove (*Columba livia*) quadrate is much higher than the squamosal capitulum. The squamosal capitulum in most Columbidae has a rostrocaudally elongate oval shape with a convex articular surface, except dodo. The squamosal capitulum of the dodo quadrate has a rounded shape with a convex articular surface. The otic capitulum also has a rostrocaudally elongate oval shape with a slight convex or flat articular surface, facing mediodorsally. However, the otic capitulum on the dodo quadrate appears a round or square outline with a convex articular surface.

Based on the shape variance, the orbital process of Columbidae quadrate could be grouped into two types: The first one shows a robust orbital process with a slightly high ratio and blunt tip, pointing rostrodorsally, such as Dodo. The second type shows a slender and elongate orbital process with a high aspect ratio. Its tip mostly is round or blunt, but the tip of the oribtal process is pointed on the Grey-fronted dove (*Leptotila rufaxilla*) quadrate. On some Columbidae quadrate (e.g., Large green pigeon and Rock dove), a flat crest or ridge is laterally located at the orbital process, close to the tip. The basiorbital fossa is relatively shallow and broad on most Columbidae quadrate; however, it is deep and broad on the dodo quadrate and deep and narrow on the Large green pigeon quadrate, respectively.

In Columbidae, the dorsal or ventral margin of the quadratojugal cotyle is usually not well-developed with a shallow articular surface, shown in a saddle-like shape. However, the quadratojugal cotyle of dodo quadrate exhibits a cup-like shape with a circular margin and shallow articular surface. The quadratojugal cotyle of Columbidae quadrate generally faces laterally but it only faces more rostrolaterally on the Rock dove quadrate. The articular connection to the pterygoid on Columbidae quadrates usually has two portions, pterygoid condyle and orbitopterygoid facet, and these two articulations are close to each other. However, the orbitopterygoid facet is absent on the quadrate of dodo, Grey-fronted dove, and the Tooth-billed pigeon (*Didunculus strigirostris*). The pterygoid condyle in Columbidae is usually adjacent to the medial condyle with a distinct distance, such as Tooth-billed pigeon, Rock dove, and Grey-fronted dove. The pterygoid condyle is not rostrally standing and mostly appears to be a round shape with a flat articular surface. However, on the quadrate of the Dodo and Large green pigeon, the pterygoid condyle shows an oval outline, while the pterygoid condyle is rostrally standing on the Grey-fronted dove quadrate.

Surprisedly, the mandibular process on Columbidae quadrate does not have the similar scenario as their closest relatives (e.g., Mesitornithidae and Pterocliformes) or Otidimorphae – it only has two condyles (medial and lateral condyle) lateromedially lining in with a wide and shallow intercondylar sulcus but not in Dodo. On the Dodo quadrate, these two condyles array rostrocaudally. The medial condyle has a rostrocaudally elongate oval shape with a standing convex articular surface in most Columbidae, but it has a relatively flat articular surface on the Rock dove quadrate only. On the other hand, the medial condyle of the Tooth-billed pigeon significantly elongates rostrocaudally, similar as that of parrot (Psittaciformes, see dorsal view of parrot quadrates in Plate 41-42). The lateral condyle exhibits a rounded shape with a flat articular surface on Dodo and Grey-fronted dove quadrate, while it exhibits an oval shape with flat articular surface on other Columbidae quadrates. On the mandibular process of the Tooth-billed pigeon quadrate, the lateral condyle significantly elongates rostrocaudally with a convex articular articular surface.

The quadrate body of Columbidae is curved due to a curved medial crest and lateral crest (but found on the Rock dove and Grey-fronted dove only). The caudomedial depression, caudally located at the quadrate body, is shallow and unclear. In Columbidae, avian pneumatic foramen is located at four positions of the quadrate body. The rostromedial foramen is medially located at the quadrate body, encompassed by the medial crest and the orbital process on the quadrate of Dodo and Grey-fronted dove. The basiorbital foramen is medially located at the orbital process, close to the pterygoid condyle and basiorbital fossa on the Black-chinned fruit dove (*Ptilinopus leclancheri*) and Rock dove quadrates. The postcapitular foramen is caudally located at the otic process on the Tooth-billed pigeon, Dodo, and Black-chinned fruit dove quadrates. Different with all other avian quadrates, a pneumatic foramen (dorsal foramen) is located at the centre of the intercapitular incisure on most Columbidae quadrates, such as Black-chinned fruit dove, Rock dove, and Grey-fronted dove.

**Gruiformes:** Heliornithidae (Plate 21)

In Heliornithidae (African finfoot, *Podica senegalensis*), the otic process has a deep and clear intercapitular incisure between squamosal and otic capitulum. The squamosal capitulum is as high as the otic capitulum in rostral view and it has an equal size with the otic capitulum. Both squamosal capitulum and otic capitulum have round shapes with convex articular surfaces, facing rostrolaterally and dorsomedially, respectively.

The orbital process of the African finfoot quadrate is robust and elongate with a slight high aspect ratio and it has a flat and slant tip, pointing mediorostrally. Besides, Heliornithidae quadrate exhibits a deep and distinct basiorbital fossa.

The quadratojugal cotyle in Heliornithidae has an oval shape with a thick, circular and complete margin (though the dorsal margin of the contact is inward) and a deep articulation, close to the lateral condyle. The contact on quadrate with pterygoid on Heliornithidae quadrate shows two articular surfaces: one (pterygoid condyle) is rostrally protruding, dorsally located at the medial condyle with a distinct and deep gap. It appears an oval shape with a convex articular surface. The other (orbitopterygoid facet) is medially located at the orbital process, close to the base of the orbital process. It shows a distinct oval shape with a flat articular surface for articulation.

The mandibular process on Heliornithidae quadrate shows three condyles, arranging in a L shape. The medial condyle shows a rectangular outline with a flat articular surface, and a lateral trochlea is present as a tiny projection with a flat articular surface. The lateral condyle is continuous with the caudal condyle and has and lateromedially elongate oval shape with a slightly convex articular surface. The caudal condyle is an elongate bulge caudally attached with the quadrate body.

The quadrate body in Heliornithidae is relatively curved with a curved lateral crest. A shallow but unclear caudomedial depression is caudally located at the quadrate body, between two capitula. In Heliornithidae, the pneumatic foramen is absent on its quadrate.

**Gruiformes:** Sarothruridae (Plate 21)

In Sarothruridae (Buff-spotted flufftail, *Sarothrura elegans*), the otic process has a shallow and wide intercapitular incisure between two capitula. The squamosal capitulum is higher than the otic capitulum, but it has a slightly smaller size than that of the otic capitulum. Both squamosal capitulum and otic capitulum on the Buff-spotted flufftail quadrate show rectangle-with rounded-corner shapes with flat articular surfaces, facing rostrolaterally and dorsomedially, respectively.

The orbital process of the Buff-spotted flufftail quadrate appears to be robust but short with a lower aspect ratio and has a flat tip, pointing rostrally. The basiorbital fossa of the Sarothruridae quadrate is deep and broad.

The quadratojugal cotyle of the Sarothruridae quadrate exhibits an oval shape with a thick and complete margin (though the dorsal margin of the contact is slightly inward). It has a deep articulation with the jugal bar and is adjacent to the lateral condyle. Unlike its close relatives (Heliornithidae), a thick and elongate bulge (submeatic process) is present and caudodorsally attached with the quadratojugal cotyle on the Buff-spotted flufftail quadrate. Like Heliornithidae, the articulation between quadrate and pterygoid on the Sarothruridae quadrate exhibits two contacts (pterygoid condyle and orbitopterygoid facet): the pterygoid condyle is rostrally protruding and above the medial condyle with a distinct and deep gap. It appears an oval shape with a convex articular surface. The orbitopterygoid facet is medially located at the orbital process, close to the base of the orbital process with a flat articular surface for attachment.

The mandibular process on Sarothruridae quadrate has three condyles, arranging in a L shape. The medial condyle has an oval shape with a convex articular surface, and the lateral trochlea is present, as shown in triangular outline with a flat articular surface. The medial condyle also has a distinct medial edge with its quadrate body. The lateral condyle is not distinct with the caudal condyle, and it has a lateromedially elongate oval shape with a slightly convex articular surface. The caudal condyle is a thick and rounded bulge caudomedially attached with the quadrate body.

The quadrate body in Sarothruridae is medially curved due to a curved medial crest. A shallow and unclear caudomedial depression is caudally located at the quadrate body, between two capitula. Like its sister group (Heliornithidae), the pneumatic foramen is absent on Sarothruridae quadrate.

**Gruiformes:** Rallidae (Plate 21 and 22)

In most Rallidae, the otic process has a deep and wide intercapitular incisure between two capitula but not found on Takahē (*Porphyrio hochstetteri*) and Inaccessible Island rail (*Atlantisia rogersi*) quadrates. The Takahē quadrate shows a deep but narrow intercapitular incisure while the Inaccessible Island rail quadrate exhibits a shallow and wide intercapitular incisure. The squamosal capitulum is only slightly higher than the otic capitulum, but the shape and its size of two capitula are disparity and variety among this clade. In Takahē, the squamosal capitulum is much larger than the otic capitulum. Its squamosal capitulum has a rounded shape with a convex articular surface, facing dorsally. Its otic capitulum has a rostrocaudally elongate oval with a flat articular surface, pointing dorsomedially. On the quadrates of the Tasmanian nativehen (*Tribonyx mortierii*) and Weka (*Gallirallus australis*), the squamosal capitulum shows equal size with the otic capitulum. The squamosal capitulum of the Tasmanian nativehen quadrate exhibits an oval shape with a convex articular surface, facing dorsolaterally. Its otic capitulum shows a rounded shape with a flat articular surface, facing caudodorsally. On Weka quadrate, the squamosal capitulum appears to be an oval shape (or rectangle shape with rounded angle) with a convex articular surface. Its otic capitulum appears a rectangle-with-rounded-angle shape with a flat articular surface, facing caudodorsally. On the quadrates of the Virginia rail (*Rallus limnicola*), Slaty-breasted rail (*Lewinia striata*), and Inaccessible Island rail, the otic capitulum is larger than the squamosal capitulum and all shows a rectangle-with-rounded-angle shape, facing caudodorsally. On the Virginia rail quadrate, the squamosal capitulum appears an elongate diamond shape and faces rostrodorsally. On the Slaty-breasted rail quadrate, the squamosal capitulum exhibits a rounded shape and faces dorsally. On the Inaccessible Island rail quadrate, the squamosal capitulum shows an elongate oval shape and points rostrodorsally.

The orbital process of the Rallidae quadrate is mostly robust and elongate with a high aspect ratio, pointing rostrodorsally. The Weka quadrate, however, shows a distinct crest on the orbital process extending from the ventral margin to the tip of the orbital process. In Rallidae, the tip of the orbital process shows various shapes: The Takahē quadrate shows a round and pointed tip, while most of other Rallidae quadrates appear to a flat tip. Surprisingly, the Takahē quadrate exhibits a standing orbital crest on the centre lateral side of the orbital process. The Rallidae quadrate mostly shows a deep and broad basiorbital fossa, but not found on the Tasmanian nativehen quadrate whose basiorbital fossa is shallow.

The quadratojugal cotyle of the Rallidae quadrate shows an oval or cup-like shape with a thick margin and a deep joint. It is mostly close to the lateral condyle with a small gap. However, the quadratojugal cotyle is adjacent to the lateral condyle on the quadrate of the Virginia rail, Slaty-breasted rail, and Tasmanian nativehen.

For the joint articulation with pterygoid, only Takahē and Tasmanian nativehen quadrates exhibit a single articulation with pterygoid (pterygoid condyle), differing from most Rallidae quadrates which have two articulations with pterygoid (pterygoid condyle and orbitopterygoid facet). The pterygoid condyle of Rallidae quadrates appears an oval shape with a convex articular surface, and is rostrally protruding, significantly separated from the medial condyle. The orbitopterygoid facet is medially located at the orbital process, close to the base of the orbital process, and it shows a flat articular surface for pterygoid attaching.

The mandibular process exhibits three condyles, arranging in a L- or triangular shape. Different with most Rallidae quadrate, the mandibular process of the Weka quadrate shows an inward curved margin between medial and lateral condyles. Only on the Takahē and Slaty-breasted rail quadrates, a clear fossa is present at the centre of the mandibular process. Like most of other Gruiformes, the medial condyle shows an oval outline with a saddle articular surface, separating into medial and lateral facet (lateral trochlea). The medial condyle also has a clear edge with quadrate body in all Gruiformes. The lateral condyle is continuous with the caudal condyle, and it has an elongate shape in a rectangle-with-rounded-corner with a convex articular surface. The caudal condyle is a thick and elongate bulge and caudomedially positioned along the quadrate body.

The quadrate body of Rallidae is thick and bears a right-angle lateral crest and medially curved medial crest. Similar as other Gruiformes, a shallow and unclear caudomedial depression is caudally located at the quadrate body. Unlike Heliornithidae and Sarothruridae, one single pneumatic foramen (rostromedial foramen) is medially located at quadrate body on most Rallidae quadrate – it is ventrally present at the otic process and above the basiorbital fossa, facing medially. This foramen is absent on *Atlantisia rogersi* quadrate, and appears to be a fossa on *Tribonyx mortierii* quadrate. Also, another clear fossa (basiorbital fossa) is medially located at the orbital process and close to the pterygoid condyle on *Tribonyx mortierii* quadrate.

**Gruiformes:** Psophidae (Plate 22)

On the quadrate of Psophidae (*Psophia crepitans*), it has a clear but shallow intercapitular incisure between the squamosal and otic capitulum. The squamosal capitulum is as high as the otic capitulum and it is slightly larger than otic capitulum. The squamosal capitulum has a lateromedially elongate oval shape with a convex articular surface, and it faces dorsally. The otic capitulum in Psophidae has a rostrocaudally elongate oval shape (though the medial margin of this facet is slightly inward) with a flat articular surface, and it faces caudodorsally.

The orbital process in *Psophia crepitans* is robust, elongate and straight with a high aspect ratio and a flat tip, pointing mediorostrally. Besides, Psophidae has a deep and wide basiorbital fossa.

The quadratojugal cotyle in Psophidae has a cap-like shape with a thick and circular margin. It has a deep fossa and is adjacent to the lateral condyle. The pterygoid condyle in *Psophia crepitans* is rostrally protruding and above the medial condyle with a distinct and deep gap. It has an oval or rounded shape with a slightly convex articular surface.

The mandibular process has three condyles, arranging in a L-shape. The medial condyle shows a triangular shape with a saddle articular surface, separating into medial and lateral facet (lateral trochlea). The lateral condyle is continuous with the caudal condyle, and it has an elongate shape in a rectangle-with-rounded-corner with a convex articular surface. The caudal condyle appears to be an elongate thick bulge and is positioned caudally along the quadrate body.

In Psophidae, the quadrate body is medially curved due to a curved medial crest. A shallow caudomedial depression caudally shows on the quadrate body, between two capitula. On the Psophidae quadrate, the pneumatic foramen (postcapitular foramen) is caudally located at the otic process (at the middle position between two capitula), and it shows a relatively large opening.

**Gruiformes:** Aramidae (Plate 22)

On the quadrate of Aramidae (*Aramus guarauna*), its otic process has a deep and narrow intercapitular incisure between the squamosal and otic capitulum. Unlike most avian quadrates, the squamosal capitulum is lower than the otic capitulum in rostral view and it is also smaller than otic capitulum. The squamosal capitulum has a rounded or squared shape with a slightly convex articular surface, facing rostrolaterally. The otic capitulum has a rostrocaudally elongate oval shape (though the medial margin is slightly inward) with a flat articular surface, facing caudodorsally.

The orbital process in *Aramus guarauna* is robust, elongate and straight with a high aspect ratio, and its tip is flat and slant, pointing rostrodorsally. Besides, Psophidae has a relatively deep and wide basiorbital fossa.

The quadratojugal cotyle in Aramidae has a cap-like shape with a thick and circular margin. It has a deep joint and is adjacent to the lateral condyle. A protrusion with a flat articular surface is rostroventrally located at the margin of the quadratojugal cotyle on Aramidae quadrate. The articulation on *Aramus guarauna* quadrate has two articular contacts with pterygoid: pterygoid condyle is rostrally standing and above the medial condyle with a distinct and deep gap. It has an oval shape with a convex articular surface. The orbitopterygoid facet, medially located at the orbital process, has an oval shape with a flat articular surface.

The mandibular process has three condyles, arranging in a L-shape. The medial condyle shows an oval shape with a saddle articular surface, separating into medial and lateral facet (lateral trochlea). The lateral condyle is continuous with the caudal condyle, and it has an elongate rectangle-with-rounded-corner shape with a convex articular surface. The caudal condyle is a thick and oval bulge caudally attached with the quadrate body.

In Aramidae, the quadrate body is wide with a relatively straight medial crest and right-angle lateral crest. In the caudal view, there is a shallow caudomedial depression between two capitula. Different with other Gruiformes, a subcapitular tubercle is rostroventrally located at the squamosal capitulum, and it has a bulge-like protrusion. In Aramidae, the pneumatic foramen (postcapitular foramen) is caudally present at the otic process (at the middle position of two capitula), but its opening appears to be relatively smaller than one found on *Psophia crepitans* quadrate.

**Gruiformes:** Gruidae (Plate 22)

On the Gruidae quadrate, the intercapitular incisure between squamosal and otic capitulum is shallow and narrow on the otic process. The squamosal capitulum is higher and larger than the otic capitulum. The squamosal capitulum shows a rectangular shape with a convex articular surface, facing rostrodorsally. The otic capitulum shows a rostrocaudally elongate oval shape with a flat articular surface, facing dorsomedially.

Like other Gruiformes, the orbital process is robust, elongate and straight with a high aspect ratio, but its tip is distinct with any other Gruidae – its tip is flat and slant with a rostrolaterally elongate protrusion, pointing rostrally. The basiorbital fossa of the Gruidae quadrate is shallow and wide.

The quadratojugal cotyle of the Gruidae quadrate exhibits an oval shape with a thick and circular margin. It has a deep articular surface and is adjacent to the lateral condyle. The pterygoid condyle on the Gruidae quadrate is rostrally standing and dorsally separated from the medial condyle with a distinct and deep gap. It has a lateromedially elongate oval shape with a convex articular surface. On the quadrate of *Leucogeranus leucogeranus*, the second articulation with the pterygoid (orbitopterygoid facet) is present and is medially located at the orbital process, close to the pterygoid condyle with an oval outline and a flat articular surface.

The mandibular process on Gruidae quadrate has three condyles, arranging in an oblique T shape. The medial condyle shows an oval shape with a saddle articular surface, separating into medial and lateral facet (lateral trochlea). The lateral condyle is continuous with the caudal condyle, and it has an elongate rectangle-with-rounded-corner shape with a convex articular surface. The caudal condyle is a thick, slant bulge and positioned caudomedially attached along the quadrate body.

In Gruidae, the quadrate body is wide and slightly medially curved with a relative curved medial crest. A shallow caudomedial depression is caudally located at the otic process, between two capitula. Similar as the Aramidae quadrates but slightly different with this group, two types of the subcapitular tubercle are ventrolaterally located at the squamosal capitulum: one appears to a tongue-like shape with a flat surface, rostroventrally located at the squamosal capitulum; the other shows a tiny protrusion, caudoventrally located at the squamosal capitulum. On the Gruidae quadrate, the postcapitular foramen shows on all the specimens in this study, and it is caudally located at the otic process with a relatively larger opening. The caudomedial foramen, accommodated at the caudomedial depression, are only found on the *Balearica pavonine* quadrate and faces medially. The basiorbital foramen medially shows on the quadrate body of *Leucogeranus leucogeranus* only with a small opening.

**Phoenicopteriformes:** Phoenicopteridae (Plate 23)

In Phoenicopteridae, the otic process has a shallow and wide intercapitular incisure between the squamosal and otic capitulum. The squamosal capitulum is significantly higher than the otic capitulum but it has an equal size with the otic capitulum. The squamosal capitulum has a rounded shape and the otic capitulum has a lateromedial elongate oval shape with a convex articular surface in *Phoenicopterus roseus*, while it has a rounded shape with a slightly convex articular surface in *Phoenicoparrus jamesi*. It faces dorsally (*Phoenicopterus roseus*) or dorsolaterally (*Phoenicoparrus jamesi*) in otic capitulum of the otic process.

The orbital process in Phoenicopteridae is robust and elongate with a high aspect ratio and its ventral margin is significantly broad (Saiff, 1978). Its tip is rounded (*Phoenicopterus roseus*) or flat (*Phoenicoparrus*), pointing rostrodorsally. On the surface of the orbital process in Phoenicopteridae, a standing orbital crest is present with a ridge shape, extending from its tip to the base of the orbital process (Saiff, 1978), especially in *Phoenicoparrus*. Besides, Phoenicopteridae has a broad basiorbital fossa, but it is much deeper in *Phoenicoparrus* than that in *Phoenicopterus*.

The quadratojugal cotyle has a cup-like shape with a thick and circular and margin. It has a deep articular joint and is close to the lateral condyle (Saiff, 1978). In Phoenicopteridae, the pterygoid condyle is rostromedially protruding in *Phoenicopterus*, but it is blunt in *Phoenicoparrus*. The pterygoid condyle in Phoenicopteridae is dorsally located at the medial condyle with a broad gap, and it has a slender oval shape (Mayr, 2015) with a slightly convex articular surface.

The mandibular process has three condyles, arranging in T shape, and it has a deep fossa or a distinct intercondylar sulcus at the centre position (Saiff, 1978). The medial condyle appears to be an oval shape with a saddle articular surface, separating into medial and lateral facet (lateral trochlea) (Saiff, 1978). The medial facet of the medial condyle is prominent with a clear convex articular surface, while the lateral facet shows a flat articular surface, instead. The medial condyle appears a clear margin with the quadrate body. The lateral condyle has an oval shape with a slightly flat surface, while the caudal condyle is a rounded bulge caudomedially attached with the quadrate body with a smooth articular surface.

In Phoenicopteridae, the quadrate body is relatively straight and wide with a straight lateral and medial crest. The caudomedial depression is shallow and unclear on the Phoenicopteridae quadrate, especially that of *Phoenicoparrus*. On flamingo quadrate, a subcapitular tubercle is ventrolaterally located at the squamosal capitulum (Mayr, 2015), and it is a rounded and tiny protrusion with a flat articular surface. In Phoenicopteridae, the pneumatic foramen (caudomedial foramen) is caudomedially located at the caudomedial depression (Saiff, 1978), facing caudomedially.

**Podicipediformes:** Podicipedidae (Plate 23)

On grebe (Podicipedidae) quadrate, the oitc process usually has a shallow and wide intercapitular incisure between the squamosal and otic capitulum, but the intercapitular incisure is significantly deep in *Podilymbus gigas*. The squamosal capitulum is as high as the otic capitulum, but it only has the similar size with the otic capitulum in *Podiceps taczanowskii*. The squamosal capitulum is much larger in most grebes, such as *Rollandia Rolland* and *Podilymbus gigas*.

The otic process in Podicipedidae shows a huge morphological diversity: in *Rollandia Rolland*, the squamosal capitulum has two facets contacting with cranium, the rostral and caudal facet. The rostral facet has a slender diamond shape with a slightly convex articular surface, while the caudal facet has a dorsoventrally elongate ovois shape. A clear incisure is present between these two facets. Its otic capitulum has a rectangle-with-rounded-corner shape with a flat articular surface, pointing dorsomedially. In *Podiceps taczanowskii*, the squamosal capitulum has a triangle shape (though lateral margin is curved) with a convex articular surface, facing dorsally. Its otic capitulum has a lateromedially elongate oval shape with a slightly convex articular surface, facing in dorsally. In *Podilymbus gigas*, the squamosal capitulum has a kidney-like shape (a curved lateral margin) with a convex articular surface, facing dorsally. Its otic capitulum has a rounded shape, pointing caudodorsally.

Unlike its sister group (Phoenicopteridae), the orbital process in Podicipedidae is slender and elongate with a high aspect ratio. Its tip is flat and turns into horizontal (in *Podiceps taczanowskii*) or subhorizontal (*Rollandia rolland* and *Podilymbus gigas*). The orbital crest is only present in *Podilymbus gigas* with a L shape close to the tip of the orbital process. Besides, the basiorbital fossa in Phoenicopteridae is deep and wide, but it is shallow in *Podilymbus gigas*.

The quadratojugal cotyle in Podicipedidae has a cup-like shape with a thick and circular margin. It is a deep articulation and close to the lateral condyle. The quadratojugal cotyle is relatively laterally standing in grebe than that of flamingo (Mayr, 2015). The pterygoid condyle in Podicipedidae is rostrally protruding (Mayr, 2015), and it is dorsomedially located at the medial condyle with a small gap. It has either an oval shape or rounded shape (only *Podiceps taczanowskii*) with a slightly convex articular surface.

The mandibular process on Podicipedidae quadrate has three condyles, arranging in a L shape, and a clear fossa is present between the medial and lateral condyle in all grebes, especially in *Podilymbus gigas*. Similar as Phoenicopteridae, the medial condyle shows an oval outline with a saddle articular surface, separating into medial and lateral facet (lateral trochlea). The medial facet is prominent with a convex articular surface and a rostral tip (Mayr, 2015), while the lateral facet appears a flat articular surface instead. Both these two facets appear a clear margin with the quadrate body. The lateral condyle shows an oval or rounded shape with a slightly flat articular surface. The caudal condyle is an elongate and slender protrusion and caudally positioned along the quadrate body in most grebes. However, in *Podiceps taczanowskii*, its caudal condyle has a rounded shape and relatively thicker.

In Podicipedidae, the quadrate body is relatively straight and slender with a straight lateral and medial crest. The caudomedial depression is shallow but distinct on the grebe quadrates. The subcapitular tubercle is ventrolaterally located at the squamosal capitulum (Mayr, 2015) on some grebes, such as *Podiceps taczanowskii*, with a slender and standing ridge. In Podicipedidae, the pneumatic foramen is absent, different with its sister group.

**Charadriiformes:** Burhinidae (Plate 24)

In Burhinidae (*Burhinus senegalensis*), its oitc process has a shallow but clear intercapitular incisure between the squamosal and otic capitulum. The squamosal capitulum is higher than the otic capitulum and it has larger size with the otic capitulum. The squamosal capitulum has a rounded or squared shape with a rostrocaudally bending and smooth articular surface, pointing dorsolaterally. The otic capitulum has a lateromedially elongate oval shape, facing dorsomedially.

The orbital process in Burhinidae is robust but slightly short with a low aspect ratio, and its tip is broad and flat, pointing rostrodorsally. Besides, Burhinidae has a shallow but broad basiorbital fossa.

The quadratojugal cotyle of Burhinidae quadrate is slightly laterally standing and has a cup-like shape with a thick and circular margin (though its dorsal margin is slightly inward). It has a deep articulation with a jugal and is close to the lateral condyle. In *Burhinus senegalensis*, its pterygoid condyle is rostrally standing and is dorsally located at the medial condyle with a wide and deep gap. It has a slender oval shape with a convex articular surface.

The mandibular process on Burhinidae quadrate has three condyles, arranging in a L or triangle shape with a clear fossa at the centre position. The medial condyle has an oval shape with a saddle articular surface, separating into medial and lateral facet (lateral trochlea). The medial facet is prominent with a clear convex articular surface, while the lateral facet shows a flat articular surface. Both these two facets appear a clear margin with the quadrate body. The lateral condyle shows a rounded shape and bears a convex articular surface. The caudal condyle appears to be an elongate and thick protrusion and is caudally positioned along the quadrate body with a smooth articular surface.

In Burhinidae, the quadrate body is relatively curved and slender with a curved lateral and medial crest. A shallow caudomedial depression and a deep caudomedial fossa is caudally present at the quadrate body. However, due to a significantly twisted tympanic crest, the caudomedial fossa faces more caudomedially instead of caudally. The subcapitular tubercle is laterally located at the squamosal capitulum with a round shape and a convex surface. In Burhinidae, the pneumatic foramen (postcapitular foramen) is caudally located at the otic process, with a larger opening. Also, the caudomedial fossa abovementioned might be also related to the avian air sac system.

**Charadriiformes:** Charadriidae (Plate 24)

The quadrate of Charadriidae (*Charadrius vociferus*) has a narrow but deep intercapitular incisure on its otic process. The squamosal capitulum is slightly higher than the otic capitulum and it also has a larger size with the otic capitulum. The squamosal capitulum has two articulations with cranium, the lateral and medial facet. The lateral facet has a rounded shape with a rostrocaudally bending and convex articular surface, while the medial facet has an oval shape with a flat articular surface. There is not a clear incisure between these two facets. The otic capitulum has a rounded shape, facing in dorsally. Below the otic capitulum, there is a clear and deep notch.

The orbital process in Charadriidae is robust and slightly elongate with a slightly higher aspect ratio, and its tip is broad and flat, pointing rostrodorsally. Besides, Charadriidae has a relatively deep and broad basiorbital fossa.

The quadratojugal cotyle of Charadriidae quadrate is slightly laterally standing and has a cup-like shape with a thick and circular margin (though its dorsal and ventral margins are slightly inward). It has a deep joint contacting with the jugal bone and is close to the lateral condyle. In *Charadrius vociferus*, the articular surface on quadrate has two connections with the pterygoid: the pterygoid condyle is rostrally standing and above the medial condyle with a small gap. It has an oval shape with a convex articular surface; the orbitopterygoid facet is ventrally located at the orbital process (close to the base of the orbital process), and it has an oval shape with a smooth articular surface.

The mandibular process has three condyles, arranging in a right-triangular shape with a shallow fossa at the centre position. The medial condyle has an oval shape with a saddle articular surface, separating into medial and lateral facet (lateral trochlea). The medial facet has a clear convex articular surface, while the lateral facet appears a flat articular surface. Both these two facets show a clear margin with the quadrate body. The lateral condyle has an oval shape with a slightly convex articular surface. The caudal condyle appears to be an elongate thick bulge and is caudomedially positioned along the quadrate body with a smooth articular surface.

In Charadriidae, the quadrate body of is slightly curved with a right-angled lateral crest and curved medial crest. The caudomedial depression is unclear or not well-developed, but the caudomedial fossa is clear and deep, facing caudomedially. The subcapitular tubercle is caudolaterally located at the squamosal capitulum with a rounded and flat surface. In Charadriidae, the pneumatic foramen (basiorbital foramen) is medially located at the quadrate body, close to the base of the orbital process. Similar as Burhinidae, the caudomedial fossa abovementioned might be capable of air exchange in the avian air sac system.

**Charadriiformes:** Recurvirostridae (Plate 24)

In Recurvirostridae (*Recurvirostra avosetta*), the intercapitular incisure is absent on the otic process, indicating that two capitula are continuous or close to each other. The squamosal capitulum is as high as the otic capitulum and it is slightly smaller than the otic capitulum. The squamosal and otic capitulum all have lateromedially elongate oval shapes with convex articular surfaces, pointing rostrodorsally and caudodorsally, respectively. Below the otic capitulum, there is a clear and deep notch.

The orbital process in Recurvirostridae is robust and extremely elongate with a significantly higher aspect ratio, and its tip is thick, broad and rounded, pointing rostrally. Besides, Recurvirostridae has a relatively deep and broad basiorbital fossa.

The quadratojugal cotyle in Recurvirostridae is slightly laterally standing and it has a cup-like shape with a thick and circular margin (though its dorsal and ventral margins are slightly inward). It has a deep articulation and is close to the lateral condyle. Its caudal (or caudodorsal) margin is prominent and laterally standing. In *Recurvirostra avosetta*, the articulations with pterygoid on quadrate has two connections: one (pterygoid condyle) is rostrally standing and abobve the medial condyle with a small gap. It has an oval shape with a flat articular surface. The other (orbitopterygoid) is ventrally located at the orbital process and has a flat articular surface.

The mandibular process exhibits three condyles, arranging in a triangular outline and bears a shallow fossa at the centre position. The medial condyle has an oval shape with a saddle articular surface, separating into medial and lateral facet (lateral trochlea). Both facets show flat articular surfaces and clear margins with the quadrate body. The lateral condyle has a rounded shape with a slightly convex articular surface. The caudal condyle appears to be an elongate and thick bulge and is caudomedially positioned along the quadrate body with a smooth articular surface.

In Recurvirostridae, the quadrate body is slightly straight with a right-angled lateral crest and straight medial crest. In *Recurvirostra avosetta*, the caudomedial depression is either shallow or unclear between two capitula. The caudomedial fossa is relatively deep and faces more caudomedially. The subcapitular tubercle is caudolaterally located at the squamosal capitulum with a dorsoventrally elongate oval shape and a convex surface. The pneumatic foramen (rostromedial foramen) on the Recurvirostridae quadrate is medially located at the quadrate body and above the basiorbital fossa. It is relatively large and surrounded by the medial crest and orbital process. Besides, the caudomedial fossa abovementioned might be also related the air exchange.

**Charadriiformes:** Haematopodidae (Plate 24)

In Haematopodidae (*Haematopus ostralegus*), the intercapitular incisure is absent on the otic process. The squamosal capitulum is slightly higher than the otic capitulum but it has an equal size with the otic capitulum.

Similar as Recurvirostridae, both squamosal and otic capitulum all show lateromedially elongate oval outline with convex articular surfaces, facing dorsolaterally and dorsomedially, respectively. Below the otic capitulum, there is a clear and deep notch.

The orbital process in Haematopodidae is robust and elongate with a significantly higher aspect ratio, and its tip is thick, broad and blunt, facing rostrally. Besides, Haematopodidae has a deep and narrow basiorbital fossa.

The quadratojugal cotyle in Haematopodidae is slightly laterally standing and it has a cup-like shape with an oval margin (though its rostrodorsal margin is slightly inward). It has a deep articulation and is adjacent to the lateral condyle. Its caudal (or caudodorsal) margin is prominent and dorsally standing. In *Haematopus ostralegus*, the pterygoid condyle is not rostrally protruding and is separated from the medial condyle with a deep gap. It has a lateromedially elongate oval shape (or rectangle-with-rounded-corner shape) with a flat articular surface.

The mandibular process has three condyles, arranging in a Z-shape. The medial condyle shows a lateromedially elongate oval shape with a saddle articular surface, separating into medial and lateral facet (lateral trochlea). Both facets appears to be prominent with a flat articular surface and bear clear margin with the quadrate body. The lateral condyle shows a rounded shape with a smooth articular surface. The caudal condyle appears to be a thick protrusion and is caudomedially positioned along the quadrate body with a concave articular surface.

In Haematopodidae, the quadrate body is slightly straight with a right-angled lateral crest and straight medial crest. In *Haematopus ostralegus*, the caudomedial depression is shallow and caudally located at the otic process. The subcapitular tubercle is caudolaterally located at the squamosal capitulum with a dorsoventrally elongate oval shape and a convex surface. On the Haematopodidae quadrate, the pneumatic foramen (postcapitular foramen) is caudally located at the otic process with a relatively large opening.

**Charadriiformes:** Pedionomidae (Plate 24)

In Pedionomidae (*Pedionomus torquatus*), the intercapitular incisure is shallow and narrow between two capitula. The squamosal capitulum is higher than the otic capitulum, and it has an equal or a slightly smaller size than the otic capitulum. The squamosal capitulum has a rounded shape with a convex articular surface, pointing dorsolaterally. The otic capitulum has a rostrocaudally elongate oval shape with a flat articular surface, and it faces caudodorsally.

The orbital process in *Pedionomus torquatus* is robust but slightly short with a relatively lower aspect ratio. Its tip is broad and blunt, pointing rostrodorsally. Besides, Pedionomidae has a shallow and broad basiorbital fossa.

The quadratojugal cotyle in Pedionomidae is slightly laterally standing and has a cup-like shape with a thick and circular margin (though its dorsal and ventral margin is slightly inward). It has a deep joint and is adjacent to the lateral condyle. The pteyrogid condyle in *Pedionomus torquatus* is rostrally standing and above the medial condyle with a wide and deep gap. It has an oval shape with a convex articular surface.

The mandibular process has three condyles, arranging in a triangular shape with a shallow fossa at the centre position. The medial condyle has a rounded shape with a slightly convex articular surface. The lateral condyle appears a rounded shape with a slightly convex articular surface, while the caudal condyle shows a slant, elongate and thick bulge caudally attached with the quadrate body with a smooth articular surface.

In Pedionomidae, the quadrate body is slightly curved due to its right-angled lateral crest and curved medial crest. The caudomedial depression is shallow, but the caudomedial fossa is deeply accommodate at the caudomedial depression, facing caudally. In Pedionomidaem, two pneumatic foramina are located at the quadrate body: one (rostromedial foramen) is medially located at the otic process, surrounded by the medial crest and orbital process; the other (postsquamosal capitulum foramen) is caudally located at the squamosal capitulum. The abovementioned caudomedial fossa likely links with the cranial avian air sac system.

**Charadriiformes:** Jacanidae (Plate 24)

In Jacanidae, the otic process has an unclear and shallow intercapitular incisure between the squamosal and otic capitulum. The squamosal capitulum has the same height and size as the otic capitulum. The squamosal capitulum has a rounded shape with a convex articular surface, pointing rostrodorsally. The otic capitulum has a rostrocaudally elongate oval or rectangle-with-rounded-corner shape with a flat articular surface, pointing dorsomedially.

The orbital process in Jacanidae is robust and elongate with a slightly higher aspect ratio. Its tip is rounded and points rostrally. Besides, Jacanidae quadrate has a shallow and broad basiorbital fossa.

The quadratojugal cotyle in Jacanidae is slightly laterally standing and it has a cup-like shape with a thick and oval margin (though its dorsal margin is slightly inward). It has a deep fossa to connect with the jugal bar and is adjacent to the lateral condyle. On Jacanidae quadrate, the articular surface with pterygoid has two connections: one (pterygoid condyle) is standing and separated from the medial condyle with a wide and deep gap. It has an oval shape with a convex articular surface; the other (orbtiopterygoid facet) has an oval shape with a flat articular surface, medially located at the orbital process and close to the base of the orbital process.

The mandibular process exhibits three condyles, arranging in a L shape with a shallow fossa at the centre position. The medial condyle shows a rostrocaudally elongate oval outline with a slightly convex articular surface. It has a clear margin with the quadrate body. The lateral condyle shows an oval shape with a slightly convex articular surface. The caudal condyle appears to be a thick and rounded bulge and is caudomedially positioned along the quadrate body with a convex articular surface.

In Jacanidae, the quadrate body is slightly curved due to its right-angled lateral crest and curved medial crest. The caudomedial depression, caudally located at the quadrate body, is narrow and deep. In Jacanidae, only one pneumatic foramen (rostromedial foramen) is medially located at the quadrate body, as shown in a relatively large opening. Iit is surrounded by the medial crest and orbital process.

**Charadriiformes:** Scolopacidae (Plate 25)

The quadrate of Scolopacidae shows a great morphological diversity on each character, such as muscle attachment, shape of articulation contact, or pneumaticity. For instance, most of Scolopacidae quadrate has a narrow and shallow intercapitular incisure on the otic process, but not in *Scolopax rusticola* whose intercapitular incisure is absent. The squamosal capitulum is higher than the otic capitulum in *Limosa lapponica* and *Numenius phaeopus*, while it has same height as the otic capitulum in *Lymnocryptes minimus* and *Scolopax rusticola*. The squamosal capitulum is much smaller than the otic capitulum in most taxon but not in *Lymnocryptes minimus* whose squamosal capitulum has an equal size as the otic capitulum.

The squamosal capitulum in Scolopacidae usually has a lateromedial elongate oval shape with a slightly convex articular surface, facing dorsolaterally, though it has a smooth articular surface in *Lymnocryptes minimus* and a convex surface in *Numenius phaeopus*. However, in *Limosa lapponica*, its squamosal capitulum has a round shape with a convex articular surface, facing dorsally. The otic capitulum has a rounded shape, facing dorsally or caudodorsally in *Limosa lapponica* and *Scolopax rusticola*, while it has a lateromedially elongate oval shape, facing either dorsomedially in *Lymnocryptes minimus* or caudodorsally in *Numenius phaeopus*. Besides, a clear notch is ventromedially located at the otic capitulum in all Scolopacidae.

In Scolopacidae, the orbital process is robust and elongate and usually faces rostrally, but it faces dorsolaterally in *Scolopax rusticola* only. The tip of the orbital process in sandpiper quadrates has different shape taxa by taxa. For instance, it is rounded in *Lymnocryptes minimus* or pointed in *Scolopax rusticola*. In *Limosa lapponica* and *Numenius phaeopus*, it is flat. Usually, the basiorbital fossa in Scolopacidae is shallow and broad, but it is relatively deeper than other taxon in *Limosa lapponica*.

The quadratojugal cotyle in Scolopacidae is slightly laterally standing and has a cup-like shape with a thick and oval margin (though its dorsal and ventral margin is slightly inward). It has a deep fossa and is usually separated from the lateral condyle with a tiny gap, except *Lymnocryptes minimus* whose quadratojugal cotyle is adjacent to the lateral condyle. Different with most avian quadrates, the quadratojugal cotyle in *Scolopax rusticola* faces ventrolaterally, instead of laterally. The articular contact with pterygoid on Scolopacidae quadrate also has two articulations: one (pterygoid condyle) is slightly standing and it is dorsally located at the medial condyle with a clear and deep gap. It has either a lateromedially elongate oval shape in most Scolopacidae or a rounded shape in *Scolopax rusticola*. The other (orbitopterygoid facet) is a smooth articular surface medially located at the orbital process close to the base of the orbital process.

The mandibular process in Scolopacidae exhibits three condyles, arranging in a triangular shape, and there is a shallow fossa at the centre position in most Scolopacidae. The medial condyle usually shows a lateromedially elongate oval outline with a prominent convex articular surface (especially in *Numenius*), separating into lateral and medial facet (lateral torchlea). In *Lymnocryptes*, the medial condyle shows a rounded outline. The medial conylde shows a clear margin with the quadrate body in all sandpiper quadrates. The lateral condyle appear to be either a rounded shape (*Limosa*) or lateromedial elongate oval outline (e.g., *Lymnocryptes*, *Scolopax*, and *Numenius*) with a slightly convex articular surface in most sandpiper. In *Lymnocryptes*, the lateral condyle shows a flat articular surface. The caudal condyle appears to be a thick and rounded bulge and is caudomedially positioned along the quadrate body with a convex articular surface in most sandpiper. In *Lymnocryptes*, the caudal condyle appears a flat articular surface.

In Scolopacidae, the quadrate body is straight due to its right-angled lateral crest and slightly straight medial crest. The caudomedial depression is shallow and narrow on most sandpiper quadrate, but not on the Eurasian woodcock (*Scolopax*) quadrate. The caudomedial fossa is accommodated at the caudomedial depression on the Jack snipe (*Lymnocryptes*) quadrate only. The subcapitular tubercle only appears on the Jack snipe and Bar-tailed godwit (*Limosa*) quadrates, and it is laterally or rostrally located at the squamosal capitulum with a tiny and rounded protrusion. On the Scolopacidae quadrate, the avian pneumatic foramen is located at four positions: the postsquamosal capitulum foramen is caudally located at the squamosal capitulum on Eurasian woodcock quadrate only. The rostromedial foramen is medially located at the quadrate body and is above the basiorbital fossa on most Scolopacidae quadrates. The caudomedial foramen is accommodated at the caudomedial depression on the Eurasian whimbrel (*Numenius*) and Jack snipe (as a fossa abovementioned) quadrates. The postcapitular foramen is caudally located at the otic process of the Bar-tailed godwit and Eurasian woodcock quadrates.

**Charadriiformes:** Turnicidae (Plate 25)

In buttonquail (Turnicidae), it has a broad and shallow intercapitular incisure on its otic process. The squamosal capitulum is significantly higher than the otic capitulum, but it is slightly smaller than the otic capitulum. The squamosal capitulum has a rounded shape with a convex articular surface, pointing dorsally. The otic capitulum has a lateromedially elongate oval shape with a flat articular surface, facing dorsomedially.

The orbital process in buttonquail is slender and elongate with a slightly higher aspect ratio. Similar as that of *Scolopax rusticola* quadrate, the rostral portion gently dorsally twists, facing dorsally, and its tip is rounded. Besides, Turnicidae has a shallow and narrow basiorbital fossa.

The quadratojugal cotyle in Turnicidae is laterally standing and has a cup-like shape with a thick and oval margin. It has a slightly deep fossa and is adjacent to the lateral condyle. The articular surface with pterygoid on buttonquail quadrate has two contacts: one (pterygoid condyle) is rostrally standing and is rostrodorsally located at the medial condyle with a wide gap. It has a rounded shape with a slightly convex articular surface; the other (orbitopterygoid facet) is medially located at the orbital process and has a distinct oval shape with a flat articular surface.

The mandibular process has three condyles, arranging in a L-shape with a shallow fossa at the centre position. The medial condyle shows a lateromedially elongate oval shape with a convex articular surface and bears a clear margin with the quadrate body. The lateral condyle shows a lateromedially elongate oval outline with a convex articular surface. The caudal condyle appears to be a thick bulge and is caudomedially positioned along the quadrate body with a convex articular surface.

In Turnicidae, the quadrate body is slender and curved with its right-angled lateral crest and a curved medial crest. The caudomedial depression is wide and deep, and the caudomedial fossa is also deep as shown on the quadrate body. Even though the pneumatic foramen on the Turnicidae is absent, two clear fossae likely work similar function as pneumatic foramen in air sac system: the caudomedial fossa is caudomedially located at the quadrate body, and the rostromedial fossa is ventromedially located at the otic process and surrounded by the medial crest and a distinct ridge.

**Charadriiformes:** Stercorariidae (Plate 25)

On the otic process of skua quadrate (Stercorariidae), the intercapitular incisure is shallow and narrow. The squamosal capitulum is slightly higher than the otic capitulum, but it has an equal size as its otic capitulum. Both squamosal and otic capitulum has a lateromedially elongate oval shape with a convex articular surface, facing dorsolaterally and dorsomedially, respectively.

The orbital process in skua is robust and relatively short with a slightly lower aspect ratio than most Charadriiformes. The tip of the orbital process is flat and broad, facing rostrodorsally, and its rostral portion is slightly twisted. Besides, the basiorbital fossa is slightly deep and broad in Stercorariidae.

The quadratojugal cotyle is laterally standing and has a cup-like shape with a thick oval margin. It has a markedly deep fossa and is dorsally located adjacent to the lateral condyle. The pterygoid condyle is rostrally standing and is close to the medial condyle with a small gap. It has a rounded shape with a convex articular surface.

The mandibular process exhibits three condyles, arranging in a L-shape with a shallow fossa at the centre position. The medial condyle shows a rostrocaudally rectangular outline with a prominent articular surface. Also, it shows a distinct margin with the quadrate body. The lateral condyle shows a lateromedially elongate oval shape with a convex articular surface. The caudal condyle appears to be a thick bulge and is caudally positioned along the quadrate body with a convex articular surface.

In Stercorariidae, the quadrate body is wide and slightly straight with its right-angled lateral crest and a minorly curved medial crest. The caudomedial depression is significantly unclear and shallow on the skua quadrate. Two pneumatic foramina are present on the Stercorariidae quadrate: the basiorbital foramen is medially located at the quadrate body, close to the pterygoid condyle and base of the orbital process; the postotic capitulum foramen is caudoventrally located at the otic capitulum.

**Charadriiformes:** Glareolidae (Plate 26)

The quadrate in Glareolidae does not have a clear or distinct intercapitular incisure between squamosal and otic capitulum, but each capitulum is far from each other. The squamosal capitulum is significantly higher than the otic capitulum and has a larger size than the otic capitulum. The squamosal capitulum has a rounded shape with a convex articular surface, facing dorsolaterally, and it is rostrocaudally bending. The otic capitulum has a rostrocaudally elongate oval shape with a slightly convex articular surface, pointing caudomedially.

The orbital process in Glareolidae is thin and robust, but it is relatively short with a low aspect ratio. Its tip is flat with a sharp edge, facing rostrally. Besides, Glareolidae quadrate has a shallow and narrow basiorbital fossa.

The quadratojugal cotyle is laterally standing and shows a cup-like shape with a thick and oval margin. It also has a deep articular articular surface and is adjacent to the lateral condyle. The ventral margin of the quadratojugal cotyle expands ventrally, forming a flat articular surface. The articulation with pterygoid on Glareolidae quadrate has two contacts: one (pterygoid condyle) is slightly standing and is above the medial condyle with a wide gap. It has a rounded shape with a slightly convex articular surface. The other (orbitopterygoid facet) isa distinct bulge with a flat articular surface, and it is medially located at the orbital process and is close to the base of the orbital process.

The mandibular process exhibits three condyles, arranging in a boomerang-like shape. A deep fossa or a clear intercondylar sulcus is located between the medial and lateral condyle. Both medial and lateral condyles show an oval outline with prominent articular surfaces, but they are elongate in rostrocaudal and lateromedial, respectively. The caudal condyle appears to be not well-developed, as shown in a tiny protrusion, and is caudally positioned along the quadrate body with a flat articular surface.

In Glareolidae, the quadrate body is only slightly medially curved due to its right-angled lateral crest and a curved medial crest. The caudomedial depression is shallow and wide, but a deep fossa is accommodated at the caudomedial depression. The Glareolidae quadrate is highly pneumatic: the postcapitular foramen is caudally locate at the otic process. The rostromedial and basiorbital foramina are medially located at the quadrate body. The caudomedial foramen is accommodated at the caudomedial depression.

**Charadriiformes:** Alcidae (Plate 26)

In Alcidae (*Uria aalge*), the otic process has a slightly narrow intercapitular incisure between two capitula. The squamosal capitulum is slightly higher and has a larger size than the otic capitulum. The squamosal capitulum has a lateromedially elongate oval shape with a convex articular surface, facing dorsolaterally. The otic capitulum has a rostrocaudally elongate oval shape and bears a convex articular surface, pointing dorsomedially.

The orbital process in Alcidae is robust and slender with a high aspect ratio, and its tip is flat, pointing rostrally. Besides, Alcidae quadrate has a wide and deep basiorbital fossa.

The quadratojugal cotyle is laterally standing and has a cup-like shape with a thick, oval margin. It has a deep joint connecting with jugal bone and is adjacent to the lateral condyle. The ventral margin of the quadratojugal cotyle expands ventrally, forming a flat articular surface. The pterygoid condyle in Alcidae is rostrally standing and has a rounded shape with a convex articular surface. It is dorsally located at the medial condyle with a wide and deep gap.

The mandibular process exhibits three condyles, arranging in a L-shape. The medial condyle shows a rounded shape with a convex articular surface, and it shows a clear margin with the quadrate body. The lateral condyle shows a lateromedially elongate oval outline with a convex articular surface. It shows a clear rostral margin with the quadrate body. The caudal condyle appears to be an elongate and slant protrusion, and is caudomedially positioned along the quadrate body with a convex articular surface.

In Alcidae, the quadrate body is slightly straight due to its right-angled lateral crest and a minorly curved medial crest. In the caudal view, neither caudomedial depression nor caudomedial fossa is absent in Alcidae. Also, the pneumatic foramen is absent on the Alcidae quadrate.

**Charadriiformes:** Laridae (Plate 26)

The otic process of Laridae quadrates exhibit a wide and deep intercapitular incisure between two capitula. The squamosal capitulum is significantly higher than the otic capitulum, especially in *Rynchops niger*. In *Chroicocephalus novaehollandiae*, the squamosal capitulum shows an equal size to the otic capitulum, whereas in *Rynchops* it is slightly larger than the otic capitulum. The squamosal capitulum shows a rounded shape and it is oriented rostrodorsally. The otic capitulum shows a lateromedially elongate oval outline and points dorsally. Two capitula show convex articular surfaces in *Chroicocephalus*, but they show flat articular surfaces in *Rynchops*.

The orbital portion (orbital process and basiorbital fossa) is completely different between two studied Laridae quadrates. In *Chroicocephalus*, the orbital process is slightly elongate and thick with a higher aspect ratio. Its tip is flat and slant, facing rostrodorsally. On the other hand, the orbital process in *Rynchops* is thin and relatively short with a lower aspect ratio. Its tip is sharp, pointing rostrodorsally. Besides, the basiorbital fossa in *Chroicocephalus novaehollandiae* is narrow and deep, while it is narrow and shallow in *Rynchops*.

Like other Charadriiformes, the quadratojugal cotyle of Laridae quadrate is laterally standing, but it is more ventrally standing in *Rynchops*, making its quadratojugal cotyle lower than the medial condyle. It has a cup-like shape with a thick and oval margin (though the ventral margin is slightly inward in *Rynchops*). It has a deep articular contact with the jugal bone and is adjacent to the lateral condyle. Similar as most Charadriiformes, the articulations with pterygoid on quadrate in Laridae has two contacts: one (pterygoid condyle) is rostrally standing and is separated from the medial condyle with a clear and deep gap. It has a rounded shape with a convex articular surface; the other is medially located at the orbital process and has a rounded shape with a flat articular surface.

The mandibular process exhibits three condyles, arranging in a triangular shape with a distinct fossa (relatively deep in *Rynchops*). The medial condyle shows a rostrocaudally elongate oval outline with a convex articular surface. It shows a clear margin with the quadrate body. The lateral condyle shows a lateromedially elongate oval shape with a convex articular surface. The caudal condyle appears to be a rounded, slant bulge. It is caudally positioned in *Chroicocephalus* and caudomedially located in *Rynchops*, respectively.

The quadrate body of the silver gull (*Chroicocephalus*) is slightly straight and bears a right-angled lateral crest and a straight medial crest. The quadrate body of the Black skimmer (*Rynchops*) is laterally curved and bears a curved lateral crest and a straight medial crest. The caudomedial depression is shallow and wide on the Laridae quadrate. Two pneumatic foramina are present on the Laridae quadrate: the caudomedial foramen is accommodated at the caudomedial depression on both Laridae quadrates in this study; the basiorbital foramen is medially located at the quadrate body on the silver gull quadrate only, close to the pterygoid condyle.

**Eurypygiformes:** Eurypygidae (Plate 27)

In Eurypygidae (*Eurypyga helias*), the otic process shows a shallow and distinct intercapitular incisure between two capitula. The squamosal capitulum exhibits a dorsoventrally elongate oval outline with a convex articular surface, pointing rostrodorsally. The otic capitulum exhibits a rounded shape with a flat articular surface, facing caudodorsally.

The orbital process is elongate and slender with a high aspect ratio. Its tip is flat, broad and is tapered into horizonal, pointing rostrally. The basiorbital fossa is shallow and broad.

The quadratojugal cotyle is slightly standing laterally and shows a cup-like shape with a thick and oval margin. It has a deep fossa and is dorsally adjacent to the lateral condyle. The rostroventral margin of the quadratojugal cotyle expands slightly, forming a flat articular surface for jugal bar. The pterygoid condyle is slightly standing rostrally and is significantly dorsomedially located at the medial condyle with a narrow gap. It shows a lateromedially elongate oval shape with a convex articular surface.

The mandibular process shows three condyles, arranging in a L- or triangular shape, and bears a shallow fossa at the centre position. Both medial and lateral condyles show rostrocaudally elongate oval outline with convex articular surfaces. The caudal condyle is an elongate, thick bulge and caudally positioned along the quadrate body. It shows a smooth articular surface.

In Eurypygidae, the quadrate body is relatively straight and bears a right-angle lateral and medial crests. The caudomedial depression is shallow and unclear between two capitula. Only one pneumatic foramen (rostromedial foramen) is medially located at the quadrate body, surrounded by the medial crest and the orbital process. The caudomedial fossa, which is located at the caudomedial depression, might also be related to the air sac system in bird skull.

**Eurypygiformes:** Rhynochetidae (Plate 27)

In Rhynochetidae (*Rhynochetos jubatus*), the otic process has a shallow and distinct intercapitular incisure between the squamosal and otic capitula. Similar as its sister group (Eurypygidae), the squamosal capitulum is as high and large as the oitc capitulum.

The squamosal capitulum shows a dorsoventrally elongated rectangle-with-rounded-corner outline (though the lateral margin is inward) with a convex articular surface and is oriented rostrodorsally. The otic capitulum shows a rounded shape with a convex articular surface and faces dorsally.

The orbital process is slightly robust, elongate, and exhibits a high aspect ratio. Its tip is flat, broad, and tapers into subvertical, pointing rostrodorsally. Besides, the basiorbital fossa is shallow and broad.

The quadratojugal cotyle is slightly standing laterally and has a cup-like shape with a thick and oval margin. It has a deep articulation and is dorsally adjacent to the lateral condyle. The rostroventral margin of the quadratojugal cotyle expands, forming a flat articular surface for jugal bar. The articulation with pterygoid shows two connections: pterygoid condyle is standing slightly and is dorsally separated from the medial condyle with a wide gap. It shows a lateromedially elongate oval shape with a convex articular surface. The orbitopterygoid facet is medially located at the orbital process with a flat articular surface.

The mandibular process shows three condyles, arranging in a L- or triangular shape, and bears a deep fossa at the centre position. The medial condyle shows a L-shape with a saddle articular surface, separating into medial and lateral parts (lateral trochlea). The lateral trochlea appears to be a prominent convex articular surface and is much larger than the medial part, while the medial part shows a flat articular surface. The lateral condyle shows a rounded or squared shape with a convex articular surface. The caudal condyle appears to be a titled, thick bulge and is caudally positioned along the quadrate body. It has a convex articular surface and faces caudoventrally.

In Rhynochetidae, the quadrate body is straight, slender and bears a right-angle lateral crest and a minorly curved medial crest. The caudomedial depression is shallow, and two pneumatic foramina are present on the Kagu quadrate. The postsquamosal capitulum foramen is caudoventrally located at the squamosal capitulum with a tiny opening. The rostromedial foramen is medially located at the quadrate body and is surrounded by the medial crest and the orbital process.

**Phaethontiformes:** Phaethontidae (Plate 27)

The quadrate of Phaethontidae (*Phaethon lepturus*) has a shallow and narrow intercapitular incisure between two capitula on the otic process. Unlike most avian quadrate, the otic capitulum is slightly higher and larger than the squamosal capitulum. The squamosal capitulum shows a lateromedially elongate oval shape with a convex articular surface, and it points dorsolaterally. The otic capitulum also has a lateromedially elongate oval shape with a markedly convex articular surface, facing dorsally. It is slightly located rostrally at the otic process.

The orbital process is robust, slight short and bears a low aspect ratio. Its tip is flat, slant and broad, pointing rostrally. Besides, the basiorbital fossa is shallow and broad.

The quadratojugal cotyle is laterally standing and has a cup-like shape with a thick and circular margin (though its dorsal and ventral margins are slightly inward). It has a deep articulation (Saiff, 1978) and is adjacent to the lateral condyle. The pterygoid condyle is slightly standing and has a rounded shape with a convex articular surface. It is dorsally located at the medial condyle with a small gap, but it is medially adjacent to the medial condyle.

The mandibular process has three condyles, arranging in a L-shape with a shallow fossa at the centre position. Similar as Rhynochetidae, the medial condyle in Phaethontidae also has a L-like shape with a saddle articular surface, separating into medial and lateral parts (lateral trochlea). The medial part has an oval shape with a prominently convex articular surface (Saiff, 1978), and it is much larger than the lateral part. On the other hand, the lateral part only has a flat articular surface. The lateral condyle is continuous with the caudal condyle with a J-shape, and it has a convex articular surface. The caudal condyle is a slightly slant (sub-horizontal) and slender bulge caudally attached with quadrate body, and it has a cashew shape with a flat articular surface (Saiff, 1978).

In Phaethontidae, the quadrate body is laterally curved due to a curved lateral crest and a minorly curved medial crest. The caudomedial depression is absent on the white-tailed tropicbird (*Phaethon lepturus*) quadrate due to a lack of the tympanic crest. Only one pneumatic foramen (basiorbital foramen) is medially present on the quadrate body of the white-tailed tropicbird – it is close to the pterygoid condyle and is relatively large (Saiff, 1978).

**Gaviiformes:** Gaviidae (Plate 27)

In Gaviidae, the otic process does not have a clear intercapitular incisure between two capitula. The squamosal capitulum is as high as the otic capitulum in *Gavia stellata*, but it is slightly lower than the otic capitulum in *G. arctica*. It has an equal size as the otic capitulum. In both species. The squamosal capitulum has a rostrocaudally elongate oval shape with a convex articular surface in both two species, and it faces rostrodrossally. The otic capitulum has a lateromedially elongate oval shape in *G. stellata* but it has a rostrocaudally elongate oval in *G. arctica*. It has a convex articular surface and points either dorsomedailly in *G. stellata* or dorsocaudally in *G. arctica*.

The orbital process in Gaviidae is significantly slender and elongate with a high aspect ratio. Its tip is pointed and faces rostrally. It has a distinct ridge ventrally located at the orbital process. Besides, the basiorbital fossa is shallow and narrow in Gaviidae.

The quadratojugal cotyle shows a cup-like shape with a thick and circular margin. Its rostrodorsal margin is prominent and laterally protruding. It has a deep fossa and is adjacent to the lateral condyle. Unlike most avian groups, the quadratojugal cotyle slightly faces ventrolaterally. The articular contact with pterygoid on Gaviidae quadrate has two connections: one (pterygoid condyle) is rostrally standing. It is dorsally located at the medial condyle with a small gap, but it is medially adjacent to the medial condyle. It has a lateromedially slender oval shape with a convex articular surface. The other (orbitopterygoid facet) is medially located at the orbital process with a flat and oval articulation.

The mandibular process shows three condyles, arranging in a twisted L shape (medial margin is inwardly curved) with a shallow fossa at the centre position (only in *G. stellata*). The medial condyle in Gaviidae has a lateromedially elongate oval shape with a saddle articular surface, separating into medial and lateral parts (lateral trochlea). The medial part has an oval shape with a prominently convex articular surface, and it is much larger than the lateral part. On the other hand, the lateral part only has a flat articular surface. Also, the medial condyle has a clear margin with the quadrate body. The lateral condyle has a rounded shape with a convex articular surface. The caudal condyle is a slightly slant (sub-horizontal) and slender protrusion caudally attached with quadrate body, and it has a flat articular surface.

In Gaviidae, the quadrate body is slightly straight and slender due to a right-angle lateral and medial crest. The caudomedial depression is absent on the Gaviidae quadrate due to a lack of the tympanic crest. The pneumatic foramen is also absent in Gaviidae. However, two distinct fossae are located at the quadrate body: one is rostroventrally located at the otic process, and the other is laterally located at the quadrate body, below the otic process.

**Sphenisciformes:** Spheniscidae (Plate 27-28)

In Spheniscidae, the otic process has a distinct and deep intercapitular incisure between the squamosal and otic capitulum (Saiff, 1976), especially in *Eudyptes chrysolophus* and *Icadyptes salasi* (Ksepka et al., 2008). However, it is shallow in *Paraptenodytes antarcticus* (Bertelli et al., 2006). The squamosal capitulum is as high as the otic capitulum in extant penguins (*Eudyptes chrysolophus* and *Spheniscus humboldti* in this study), but it is slightly lower than the otic capitlum in extinct taxon, such as *Madrynornis mirandus* (Degrange et al., 2018) and *Kairuku waitaki* (Ksepka et al., 2012). The squamosal capitulum is much larger than the otic capitulum in *Spheniscus humboldti* and *Paraptenodytes antarcticus* (Bertelli et al., 2006), but it is slightly smaller in *Eudyptes chrysolophus* and *Madrynornis mirandus*.

The squamosal capitulum has a cashew outline in *Spheniscus humboldti* while it has a rectangle-with-rounded-corner outline in *Eudyptes chrysolophus* and *Madrynornis mirandus*. The outline of the squamosal capitulum is medially curved only in extant species. The squamosal capitulum in penguin usually has a convex articular surface and faces dorsolaterally or dorsally. The otic capitulum, on the other hand, has a rostrocaudally elongate oval shape in Spheniscidae, and it has a relatively convex articular surface, facing dorsomedially. A distinct and wide notch is medioventrally located at the otic capitulum only on extant penguin quadrates.

The orbital process in Spheniscidae is robust with a relative high aspect ratio, and its tip turns to become a pointed shape (*Spheniscus humboldti*), a rounded shape (*Paraptenodytes antarcticus*), a flat shape (*Eudyptes chrysolophus* and *Icadyptes salasi*), or a hammer-like shape (*Madrynornis mirandus*), pointing rostrodorsally. A distinct ridge, orbital crest, is laterally present on the orbital process, extending form its tip to the base of the orbital process in either extant penguins or extinct taxon (Bertelli et al., 2006; Degrange et al., 2018; Ksepka et al., 2008). Besides, the basiorbital fossa in most Spheniscidae is deep and slightly wide, but it is much shallower in *Madrynornis mirandus* than other penguin quadrates.

The quadratojugal cotyle in Spheniscidae has a cup-like shape with a thick and oval margin. It has a significantly deep articular fossa (Degrange et al., 2018; Ksepka et al., 2008) and is adjacent to the lateral condyle (Saiff, 1976). There is a distinct ridge dorsally located at the quadratojugal cotyle in either extant penguin or extinct species (e.g., *Paraptenodytes antarcticus*, Bertelli et al., 2006). The pterygoid condyle is rostrally standing, and it has an oval shape or a knob-like structure with a convex articular surface (Bertelli et al., 2006; Degrange et al., 2018). It is dorsally located and distinctly separated from the medial condyle.

The mandibular process on Spheniscidae quadrate has three condyles, arranging in a L-shape, and bears a shallow fossa in *Spheniscus humboldti* or a shallow intercondylar sulcus between medial and lateral condyles (Saiff, 1976). The medial condyle shows a great morphological diversity among Spheniscidae: in *Spheniscus humboldti*, the medial condyle shows a lateromedially elongate oval or rectangle-with-rounded-corners shape with a saddle articular surface, separating into medial and lateral parts (lateral trochlea). The medial part has an oval shape with a prominent convex articular surface, and it is much larger than the lateral part. On the other hand, the lateral part only has a flat articular surface. However, in *Eudyptes chrysolophus* and *Madrynornis mirandus*, the medial condyle has a rostrocaudally elongate oval shape with a prominently convex articular surface, and it is not bifurcated into two articular surfaces with the mandible. In all penguin quadrates, the medial condyle shows a clear margin with the quadrate body and rostrally expands to form a tip (Saiff, 1976). The lateral condyle is confluence with the caudal condyle in extant species but not in *Madrynornis mirandus* whose lateral condyle is distinct with the caudal condyle. The lateral condyle has a rounded shape with a slightly convex articular surface in *Spheniscus humboldti* and *Madrynornis mirandus*, but it has an oval shape with a flat articular surface in *Eudyptes chrysolophus*. The caudal condyle has a thick and elongate protrusion caudally attached with quadrate body, and it has a slightly convex articular surface, facing caudally. However, the caudal condyle in *Icadyptes salasi* is relatively large and has a rounded shape with a convex articular surface (Fig. 6 in Ksepka et al., 2008).

The Spheniscidae quadrate body is slightly straight due to a right-angle lateral and minorly curved medial crest, but it is much wider on *Eudyptes chrysolophus* quadrate. The caudomedial depression is absent on the Spheniscidae quadrate due to a lack of the tympanic crest. On most Spheniscidae quadrate, the subcapitular tubercle is rostroventrally located at the squamosal capitulum with a distinct protrusion. However, the subcapitular tubercle is lateroventrally located at the squamosal capitulum with a rounded shape on *Icadyptes salasi* (Ksepka et al., 2008) and *Paraptenodytes antarcticus* (Bertelli et al., 2006) quadrates, and it is much closer to the squamosal capitulum than other penguin quadrates. The pneumatic foramen is also absent in either extant penguin quadrates or their extinct relatives (Bertelli et al., 2006; Ksepka et al., 2008; Saiff, 1976).

**Procellariiformes:** Diomedeidae (Plate 28)

The Diomedeidae (*Phoebetria palpebrata*) quadrate has a wide intercapitular incisure between two capitula on its otic process. The squamosal capitulum is much higher and larger than the otic capitulum. The squamosal capitulum has a lateromedially elongate oval shape with a convex articular surface, and it faces dorsally. The otic capitulum has a rounded or squared shape with a convex articular surface, pointing medially or dorsomedially.

The orbital process in *Phoebetria palpebrata* is robust and elongate with a high aspect ratio. Its tip is flat and broad, and turns to be medially slant, facing rostrally. Besides, the basiorbital fossa in Diomedeidae is deep and wide.

The quadratojugal cotyle of Diomedeidae quadrate has a cup-like shape with a thick and circular margin. It has a deep fossa, adjacent to the lateral condyle. Its ventral margin ventrally expands, forming a flat articular surface for jugal bar attachment. The pterygoid condyle is rostrally standing, and it has an oval or rounded shape with a convex articular surface. It is dorsally located at the medial condyle with a small gap, but it is medially adjacent to the medial condyle. Unlike most other avian quadrates, the pterygoid condyle in *Phoebetria palpebrata* is located much medially at the quadrate body.

The mandibular process on Diomedeidae quadrate has three condyles, arranging in a L-shape with a shallow fossa at the centre position. The medial condyle in Diomedeidae has a lateromedially elongate rectangle shape with a saddle surface, separating into medial and lateral parts (lateral trochlea). The medial part has an oval shape with a prominently convex articular surface, and it is much larger than the lateral part. The lateral part only has a flat articular surface. Also, the medial condyle has a clear margin with the quadrate body and a distinct rostral tip. The lateral condyle has a rounded shape with a convex articular surface, while the caudal condyle is a thick, slant and elongate protrusion caudally attached with the quadrate body and has a slightly convex articular surface.

The Diomedeidae quadrate body is slightly straight due to a right-angle lateral and a minorly curved medial crest. The caudomedial depression is absent on the Light-mantled Albatross (*Phoebetria palpebrata*) quadrate. Only one pneumatic foramen (basiorbital foramen) is medially located at the Light-mantled Albatross quadrate body, close to the base of the orbital process and pterygoid condyle.

**Procellariiformes:** Oceanitidae (Plate 28)

In Oceanitidae (*Pelagodroma marina*), the oitc process has a wide intercapitular incisure between the squamosal and otic capitulum (Piro and Hospitaleche, 2019). The squamosal capitulum is as high as otic capitulum, but the otic capitulum is much larger than the otic capitulum. The squamosal capitulum has a rounded or squared shape with a rostrocaudally convex articular surface, and it faces dorsolaterally. The otic capitulum has a lateromedially elongate and slender oval shape with a flat articular surface and is oriented caudodorsally (Piro and Hospitaleche, 2019). A shallow and wide notch is ventrally located at the otic capitulum.

Similar as Diomedeidae (*Phoebetria palpebrata*), the orbital process is robust and elongate with a high aspect ratio in *Pelagodroma marina*. Its tip is flat and broad, and it turns to be medially slanted (close to sub-horizonal), pointing rostrally. Besides, Oceanitidae quadrate has a deep and wide basiorbital fossa.

Unlike other Procellariiformes or Phaethoquornithes (Eurypygimorphae+ Aequornithes), the quadratojugal cotyle in Oceanitidae has a saddle-like shape (dorsal and ventral margins are not well-developed) with a thick margin and shallow fossa, adjacent to the lateral condyle. The pterygoid condyle is only slightly standing and has an oval shape with a slightly convex articular surface (Piro and Hospitaleche, 2019). The pterygoid condyle is dorsally separated from the medial condyle with a small gap.

The mandibular process on Oceanitidae quadrate has three condyles, arranging in a L- or a boomerang-like shape with a shallow fossa at the centre position. The medial condyle in Oceanitidae has a rostrocaudally elongate oval shape with a prominently convex articular surface. The lateral condyle has a lateromedially elongate oval shape with a slightly convex articular surface. The caudal condyle has an elongate protrusion, but it is not as standing as some waterbirds, such as *Phoebetria palpebrata*. It aligns horizonally and is caudally attached with the quadrate body.

The Oceanitidae quadrate body is slightly curved due to a curved lateral and medial crest. The subcapitular tubercle is ventrolaterally located at the squamosal capitulum, as shown in a tiny oval outline with a slightly convex surface. The caudomedial depression is absent due to a lack of the tympanic crest. One the White-faced storm petrel (*Pelagodroma marina*) quadrate, one pneumatic foramen (basiorbital foramen) is medially located at the quadrate body with a narrow opening, and it is far from the pterygoid condyle (Piro and Hospitaleche, 2019).

**Procellariiformes:** Hydrobatidae (Plate 28)

In Hydrobatidae (*Hydrobates leucorhous*/*Oceanodroma leucorhoa*), the otic process has a wide and shallow intercapitular incisure between two capitula. The squamosal capitulum is much higher and larger than the otic capitulum. Both squamosal and otic capitulum have a lateromedially elongate oval shape with a slightly convex articular surface, pointing dorsolaterally and dorsomedially, respectively. A wide and shallow notch is ventrally located at the otic capitulum.

In *Hydrobates leucorhous*, the orbital process is robust and elongate with a high aspect ratio. Its tip is flat and broad, and it turns to be medially slanted (close to sub-horizonal), pointing rostrodorsally. Besides, Hydrobatidae quadrate has a deep and wide basiorbital fossa.

The quadratojugal cotyle in Hydrobatidae has a cup-like shape with a thick and circular margin (though its dorsal and ventral margins are slightly inward). It has a deep articular fossa and is adjacent to the lateral condyle. The pterygoid condyle is only slightly standing and has a slender oval shape with a slightly convex articular surface. It is dorsally located at the medial condyle with a deep gap, but it is medially adjacent to the medial condyle.

The mandibular process on Hydrobatidae quadrate has three condyles, arranging in a L shape with a shallow fossa at the centre position. The medial condyle in Hydrobatidae has a lateromedially elongate rectangle or oval outline with a saddle articular surface, separating into medial and lateral parts (lateral trochlea). The medial part has an oval shape with a prominently convex articular surface, and it is much larger than the lateral part. The lateral part only has a flat articular surface. Also, the medial condyle has a clear margin with the quadrate body and a rostral tip. The lateral condyle has a lateromedially elongate oval shape with a slightly convex articular surface. The caudal condyle is a tiny and elongate protrusion, but it is not as standing as some waterbird quadrates. It aligns horizontally and is caudally attached with its quadrate body.

The Hydrobatidae quadrate body is slightly curved due to a curved lateral and medial crest. Like Oceanitidae, the subcapitular tubercle is also ventrolaterally located at the squamosal capitulum on the Leach's storm petrel (*Hydrobates leucorhous*) quadrate, and it appears a slender oval outline with a slightly convex articular surface. The caudomedial depression is absent due to a lack of the tympanic crest. One Hydrobatidae, one pneumatic foramen (basiorbital foramen) is medially located at the quadrate body as shown in a relatively large opening.

**Procellariiformes:** Procellariidae (Plate 28-29)

In Procellariidae, the otic process shows a wide intercapitular incisure between the squamosal and otic capitulum (Saiff, 1974). The squamosal capitulum is higher than the otic capitulum on most Procellariidae quadrate (e.g., *Ardenna tenuirostris* and *Puffinus puffinus*) but it is as elevated as the otic capitulum on the common diving petrel (*Pelecanoides urinatrix*) and the blue petrel (*Halobaena caerulea*) quadrates. Generally, the squamosal capitulum is much larger than the otic capitulum except for the common diving petrel quadrate. The squamosal capitulum usually exhibits a rostrocaudally elongate oval shape with a slightly convex articular surface, facing dorsolaterally. However, the squmosal capitulum of the common diving petrel and blue petrel quadrate shows a lateromedial elongate ovioid shape with a convex articular surface and faces dorsolaterally. The otic capitulum usually shows a lateromedially elongated shape with a convex articular surface, facing dorsally or caudodorsally. Surprisingly, the otic capitulum of the Short-tailed shearwater (*Ardenna tenuirostris*) shows a rostrocaudally elongate oval shape with a convex articular surface, facing dorsomedially.

The orbital process of the Procellariidae quadrate is robust and elongate with a high aspect ratio, pointing rostrally. Its tip is flat and slant (sub- horizonal) in most Procellariidae. Nevertheless, the tip of the orbital process is pointed and turns to be horizontal on the common diving petrel quadrate. The basiorbital fossa is wide on the Procellariidae quadrate and significantly deeper in a few species, such as Manx shearwater (*Puffinus puffinus*) and common diving petrel.

The quadratojugal cotyle of the Procellariidae quadrate appears a cup-like shape with a thick and oval margin, and its dorsal and ventral margin is slightly inward. It has a deep articulation and is adjacent to the lateral condyle (Saiff, 1974). The pterygoid condyle is standing rostrally on Procellariidae quadrate, as shown in an oval shape with a distinct convex articular surface. However, the pterygoid condyle appears an oval shape with a flat articular surface on the common diving petrel quadrate. Like other procellariform quadrates, the pterygoid condyle is dorsally separated from the medial condyle with a wide gap on the Procellariidae quadrate, but it is medially adjacent to the medial condyle.

In Procellariidae, the mandibular process has three condyles, arranging in a L shape with a shallow fossa (intercondylar sulcus) at the centre position (Saiff, 1974). The medial condyle shows a L shape outline with a saddle surface, separating into medial and lateral parts (lateral trochlea). The medial part has a rostrocaudally elongate oval shape with a significantly prominent convex articular surface, while the lateral part only appears a flat articular surface (Saiff, 1974). Besides, the medial condyle has a distinct margin with the quadrate body. The lateral condyle is confluent with the caudal condyle on all Procellariidae quadrate, forming a cashew-like or a rectangle-with-rounded-corner shape with a convex articular surface. The lateral condyle shows a rounded shape with a convex articular surface, while the caudal condyle appears to be an elongate protrusion. The caudal condyle turns to be slant on most Procellariidae quadrate. However, in White-headed petrel (*Pterodroma lessonii*), common diving petrel, and blue petrel, the caudal condyle appear to be horizontally positioned along the quadrates.

The Procellariidae quadrate body is slightly straight due to a right-angle lateral and a minorly curved medial crest. The caudomedial depression is also absent on the Procellariidae quadrate due to a lack of the tympanic crest. The subcapitular tubercle is present on a few Procellariidae quadrate, such as Northern fulmar (*Fulmarus glacialis*), Manx shearwater, Common diving petrel, and Southern giant petrel (*Macronectes giganteus*). It is lateroventrally located at the squamosal capitulum with a standing oval shape. The Manx shearwater and Common diving petrel quadrates lack any pneumatic foramen. The pneumaticity still appears on some Procellariidae quadrates. For instance, the basiorbital foramen is medially located at the quadrate body and close to the pterygoid condyle with a large size on most Procellariidae quadrates (Saiff, 1974). The postcapitular foramen is caudally located at the otic process with a large size on the Northern fulmar and Southern giant petrel quadrates.

**Ciconiiformes:** Ciconiidae (Plate 30)

In Ciconiidae, the otic process does not have a clear intercapitular incisure between the squamosal and otic capitulum, but these two capitula are widely separated to each other. The squamosal capitulum is slightly higher but much larger than the otic capitulum. The squamosal capitulum has a rounded shape with a rostrocaudlly bending articular surface and it faces dorsally or rostrodorsally. The otic ccaptiulum shows a rostrocaudally elongate oval shape with a slightly convex articular surface, facing caudodorsally.

The orbital process of Ciconiidae quadrate is robust, elongate and significantly thick with a slightly high aspect ratio (Saiff, 1978). The tip of this process is flat and broad, pointing rostrally (Oliveira et al., 2019). The basiorbital fossa of Ciconiidae quadrate is shallow and wide (Oliveira et al., 2019).

The quadratojugal cotyle in Ciconiidae has a cup-like shape with a thin margin. It has a deep fossa, but it is distinct to the lateral condyle with a small gap. The ventral margin of the quadratojugal cotyle ventrolaterally expands, forming a flat surface for bone articulation. The articulation with pterygoid on Ciconiidae quadrate varies in different species. In *Ciconia cicionia*, the articulating process with pterygoid rostrally projects with a round shape and slightly convex articular surface, and it is widely separated with the medial condyle. In *Leptoptilos crumenifer*, however, this articulation has two connections: one (pterygoid condyle) is dorsally separated from the medial condyle with a wide gap, but it is medially adjacent to the medial condyle. It has a slender oval shape with a slightly convex articular surface; the other (orbitopterygoid facet) is medially located at the orbital process with a flat articular surface and it is close to the base of the orbital process.

The mandibular process on Ciconiidae quadrate has three condyles, arranging in a triangle shape with a shallow fossa at the centre position. The medial condyle has a lateromedially elongate oval shape with a saddle articular surface, separating into medial and lateral parts (lateral trochlea). The medial part has an oval shape with a prominently convex articular surface, and it is much larger than the lateral part. The lateral part only appears a square outline with a flat articular surface. The lateral condyle shows a rostrocaudally elongate oval shape with a convex articular surface (Oliveira et al., 2019). The caudal condyle has a thick elongate protrusion with a slightly convex articular surface and aligns horizontally.

The Ciconiidae quadrate body is slightly straight and wide due to a minorly curved lateral and medial crest. The caudomedial depression is shallow and wide. The subcapitular tubercle is ventrolaterally located at the squamosal capitulum on the Ciconiidae quadrate and it has a wide and oval shape with a slightly convex surface. Two pneumatic foramina are present on the Ciconiidae quadrate: the postcapitular foramen is caudally located at the otic process with a relatively large size; the basiorbital foramen is medially located at the quadrate body and below the basiorbital fossa (Saiff, 1978).

**Suliformes:** Fregatidae (Plate 30)

In Fregatidae (*Fregata aquila*), the otic process has a shallow intercapitular incisure, widely separating two capitula (Saiff, 1978). The squamosal capitulum is slightly lower than otic capitulum, but it has an equal size as the otic capitulum. Both squamosal and otic capitulum have lateromedially elongate oval shapes with slightly convex articular surfaces, facing dorsally.

The orbital process in *Fregata aquila* is robust and elongate with a high aspect ratio. Its tip is flat and broad and turns to be slant (or sub-vertical), pointing rostrally. Besides, Fregatidae quadrate has a shallow and wide basiorbital fossa.

The quadratojugal cotyle in Fregatidae is laterally standing and it has a cup-like shape with a thick and circular margin (thought its caudal margin is inward or underdeveloped). It has a deep joint articulation, adjacent to the lateral condyle. The rostral margin of the quadratojugal cotyle slightly expands laterally, forming a flat surface for jugal attaching. In Fregatidae, two articulations on its quadrate connect with pterygoid: one (pterygoid condyle) has a rectangle-with-rounded-corner shape with a flat articular surface. It is dorsally located at the medial condyle with a short gap, but it is medially adjacent to the medial condyle. Also, compared with that of other avian clades, the pterygoid condyle of Fregatidae quadrate is significantly located medially. The other, orbiopterygoid facet, is medially located at the orbital process, close to the base of the orbital process. It has a distinct oval shape with a flat articular surface.

The mandibular process on Fregatidae quadrate has three condyles, arranging in a L shape with a deep fossa at the centre position (Saiff, 1978). The medial condyle has a rostrocaudally elongate oval outline with a prominent convex articular surface. Also, it has a clear margin with the quadrate body. The lateral condyle has a lateromedial elongate oval shape with a convex articular surface. The caudal condyle is a thick, elongate but slant protrusion and has a slightly convex articular surface, facing caudally.

The Fregatidae quadrate body is lateromedially wide and slightly medially curved due to a minorly curved medial crest. Only one pneumatic foramen (basiorbital foramen) is medially located at the quadrate body (Saiff, 1978) and dorsally located at the pterygoid condyle with a relatively large size.

**Suliformes:** Sulidae (Plate 30)

In Sulidae, the otic process has a wide and deep intercapitular incisure between its squamosal and otic capitulum (Saiff, 1978). The squamosal capitulum is as high as otic capitulum, but it is larger than otic capitulum. Both capitula have a rostrocaudally elongate oval shape with a convex articular surface, facing dorsally.

The orbital process in Sulidae is robust with a slightly lower aspect ratio, and it has a triangle shape in lateral view. Its tip is pointed, facing rostrally. The basiorbital fossa is deep and wide in *Sula dactylatra*, but it is shallow and narrow in *Morus bassanus*.

The quadratojugal cotyle of Sulidae has a cup-like shape with a thick and circular margin. It has a deep articulation, close to the lateral condyle. Like other Suliformes (e.g., *Fregata aquila* and *Leucocarbo atriceps*), the rostral margin of the quadratojugal cotyle slightly laterally and rostrally expand, forming an attachment for jugal bar. Similar as other Suliformes, the pterygoid condyle of Sulidae quadrate is much ventromedially adjacent to the medial condyle. It shows a dorsoventrally elongate oval shape with a convex articular surface. Like Phalacrocoracidae (see below), a distinct crest is caudomedially located at the pterygoid condyle.

The mandibular process on Sulidae quadrate has three condyles, arranging in a V or U shape with a deep fossa at the centre position size (Saiff, 1978). The medial condyle has a significantly concave articular surface, forming a clear notch below the quadrate body. It has two articular surfaces, separating into medial and lateral part (lateral trochlea). The medial part has a rostrocaudally elongate oval shape with a prominently convex articular surface. The lateral part only has a flat articular surface with distinct margin with the quadrate body. The lateral condyle is rostrocaudally elongate with an oval shape and a flat articular surface. The caudal condyle is a thick protrusion and has a slightly convex articular surface, facing caudally.

The Sulidae quadrate body is straight and lateromedially wide due to a minorly curved lateral and medial crest. Either the caudomedial depression or caudomedial fossa is absent on its quadrate. The subcapitular tubercle on Sulidae quadrate is a tiny protrusion, ventrolaterally located at the squamosal capitulum. In Sulidae, one pneumatic foramen (basiorbital foramen) is medially located at the quadrate body, and it is below the basiorbital fossa with a relatively large size (Saiff, 1978).

**Suliformes:** Anhingidae (Plate 30)

The otic process of the Anhingidae (*Anhinga anhinga*) quadrate exhibits a narrow intercapitular incisure between squamosal and otic capitulum (Saiff, 1978). Unlike most avian quadrates, these two capitula rostrocaudally line up on Anhingidae quadrate, instead of lateromedially. The squamosal capitulum is as high as otic capitulum, but the otic capitulum is significantly larger than the squamosal capitulum. The squamosal capitulum shows a rostrocaudally elongate oval shape with a convex articular surface, facing rostrodorsally. The otic capitulum shows a squared shape with a flat articular surface, facing dorsally.

In *Anhinga anhinga*, the orbital process is robust but thin with a slightly high aspect ratio and a triangle shape in lateral view. Its tip is pointed and faces rostrally. Besides, the basiorbital fossa is wide and deep in *Anhinga anhinga* quadrate.

The quadratojugal cotyle of Anhingidae quadrate has a cup shape with a thick and circular margin. It shows a significantly deep fossa and is adjacent to the lateral condyle. However, different with other suliform quadrates, the rostral margin of the quadratojugal cotyle does not rostrally expand to form a surface for bone attachment in *Anhinga anhinga*. The pterygoid condyle of Anhingidae quadrate is adjacent to the medial condyle, unlike that of other sulliform quadrates, and it appears an oval shape with a flat articular surface. There is a distinct ridge/crest caudomedially located at the pterygoid condyle.

The mandibular process on Anhingidae quadrate has three condyles, arranging in a V shape with a deep fossa at the centre position. The medial condyle has a rostrocaudally elongate oval shape with a prominent convex articular surface (Saiff, 1978). The lateral condyle has a lateromedially elongate oval shape with a slightly flat articular surface. The caudal condyle shows a thick and elongate protrusion and has a convex articular surface, facing caudally.

In Anhingidae, the quadrate body is straight and wide due to a straight lateral and medial crest. Either the caudomedial depression or caudomedial fossa is absent on its quadrate. The subcapitular tubercle on Anhingidae quadrate appear a platform-like protrusion, rostroventrally located at the squamosal capitulum. In Anhingidae, only one pneumatic foramen (rostromedial foramen) is medially present on its quadrate body (Saiff, 1978). It is compressed by the medial crest and a distinct ridge with a narrow and slender opening.

**Suliformes:** Phalacrocoracidae (Plate 31)

In Phalacrocoracidae (*Leucocarbo atriceps*), its quadrate has a narrow but very deep intercapitular incisure on the otic process (Saiff, 1978). The squamosal capitulum is higher and larger than the otic capitulum in the rostral view. Both two capitula show rostrocaudally elongate oval outlines with convex articular surfaces, facing dorsally and dorsomedially, respectively.

The orbital process of the *Leucocarbo atriceps* quadrate is significantly slender with a high aspect ratio (Saiff, 1978). The tip of its orbital process is pointed, facing rostrally. The basiorbital fossa on the Phalacrocoracidae quadrate is shallow and narrow.

The quadratojugal cotyle is laterally protruding in Phalacrocoracidae and it has a cup-like shape with a thick and droplet-like margin. The cotyle of the *Leucocarbo* quadrate also shows a deep articular fossa, adjacent to the lateral condyle. Like Fregatidae quadrate (*Fregata aquila*), the rostral margin of the quadratojugal cotyle expands rostrolaterally, forming a flat articular surface for jugal. Unlike most avian quadrates, the pterygoid condyle of Phalacrocoracidae quadrate is much dorsomedially located at the medial condyle with a deep and wide gap. Its pterygoid condyle appears a rounded outline with a slightly convex articular surface. A distinct crest is caudomedially located at the pterygoid condyle.

The mandibular process on Phalacrocoracidae quadrate has three condyles, arranging in a V shape with a deep fossa at the centre position. The medial condyle shows a rostrocaudally elongate lens-like outline with a prominent convex articular surface. It has a clear margin with the quadrate body. The lateral condyle has a lateromedially elongate lens-like shape with a flat articular surface. The caudal condyle shows a thick protrusion with a slightly convex articular surface, facing caudally.

Different with most avian quadrates (e.g., straight or slightly medially curved body shape), the quadrate body of Phalacrocoracidae quadrate is extremely medially curved due to a significantly curved medial crest. Neither the caudomedial depression nor caudomedial fossa is present on its quadrate. The subcapitular tubercle on the Phalacrocoracidae quadrate exhibits a tiny spike-like shape, ventrolaterally located at the squamosal capitulum. In Phalacrocoracidae, only one pneumatic foramen (basiorbital foramen) is medially located at the quadrate body (Saiff, 1978), and it is close to the basiorbital fossa with a relatively small size.

**Pelecaniformes:** Threskiornithidae (Plate 31)

The otic process of the Threskiornithidae (*Eudocimus ruber*) quadrate has a narrow but deep intercapitular incisure between squamosal and otic capitulum, and these two capitula are close to each other. The squamosal capitulum is slightly larger than the otic capitulum. The squamosal capitulum has a rostrocaudally elongate oval shape, facing dorsolaterally. The otic capitulum has a gourd-like shape (its lateral and medial margin is curved and inward) with a flat articular surface, facing dorsally. Below the otic capitulum, there is a clear notch.

On the Scarlet ibis (*Eudocimus ruber*) quadrate, the orbital process is rostrally elongate and dorsoventrally deep, and its tip is slant and flat, pointing rostrally. A tiny spike is ventrally located at the orbital process. The basiorbital fossa of the scarlet ibis quadrate is wide and shallow.

The quadratojugal cotyle of the Threskiornithidae quadrate appears a cup shape with a thick margin, although its rostral margin is slightly inward. Its cotyle is a deep articulation, adjacent to the lateral condyle. The ventral margin of the quadratojugal cotyle ventrolaterally expands, forming a flat articular surface for the jugal bar. The pterygoid condyle of the Scarlet ibis quadrate is dorsally located at the medial condyle with a wide and deep gap, and it has an oval shape with a convex articular surface.

The mandibular process on Threskiornithidae quadrate has three condyles, arranging in a triangle shape. The medial condyle has a lens-like shape with a saddle articular surface, separating into medial and lateral part (lateral trochlea). The medial part shows a rostrocaudally elongate and oval outline with a prominent convex articular surface, while the lateral part shows a triangle outline with a flat articular surface. The lateral condyle is lateromedially elongate and exhibits a slender and oval shape with a flat articular surface. The caudal condyle is a thick but short protrusion with a flat articular surface.

In Threskiornithidae, the quadrate body is straight due to a slightly curved lateral and medial crest. The lateral crest is significantly standing and thick on the scarlet ibis quadrate, and its caudomedial depression is wide and deep. Far from the otic process, there is a thick protrusion caudally located at the quadrate body (Saiff, 1978). The subcapitular tubercle of Threskiornithidae quadrate shows a rounded shape with a convex surface, laterally located at the squamosal capitulum. Only pneumatic foramen (caudomedial foramen) is medially located at the quadrate body of scarlet ibis, and it is compressed by the medial and tympanic crest, facing medially.

**Pelecaniformes:** Ardeidae (Plate 31)

In Ardeidae, the otic process exhibits a wide and deep intercapitular incisure between squamosal and otic capitulum (Saiff, 1978). The squamosal capitulum is as elevated as the otic capitulum, but larger than the otic capitulum. The squamosal capitulum has a rostrocaudally elongated oval outline (though its lateral margin is dorsally inward), facing rostrodorsally. The squamosal capitulum is much lateromedially slender in *Ardea alba* than that in *Tigrisoma lineatum* and *Ixobrychus minutus*. The otic capitulum usually shows a rostrocaudally elongate oval outline facing caudodorsally, but the otic capitulum of *Tigrisoma* quadrate shows a rounded shape.

The orbital process of the Ardeidae quadrate is significantly elongate with a robust base and a high aspect ratio (Saiff, 1978). Generally, the orbital process in Ardeidae is thin, but it is much thicker in *Tigrisoma*. The tip of the orbital process has a spoon-like or a rounded outline, pointing rostrodorsally. The basiorbital fossa of the Ardeidae quadrate is wide and shallow.

The quadratojugal cotyle of the Ardeidae quadrate appears a cup shape with a thick and circular margin. It has a deep articulation, adjacent to the lateral condyle (Saiff, 1978). The pterygoid condyle of Ardeidae is mostly dorsomedially located at the medial condyle, and it has a rounded outline with either a slightly convex articular surface (*Ixobrychus* and *Tigrisoma* quadrate) or a flat articular surface (*Ardea alba*). However, the articulation with pterygoid on *Ixobrychus* quadrate shows the second contact: the orbitopterygoid facet is medially located at the base of the orbital process and shows an oval shape with a flat articular surface.

The mandibular process on the Ardeidae quadrate has three condyles, arranging in a U shape with a deep fossa at the centre position. The medial condyle usually shows a rostrocaudally elongate oval shape with a prominent convex articular surface. However, the medial condyle of *Ardea* quadrate shows two articulation with the lower jaw, separating into medial and lateral part (lateral trochlea). The lateral trochlea of *Ardea* quadrate is tiny and has a triangle outline with a flat articular surface. The lateral condyle has a rounded shape (*Ardea alba* and *Tigrisoma lineatum*) or an oval shape (*Ixobrychus minutus*) with a convex articular surface. The caudal condyle is a thick but short protrusion, aligned sub-horizontally or horizontally in caudal view. It has a convex articular surface facing caudally or caudoventrally.

In Ardeidae, the quadrate body is relatively straight and wide due to a curved lateral and a right-angle medial crest. The caudomedial depression is wide and deep, caudally located at the quadrate body. The subcapitular tubercle on Ardeidae quadrate exhibits a tiny spike-like protrusion, ventrolaterally located at the squamosal capitulum. Only one single pneumatic foramen (caudomedial foramen) is caudomedially located at the Ardeidae quadrate body (Saiff, 1978). It is accommodated in the caudomedial depression, compressed by the medial and tympanic crest, and it faces medially due to the strongly twisted tympanic crest.

**Pelecaniformes:** Scopidae (Plate 32)

In Scopidae (*Scopus umbretta*), the otic process has a shallow and wide intercapitular incisure between squamosal and otic capitulum. The squamosal capitulum is higher than the otic capitulum, but the otic capitulum is much larger than the squamosal capitulum. The squamosal capitulum shows a rounded shape, and its articular surface is rostrocaudally convex, pointing rostrally. The otic capitulum has a lateromedially elongate oval shape with a convex articular surface, facing dorsally.

The orbital process of *Scopus* quadrate is robust, thick, and elongate with a high aspect ratio. Its tip is flat and broad, pointing rostrally. The basiorbital fossa of the Scopidae quadrate is shallow and narrow.

The quadratojugal cotyle of the Scopidae quadrate appears a cup shape with a thick and circular margin. It has a deep fossa and is close to the lateral condyle. The rostroventral margin of the quadratojugal cotyle slightly expands ventrolaterally, forming a flat articular surface for the jugal bar. Though the pterygoid condyle of the *Scopus* quadrate is dorsally separated from the medial condyle with a shallow gap, it is medially adjacent with the medial condyle (Elzanowski et al., 2001). It has an oval shape with a flat articular surface.

The mandibular process on the Scopidae quadrate has three condyles, arranging in a curved triangle shape with a shallow fossa at the centre position. The medial condyle has a lateromedially elongate oval shape with a saddle articular surface, separating into medial and lateral part (lateral trochlea). The medial part shows a rostrocaudally elongate oval shape with a convex articular surface, while the lateral part has a lens-like outline with a flat articular surface. The lateral condyle is confluent with the caudal condyle, and it has a lateromedially elongate oval shape with a slightly convex articular surface. The caudal condyle appears an elongate protrusion with a slightly concave articular surface.

In Scopidae, the quadrate body is relatively straight with a right-angle lateral and medial crest. The caudomedial depression is shallow and wide. Only one pneumatic foramen (rostromedial foramen) is medially present on the quadrate body of the Hamerkop (*Scopus umbrette*), surrounded by the medial crest and the orbital process.

**Pelecaniformes:** Balaenicipitidae (Plate 32)

In Balaenicipitidae (*Balaeniceps rex*), the otic process has a shallow and wide intercapitular incisure between two capitula. The squamosal capitulum is as elevated and large as the otic capitulum. The squamosal capitulum shows a lateromedially elongate oval shape with a convex articular surface, facing dorsally. The otic capitulum appears a lateromedially elongate oval shape, facing dorsally. However, unlike most avian quadrate, the otic capitulum has two articulations with cranium, dividing into two oval facets with a convex articular surface.

The orbital process of the *Balaeniceps* quadrate is thin but dorsoventrally deep with a lower aspect ratio. Its tip is flat and broad, pointing rostrally. The Balaenicipitidae quadrate has a shallow and broad basiorbital fossa.

The quadratojugal cotyle of the Balaenicipitidae quadrate has a cup shape with a thick and oval margin. It has a deep articular surface and is close to the lateral condyle with a small gap. Similar as most Pelecaniformes, the rostroventral margin of the quadratojugal cotyle expands ventrolaterally, forming a flat articular surface for the jugal bone. The articulation with pterygoid on the *Balaeniceps* quadrate has two connections: one (pterygoid condyle) is separated from the medial condyle with a distinct gap, but it is also medially adjacent to the medial condyle (Elzanowski et al., 2001). It has a triangle-with-rounded-corner shape with a slightly convex articular surface. The other (orbiopterygoid facet) is medially located at the orbital process and shows an oval shape with a flat articular surface. A distinct and thick crest is medially located at the quadrate body, compared to the pterygoid condyle.

The mandibular process on the Balaenicipitidae quadrate has three condyles, arranging in a T shape with a clear fossa at the centre position. The medial condyle shows a significantly concave articular surface, dividing into medial and lateral part (Saiff, 1978). The medial part is strongly slender with a rostrocaudally prominent convex articular surface, while the lateral part has a slightly flat articular surface. The lateral condyle is confluent with the caudal condyle, and it has a rounded shape with a convex articular surface. The caudal condyle appears a thick and slanted with a convex articular surface, facing caudoventrally.

In Balaenicipitidae, the quadrate body is relatively straight and wide with a right-angle lateral and medial crest. In the caudal view, there is a shallow and wide caudomedial depression. Only one pneumatic foramen (basiorbital foramen) is medially located at the Shoebill (*Balaeniceps rex*) quadrate body, close to the base of the orbital process. It is surrounded by the medial crest and the orbital process, as shown in a relatively large size.

**Pelecaniformes:** Pelecanidae (Plate 32)

In Pelecanidae (*Pelecanus occidentalis*), the oitc process has a shallow intercapitular incisure. Unlike most avian quadrates, the otic capitulum is much elevated and larger than the squamosal capitulum. The squamosal capitulum shows a rostrocaudally elongate oval shape and it faces laterally different with most of the avian quadrates which face dorsally or dorsolaterally. The otic capitulum exhibits two small articulations connecting with the cranium, and both appear a rounded shape, pointing dorsally and dorsomedially, respectively. Between these two rounded articulations, a narrow and distinct insecure is present.

The orbital process of the *Pelecanus* quadrate appears robust and thick with a triangle shape in lateral view. Its tip is flat and points rostrally. The basiorbital fossa of the Pelecanidae quadrate is deep and broad.

Unlike most Ardeae (Eurypygimorphae and Aequornithes), the quadratojugal cotyle of the Pelecanidae quadrate does not shows a distinct margin. Instead, it shows a rounded shape with a significantly flat articular surface, close to the lateral condyle with a small gap. The pterygoid condyle of the *Pelecanus* quadrate is adjacent to the medial condyle (Saiff, 1978) and shows a rectangle-with-rounded-corner shape with a flat articular surface.

The mandibular process of the Pelecanidae quadrate has three condyles, arranging in a L shape with a deep fossa at the centre position (Saiff, 1978). The medial condyle has a slender and rostrocaudally elongate rectangle-with-rounded-corner shape while the lateral condyle exhibits a rounded shape. The caudal condyle appears a tiny protrusion, facing caudoventrally.

In Pelecanidae, the quadrate body is slightly laterally curved due to a curved lateral and a straight medial crest. The caudomedial depression is absent on the Brown pelican (*Pelecanus occidentalis*) quadrate. Instead, a large pneumatic foramen (postcapitular foramen) is caudally located at the otic process (Saiff, 1978). Besides, the Pelecanidae quadrate is highly pneumatic (Saiff, 1978): one fossa is laterally located at the quadrate body, close to the orbital process.

**Opsithocomiformes:** Opsithocomidae (Plate 33)

The otic process of Opsithocomidae (*Opisthocomus hoazin*) quadrate shows a shallow and narrow intercapitular incisure between two capitula. The squamosal capitulum is as elevated and large as the otic capitulum. The squamosal capitulum has a rostrocaudally elongate oval outline, facing dorsolaterally. The otic capitulum has a lateromedially elongate oval shape with a flat articular surface, and it faces dorsally.

The shape of the orbital process of the hoatzin quadrate is similar as several subscales of the galliform quadrates (Numididae and Phasianidae) – it is elongate and slender with a high aspect ratio, and its tip points rostrodorsally. The significant difference between the galliform quadrate and the hoatzin quadrate is that the orbital process of the hoatzin quadrate is significantly dorsoventrally deeper than that of the galliform quadrate. The basiorbital fossa is wide and shallow on the Opsithocomidae quadrate.

The quadratojugal cotyle of the Opsithocomidae quadrate also looks similar as some galliform quadrates (e.g., Numididae and Phasianidae), showing a saddle-like shape (the rostral and caudal margin is underdeveloped) with a shallow articulation. Unlike that of the galliform quadrate, the quadratojugal cotyle of the *Opisthocomus* quadrate is laterally protruding, adjacent to the lateral condyle. The pterygoid condyle on Opsithocomidae quadrate is not rostrally standing, and it is above the medially condyle with a wide and shallow. It exhibits a round shape with a flat articular surface.

The mandibular process of the Opsithocomidae quadrate has three condyles, arranging in a L or a boomerang shape. The medial condyle has a rostrocaudally elongate oval outline while the lateral condyle shows a lateromedially elongate oval outline. The caudal condyle appears a short and thick bulge with a flat articular surface.

In Opsithocomidae, the quadrate body is relatively curved with a right-angle lateral crest and a minorly curved medial crest. In the caudal view, the caudomedial depression is shallow. The subcapitular tubercle is rostrolaterally located at the squamosal capitulum of the Opsithocomidae quadrate, and it exhibits a mound-like shape. Either a fossa or a pneumatic foramen does not present on the Opsithocomidae quadrate.

**Accipitriformes:** Cathartidae (plate 33)

In Cathartidae (*Cathartes burrovianus*), the otic process shows a shallow intercapitular incisure between squamosal and otic capitulum. The squamosal capitulum is slightly elevated than the otic capitulum, but it has an equal size as the otic capitulum. The squamosal capitulum has a lateromedially elongate oval shape, facing dorsolaterally. The otic capitulum also has a lateromedially elongate oval shape, but it has a flat articular surface and faces caudodorsally or dorsally. Below the otic capitulum, there is a clear notch on the quadrate body.

The orbital process of the Cathartidae quadrate is elongate and straight with a high aspect ratio. Its tip is flat, broad and vertical, pointing rostrally. The Cathartidae quadrate exhibits a deep and wide basiorbital fossa.

The quadratojugal cotyle of the Cathartidae quadrate shows a laterally standing cup shape with a thick margin. It has a deep fossa and is close to the lateral condyle. The ventral margin of the quadratojugal cotyle expands ventrolaterally, forming a flat articular surface for jugal bar. The connection with pterygoid on the *Cathartes* quadrate has two articulations: the pterygoid condyle is dorsally located at the medial condyle with a short but deep gap. It has a rounded shape with a flat articular surface, facing rostrally; the orbitopterygoid facet is medially located at the orbital process, close to the base of the orbital process, and it has a rounded shape with a flat articular surface.

The mandibular process on the Cathartidae quadrate has three condyles, arranging in a L or triangle shape with a distinct intercondylar sulcus between medial and lateral condyle (Saiff, 2006), and a deep fossa at the centre position. Different from other accipitriform quadrates, the medial condyle of the Cathartidae quadrate has two articulations with lower jawbone, medial and lateral part (lateral trochlea). The medial part has a rostrocaudally elongate oval outline with a prominent convex articular surface. The lateral part has a rounded or square shape with a flat articular surface. Both these parts have a distinct margin with the quadrate body. The lateral condyle has a rectangle-with-rounded-corner outline, while the caudal condyle appears an elongate and thick protrusion, sub-horizontally aligning with a curved articular surface.

The Cathartidae quadrate body is relatively straight with a right-angle lateral and a slightly straight medial crest. The caudomedial depression is shallow and wide, while the caudomedial fossa appears to be distinct and deep. The subcapitular tubercle is rostroventrally located at the squamosal capitulum on the Cathartidae quadrate, as shown in a linear ridge. On the Lesser yellow-headed vulture (*Cathartes burrovianus*) quadrate, only one pneumatic foramen (basiorbital foramen) present as shown in medial surface of the quadrate body. The caudomedial fossa abovementioned is possibly related to the air sac system.

**Accipitriformes:** Pandionidae (Plate 33)

In Pandionidae (*Pandion haliaetus*), the otic process has a shallow and wide intercapitular incisure between two capitula. The squamosal capitulum is significantly elevated and slightly larger than the otic capitulum. The squamosal capitulum has a rounded outline, facing dorsolaterally. The otic capitulum has a lateromedially elongate oval outline, facing dorsally.

The orbital process of the *Pandion* quadrate is elongate and straight with a high aspect ratio. Its tip is flat, broad and vertical, pointing rostrally. The basiorbital fossa of the Pandionidae quadrate is slightly deep and broad.

The quadratojugal cotyle of the osprey quadrate shows a cup shape with a thick margin and it has a deep articular surface, adjacent to the lateral condyle. The rostroventral margin of the quadratojugal cotyle slightly expands ventrally, forming a flat articular surface for jugal bone. The articulation with pterygoid on the Pandionidae quadrate has two connections: one (pterygoid condyle) is above the medial condyle with a wide and shallow gap. It has an oval shape with a flat articular surface; the other (orbitopterygoid facet) is medially located at the orbital process and is close to the base of the orbital process. It has an oval shape with a flat articular surface.

The mandibular process of the Pandionidae quadrate has three condyles, arranging in a L shape with a shallow fossa at the centre position. The medial condyle shows a rostrocaudally elongate oval shape. The lateral condyle is confluent with the caudal condyle, and it exhibits a slender rectangle-with-rounded-corner outline. The caudal condyle appears a short and thick protrusion with a flat articular surface.

In Pandionidae, the quadrate body is relatively straight with a minorly curved lateral and medial crest. The caudomedial depression is wide and shallow. Either pneumatic foramen or fossa which is related to air sac system is absent on the osprey (*Pandion haliaetus*) quadrate in this study due to the juvenile or sub-adult specimen (FMNH 437336). The adult specimen (FMNH 466261) clearly shows a basiorbital foramen, similar to other accipitriform quadrates.

**Accipitriformes:** Sagittariidae (Plate 33)

In Sagittariidae (*Sagittarius serpentarius*), the otic process has a shallow intercapitular incisure between squamosal and otic capitulum, different with the previous observation (Saiff, 2006). The squamosal capitulum is significantly elevated than the otic capitulum, but it is smaller than the otic capitulum. The squamosal capitulum shows a lateromedially elongate oval shape, facing dorsally. The otic capitulum shows a lateromedially elongate parallelogram outline with a flat articular surface, facing dorsally. Below the otic capitulum, there is a wide and deep notch on the quadrate body.

The orbital process of the *Sagittarius* quadrate is robust, and its tip is sharp, facing rostroventrally. The basiorbital fossa is shallow and narrow in Sagittariidae.

The quadratojugal cotyle of the Sagittariidae quadrate is laterally standing and exhibits a cup shape with a thick margin. It has a shallow articulation and is adjacent to the lateral condyle. The rostroventral margin of the quadratojugal cotyle slightly expands ventrally, forming a flat articular surface for jugal bar attaching. The articulation with pterygoid on the *Sagittarius* quadrate has two contacts: one (pterygoid condyle) is above the medial condyle with a distinct gap, and it has a slender oval shape with a flat articular surface. The other (orbitopterygoid facet) is medially located at the orbital process, close to the base of the orbital process, and it has an oval shape with a flat articular surface.

The mandibular process of the Sagittariidae quadrate has three condyles, arranging in a triangle shape with a clear groove (intercondylar sulcus) between medial and lateral condyle. The medial condyle shows a rostrocaudally elongate oval shape. The lateral condyle is confluent with the caudal condyle and has a rounded outline. The caudal condyle appears a tiny protrusion with a flat articular surface.

In Sagittariidae, the quadrate body is slender and curved with a curved lateral and medial crest. The caudomedial depression is located caudally at the otic process and it is wide and shallow. The caudomedial fossa, encompassed by the medial crest and a tympanic crest, is relatively narrow and shallow. Two pneumatic foramina present on the *Sagittarius* quadrate (Saiff, 2006): one (basiorbital foramen) is medially located at the quadrate body with two tiny openings; the other (postcapitular foramen) is caudally located at the otic process, close to the middle position between two capitula. The caudomedial fossa abovementioned might be also part of the air sac system in bird.

**Accipitriformes:** Accipitridae (Plate 33-34)

In Accipitridae, the otic process has a shallow intercapitular incisure between two capitula in *Elanus caeruleus*, *Buteo rufofuscus* and *Necrosyrtes monachus*, but it has a deep and wide intercapitular incisure in *Accipiter nisus* and *Circus macrourus*. In *Parabuteo unicinctus*, the intercapitular incisure is absent. The squamosal capitulum is significantly elevated than the otic capitulum, but it usually has a smaller size than the otic capitulum. It has an equal size with the otic capitulum on *Parabuteo* quadrate only.

The squamosal capitulum of the Accipitridae quadrate shows a huge range of the morphological diversity: it has a lateromedially elongate oval outline, facing dorsolaterally in *Elanus caeruleus*, *Parabuteo unicinctus*, and *Necrosyrtes monachus*. The squamosal capitulum of the Jackal buzzard (*Buteo rufofuscus*) quadrate shows a rostrocaudally elongated oval outline and faces dorsolaterally. In *Accipiter nisus* and *Circus macrourus*, their squamosal capitula have rounded shapes, facing dorsolaterally. The otic capitulum of the Accipitridae quadrate usually shows a rostrocaudally elongate oval shape, pointing dorsomedially or dorsally. However, the otic capitulum of the Harris’s hawk (*Parabuteo unicinctus*) quadrate shows a rounded shape, instead. Below the otic capitulum, there is a wide but shallow notch on the quadrate body.

Similar to their sister group (Sagittariidae), the orbital process of the Accipitridae quadrate is robust with a slightly high aspect ratio. Its tip is sharp and points rostrally or rostroventrally. The Accipitridae quadrate has a relative deep and wide basiorbital fossa.

The quadratojugal cotyle of the Accipitridae quadrate is laterally standing and exhibits a cup-like shape with a thick margin. It usually has a deep articulation and is adjacent to the lateral condyle except Black-winged kite (*Elanus caeruleus*). The quadratojugal cotyle of the Black-winged kite quadrate is shallow and its rostroventral margin slightly expands ventrally, forming a flat articular surface for the jugal bone. The pterygoid condyle of the Accipitridae quadrate is dorsally located at the medial condyle with a distinct gap, and it appears an oval outline with a slightly convex articular surface. The articulation with pterygoid of the *Necrosyrtes monachus* quadrate, however, has two surfaces: the pterygoid condyle looks similar as that of other Accipitridae, and the orbitopterygoid facet is medially located at the orbital process, close to the base of the orbital process, and shows a slender oval shape with a flat articular surface.

The mandibular process on the Accipitridae quadrate has three condyles, arranging in a L shape with a shallow fossa at the centre position. The medial condyle shows a rostrocaudally elongate oval shape with a convex articular surface. The lateral condyle is confluent with the caudal condyle, and it also has a lateromedially elongate oval shape with a flat articular surface. The caudal condyle is not well-developed in Accipitridae, and it only appears a tiny protrusion with a flat articular surface.

In Accipitridae, the quadrate body is usually curved and slender with a curved lateral and medial crest. The caudomedial depression is caudally located at the otic process, and it is wide and shallow encompassed by the medial and tympanic crest. The caudomedial fossa is generally shallow on most Accipitridae quadrate, but it is significantly deep on Harris’s hawk and Pallid harrier (*Circus macrourus*) quadrates. The subcapitular tubercle on the Accipitridae quadrate appears a tiny mound and is ventrolaterally located at the squamosal capitulum.

The pneumatic foramen of the Accipitridae quadrate also exhibits great diversity in its positions: the basiobrital foramen, medially located at the quadrate body and close to basiobrital fossa, presents on all Accipitridae quadrate with the various size of opening. On Hooded vulture (*Necrosyrtes monachus*) quadrate, three extra pneumatic foramina present on its quadrate body: two are caudally located at the otic process (postsquamosal capitulum foramen and postcapitular foramen), while the other (rostromedial foramen) is medially located at the quadrate body, above the basiorbital fossa. The caudomedial fossa abovementioned might be related to the air sac system on most Accipitridae quadrate, but not found on Jackal buzzard and Hooded vulture quadrate.

**Strigiformes:** Tytonidae (Plate 35)

In Tytonidae, the intercapitular incisure is absent on the otic process, but each capitulum is far from each other, especially in Barn owl (*Tyto alba*). The squamosal capitulum of the Tytonidae quadrate is much elevated and larger than the otic capitulum. The squamosal capitulum has a rostrocaudally elongate rectangle-with-rounded-corner outline in Bay owl (*Phodilus badius*), pointing dorsally. The squamosal capitulum of the Barn owl quadrate, however, shows a T shape and it is compsed by two articulations: one shows a rostrocaudally elongate rectangle-with-rounded-corner outline with a convex articular surface, facing dorsally; the other has a lens-like shape with a flat articular surface, facing rostrodorsally. Unique to other avian otic process, the otic capitulum of the Tytonidae quadrate significantly prolonged caudomedially, forming a “process” -like structure (Shufeldt, 1900). In both taxa, the otic capitulum has two articular surfaces connecting with the cranium: one shows dorsoventrally elongate oval outline, facing caudomedially; the other, laterally located at the abomined one, also shows an oval shape, but points rostromedially.

The orbital process of the Tytonidae quadrate is dorsoventrally slender with either a flat tip in Bay owl or a sharp tip in Barn owl. The basiorbital fossa is shallow and wide.

The quadratojugal cotyle of the Tytonidae quadrate is laterally standing and shows a cup-like outline with a thick margin, and it is adjacent to the lateral condyle. It has a deep fossa on Barn owl quadrate but exhibits a flat articular surface on Bay owl quadrate. The articulation with pterygoid of the Tytonidae quadrate has two connections: one (pterygoid condyle) is dorsally adjacent to the medial condyle without any distinct gaps and shows a rounded shape. The other (orbitopterygoid facet) is medially located at the orbital process, close to the basiorbital fossa, and it appears a slender oval outline with a flat articular surface.

The mandibular process on the Tytonidae quadrate has three condyles, arranging in a L or a boomerang shape with a shallow and wide intercondylar sulcus between medial and lateral condyle. The medial condyle shows a rostrocaudally elongate oval shape with a convex articular surface. The lateral condyle is confluent with the caudal condyle, forming a rectangle-with-rounded-corner outline. The caudal condyle faces caudally.

In Tytonidae, the quadrate body is medially curved due to its strongly curved medial crest. The caudomedial depression, caudally located at the otic process, is wide and deep. One pneumatic foramen (caudomedial foramen), encompassed by the medial and tympanic crest, is caudomedially located at the Tytonidae quadrate body with a large opening.

**Strigiformes:** Strigidae (Plate 35)

Similar to its sister group (Tytonidae), the intercapitular incisure is absent on the Strigidae quadrate, and each capitulum is also far from each other. The squamosal capitulum is significantly elevated than the otic capitulum on the Strigidae quadrate. The squamosal capitulum shows an equal (in *Strix aluco*) or slightly larger (in *Ninox novaeseelandiae*) size than the otic capitulum. The otic process of the Strigidae quadrate has a great morphological diversity: in Tawny owl (*Strix aluco*), the squamosal capitulum shows a rostrocaudally elongate lens-like outline, facing dorsally. Its oitc capitulum has two articulations with the cranium: one appears an oval shape with a flat articular surface, facing medially; the other has a slender lens-like shape with a flat articular surface, pointing rostrodorsally. In Morepork (*Ninox novaeseelandiae*), the squamosal capitulum shows a lateromedially elongate oval outline, facing caudodorsally. Its oitc capitulum exhibits a number eight outline with a convex articular surface, and it faces dorsomedially. Similar as Tytonidae quadrate, the otic capitulum of the Strigidae quadrate is significantly caudomedially prolonged (Norberg, 1977; Shufeldt, 1900).

Like Tytonidae, the orbital process of the Strigidae quadrate is dorsoventrally slender and elongate, but its tip appears to be pointed in Morepork or hook-like in Tawny owl. The basiorbital fossa is shallow and wide.

The quadratojugal cotyle of the Strigidae quadrate is also laterally standing and adjacent to the lateral condyle. It shows a cup-like outline with a thick margin and has a deep fossa on Morepork quadrate or a shallow articular surface on Tawny owl quadrate. The articulation with pterygoid on the Strigidae quadrate has two attachments: the pterygoid condyle is not rostrally standing and shows a slender oval outline. It is adjacent to the medial condyle in Morepork or separated from the medial condyle with a wide gap in Tawny owl. The orbitopterygoid facet is medially located at the orbital process, close to the basiorbital fossa, it exhibits a slender oval outline with a flat articular surface.

The mandibular process on the Strigidae quadrate has three condyles, arranging in a L or a boomerang shape with a shallow and wide intercondylar sulcus between medial and lateral condyle (Choudhary et al., 2021). It has a deep fossa at the centre of the mandibular process on Morepork quadrate.The medial condyle shows a rostrocaudally elongate oval shape with a convex articular surface. The lateral condyle is confluent with the caudal condyle on Tawny owl quadrate only, forming a parallelogram outline. In Morepork, the lateral condyle shows a lateromedially elongate oval shape with a slightly convex articular surface. The caudal condyle appears a tiny and curved protrusion.

Similar to Tytonidae, the quadrate body of the Strigidae quadrate is medially curved due to its strongly curved medial crest. The caudomedial depression, caudally located at the otic process, is wide and relatively deep. In Strigidae, one pneumatic foramen (caudomedial foramen), encompassed by the medial and tympanic crest, is caudomedially located at the quadrate body with a relatively large size.

**Coliiformes:** Coliidae (Plate 36)

The otic process of the Coliidae quadrate shows a wide intercapitular incisure between two capitula. The squamosal capitulum is significantly elevated than the otic capitulum in rostral view and has a larger size that the otic capitulum. Both two capitula have lateromedially elongated oval outline, facing dorsally (squamosal capitulum) and medially (otic capitulum), respectively.

The orbital process of the Coliidae quadrate is relatively short and has a rounded tip, pointing rostrodorsally. The basiorbital fossa of the Coliidae quadrate s shallow and narrow.

The quadratojugal cotyle of the Coliidae quadrate is laterally protruding and has an oval outline, adjacent to the lateral condyle. However, the margin of the quadratojugal cotyle is not well-developed on the Blue-naped mousebird (*Urocolius* *macrourus*) quadrate, and it appears a flat articular surface, unlike a deep or shallow articulation. The quadratojugal cotyle of the Speckled mousebird (*Colius striatus*) quadrate exhibits a deep articulation with a relatively small size. On the Coliidae quadrate, the pterygoid condyle is not rostrally standing and dorsally adjacent to the medial condyle without any distinct gap. It has a rounded shape with a convex articular surface.

The mandibular process of the Coliidae quadrate has three condyles, arranging in L shape in ventral view with a deep furrow between medial and lateral condyle. The medial condyle shows a rectangle-with-rounded-corner outline with a prominent convex articular surface. The lateral condyle is continuous with the caudal condyle, forming a rectangle-with-rounded-corner outline with a flat articular surface. The caudal condyle appears a thick and short protrusion.

The quadrate body of the Coliidae is medially curved due to its strongly curved medial crest. In the caudal view, the caudomedial depression is wide and deep. The subcapitular tubercle of the Coliidae quadrate shoes a standing but slender oval outline, rostrolaterally below the squamosal capitulum. On the Coliidae quadrate, the pneumatic foramen is mostly located at the caudal side of the otic process with numerous tiny openings (e.g., postsquamosal capitulum foramen, postotic capitulum foramen, and postcapitular foramen). The Blue-naped mousebird quadrate, one extra pneumatic foramen caudomedially located at the quadrate body, encompassed by the medial and tympanic crest, and it has significantly large size, facing medially.

**Leptosomiformes:** Leptosomidae (Plate 36)

The otic process of the Leptosomidae quadrate has a distinct intercapitular incisure between two capitula. The squamosal capitulum is slightly elevated than the otic capitulum, but it shows a similar size as the otic capitulum. The squamosal capitulum shows a diamond shape with a convex articular surface, facing caudodorsally. The otic capitulum exhibits a lateromedially elongated oval outline, facing dorsomedially.

The orbital process of the Cuckoo-roller (*Leptosomus discolor*) quadrate is elongated and robust with a flat tip, facing rostrally. The Leptosomidae quadrate has a shallow and wide basaiorbital fossa.

The quadratojugal cotyle of the Leptosomidae quadrate is laterally protruding, adjacent to the lateral condyle, and it has a cup shape with a thick margin and a deep fossa. The ventral margin of the quadratojugal cotyle expands ventrally, forming a flat articular surface for jugal bar attaching. On the Leptosomidae quadrate, the pterygoid condyle is not rostrally standing and dorsally located at the medial condyle with a short but distinct gap. It shows a rounded outline with a convex articular surface.

The mandibular process of the Leptosomidae quadrate has three condyles, arranging in L shape in ventral view with a shallow and wide fossa at the centre position. The medial condyle shows a rostrocaudally elongated oval outline with a deep articular surface. The lateral condyle is confluent with the caudal condyle, forming a cashew-nut outline with a flat articular surface. The caudal condyle appears a thick and caudally standing protrusion.

The Leptosomidae quadrate body exhibits a curved shape due to a strongly curved lateral and medial crest. In the caudal view, the caudomedial depression is wide and shallow. The subcapitular tubercle of the *Leptosomus discolor* quadrate is lateroventrally located at the squamosal capitulum and seems to merge with the squamosal capitulum. On the Leptosomidae quadrate, two pneumatic foramens present at the medial side of the quadrate body: the rostromedial foramen, encompassed by the medial crest and the orbital process, shows a large size. The other, basiorbital foramen, is located close to the pterygoid condyle with a small opening.

**Trogoniformes:** Trogonidae (Plate 36)

On the Trogonidae quadrate, the otic process does not show a clear intercapitular incisure between two capitula. The two capitula are remarkably separated, and the squamosal capitulum is larger in size than the otic capitulum. Distinctly, in Trogonidae the squamosal capitulum shows a dolosse outline, having three “legs” extending in rostral, lateral and medial direction. The rostral and lateral “legs” form a concave articular surface, while the lateral and medial “legs” form a rectangle-with-rounded-corners outline with a flat articular surface. The otic capitulum shows two articulations and bears a distinct cleavage between these two articulations. Both two articulations are lateromedially elongate oval outline with a flat articular surface and are oriented dorsally or dorsomedially.

The orbital process of the Trogonidae quadrate is elongate with a flat and slant tip, pointing rostrodorsally. The tip of the orbital process tapers vertically or subvertically in *Trogon*, while it tapers horizontally in *Apaloderma narina*. The Trogonidae quadrate has a shallow and narrow basiorbital fossa.

Unlike other avian quadrate, the quadratojugal cotyle of the Trogonidae quadrate is laterally standing and shows a rectangle-with-rounded-corners outline with a flat articular surface, adjacent to the lateral condyle. This articular surface is oriented dorsolaterally, instead of facing laterally. The pterygoid condyle is rostrally protruding and dorsally separated from the medial condyle with a deep gap. This condyle bears a rounded outline and a significantly convex articular surface.

The mandibular process exhibits three condyles, arranging in L-shape in ventral view, and bears a shallow fossa at the centre position. The medial condyle shows a rounded outline with a prominent articular surface. The lateral condyle is continuous with the caudal condyle, forming a rectangle-with-rounded-corner outline with a slightly flat articular surface. The caudal condyle appears to be a thick and short protrusion.

The quadrate body of Trogonidae is slightly straight and bears a right-angle lateral crest and a slightly curved medial crest. In the caudal view, neither the caudomedial depression nor caudal fossa shows on the quadrate body. In Trogonidae, one single pneumatic foramen (postcapitular foramen) is caudally located at the otic process with a relatively large size, facing caudomedially.

**Bucerotiformes:** Upupidae (Plate 37)

The quadrate of Upupidae (*Upupa epops*) shows a packed otic process without a clear intercapitular incisure. The squamosal capitulum is slightly higher than the otic capitulum and shows a larger size than the otic capitulum. The squamosal capitulum has two distinct articular surfaces connecting with the cranium, and these two articulations form a rectangle-with-rounded-corners outline with a concave articular surface, facing caudodorsally. The otic capitulum appears a lateromedially elongated outline with a convex articular, pointing dorsomedially.

The orbital process of the Hoopoe (*Upupa epops*) quadrate is slender and elongated with a high aspect ratio. On the lateral surface of the orbital process, a distinct crest (not the orbital crest) extends from the tip of the orbital process to its base. The tip of the orbital process on *Upupa* quadrate shows flat, facing rostrodorsally. The Upupidae quadrate has a shallow and narrow basiorbital fossa.

The quadratojugal cotyle is laterally standing and exhibits a cup-like shape with a thick margin and a shallow fossa. The ventral margin expands rostroventrally, forming a flat articular surface for jugal bone. Also, the quadratojugal cotyle is dorsally adjacent to the lateral condyle. The pterygoid condyle is rostrally protruding and is dorsally adjacent to the medial condyle. The pterygoid condyle shows a circular outline with a convex articular surface.

On the Upupidae quadrate, the mandibular process has three condyles, arranging in L or T shape in ventral view with a shallow fossa at centre position. The medial condyle shows a parallelogram outline and rostrocaudally elongated with a prominent convex articular surface. The lateral condyle is continuous with the caudal condyle, forming an elongate oval outline with a flat articular surface. The caudal condyle appears to be an elongated protrusion at the caudal side of the quadrate body.

The Upupidae quadrate body is medially curved due to its strongly curved medial crest. The caudomedial depression is narrow but deep between two capitula. On the Upupidae quadrate, three positions exhibit pneumatic foramen: the caudomedial foramen is medially located at the quadrate body, encompassed by the medial and tympanic crest, and it has significantly large size, facing medially. The basiorbital foramen is also medially located at the quadrate body, but it is dorsally close to the pterygoid condyle with a tiny opening. The postcapitular foramen is caudally located between two capitula with several tiny pneumatic openings on the caudomedial depression.

**Bucerotiformes:** Bucorvidae (Plate 37)

The quadrate of Abyssinian ground hornbill (*Bucorvus abyssinicus*) shows a distinct but narrow intercapitular incisure on the otic process. The squamosal capitulum is slightly elevated and has significantly larger size than the otic capitulum. The squamosal capitulum of the Bucorvidae quadrate is similar as that of the Trogonidae quadrate, showing a dolosse outline with three “legs” extending medially, laterally, and caudally, respectively. The medially and lateral “legs” form a concave articular surface, and the medial leg appear a distinct ridge at the centre of the otic process. The caudal “leg” has a rounded outline. The otic capitulum exhibits a latermedially elongated lens-like outline with a convex articular surface, facing dorsally or dorsocaudally.

The orbital process of the Bucorvidae quadrate is significantly elongate but rostrodorsally deep. It has a flat tip, pointing rostrally. The basiorbital fossa on the Bucorvidae quadrate is shallow and narrow. The quadratojugal cotyle of the Bucorvidae quadrate shows a cup-like shape with a thick margin and a deep fossa, and it is dorsally adjacent to the lateral condyle. The ventral margin expanded ventrally, forming a flat aricular surface for jugal bone. The pterygoid condyle is not rostrally standing and is dorsally located at the medial condyle with a deep gap. It has a rectangle outline with a flat articular surface.

The mandibular process on the Bucorvidae quadrate has three condyles, arranging in a L or a hooked shape with a shallow fossa at the centre. The medial condyle has two articulations with the lower jawbone, separating into medial and later part (lateral trochlea). The medial part shows a rostrocaudally elongate oval outline with a convex articular surface, while the lateral trochlea appears an oval outline with a flat articular surface. The lateral condyle exhibits a lateromedially elongate oval outline with a flat articular surface. The caudal condyle appears a thick protrusion from the caudal side of the quadrate body.

On the Bucorvidae quadrate, its quadrate body is slightly straight due to its minorly curved lateral and medial crest. In the caudal view, the caudomedial depression or the caudomedial fossa is absent on the quadrate body. Only one single pneumatic foramen (rostromedial foramen) shows on the medial side of the quadrate body, encompassed by the medial crest and the orbital process.

**Bucerotiformes:** Phoeniculidae (Plate 37)

Generally speaking, the Phoeniculidae quadrate is morphologically similar to the Upupidae quadrate. For instance, the green wood hoopoe (*Phoeniculus purpureus*) quadrate does not show a distinct intercapitular incisure on the otic process. The squamosal capitulum is slightly elevated and larger than the otic capitulum. Different from that of the Upupidae quadrate, the squamosal capitulum on the Phoeniculidae quadrate shows a rostrocaudally elongated rectangle outline, while the otic cpaiutlum shows a lateromedially elongated rectangle outline with a flat articular surface.

The orbital process of the green wood hoopoe quadrate is relatively elongate and slender, and has a slant tip with a flat head, pointing rostrally. Its tip twist horizontally, unlike that of the hoopoe which shows a closely vertical. The basiorbital fossa of the Phoeniculidae quadrate is shallow and narrow. On the Phoeniculidae quadrate, the quadratojugal cotyle has a cup-like shape with a thick margin and a deep fossa, and it is dorsally adjacent to the lateral condyle. The ventral margin ventrally expanded, forming a flat articular surface for jugal bar contacting. The pterygoid condyle is dorsally adjacent to the medial condyle, and it has a rounded shape with a convex articular surface.

The mandibular process on the Phoeniculidae quadrate has three condyles, arranging in a T shape with a shallow fossa at the centre position. Unlike that of the hoopoe quadrate, the medial condyle of the green wood hoopoe quadrate show two articulations with the lower jawbone, separating into medial and later part (lateral trochlea). The medial part exhibits a rostrocaudally elongated oval outline, while the lateral part exhibits an arch-like shape with a flat articular surface. The lateral condyle is continuous with the caudal condyle, forming a lens-like or oval outline with a flat articular surface. The caudal condyle appears to be a thick and standing protrusion, pointing medially.

On the Phoeniculidae quadrate, its body is medially curved due to its right-angle lateral crest and a curved medial crest. The caudomedial depression is narrow but relatively deep, and the caudomedial fossa is accommodated at the caudomedial depression. In Phoeniculidae, one pneumatic foramen (postcapitular foramen) is caudally located at the otic process with a large opening. The abovementioned caudomedial fossa is likely related to the air sac system in bird skull.

**Coraciformes:** Meropidae (Plate 37)

The quadrate of Meropidae (*Merops orientalis*) show a packed capitula without a clear intercapitular incisure on the otic process. The squamosal capitulum is as high and large as the otic capitulum. The squamosal capitulum shows a lateromedially elongated oval outline and face dorsally. The otic capitulum is composed of two small lateromedially elongated oval facets with flat articular surfaces, and two facets point dorsomedially. There is an unclear incisure between these two facets.

The orbital process of the Asian green bee-eater (*Merops orientalis*) quadrate is robust with a low aspect ratio and shows a triangle outline in lateral view. On the middle of its dorsal margin, a distinct protrusion forms a small crest. The tip of the orbital process is pointed, facing rostrally. The Meropidae quadrate exhibits a deep and narrow basiorbital fossa.

Similar to that of the Trogonidae quadrates, the quadratojugal cotyle of the Meropidae quadrate is laterally standing and shows a rectangle-with-rounded-corners outline, dorsally close to the lateral condyle. Unlike the quadratojugal cotyle of the Trogonidae quadrates or some nightjar, it shows a shallow fossa and faces laterally on the Meropidae quadrate. The pterygoid condyle is dorsally located at the medial condyle with a wide and shallow gap. It has an oval outline with a convex articular surface.

The mandibular process on the Meropidae quadrate has three condyles, arranging in a L shape with a deep fossa (intercondylar sulcus) at the centre position. The medial condyle shows a rostrodorsally elongated oval shape with a significantly convex articular surface. The lateral condyle merges with the caudal condyle, forming a lateromedially elongated oval outline with a flat articular surface.

The quadrate body of the Asian green bee-eater is medially curved due to its right-angle lateral crest and a curved medial crest. In the caudal view, the caudomedial depression is wide and shallow. The subcaptiular tubercle is located below the squamosal capitulum with a tiny oval shape on the Meropidae quadrate. In Meropidae, two pneumatic foramens medially present on the quadrate body: the rostromedial foramen is encompassed by the medial crest and the orbital process, showing an elongate and slender opening; the basiorbital foramen is close the pterygoid condyle with an elongate opening.

**Coraciformes:** Coraciidae (Plate 37)

The Coraciidae quadrate shows a clear intercapitular incisure between two capitula on the otic process. The squamosal capitulum is elevated and larger than the otic capitulum. The squamosal capitulum shows a lateromedially slender oval shape with a prominent convex articular surface facing dorsally or dorsocaudally. The otic capitulum shows a rectangle outline with a flat articular surface, and it points dorsally or dorsomedially.

The orbital process of the Indian roller (*Coracias benghalensis*) quadrate is elongate with a triangle outline in lateral view. A distinct linear ridge extends from the tip of the orbital process. The tip of the orbital process is pointed, facing rostrally. Besides, the Coraciidae quadrate has a shallow and wide basiorbital fossa.

The quadratojugal cotyle of the Coraciidae quadrate shows a cup-like shape with a thick margin and a deep fossa, and it is adjacent to the lateral condyle. The ventral margin of the quadratojugal cotyle expanded ventrally, forming a flat articular surface for jugal bar. The pterygoid condyle is rostrally standing and dorsally located at the medial condyle with a wide gap. It has a rounded shape with a significantly articular surface.

The mandibular process on the Coraciidae quadrate has three condyles, arranging in a L shape with a deep fossa (intercondylar sulcus) at the centre position. The medial condyle has a lens-like outline and rostrocaudally elongate with a significantly convex articular surface. The lateral condyle shows a rectangle outline with a flat articular surface, while the caudal condyle appears to be a thick bulge and horizontally aligned.

The Coraciidae quadrate body is medially curved due to its right-angle lateral crest and a curved medial crest. In the caudal view, the caudomedial depression is wide and shallow. In Coraciidae, only one pneumatic foramen medially appears on the quadrate body: the basiorbital foramen is located at the basiorbital fossa with a large opening.

**Coraciformes:** Brachypteraciidae (Plate 37)

On the Brachypteraciidae quadrate, the otic process exhibits a packed capitula without a clear intercapitular incisure between them. The squamosal capitulum is as elevated and large as the otic capitulum. The squamosal capitulum shows a lateromedially elongate rectangle shape, facing laterodorsally. The otic capitulum shows a rounded or diamond-like shape and it faces dorsomedially.

The orbital process of the Pitta-like ground roller (*Atelornis pittoides*) quadrate is elongate with a triangle outline in lateral view. A distinct mound is dorsally located at the midpoint of the orbital process. The tip of the orbital process appears pointed and blunt on the Brachypteraciidae quadrate and faces rostrally. The basiorbital fossa of the Brachypteraciidae quadrate is shallow and wide.

The quadratojugal cotyle of the Brachypteraciidae quadrate has a cup-like shape with a thick margin and a deep articular surface, and it is dorsally located at the lateral condyle. The rostroventral margins expand rostroventrally, forming a flat articular surface for jugal bar contacting. The pterygoid condyle is rostrally standing and is dorsally separated from the medial condyle with a wide gap. It has a rounded shape with a significantly convex articular surface and small size.

The mandibular process on the Brachypteraciidae quadrate has three condyles, arranging in a L shape with a shallow fossa/furrow (intercondylar sulcus) between medial and lateral condyle. The medial condyle shows a rostrocuadally elongated lens-like shape with a prominent convex articular surface. The lateral condyle is confluent with the caudal condyle, forming a rectangle outline with a flat articular surface. The caudal condyle appears a thick bulge.

The Brachypteraciidae quadrate body is relatively straight due to its right-angle lateral crest and a minorly curved medial crest. In the caudal view, the caudomedial depression is narrow and shallow. The subcapitular tubercle on the Brachypteraciidae quadrate is ventrolaterally located at the squamosal capitulum and it appears a tiny and slender oval shape. In Brachypteraciidae, only one pneumatic foramen (postcapitular foramen) is caudally located at the otic process.

**Coraciformes:** Todidae (Plate 38)

The Todidae quadrate shows a packed capitula on the otic process with a clear intercapitular incisure between these two capitula. The squamosal capitulum is as elevated and large as the otic capitulum. The squamosal capitulum shows a rounded shape with a convex articular surface and points rostrodorsally. The otic capitulum has a rostrocaudally elongate rectangle outline with a flat articular surface, facing dorsomedially.

The orbital process of the Puerto Rican tody (*Todus mexicanus*) is relatively short and a triangle outline in lateral view. Similar as that of the Brachypteraciidae quadrate, a distinct rounded ridge is dorsally located at the orbital process. The Todidae quadrate has a deep and wide basiorbital fossa.

The quadratojugal cotyle on the Todidae quadrate shows a cup-like shape (though its rostral and caudal margin is not well-developed) with a thick margin and a deep fossa. It is dorsally close to the lateral condyle. The ventral margin expands rostroventrally, forming a flat articular surface for jugal bar attaching. The pterygoid condyle is rostrally standing and dorsally separated from the medial condyle with a wide and shallow gap. It exhibits a rounded shape with a convex articular surface.

The mandibular process on the Todidae quadrate has three condyles, arranging in a boomerang shape with a deep furrow (intercondylar sulcus) between medial and lateral condyle. The medial condyle shows a rostrocaudally elongate lens-like outline with a prominent convex articular surface. The lateral condyle exhibits a lateromedially elongate oval shape with a flat articular surface. The caudal condyle appears a thick protrusion.

The Todidae quadrate body is medially curved due to its right-angle lateral crest and a minor curved medial crest. The caudomedial depression is narrow and shallow on the Todidae quadrate. In Todidae, one pneumatic foramen (postcapitular foramen) is caudally located at the otic process, as shown in a tiny opening.

**Coraciformes:** Momotidae (Plate 38)

On the Momotidae quadrate, the otic process has a distinct but shallow intercapitular incisure between two capitula. The squamosal capitulum is elevated than the otic capitulum, but it has a smaller size than the otic capitulum. The squamosal capitulum exhibits a rostrocaudally elongate rectangle outline with a convex articular surface, facing dorsally. The otic capitulum shows a cashew-like shape with a convex articular surface, and it faces dorsomedially.

The orbital process of the Amazonian motmot (*Momotus momota*) is significantly elongate and slender with a high aspect ratio. Also, a distinct ridge is dorsolaterally located at the orbital process, close to the tip. The tip of the orbital process appears a slant tip (sub-horizontal) with a rounded head, facing rostrally. The Momotidae quadrate has a slightly deep but narrow basiorbital fossa.

The quadratojugal cotyle of the Momotidae quadrate has a cup-like shape with a thick margin and a deep fossa, and it is adjacent to the lateral condyle. The ventral margin expands rostroventrally, forming a flat articular surface for jugal bar. The pterygoid condyle is slightly standing and separated from the medial condyle with a wide gap. It shows a slender oval shape with a convex articular surface.

The mandibular process on the Momotidae quadrate has three condyles, arranging in a L shape with a shallow fossa (intercondylar sulcus) at the centre position. The medial condyle shows a rostrocaudally elongate lens-like outline with a prominent convex articular surface. The lateral condyle exhibits a cashew shape with a flat articular surface, while the caudal condyle appears a significantly thick bulge.

The Momotidae quadrate body is medially curved due to its right-angle lateral crest and a curved medial crest. The caudomedial depression is narrow and deep on the Amazonian motmot quadrate. In Momotidae, one pneumatic foramen is caudally located at the otic process as shown in a relatively large opening.

**Coraciformes:** Alcedinidae (Plate 38)

The Alcedinidae quadrate shows a packed capitula on the otic process with a narrow and shallow intercapitular incisure between two capitula. The squamosal capitulum is as elevated as the otic capitulum and it is significantly larger than the otic capitulum. Similar as that of the Bucorvidae and Trogonidae quadrate, the squamosal capitulum of the Alcedinidae quadrate exhibits a dolosse outline with three “legs” extending medially, laterally, and caudally, respectively. The medial and lateral “legs” form a concave articular surface, and the caudal “leg” appears a square or rounded outline with a flat articular surface. The otic capitulum shows a rounded or square shape with a flat articular surface, and it faces dorsomedially.

The orbital process of the kingfisher (*Alcedo atthis*) quadrate is significantly short with a low aspect ratio. The orbital process shows an obtuse triangle outline in lateral view, and its tip is blunt and rounded, pointing rostrally. The basiorbital fossa is shallow and narrow on the Alcedinidae quadrate.

The quadratojugal cotyle of the Alcedinidae quadrate has a cup-like shape, but its dorsorostral margin is slightly inward. The quadratojugal cotyle shows a deep fossa, and it is adjacent to the lateral condyle. The pterygoid condyle is rostrally standing and dorsally adjacent to the medial condyle. It has a rounded shape with a significantly convex articular surface.

The mandibular process on the Alcedinidae quadrate has three condyles, arranging in a L shape with a shallow furrow (intercondylar sulcus) between medial condyle and lateral-caudal condyle. The medial condyle has two articulation surfaces with the lower jawbone, separating into medial and lateral part (lateral trochlea). The medial part shows a rostrocaudally elongate oval shape with a prominent convex articular surface. The lateral part has a rounded outline with a flat articular surface. The lateral condyle merges with the caudal condyle, forming a rectangular outline with a flat articular surface. The caudal condyle appears a thick bulge.

The quadrate body of Alcedinidae is relatively straight due to its right-angle lateral and medial crest. The caudomedial depression is narrow but relatively deeper on the Alcedinidae quadrate. The subcapitular tubercle of the Alcedinidae quadrate is laterally ventrolaterally located at the squamosal capitulum with a slender oval shape. In Alcedinidae, two pneumatic foramens (postotic capitulum foramen) are caudomedially located at the otic capitulum.

**Piciformes:** Galbulidae (Plate 38)

On the otic process of the Galbulidae quadrate, the intercapitular incisure is shallow and narrow between two capitula. The squamosal capitulum is as high and large as the otic capitulum. The squamosal capitulum shows a lateromedially wide oval shape with a standing convex articular surface, facing dorsolaterally. The otic capitulum has a number-eight outline with a flat articular surface and faces dorsomedially.

The orbital process of the Paradise jacamar (*Galbula dea*) quadrate is robust and thick, and its tip is blunt and obtuse, pointing rostrally. The Galbulidae quadrate shows a deep and narrow basiorbital fossa.

The quadratojugal cotyle of the Galbulidae quadrate has a cup-like shape with an oval outline and a deep fossa, and it is dorsally close to the lateral condyle with a small gap. Its rostroventral margin expands rostroventrally, significantly forming a flat articular surface for jugal bar attaching. The pterygoid condyle is rostrally standing, and it is above the medial condyle with a wide gap. It exhibits a rounded shape with a convex articular surface.

The mandibular process on the Galbulidae quadrate has three condyles, arranging in L shape with a shallow furrow (intercondylar sulcus) between medial condyle and lateral condyle. The medial condyle has a rostrocaudally elongate oval shape with a prominently convex articular surface. The lateral condyle shows a rectangle outline with a minorly convex articular surface. The caudal condyle appears a thick bulge with a flat articular surface.

The Galbulidae quadrate body slightly is curved due to its right-angle lateral and minorly curved medial crest. In the caudal view, the caudomedial depression is wide but relatively deep. On the Galbulidae quadrate, four small pneumatic foramens present: two (postcapitular foramen and postotic capitulum foramen) are caudally located at the otic process. The other two, rostromedial foramen and basiobrital foramen, are medially located at the quadrate body.

**Piciformes:** Bucconidae (Plate 38)

On the Bucconidae quadrate, the intercapitular incisure is absent on the otic process. The squamosal capitulum is significantly higher and larger than the otic capitulum. The squamosal capitulum shows a rounded or square shape with a prominently convex articular surface, pointing dorsally or dorsolaterally. The otic capitulum shows a lateromedially (or dorsoventrally) elongate oval shape with a flat articular surface, facing medially.

The orbital process of the Bucconidae quadrate is elongate with a triangular outline in lateral view. A small, rounded crest is dorsally located at the orbital process, close to the midpoint of the orbital process. The tip of the orbital process exhibits a hook shape, pointing rostrally. The Bucconidae quadrate has a shallow and narrow basiorbital fossa.

The quadratojugal cotyle of the Bucconidae quadrate exhibits a cup shape with a deep articular fossa, and it is adjacent to the lateral condyle. Its ventral margin expands outward, forming a thick and flat articular surface for jugal. The pterygoid condyle slightly protrudes rostrally, and it is dorsally separated from the medial condyle with a wide gap. It has a rounded shape with a convex articular surface.

The mandibular process on the Bucconidae quadrate has three condyles, arranging in L shape with a shallow fossa (intercondylar sulcus) at the centre position. The medial condyle is much deeper than that of most avian quadrates, and it shows a rostrocaudally elongate oval shape with a prominently convex articular surface. The lateral condyle is continuous with the caudal condyle, forming a rectangle outline with a flat articular surface. The caudal condyle appears a thick and short bulge in caudal view.

The Bucconidae quadrate body is medially curved due to the significantly curved medial crest. In caudal view, the caudomedial depression is narrow and shallow. In Bucconidae, one pneumatic foramen and one fossa is present on the quadrate. The rostromedial pneumatic foramen is medially located at the quadrate body with a tiny opening, while the caudomedial fossa is located at the caudomedial depression, encompassed by the medial crest and the tympanic crest. This fossa possibly links to the avian cranial air sac system.

**Piciformes:** Indicatoridae (Plate 38)

The Indicatoridae quadrate shows a shallow and wide intercapitular incisure between two capitula on the otic process. The squamosal capitulum is significantly higher than the otic capitulum but it shows a smaller size than the otic capitulum. The squamosal capitulum exhibits a lateromedially elongate rectangular outline with a convex articular surface, and it faces dorsolaterally. The otic capitulum shows a lateromedially elongate oval shape with a flat articular surface, facing dorsomedially.

The orbital process of the Least honeyguide (*Indicator exilis*) quadrate is slender and elongate with a high aspect ratio. Its tip appears to be blunt and rounded, pointing rostrodorsally. The Indicatoridae quadrate shows a shallow but wide basiorbital fossa.

The quadratojugal cotyle of the Indicatoridae quadrate shows a saddle-like shape with a shallow articulation, adjacent to the lateral condyle. Unlike that of some Galliformes (especially for Phasianidae), the rostrodorsal and caudoventral margin of the quadratojugal cotyle is not well-developed on the Indicatoridae quadrate, and its fossa is much shallow than that of galliform quadrates. The pterygoid condyle is separated from the medial condyle with a small gap, and it exhibits an oval shape with a convex articular surface.

The mandibular process on the Indicatoridae quadrate has three condyles, arranging in L shape with a shallow fossa (intercondylar sulcus) at the centre. The medial condyle shows a rostrocaudally elongate oval shape with a prominently convex articular surface. The later condyle elongates lateromedially, exhibiting an oval shape with a flat articular surface. The caudal condyle appears a thick, short, and pointed bulge with a flat articular surface.

The Indicatoridae quadrate body is medially curved due to its curved medial crest. In caudal view, the caudomedial depression is wide and deep. In Indicatoridae, three pneumatic foramens are caudally located at the otic process, with a large opening of the postcapitular foramen and two tiny openings below each capitulum.

**Piciformes:** Picidae (Plate 39)

On the Picidae quadrate, the intercapitular incisure is absent between two capitula on the otic process. The squamosal capitulum is as elevated as the otic capitulum, but it is smaller than the otic capitulum. The shape of the two capitula is slightly different among Picidae: in Eurasian wryneck (*Jynx torquilla*), its squamosal capitulum has a rounded shape with a convex articular surface and points dorsolaterally, while its otic capitulum has a rectangular outline with a flat articular surface and faces dorsally. On the other hand, the squamosal capitulum on the European green woodpecker (*Picus viridis*) quadrate shows a dorsoventrally elongate oval outline with a convex articular surface, pointing laterally or dorsolaterally. Its otic capitulum shows a rostrocaudally elongate oval outline with a flat articular surface, facing dorsally or caudodorsally.

The orbital process of the Picidae quadrate is also morphologically different between *Jynx torquilla* and *Picus viridis*: On Eurasian wryneck quadrate, its orbital process is slender and dorsally curved, and the tip of the orbital process is blunt, pointing rostrodorsally. The orbital process of the European green woodpecker quadrate is significantly elongate with a higher aspect ratio and its tip is blunt and flat, pointing rostrally. The basiorbital fossa of the Picidae quadrate appears to be shallow and slightly wide.

The quadratojugal cotyle of Eurasian wryneck quadrate shows a saddle-like outline with a distinct dorsal margin and a shallow fossa, while the European green woodpecker quadrate shows a cup-like shape quadratojugal cotyle with a thin margin (though its ventral margin slightly shrinks inwardly) and a deep articulation. A robust protrusion caudomedially expands from the dorsal margin of the quadratojugal cotyle on European green woodpecker quadrate only. In both species, the quadratojugal cotyle is dorsally adjacent to the lateral condyle. The pterygoid condyle is clearly separated from the medial condyle with a small gap on the Picidae quadrate, and it shows an oval shape with a flat articular surface.

On the Picidae quadrate, the mandibular process has three condyles, arranging in a L shape with a shallow fossa (intercondylar sulcus) at the centre of the Eurasian wryneck quadrate, or arranging in a triangular shape with a distinct furrow (intercondylar sulcus) between medial condyle and lateral-caudal condyle on the European green woodpecker quadrate. The medial condyle shows a rostrocaudally elongate oval shape in *Jynx torquilla* but it shows a lateromedially elongate oval shape in *Picus viridis*. On the Eurasian wryneck quadrate, the lateral condyle exhibits a rounded/square shape with a flat articular surface, and the caudal condyle appears a thick and rounded bulge with a flat articular surface. However, the lateral condyle merges with the caudal condyle on the European green woodpecker quadrate, forming a slender and elegante oval outline with a flat articular surface.

The Picidae quadrate body is relatively straight due to its right-angle lateral and straight medial crest, and the European green woodpecker quadrate is significantly wider than the Eurasian wryneck quadrate body. The caudomedial depression is wide and shallow on the *Jynx torquilla* quadrate, but it is narrow and shallow on the *Picus viridis* quadrate. The subcapitular tubercle only appears on the European green woodpecker quadrate, as shown in a standing and slender muscle attachment. It is ventrolaterally located at the squamosal capitulum. On the Picidae quadrate, the pneumatic foramen only shows on the medial side of the quadrate body in *Picus viridis*, encompassed by the medial crest and a ridge.

**Piciformes:** Megalaimidae (Plate 39)

On the otic process of the Megalaimidae quadrate, the intercapitular incisure is absent between two capitula, but each capitulum are separated from each other. The squamosal capitulum is much elevated and larger than the otic capitulum. The squamosal capitulum shows a rostrocaudally elongate oval outline and faces dorsolaterally. The otic capitulum elongates lateromedially, as shown in an oval shape with a convex articular surface facing dorsally.

The orbital process of the Golden-whiskered barbet (*Psilopogon chrysopogon*) quadrate is slender and elongate with a high aspect ratio. A distinct and flat crest is located at the dorsal margin of the orbital process, close to its tip, and it slightly extends laterally. The tip of the orbital process is rounded and blunt, pointing rostrodorsally. The Megalaimidae quadrate has a shallow and narrow basiorbital fossa.

The quadratojugal cotyle of the Megalaimidae quadrate shows a saddle-like shape with a shallow fossa, adjacent to the lateral condyle. Similar as that of the Indicatoridae quadrate, the rostrodorsal margin of the quadratojugal cotyle is not well-developed on the Golden-whiskered barbet quadrate. Its pterygoid condyle is dorsally adjacent to the medial condyle and has a rounded shape with a convex articular surface.

The mandibular process on the Megalaimidae quadrate has three condyles, arranging in L shape with a shallow fossa (intercondylar sulcus) at the centre position. The medial condyle shows a rostrocaudally elongate rectangular outline with a prominently convex articular surface. The lateral condyle exhibits a rounded shape with a flat articular surface, while the caudal condyle appears to be a thick bulge with a triangular shape and a flat articular surface.

The Megalaimidae quadrate body is medially curved due the curved medial crest. In caudal view, the caudomedial depression is narrow and shallow. The subcapitular tubercle is rostrolaterally located at the squamosal capitulum, as shown in a triangular shape. In Megalaimidae, one pneumatic foramen, postcapitular foramen, presents on the caudomedial depression.

**Piciformes:** Lybiidae (Plate 39)

The quadrate of Lybiidae shows a narrow and shallow intercapitular incisure on the otic process. The squamosal capitulum is significantly higher than the otic capitulum, but it is smaller than the otic capitulum. The squamosal capitulum exhibits a dorsoventrally (or lateromedially) oval outline with a significantly convex articular surface, facing rostrolaterally. The otic capitulum exhibits a triangular outline with a flat articular surface, and it points dorsally.

The orbital process of the Lybiidae quadrate is slender and elongate with a high aspect ratio. Its tip expands outwardly, forming a fan-like shape pointing rostrally. The basiorbital fossa is shallow and wide on the Lybiidae quadrate.

On the Lybiidae quadrate, the quadratojugal cotyle has an oval shape with a shallow fossa, dorsally close to the lateral condyle with a small gap. The dorsal margin of the quadratojugal cotyle expands rostroventrally, forming a protrusion for jugal bone articulation. The pterygoid condyle on the Bearded barbet (*Lybius dubius*) quadrate is dorsally separated with the medial condyle with a small gap, and it exhibits a rounded or squared shape with a convex articular surface.

The mandibular process on the Lybiidae quadrate has three condyles, arranging in L shape in ventral view with a shallow furrow (intercondylar sulcus) at the centre position. The medial condyle has a trapezoid shape with a convex articular surface. The lateral condyle appears an oval shape with a flat articular surface. The caudal condyle shows a thick bulge with a rectangle shape in caudal view, facing caudally or caudoventrally.

The Lybiidae quadrate body is slightly straight due to its minorly curved lateral and medial crest. In caudal view, the caudomedial depression is narrow and shallow. In Lybiidae, one pneumatic foramen, postcapitular foramen, shows on the caudomedial depression with a relatively large size.

**Piciformes:** Ramphastidae (Plate 39)

The Ramphastidae quadrate shows a narrow and shallow intercapitular incisure between two capitula. Similar as that of the Lybiidae quadrate, the squamosal capitulum is much elevated than the otic capitulum on Yellow-throated toucan (*Ramphastos ambiguus*) quadrate. However, its squamosal capitulum has larger size than the otic capitulum. The squamosal capitulum exhibits a lateromedially wide oval outline with a convex articular surface, facing dorsolaterally. The otic capitulum has a rounded shape with a flat articular surface, and it points caudomedially.

Like that of most Piciformes, the orbital process of Yellow-throated toucan quadrate is slender and elongate with a significantly high aspect ratio. Its tip appears an arrow shape with a distinct ridge at the dorsal margin of the orbital process, and it points rostrally. The Ramphastidae quadrate has a shallow and slightly narrow basiorbital fossa.

The quadratojugal cotyle of the Ramphastidae quadrate has a cup-like shape with a thick margin and a deep fossa, adjacent to the lateral condyle. The ventral margin of the quadratojugal cotyle expands laterally on the Ramphastidae quadrate, forming a flat articular surface for jugal bone contacting. The pterygoid condyle is dorsally adjacent to the medial condyle with a distinct gap, and it appears an oval shape with a convex articular surface.

The mandibular process on the Ramphastidae quadrate has three condyles, arranging in L shape in ventral view. The medial condyle shows a rostrocaudally elongate oval shape with a convex articular surface. The lateral condyle is confluent with the caudal condyle, forming a rectangle outline with a flat articular surface. The caudal condyle appears a thick bulge in caudal view, and its articular surface faces caudally.

The Ramphastidae quadrate body is slightly straight and wide due to its minorly curved lateral and medial crest. In caudal view, the caudomedial depression is narrow and shallow. On the Ramphastidae quadrate, the subcapitular tubercle appears to be small mound-like structure, ventrolaterally located at the squamosal capitulum. In Ramphastidae, one tiny pneumatic foramen shows on the caudomedial depression, at midpoints between two capitula.

**Cariamiformes:** Cariamidae (Plate 40)

On the Cariamidae quadrate, the intercapitular incisure is absent between two capitula on the otic process. These two capitula are widely separated on the Black-legged seriema (*Chunga burmeisteri*) quadrate but are closely packed with each other on the Red-legged seriema (*Cariama cristata*) quadrate. In Cariamidae, the squamosal capitulum is slightly higher than the otic capitulum in rostral view but smaller than the otic capitulum. The squamosal capitulum shows a lateromedially elongate oval shape with a convex articular surface, facing dorsolaterally. On the other hand, the otic capitulum show different shape between two Cariamidae: it exhibits a rectangular outline with a flat articular surface on the Red-legged seriema quadrate, facing caudomedially. It exhibits a lateromedially elongate oval shape on the Black-legged seriema quadrate, pointing dorsomedially.

In Cariamidae, the orbital process is elongate with a flat tip, facing rostrally or rostrodorsally. In *Chunga burmeisteri*, a small mound is located at the midpoint of the dorsal margin of the orbital process. The basiorbital fossa of the Cariamidae quadrate appears to be deep and narrow, especially in *Chunga burmeisteri*. The quadratojugal cotyle of the Cariamidae quadrate is laterally standing and shows an oval shape with either a deep articular surface (*Chunga burmeisteri*) or a shallow fossa (*Cariama cristata*). The ventral margin of the quadratojugal cotyle expands laterally, forming a flat articular surface for jugal attaching. The pterygoid condyle is dorsally separated from the medial condyle with a wide and shallow gap. It exhibits an oval with a convex articular surface.

The mandibular process on the Cariamidae quadrate has three condyles, arranging in a L shape with a deep furrow (intercondylar sulcus) between medial and lateral-caudal condyle. The medial condyle shows a rostrocaudally elongate rectangular, or lens-like shape elongate with a convex articular surface. The lateral condyle is confluent with the caudal condyle, forming a rectangular or oval outline with a flat articular surface. The caudal condyle appears to be a pointed protrusion, displacing at the caudal side of the quadrate body.

The Cariamidae quadrate body is medially curved due to its right-angle lateral crest and a curved medial crest. In caudal view, the caudomedial depression between two capiutla is narrow and shallow. In Cariamidae, only one pneumatic foramen (postcapitular foramen) is caudally located at the caudomedial depression, close to the middle position between two capitula.

**Falconiformes:** Falconidae (Plate 40-41)

The falconiform quadrate mostly shows a widely separated intercapitular incisure on the otic process, but on American kestrel (*Falco sparverius*) quadrate, the intercapitular incisure is absent between two capitula. The squamosal capitulum is significantly higher than the otic capitulum on the falconiform quadrate, especially that of the American kestrel quadrate. Both two capitula on the otic process appear to be equal size to each other. The squamosal capitulum of the Falconiformes quadrate mostly shows a lateromedially elongate oval outline, facing dorsally. However, on the Barred Forest falcon (*Micrastur ruficollis*) quadrate, the squamosal capitulum splits into two articulations, rostral and caudal portion) with a deep incisure between these two facets. The rostral portion exhibits a rounded outline with a flat articular surface, while the caudal portion exhibits a rostrocaudally elongate rectangular outline with a convex articular surface. The otic capitulum shows a lateromedially elongate rectangular/oval outline with a flat articular surface, pointing dorsomedially.

The orbital process of the Falconidae quadrate is relatively short with a robust base and a low aspect ratio. The tip of the orbital process varies in shape, as seen in a rounded tip on Barred Forest falcon, a sharp tip on American kestrel, and a flat tip on Northern Crested Caracara (*Caracara Cheriway*) and Black Caracara (*Daptrius ater*). On the Falconiformes quadrate, the tip of the orbital process mostly points rostromedially, but it points rostrodorsally on Northern Crested Caracara and Black caracara quadrates. The basiorbital fossa appears to be deep and narrow on the Falconidae quadrate.

The quadratojugal cotyle of the Falconidae quadrate a rounded or oval outline with a deep fossa, dorsally close to the lateral condyle. Its dorsal margin of the quadratojugal cotyle expands medially and merges with the caudal condyle, forming an elongate and thick protrusion. Unlike that of most avian quadrate, the quadratojugal cotyle faces ventrolaterally on most falconiform quadrates. The pterygoid condyle is rostrally standing and is dorsally separated from the medial condyle with a wide and shallow gap. It appears a slender oval outline with a convex articular surface. However, the orbtiopterygoid facet, the other articulations with pterygoid on bird quadrate, is found on Northern Crested Caracara and Black Caracara quadrate only.

The mandibular process on the Falconidae quadrate has three condyles, arranging in a triangular shape in ventral view with a shallow furrow (intercondylar sulcus) between medial and lateral-caudal condyle. The medial condyle shows a rostrocaudally elongate oval shape with a convex articular surface. The lateral condyle is confluent with the caudal condyle, forming a rounded or cashew-like outline with a flat articular surface.

The Falconidae quadrate body is medially curved due to its right-angle lateral crest and curved medial crest. The caudomedial depression is wide and shallow. The subcapitular tubercle is rostroventrally located at the squamosal capitulum only on the Barred Forest falcon quadrate with a tiny oval shape. In Falconidae, three positions show the pneumatic foramen: the postcapitular foramen is caudally located at the otic process with a large opening on the Barred Forest falcon quadrate only. The postsquamosal capitulum foramen is caudally located at the squamosal capitulum on Barred Forest falcon and American kestrel quadrates. The basiorbital foramen is medially located at the quadrate body, close to the pterygoid condyle with a large opening, and it is found on most Falconidae quadrate.

**Psittaciformes:** Nestoridae, Strigopidae, Cacatuidae, and Psittacidae (Plate 41-42)

The Psittaciformes quadrate shows a distinct morphology, such as the singular condyle of the mandibular process, and thus, it is easily recognized from other avian quadrates. The otic process of the psittaciform quadrate exhibits a distinct and deep intercapitular incisure between two capitula. The squamosal capitulum is much elevated and larger than the otic capitulum. The squamosal capitulum shows an oval outline with a convex articular surface, pointing dorsally. This oval articulation elongates rostrocaudally on most parrot quadrate, but it elongates lateromedially on the Kākāpō (*Strigops habroptilus*) quadrate. The otic capitulum of the Psittaciformes quadrate generally exhibits a lateromedially wide oval shape with a flat articular surface but is not seen on the *Nestor notabilis* and *Psittacus erithacus*. On Kea (*Nestor notabilis*) quadrate, the otic capitulum appears a rounded outline with a convex articular surface. On Grey parrot (*Psittacus erithacus*) quadrate, the otic capitulum shows a rostrocaudally elongate outline. The otic capitulum of the parrot quadrate faces dorsomedially (e.g., *Nestor notabilis* and *Probosciger atermarginus*) or medially (e.g., *Strigops habroptilus* and *Psittacus erithacus*).

The orbital process significantly varies among Psittaciformes. For instance, the orbital process of the psittaciform quadrates mostly is slender and elongate with a high aspect ratio and a pointed (e.g., *Nestor notabilis*) or flat tip (e.g., *Strigops habroptilus* and *Probosciger atermarginus*), facing rostrally or rostrodrosally. On the Grey parrot quadrate, however, the orbital process is relatively shorter than other parrot quadrates with a distinct crest on the lateral surface and a downward pointed tip, facing rostroventrally. The basiorbital fossa is shallow and narrow on most Psittaciformes quadrate but is realtvielly deeper and wider on the Grey parrot quadrate.

The quadratojugal cotyle of the psittaciform quadrate is laterally standing and shows a cup-like shape with a thin margin and a deep fossa. On the Kea and Kākāpō quadrates, the caudal margin of the quadratojugal cotyle caudomedially expands, forming an elongate protrusion. The pterygoid condyle of the parrot quadrate does not rostrally protrude and it is mostly dorsally adjacent with the medial condyle. However, the pterygoid condyle on the Kākāpō quadrate is separated from the medial condyle with a small gap. The pterygoid condyle of the parrot quadrate exhibits a rounded shape with a convex articular surface.

Unlike other avian quadrates, the mandibular process on the Psittaciformes quadrate only appears one condyle (medial condyle), as shown in a rostrocaudally elongate oval outline with a prominently convex articular surface and a distinct margin with the quadrate body. On Kākāpō and Grey parrot quadrates, the medial condyle expands caudodorsally.

The psittaciform quadrate body is straight due to its right-angle lateral crest and straight medial crest. The caudomedial depression is absent or unclear on the parrot quadrate. In Psittaciformes, mostly the pneumatic foramen is caudomedially located at the quadrate body, encompassed by medial and tympanic crest. However, the pneumatic foramen is medially located at quadrate body on Kākāpō quadrate and it is restricted by the medial condyle and the orbital process.

**Passeriformes:** Acanthisittidae (Plate 42)

Different with other Passeriformes (Tyranni and Passeri), the Acanthisittidae quadrates shows its unique morphological characters, such as the shape of the orbital process and the medial condyle of the mandibular process (see below). The intercapitular incisure, a groove between the squamosal and otic capitulum, is absent on the otic process of the Acanthisittidae quadrate. The squamosal capitulum is slightly elevated than the otic capitulum, but it is as large as the otic capitulum. The squamosal capitulum exhibits a lateromedially elongate oval outline and its articular surface faces rostrodorsally. The otic capitulum exhibits a rostrocaudally elongate oval shape with a flat articular surface, facing caudodorsally.

The orbital process on the Acanthisittidae quadrate is robust with a dorsoventrally deep base, but it turns to be slender and elongate at its tip. Its tip is pointed, facing rostrally or rostromedially. A distinct crest/ridge is dorsolaterally located at the tip. The basiorbital fossa of the Acanthisittidae quadrate is wide and deep.

The quadratojugal cotyle of the Acanthisittidae quadrate shows a cup shape with a thick and oval margin, and exhibits a shallow shallow articular surface, dorsally close to the lateral condyle. The ventral of the quadratojugal cotyle expands rostroventrally, forming a flat articular surface attaching jugal bone. On the Acanthisittidae quadrate, the pterygoid condyle appears to be significantly standing rostrally, and it is dorsally separated with the medial condyle. It shows a rounded shape with a convex articular surface.

The mandibular process on the Acanthisittidae quadrate has three condyles, arranging in a L or a boomerang-like shape. The medial condyle is significantly deep and shows a rostrocaudally elongate oval shape with a prominently convex articular surface. The lateral condyle exhibits a rounded or a squared shape with a convex articular surface. The caudal condyle appears a short and thick bulge with a convex articular surface.

In Acanthisittidae, the quadrate body is relatively straight and wide with a right-angle lateral crest. The caudomedial depression of the Acanthisittidae quadrate is shallow and wide. Two pnematic foramina are caudally located at the quadrate body in Acanthisittidae: one (postcapitular foramen) is located at the middle of otic process with a clear opening. The other (caudomedial foramen) is accommodated with the caudomedial fossa on Rifleman (*Acanthisitta chloris*) quadrate only. It is encompassed by the medial and tympanic crest and shows a large opening.

**Passeriformes:** Tyranni (Suboscines, includes Calyptomenidae, Melanopareiidae, Philepittidae, Grallariidae, Pipridae, and Tyrannidae) (Plate 42-43)

The Tyranni quadrate shows a wide range of diversity on the otic process: 1) two capitula are packed with a clear intercapitular incisure between them, such as green broadbill (*Calyptomena viridis*) and Collared crescentchest (*Melanopareia torquata*) quadrate. 2) two capitula are packed without the intercapitular incisure between them, such as Thrush-like antpitta (*Myrmothera campanisona*) and Spectacled tyrant (*Hymenops perspicillatus*) quadrate. 3) two capitula are widely separated with a shallow intercapitular incisure, such as Common sunbird-asity (*Neodrepanis coruscans*) and Golden-headed manakin (*Ceratopipra erythrocephala*/*Pipra erythrocephala*) quadrate. The squamosal capitulum is higher than the otic capitulum on most Tyranni quadrate, but it is as elevated as the otic capitulum on the green broadbill quadrate. The squamosal capitulum is usually smaller than the otic capitulum, but it shows the similar size on Common sunbird-asity quadrate. The squamosal capitulum on the Suboscines quadrate generally shows a rounded or squared shape with a convex articular surface, facing dorsolaterally. It exhibits a lateromedially elongate oval shape on two suboscines (*Melanopareia torquata* and *Hymenops perspicillatus*) quadrates. The otic capitulum shows a rectangular outline with a slightly convex or a flat articular surface, and it faces dorsomedially.

In Tyranni, the orbital process is slender and elongate with a relative high aspect ratio. The tip of the orbital process on Tyranni quadrate usually appears to be flat with a distinct ridge/crest at its dorsal margin, and this ridge expands dorsolaterally but not found on the Collared crescentchest quadrate. On Collared crescentchest quadrate only, a distinct crest appears to be at the dorsal margin of the orbital process. The orbital process on the suboscine quadrate mostly faces rostrodorsally, but it points rostroventrally on common sunbird-asity quadrate. The basiorbital fossa of the suboscine quadrate is mostly shallow and narrow. It appears to be shallow and wide on the collared crescentchest quadrate, and appears to be relatively deep on the Common sunbird-asity and Thrush-like antpitta quadrates.

The quadratojugal cotyle of the Tyranni quadrate is laterally protruding and shows a saddle-like shape (the rostrodorsal and caudoventral margin is not well-developed) with a thick margin and shallow joint articulation, adjacent to the lateral condyle. The caudodorsal margin of the quadratojugal cotyle expands caudally, forming a prominent protrusion. Like all passeriform quadrates, the pterygoid condyle on the Tyranni quadrate significantly protrudes rostrally, and mostly widely separated from the medial condyle. In Tyranni, the pterygoid condyle shows a rounded outline with a convex articular surface, but it appears to an oval or a rectangular outline with a convex articular surface on the Common sunbird-asity quadrate.

The mandibular process on the suboscines quadrate has three condyles, arranging in L or boomerang-like shape with a shallow furrow (intercondylar sulcus). The medial condyle shows a rostrocaudally elongate oval shape with a convex articular surface. Although the lateral condyle is confluent with the caudal condyle on the suboscines quadrate, it exhibits a square (e.g., *Calyptomena viridis*) or an oval shape (e.g., *Neodrepanis coruscans*, *Myrmothera campanisona*, and *Ceratopipra erythrocephala*) on some suboscines quadrates. The lateral and the caudal condyle forms a rectangular shape with a flat articular surface on the collared crescentchest and the spectacled tyrant quadrates. The caudal condyle shows a short and thick bulge with a flat articular surface.

The Tyranni quadrate body is mostly slender and strongly curved due to a curved medial crest, such as green broadbill and Golden-headed manakin quadrates. The quadrate body on the Collared crescentchest quadrate is relatively wider than other Tyranni quadrates in this study. The caudomedial depression is generally shallow but it is significantly deep on the Collared crescentchest and Spectacled tyrant quadrate. The subcapitular tubercle appears a liner ridge ventrolaterally located at the squamosal capitulum on the Golden-headed manakin quadrate only. In Tyranni, the pnematic foramen is caudally located at the otic process, as shown in a clear opening (postcapitular foramen) at the midpoint of the otic process on all quadrate and two tiny opening at two capitula on the Spectacled tyrant quadrate only.

**Passeriformes:** Passeri (Oscines or songbirds, includes Menuridae, Ptilonorhynchidae, Maluridae, Campephagidae, Falcunculidae, and Corvidae, Callaeidae, Picathartidae/Eupetidae, Paridae, Hirundinidae, Leiothrichidae, Bombycillidae, Regulidae, Ploceidae, Fringillidae, and Emberizidae) (Plate 43-46)

The otic process does not show a clear intercapitular incisure between two capitula on most Passeri quadrates, but it shows a clear incisure on several Passeri quadrates (e.g, *Malurus melanocephalus*, *Edolisoma tenuirostre*, *Pyrrhocorax pyrrhocorax*, *Callaeas cinereus*, *Picathartes gymnocephalus*, *Hirundo rustica*, *Plocepasser mahali*, *Chlorodrepanis virens*, and *Emberiza calandra*). Two capitula are close to each other only on the Common cicadabird (*Edolisoma tenuirostre*) quadrate. The squamosal capitulum is usually much elevated than the otic capitulum but it is much smaller or eual to the otic capitulum. The squamosal capitulum of most Passeri quadrates shows a lateromedially elongate oval outline with a convex articular surface, and it faces rostrodorsally or dorsally. On some oscines quadrates (e.g., *Falcunculus frontatus*, *Picathartes gymnocephalus*, *Parus major*, and *Bombycilla garrulus*), the squamosal capitulum shows a rounded or squared outline. The otic capitulum usually has a lateromedially oval or rectangular outline with a flat articular surface, facing cuadodorsally or dorsally. On some oscines quadrates (e.g., *Picathartes gymnocephalus*, *Regulus ignicapillus*, *Chlorodrepanis virens*, and *Emberiza calandra*), the otic capitulum shows a rounded outline.

The orbital process of most Passeri quadrate is rostrally protruding and robust, especially on Eastern shriketit (*Falcunculus frontatus*), Red-billed chough (*Pyrrhocorax pyrrhocorax*), Hawaiʻi ʻamakihi (*Chlorodrepanis virens*), and Corn bunting (*Emberiza calandra*) quadrates. On some oscines quadrate, the orbital process is still rostrally protruding but slender, such as South Island kōkako (*Callaeas cinereus*), White-necked rockfowl (*Picathartes gymnocephalus*), and Great tit (*Parus major*) quadrates. The orbital process is relatively short on some Passeri quadrate, such as Red-backed fairywren (*Malurus melanocephalus*), Common cicadabird, Barn swallow (*Hirundo rustica*), and Common firecrest (*Regulus ignicapillus*). In Passeri, the tip of the orbital process is usually flat and turns to be slant or be horizontal, facing rostrally or dorsorostrally. On Hawaiʻi ʻamakihi quadrate, its tip significantly turns to be vertical, and on some Passeri quadrates (Red-backed fairywren, Great tit, and *Corn bunting*), the tip of the orbital process appears to be pointed. On Red-billed chough quadrate only, a distinct ridge is dorsally located at the tip of the orbital process. The basiorbital fossa of the Passeri quadrate is usually shallow and narrow, but it is absent on some oscines quadrates, such as Eastern shriketit, White-necked rockfowl, and Bohemian waxwing (*Bombycilla garrulus*). The basiorbital fossa is relatively deep on Superb lyrebird (*Menura novaehollandiae*) and Common cicadabird quadrate.

The quadratojugal cotyle significantly protrudes laterally on most Passeri quadrates and shows a cup-like shape with a deep articular surface, adjacent to the lateral condyle. However, this joint articulation is shallow on some Passeri quadrates, such as Eastern shriketit, Barn swallow, Bohemian waxwing, Common firecrest, and White-browed sparrow-weaver (*Plocepasser mahali*). In Passeri, the ventral or dorsoventral margin of the quadratojugal cotyle expands ventrally, forming an articular surface for contacting jugal. Similar as that of other passeriform quadrates, the pterygoid condyle of the Passeri quadrate significantly protrudes rostrally and widely separated from the medial condyle. It has a rounded or oval outline with a convex articular surface.

The mandibular process on oscines quadrate has three condyles, arranging in L shape, but it ranges lateromedially with an underdeveloped caudal condyle on some oscines quadrate, such as Satin bowerbird (*Ptilonorhynchus violaceus*), Eastern shriketit, and Hawaiʻi ʻamakihi. The intercondylar sulcus is relatively deep located at the centre of the mandibular process, between medial condyle and lateral-caudal condyle. The medial condyle shows a rounded or a rostrocaudally elongate oval outline with a prominent convex articular surface. On some oscines quadrate, the medial condyle is significantly deep, such as Superb lyrebird, Red-billed leiothrix (*Leiothrix lutea*), Bohemian waxwing, Common firecrest, White-browed sparrow-weaver, Hawaiʻi ʻamakihi, and Corn bunting. The lateral condyle is usually confluent with the caudal condyle and forms an oval or rectangular outline with a flat articular surface on the Passeri quadrate. However, the lateral condyle still exhibits a distinct shape on several oscines quadrate, shown in an oval outline with a convex articular surface (e.g., *Ptilonorhynchus violaceus, Malurus melanocephalus, Actinodura cyanouroptera, Leiothrix lutea, Plocepasser mahali*) or a rounded outline with a convex articular surface (e.g., *Falcunculus frontatus*, *Callaeas cinereus*, *Parus major*, *Emberiza calandra*). On most Passeri quadrate, the caudal condyle appears a short and thick bulge with a flat articular surface, while it is not well developed on a few oscines quadrate (e.g., *Ptilonorhynchus violaceus*, *Falcunculus frontatus*, *Hirundo rustica*, and *Chlorodrepanis virens*).

The Passeri quadrate body is mostly wide and relatively straight due to a right-angle lateral crest and relatively straight medial crest, while on some oscines quadrates, their quadrate body is laterally curved due to strongly curved lateral crests, such as Red-backed fairywren, Bohemian waxwing, Common firecrest, White-browed sparrow-weaver, and Hawaiʻi ʻamakihi. The caudomedial depression is wide and deep on most Passeri quadrates. Although the subcapitular tubercle is absent on the Passeri quadrates, the lateral crest protrudes dorsolaterally, forming a clear ridge or protrusion on most Passero quadrates. In most Passeri, only one pneumatic foramen (rostromedial foramen) is medially located at the quadrate body with variant size in different species, encompassed by the orbital process and the medial crest. The pneumatic foramen or pneumatic fossa is also caudally located at the otic process on several Passeri quadrate, such as Red-backed fairywren, Great tit, Bohemian waxwing, Common firecrest, and Corn bunting. Finally, the caudomedial pneumatic foramen/caudomedial fossa appears on caudomedial depression on some Passeri quadrates (e.g., *Malurus melanocephalus*, *Edolisoma tenuirostre*, *Falcunculus frontatus*, *Hirundo rustica*, *Actinodura cyanouroptera*, *Leiothrix lutea*, *Bombycilla garrulus*, *Chlorodrepanis virens*, and *Emberiza calandra*).
